# Chronic opioid-associated immune dysregulation among people living with HIV

**DOI:** 10.64898/2026.08.31.748399

**Authors:** Brent W. Bever, Michelle Underwood, Byung Park, Susan Pereira Ribeiro, Ryan R. Cook, Lynn Kunkel, P. Todd Korthuis, Christina Lancioni

## Abstract

**Objectives:** Persistent immune dysregulation contributes to chronic disease among people living with HIV (PWH), even after viral suppression with antiretroviral therapy (ART). Although chronic opioid exposure is associated with adverse clinical outcomes, its impact on immune homeostasis during ART remains incompletely understood. We investigated whether opioid use disorder (OUD) is associated with persistent systemic and cellular immune dysregulation despite ART-mediated reductions in HIV viral load (VL).

**Methods:** Peripheral blood was collected longitudinally from PWH with OUD (PWH/OUD+) and detectable HIV VL during 6 months of optimized ART (months 0, 3, and 6). A reference cohort of PWH without OUD (PWH/OUD−) and suppressed HIV VL provided a single blood sample. Immune profiling included plasma inflammatory biomarkers, multiplex cytokine analyses, spectral flow cytometry, and assessment of monocyte cytokine responses following lipopolysaccharide (LPS) stimulation. Mixed-effects models adjusted for HIV VL and VL-stratified analyses were performed.

**Results:** PWH/OUD+ exhibited persistent immune dysregulation despite reductions in HIV VL. Plasma sCD163, sCD14, fractalkine, and I-TAC remained elevated, whereas TGF-β1 was reduced. OUD was associated with expansion of CD16⁺ monocytes and altered expression of CCR2, CD38, and CD11b. CD4⁺ and CD8⁺ T cells, NK cells, and B cells also exhibited persistent alterations in markers of activation, metabolism, and trafficking. Monocytes from PWH/OUD+ displayed attenuated cytokine responses following LPS stimulation.

**Conclusions:** OUD is associated with persistent systemic and cellular immune dysfunction in PWH despite ART-mediated viral suppression, supporting opioid exposure as an independent contributor to chronic immune dysregulation that may promote inflammation, immune dysfunction, and long-term HIV-associated comorbidities.

## Introduction

Over 9 million adolescents and adults living in the US misuse opioids (either prescription or illicit) annually, with over eighty-thousand Americans dying from opioid overdose in 2022 alone (Volkow & Dye, 2025). Opioid exposure is even more common among people living with HIV (PWH), with roughly 15% of PWH experiencing an opioid use disorder (OUD) in their lifetime (Chinazo O. Cunningham, 2018). Chronic opioid exposure (COE) is associated with a significantly higher risk of disease and death from cardiovascular events, cancer, and infections (Hser et al., 2017), as well as an increased risk of dementia (Abrego-Guandique et al., 2026; Dublin et al., 2015). Among PWH, OUD is associated with HIV disease progression even among individuals with undetectable HIV viral load (VL) while on antiretroviral therapy (ART) (Azzoni et al., 2020; Trunfio et al., 2023). Regardless of opioid-exposure, PWH on ART remain at risk for chronic systemic inflammation and immune dysregulation that drive non-communicable diseases (Anzinger et al., 2014; Wilson et al., 2014; Martin et al., 2013; Kamat et al., 2012; Dominick et al., 2020) including HIV associated neurocognitive disorders (HAND). HAND encompasses a range of cognitive, motor, and behavioral problems (Robertson et al., 2009), and has been associated with increased central nervous system neuroinflammation (Hauser et al., 2007; Kurt F. Hauser et al., 2005). Therefore, any exposures that further exacerbate immune dysregulation among PWH, such as COE, are likely to contribute to long-term morbidities such as HAND, that cannot be addressed by optimization of ART alone.

Immune cells such as monocytes play an important role in patrolling the body for microbial pathogens and directing the immune response to infection and inflammation (Yáñez et al., 2017). Once activated, monocytes migrate to sites of inflammation and pass through the endothelium into inflamed tissue, where they facilitate the immune response by producing pro- or anti-inflammatory cytokines and chemokines, and serve as antigen-presenting cells for components of the adaptive immune system. Monocytes that express the Fc-receptor CD16, termed intermediate (CD14^++^CD16^+^) and non-classical (CD14^+^CD16^++^) monocytes, have been reported to cross the blood-brain barrier and to influence neuroinflammation. In addition, CCR2+ monocytes have been shown to play a role in Alzheimer’s disease by directing amyloid clearance, and CCR2 impairment in mice resulted in worse Alzheimer’s disease (Stahr & Galkina, 2022). In the context of HIV infection, CD16+ monocytes have been shown to be a reservoir for HIV (Veenhuis et al., 2023), where they cross the blood brain barrier and differentiate into pro-inflammatory macrophages that contribute to HAND (Burdo et al., 2013; Gras & Kaul, 2010; Valcour et al., 2011). Our previous work demonstrated that PWH and OUD (PWH/OUD+) with controlled VL have increased biomarkers of innate immune activation, expansion of CD16+ monocytes, and altered cytokine responses from stimulated peripheral blood mononuclear cells (PBMC) (Underwood et al., 2020). Opioids have been shown to inhibit cell growth (Vassou et al., 2008), promote apoptosis (Nair et al., 1997), and suppress phagocytosis (Ninkovi & Roy, 2012) in macrophages. However, the impact of COE on innate and adaptive immunity among PWH as they receive optimized ART and reductions in VL has not been reported, and there are limited data that provide insight into the specific cellular subsets most disrupted by opioid exposure. Here, we hypothesized that opioid exposure would be associated with sustained changes in innate immune cell phenotype and inflammatory responses among PWH, regardless of control of HIV viremia.

## Methods

### Participant recruitment and ethics statement

PWH/OUD+ were recruited to participate in an immunology sub-study through the "Comparing Treatments for HIV-Infected Opioid Users in an Integrated Care Effectiveness Study (CHOICES) Scale-Up Clinical Trials Network-067" Phase II clinical trial (Korthuis et al., 2022). The overall objectives of the CTN-067 trial were to optimize control of HIV infection through treatment with ART medications combined with treatment of underlying OUD with either extended-release naltrexone (XR-NTX) or treatment-as-usual (TAU) with medications including buprenorphine, methadone, and/or naloxone. Recruitment occurred at 7 comprehensive HIV-care clinics across the USA from 2017 to 2019 (clinicaltrails.gov NCT03275350). Adult participants confirmed to be living with HIV and meeting DSM-5 criteria for moderate or severe OUD and with VL ≥ 200 copies/ml at screening (PWH/OUD+, “CTN-067 participants”), were eligible. Individuals were excluded if they had a serious medical, psychiatric, or substance use disorder that, in the opinion of the study physician, would make study participation hazardous, compromise study findings, or prevent study completion. Participants were randomized to treatment with XR-NTX or TAU; ART regimens were initiated and/or optimized at provider discretion. All participants were followed for six months. All participants provided written informed consent for study participation, including participation in the immunology sub-study. The Advarra Institutional Review Board (IRB00000971) reviewed and approved the study and served as a single IRB for the study, with participating sites deferring to its regulatory role (Korthuis et al., 2022). A reference population of forty-seven PWH, aged 18-65 years old, and without any history or current substance dependency ("control group"; PWH/OUD-) were recruited from the OHSU HIV-comprehensive care clinic. Exclusion criteria for the control group included: pregnancy, active or history of substance abuse, current acute illness, and use of prescription opioids. All participants provided written informed consent for study participation, and ethical approval was provided through the OHSU Institutional Review Board.

### PBMC and Plasma Processing and Storage

Up to 32mL of peripheral blood was collected into CPT Vacutainer tubes (BD Biosciences, Franklin Lakes, New Jersey, USA) at three time points (M0/M3/M6) at each recruitment site. Participant clinical and sociodemographic data (eg, randomization arm; monthly urine toxicology testing; dates of XR-NTX or TAU treatment, age, biologic sex, substance use history, use of ART, smoking history, and Hepatitis B/C status) were also collected.

CPT Vacutainer tubes were centrifuged within 2 hours of collection and shipped at room temperature to our central laboratory, where PBMC and plasma isolation was completed following a standardized operating procedure (SOP) within 24 hrs of collection. PBMCs were cryopreserved at -150°C in 10% DMSO (Sigma-Aldrich, St Louis, Missouri, USA), Fetal bovine serum, with 0.1% Gentamicin. Plasma was stored at -80°C. The control cohort had plasma and PBMC collected, processed, and stored at a single timepoint, following the same protocol as CTN-067 participants (Underwood et al., 2020; 2022).

### Flow Cytometry

Cryopreserved PBMCs were thawed and transferred to a 96-well plate. Following antibody titration to maximize separation index, cells were stained with the markers listed in **sup table 1** using Brilliant Stain Buffer Plus (BD) buffer, fixed, and acquired using a Cytek Aurora Flow cytometer (Cytek Biosciences). SpectroFlo® (3.3.0) was used to perform spectral unmixing analysis using single-stained PBMC. The PeacoQC (v1.5.0) plugin was used in FlowJo (BD Biosciences) for quality control, and FlowJo was then used to identify monocytes, T cells, CD19+ B cells, and CD56+ populations, as shown in the **sup figs 1-4.** Uniform Manifold Approximation and Projection (UMAP) (v4.1.1) plugin was used in FlowJo to examine phenotypic changes over time within each cellular population.

**Table 1.**
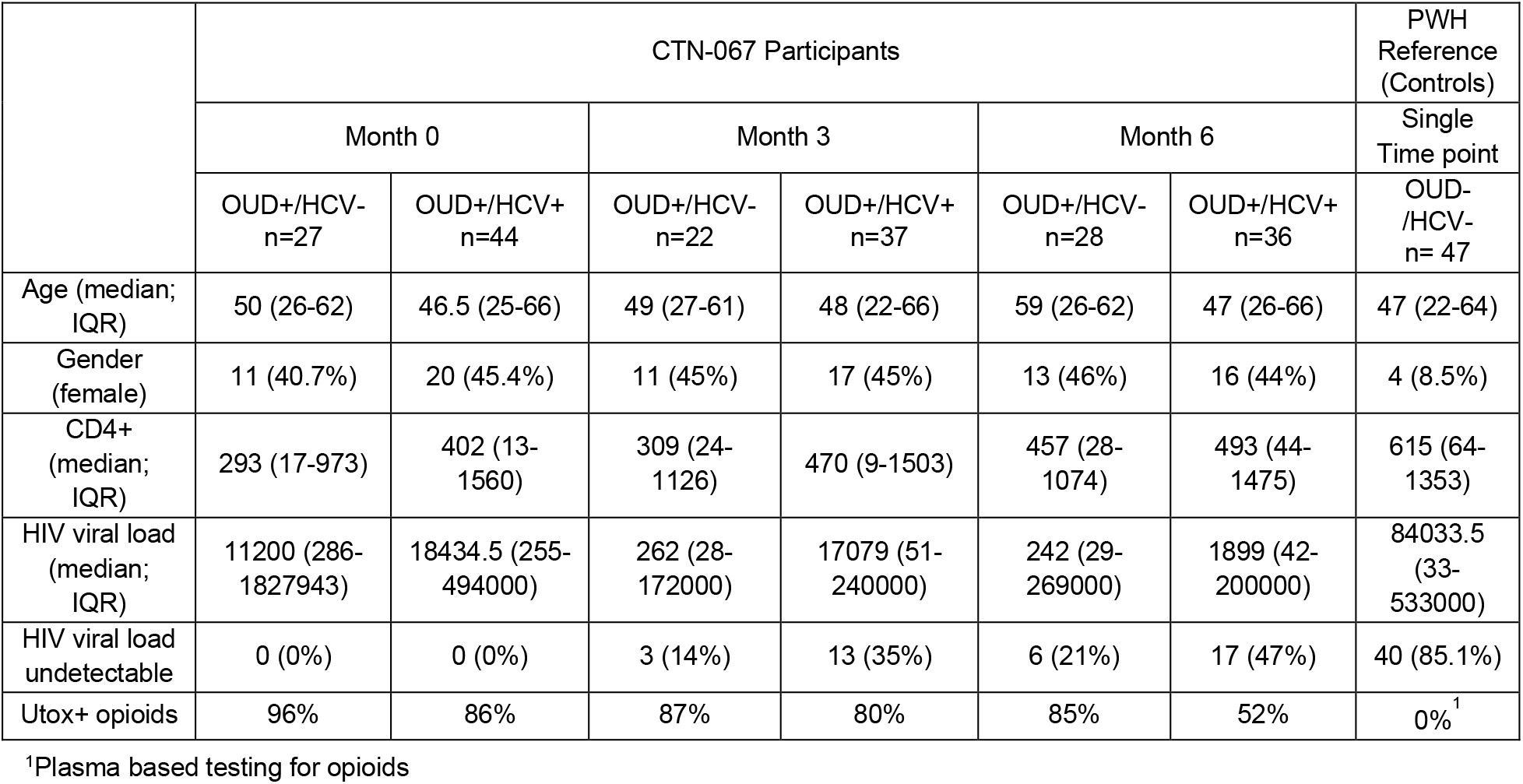
Demographic and clinical characteristics of study participants.

|  | CTN-067 Participants |  |  |  |  |  | PWH Reference (Controls) |
| --- | --- | --- | --- | --- | --- | --- | --- |
|  | Month 0 |  | Month 3 |  | Month 6 |  | Single Time point |
|  | OUD+/HCV-<br>n=27 | OUD+/HCV+<br>n=44 | OUD+/HCV-<br>n=22 | OUD+/HCV+<br>n=37 | OUD+/HCV-<br>n=28 | OUD+/HCV+<br>n=36 | OUD-/HCV-<br>n= 47 |
| Age (median; IQR) | 50 (26-62) | 46.5 (25-66) | 49 (27-61) | 48 (22-66) | 59 (26-62) | 47 (26-66) | 47 (22-64) |
| Gender (female) | 11 (40.7%) | 20 (45.4%) | 11 (45%) | 17 (45%) | 13 (46%) | 16 (44%) | 4 (8.5%) |
| CD4+ (median; IQR) | 293 (17-973) | 402 (13-1560) | 309 (24-1126) | 470 (9-1503) | 457 (28-1074) | 493 (44-1475) | 615 (64-1353) |
| HIV viral load (median; IQR) | 11200 (286-1827943) | 18434.5 (255-494000) | 262 (28-172000) | 17079 (51-240000) | 242 (29-269000) | 1899 (42-200000) | 84033.5 (33-533000) |
| HIV viral load undetectable | 0 (0%) | 0 (0%) | 3 (14%) | 13 (35%) | 6 (21%) | 17 (47%) | 40 (85.1%) |
| Utox+ opioids | 96% | 86% | 87% | 80% | 85% | 52% | 0%<sup>1</sup> |

<sup>1</sup>Plasma based testing for opioids

### CD14+ Isolation and Stimulation

CD14+ isolations and stimulations were performed on freshly isolated (not cryopreserved) PBMC samples. CD14+ positive selection was performed (Miltenyi Biotec, Auburn, CA) following manufacturer’s instructions, with purity of resulting populations demonstrated to be >90% CD3-CD19-CD14+ monocytes by flow cytometry. CD14+ monocytes were plated on a 48-well flat bottom non-ULA tissue culture plate (10^6^ cell/well). Cells were left unstimulated (resting) or stimulated with Lipopolysaccharide (LPS, ultrapure; Invivogen; 100 ng/mL) in duplicate. After 18 hours of incubation (37°C/5% Co2), cell culture supernatants were harvested and stored at -80°C.

### Plasma ELISA

Plasma cytokine and protein quantification were performed using commercially available kits, and manufacturer’s instructions were followed: soluble CD14 (sCD14; R&D Systems, Minneapolis, Minnesota, USA, Cat#QK383), Intestinal Fatty Acid-Binding Protein (I-FABP; R &D Systems, Cat# DFBP20), lipopolysaccharide-binding protein (LBP; R&D Systems, Cat# DY870-05), IL-37 (R&D Systems, Cat# DY1975-05), Occludin (NOVUS Biologicals, Centennial, CO, Cat# NBP2-80305), and soluble CD163 (sCD163; Invitrogen, Thermo-Fisher Scientific, Cat# CHCD163). Experimental duplicates were performed for each sample.

### CD14+ Culture Supernatant and Plasma Cytokine Quantification

U-PLEX assay (MESO Scale Multi-Array technology; Meso Scale Discovery; Rockville, MD, USA) was used to quantify plasma: Fraktalkine, GMCSF, ITAC, IFN-β, IFN-α, IFN-γ, IL-10, IL-12p70, IL-15, IL-17A, IL-18, IL-1β, IL-2, IL-21, IL-22, IL23, IL-27, IL-29, IL-33, IL-4, IL-6, IL-7, IL-8, IL-9, IP-10, MCP-3, MIP-1α, MIP-3α, SDF-1α, TGF-β1, TGF-β2, TGF-β3, and TNF-α. Following manufacturer’s instructions, duplicate standard curves and plasma samples (25µL plasma) were included from each study timepoint. Electrochemiluminescence was detected using MESO QuickPlex SQ qwo (Meso Scale Discovery Rockville, MD, United States). Using DISCOVERY WORKBENCH v4.0 software (Meso Scale Discovery, Rockville, MD, United States), a standard curve was generated, and mean values for each sample duplicate calculated in pg/mL.

### Statistical Analysis

For comparison of demographic and clinical characteristics (**Table 1**), the median was calculated for age, CD4+ T cell count, and HIV VL, and the IQR reported. The frequency of female biologic sex, undetectable HIV VL, and positive urine toxicology test for opioids, was also reported. Statistical analyses were performed using SAS 9.4 (SAS Institute Inc., Cary, NC, USA). For longitudinal and cross-sectional comparisons across time points (M0, M3, M6) and controls, a mixed-effects repeated-measures ANOVA model with compound symmetry or first-order autoregressive, AR(1), covariance structure was used to account for within-subject correlation. Bayesian Information Criteria (BIC) was used for selecting covariance structure. HIV VL was included as a covariate in all models using a log₁₀ transformation (log₁₀[viral load + 1]) to normalize distribution and control for the effects of viral replication. Post hoc multiple comparisons were performed using Tukey-Kramer correction. For analyses stratified by VL, GraphPad Prism (V.10.6.1) was used. Participants were categorized into high VL (HVL; >500 copies/mL) and low viral VL (LVL; ≤375 copies/mL) groups based on clinical thresholds for viral suppression. Comparisons between HVL, LVL, and control groups were performed using a non-parametric Mann-Whitney test. The level of significance alpha = 0.05 was set for each statistical analysis performed.

## Results

### Characteristics of Study Participants

Seventy-one PWH/OUD+ enrolled in the CTN067 CHOICES six-month clinical trial (Korthuis et al., 2022), and a population of forty-seven PWH/OUD- on stable ART, who served as a control, or reference population, were included. Co-infection with Hepatitis C (HEP) was common among PWH/OUD+ individuals (**Table 1**). Of 73 CTN067 participants, 57.6% were assigned to TAU and 42.3% were assigned to XR-NTX. Successful reconstitution of CD4+ T cells and reductions in HIV VL were observed by month six (M6), regardless of OUD-treatment assignment (**Table 1**). The majority of trial participants demonstrated on-going exposure to non-prescription opioids based on monthly urine-based testing, with a median duration of OUD treatment of only 2 (TAU) or fewer (XR-NTX) months. The PWH reference group demonstrated low to undetectable HIV VL, normal CD4+ T cell counts, no HCV co-infection, and no exposure to prescription or non-prescription opioids based on plasma-based testing and self-report.

### PWH/OUD maintain altered biomarkers of innate immune activation despite reductions in viral load

Chronic innate immune activation is associated with poor clinical outcomes among PWH, including HAND, and may be partially driven by HIV-associated damage to the intestinal mucosa (Nasi et al., 2020; Nyström et al., 2015; Ramendra et al., 2019; Balena et al., 2020; Franchi et al., 2019; Gill & Kolson, 2014; Tedaldi et al., 2015). As opioids may also damage the intestinal mucosa (Chopyk & Grakoui, 2020; Wang & Roy, 2017) and are associated with poor cognitive outcomes (Abrego-Guandique et al., 2026) here we quantified plasma biomarkers of innate immune activation (sCD14; sCD163), intestinal permeability (I-FABP; LBP) and IL37, an anti-inflammatory cytokine associated with inflammatory bowel disease (Porter et al., 2018; Van Der Gracht et al., 2016). In a cross-sectional analysis, after adjustment for HIV VL, sCD14 levels were significantly higher in PWH/OUD+ at M0 (adj.p = 0.001), M3 (adj.p = 0.0001), and M6 (adj.p = 0.0002) than in controls, as shown in **Figure 1a**. Soluble CD163 levels were also significantly increased in PWH/OUD+ compared with controls at M0 (adj. p = 0.0001), M3 (adj. p = 0.0001), and M6 (adj. p = 0.0001) (**Fig. 1a**). There were no significant differences in plasma levels of I-FABP, IL-37, or LBP between PWH/OUD+ and the control group (**Sup table 2**). In a longitudinal analysis adjusted for HIV VL among PWH/OUD+, there was a significant decrease in sCD163 expression from M0 to M6 (**Sup table 2**). However, there were no significant differences in plasma levels of sCD14, LBP, IL-37, and I-FABP over 6-months among individuals with PWH/OUD+ (**Sup table 2**). Our results suggest that PWH/OUD+ experience sustained innate immune activation despite successful reductions in HIV VL, when compared to PWH without opioid exposure.

**Figure 1.**
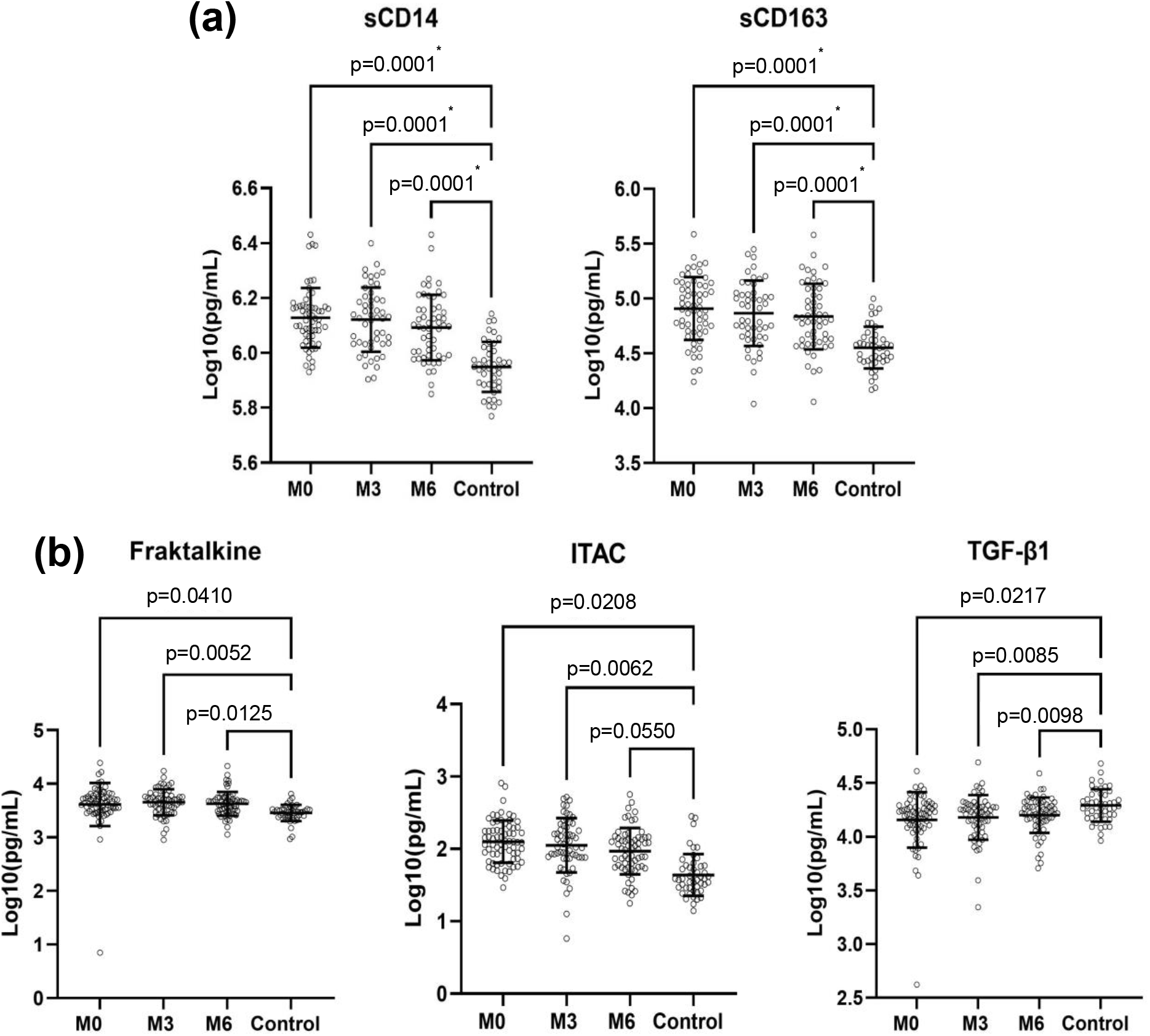
Alterations in peripheral blood plasma biomarkers and cytokines are associated with dysregulated innate immunity among PWH/OUD+. **(a)** Plasma levels of sCD14 and sCD163 were measured by ELISA from PWH/OUD+ at M0 (n=59), M3 (sCD14 n=50, sCD163 = 49), and M6 (n=54), and control participants (n=44). Shown are Log10 transformed mean results (pg/ml) with SD. Statistical analyses were performed using repeated-measures ANOVA with compound symmetry, including HIV viral load as a covariate, and Tukey–Kramer correction for multiple comparisons. **(b)** Plasma cytokine levels of Fractalkine (M0 n=71, M3 n=60, M6 n=71, control n=47), ITAC (M0 n=70, M3 n=60, M6 n=64, control n=47), and TGF-β1 (M0 n=70, M3 n=60, M3 n=60, M6 n=63, control n=47) were measured by MesoScale multiplex assay. Shown are Log10 transformed mean results (pg/ml) with SD. Statistical analyses were performed using a general linear mixed-effects model. A first-order autoregressive [AR(1)] covariance structure was applied to account for within-subject correlation, with HIV viral load included as a covariate. P values adjusted for multiple comparisons are denoted with *.

### OUD is associated with alterations in peripheral blood cytokines

Prior research suggests that opioid exposure impairs cellular chemotaxis (Eisenstein, 2019; Szabo et al., 2002; Zhang et al., 2003). Here, we compared plasma levels of 33 cytokines and chemokines that modulate cellular immune functions, including chemotaxis, among PWH/OUD+ compared with controls. In a cross-sectional analysis with HIV VL included as a covariate, fractalkine levels were significantly higher at M0 (p = 0.0410), M3 (p = 0.0052), and M6 (p = 0.0125) among PWH/OUD+ (**Figure 1b)**. Among PWH/OUD+, plasma levels of Interferon-inducible T-cell alpha chemoattractant (ITAC) were significantly increased at M0 (p = 0.0208), M3 (p = 0.0062), and M6 (p = 0.0550). Conversely, plasma levels of transforming growth factor beta-1 (TGF-β1) were significantly reduced at all three time points [M0 (p = 0.0127), M3 (p = 0.0085), and M6 (p = 0.0098)], when compared to controls. IL-8 was significantly increased at M0 and M6 when compared to the control group (**Sup table 3**). It is noted that these differences were no longer significant after applying correction for multiple comparisons, potentially due to limited sample size (**Sup table 3**). Importantly, there were no significant changes over time in plasma cytokine levels among individuals with PWH/OUD+, despite overall reductions in HIV VL (**Sup table 3**).

### OUD is associated with altered monocyte phenotypes among PWH

Previous research identified an association between OUD and expansion of CD16+ monocytes (Underwood et al., 2020), as well as alterations in the phenotypes and functions of human monocyte-derived macrophages (Barbaro et al., 2021; Murphy et al., 2019; Rao et al., 2014; Malik & Agrewala, 2025). Here, we performed detailed phenotyping of peripheral monocytes using spectral flow cytometry in a subset of PWH/OUD+ and controls, and analyzed our results using both unsupervised (**Figure 2a**) and supervised approaches (**Figure 2b-d; Sup table 4).** Although our analysis of cellular phenotypes considered HIV VL as a covariable (**Figures 2**), to provide further rigor to our approach we also performed a stratified analysis based on samples collected during periods of high VL (HVL; >500 copies/mL) versus low VL (LVL; <500 copies/mL) within the PWH/OUD+ cohort (**Figure 3; Sup table 5**).

**Figure 2.**
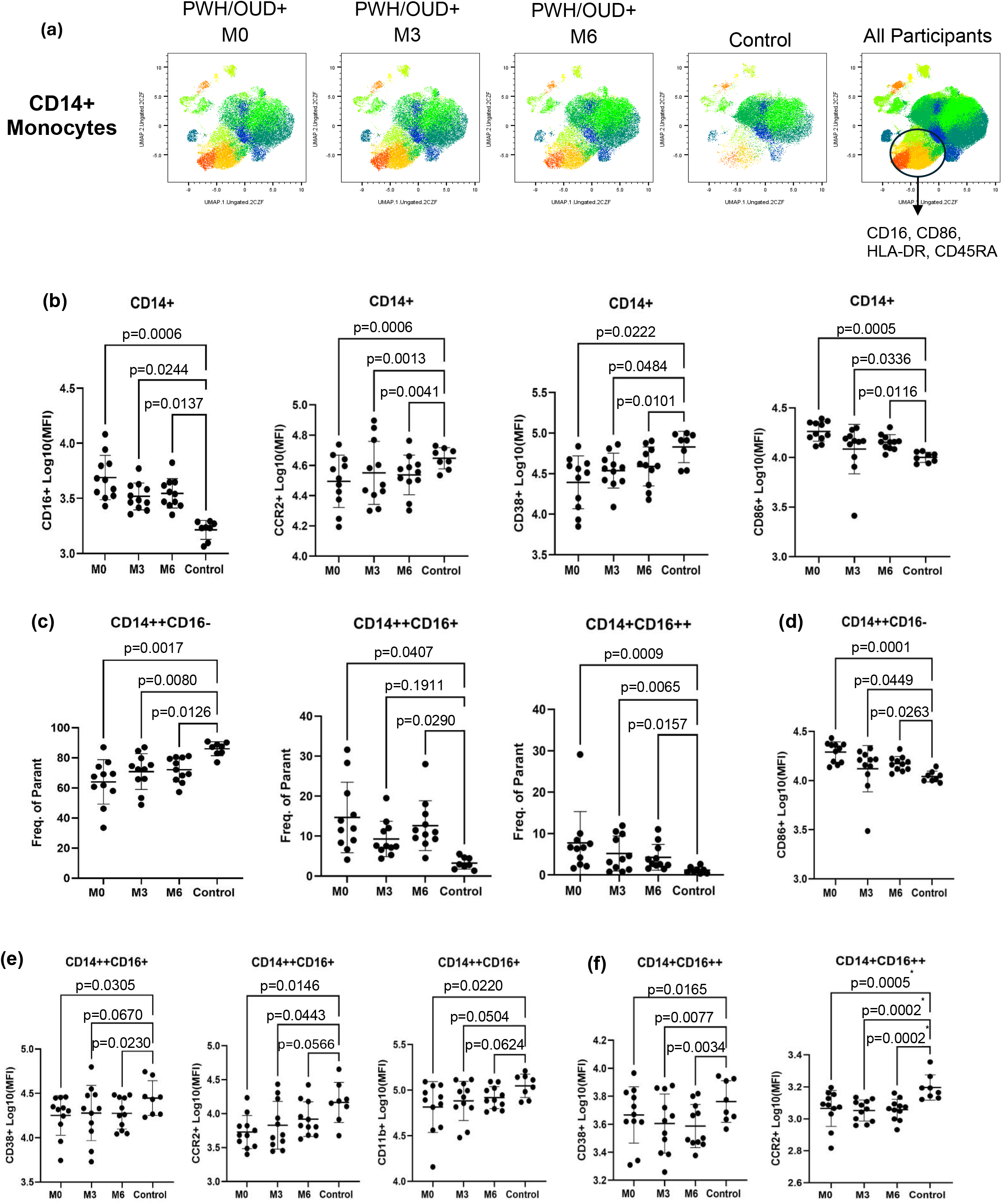
PWH/OUD+ exhibit expansion of CD16+ monocyte subsets, as well as altered expression of activation and migration-associated markers. **(a)** UMAP visualization of CD14⁺ monocyte phenotypes over time (M0, M3, M6) among PWH/OUD+ (n=11) compared to controls (n=8) with markers CD16, CD86, HLA-DR and CD45RA annotated. (b) Mean fluorescence intensity (MFI) of CD16, CCR2, CD38, and CD86 on total CD14⁺ monocytes compared to the controls. Gating was performed on total live CD3⁻CD19⁻CD56⁻CD14⁺ cells within the high-forward-scatter (FSC) cell (large cell gate populations) (sup fig 1). Shown are Log10 transformed mean results (MFI) with SD. **(c)** Shown are frequencies of classical, intermediate, and nonclassical monocytes over time when compared to controls. Frequencies were calculated from total live CD3⁻CD19⁻CD56⁻ cells based on CD14 and CD16 expression. Shown are mean percentage with SD. **(d)** MFI of CD86 on classical CD14⁺⁺CD16⁻ monocytes across all three timepoints compared to controls. Shown are Log10 transformed mean results (MFI) with SD. **(e)** MFI of markers CD38, CCR2 and CD11b on intermediate CD14⁺⁺CD16⁺ monocytes across all three timepoints when compared to controls. Shown are Log10 transformed mean results (MFI) with SD. **(f)** MFI of CD38 and CCR2 on non-classical CD14⁺CD16⁺⁺ monocytes across M0, M3, and M6 compared to controls. Shown are Log10 transformed mean results (MFI) with SD. Cross-sectional analyses across were performed using mixed-effects repeated-measures ANOVA with compound symmetry to account for within-subject correlation. HIV viral load was included as a covariate; Tukey correction was applied for multiple comparisons. P values adjusted for multiple comparisons are denoted with *.

**Figure 3.**
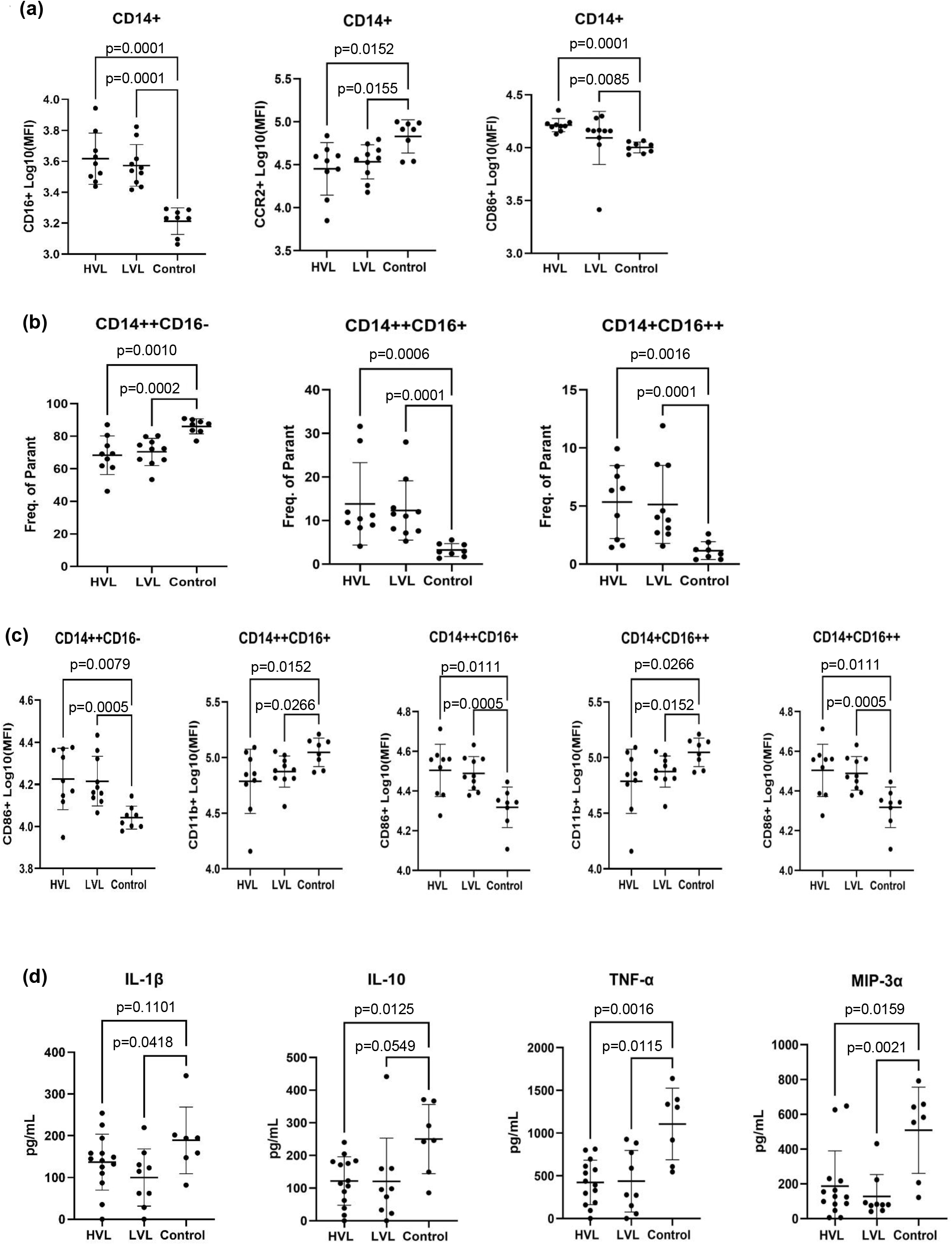
Stratification by HIV Viral load supports association between OUD and altered monocyte subsets and functional responses. **(a)** Shown are MFI of CD16, CD86 and CCR2 within CD14⁺ monocytes in collected during periods of high viral load (HVL; >477 copies/mL, Average 96367 copies/mL) and low viral load (LVL; <326 copies/mL, Average 47 copies/mL ), compared to controls. Shown are log10 transformed MFI with SD. **(b)** Frequencies of Classical CD14⁺⁺CD16⁻ , Intermediate CD14⁺⁺CD16⁺, and Non-classical CD14⁺CD16⁺⁺ monocytes in HVL and LVL groups compared to controls. Shown are mean frequencies with SD. **(c)** MFI of CD86 on CD14⁺⁺CD16⁻ monocytes, CD11b, and CD86, on CD14⁺⁺CD16⁺ and CD14⁺CD16⁺⁺ monocytes in HVL and LVL groups compared to control. Shown are log10 transformed MFI with SD.For A-C, cross-sectional analyses were performed using mixed-effects repeated-measures ANOVA with compound symmetry to account for within-subject correlation. HIV viral load (log₁₀[viral load + 1]) was included as a covariate; Tukey correction was applied for multiple comparisons. **(d)** CD14+ monocytes were isolated from PWH/OUD+ and stimulated with LPS, and cytokines were measured by Mesoscale array. Shown are mean pg/mL of IL-1β, IL-10, TNF-α, and MIP-3α production in HVL (n = 14, HVL; >599 copies/mL, Average 92347 copies/mL) and LVL (n = 9, LVL; <375 copies/mL, Average 62 copies/mL ) groups compared to control (n = 7). For statistical analysis, samples were considered independent and stratification groups based on viral load at the time of blood draw among CTN-067 participants. Comparisons performed using the nonparametric Mann-Whitney test. P values adjusted for multiple comparisons are denoted with *.

Among all CD14⁺ monocytes, CD16 expression was increased at M0 (p = 0.0006), M3 (p = 0.0244), and M6 (p = 0.0137) among PWH/OUD+ as compared to controls (**Fig. 2b**). CD16 expression was also elevated in CD14+ monocytes when stratifying by HIV VL among PWH/OUD+ when compared to controls, at periods of both high (p=0.0004) and low (p=0.0001) HIV VL (**Fig 3a**). CCR2 was decreased at M0 (p=0.0006), M3 (p=0.0013), M6 (p=0.0041) as compared to controls (**Fig 2b**). When stratifying by HIV VL among PWH/OUD+, CCR2 expression was reduced on monocytes at periods of both high (p=0.0152) and low (p=0.0155) HIV VL, when compared to controls (**Fig 3a**). CD14+ monocytes also showed decreased CD38+ expression at M0 (p=0.0222), M3 (p=0.0484), and M6 (p=0.0101) when compared to controls (**Fig 2b**), but showed no significance when stratified by HIV VL (**Sup table 4).** Finally, CD86 expression was elevated in CD14+ cells at M0 (p=0.0005), M3 (p=0.0336), and M6 (p=0.0116) (**Fig 2b**) among PWH/OUD+ when compared to controls. CD86 expression was also elevated at periods of both high (p=0.0001) and low (p=0.0085) HIV VL among PWH/OUD+, when compared to controls (**Fig 3a**).

The frequency of CD14⁺⁺CD16⁻ monocytes were reduced among PWH/OUD+ as compared to controls at M0 (adj.p = 0.0082), M3 (adj.p = 0.0369), and M6 (p = 0.0126) (**Fig. 2c**). This association was also observed at periods of both high (p=0.0010) and low (p=0.0002) HIV VL among PWH/OUD+, when compared to controls (**Fig. 3b**). Intermediate CD14⁺⁺CD16⁺ monocytes were increased at M0 (p = 0.0407), M6 (p = 0.0290) (**Fig. 2c**), and elevated at both periods of high (p=0.0006), and low (p=0.0001) HIV VL, when compared to controls (**Fig. 3b**). Nonclassical CD14⁺CD16⁺⁺ monocytes were increased at M0 (adj.p = 0.0047), M3 (adj.p = 0.0301), M6 (p = 0.0157) (**Fig. 2c**), and elevated in both periods of high (p=0.0016), and low (p=0.0001) HIV VL when compared to controls (**Fig. 3b**).

In-depth analysis of monocyte subsets revealed increased CD86 expression on classical CD14⁺⁺CD16⁻ monocytes at M0 (p = 0.0001), M3 (p = 0.0449), M6 (p = 0.0263) among PWH/OUD+ as compared to controls (**Fig. 2d**). This association was also observed at periods of high (p=0.0079) and low (p=0.0005) HIV VL among PWH/OUD+, when compared to controls (**Fig 3c; Sup table 5**). Among CD14⁺⁺CD16⁺ intermediate monocytes, CD38 expression was reduced at M0 (p = 0.0305) and M6 (p = 0.0230), and CCR2 expression was reduced at M0 (p = 0.0146), M3 (p = 0.0443) (**Fig. 2e**) when compared to controls. This association was not observed when comparisons were made between periods of high and low HIV VL and the control groups, however (**Sup Table 5).** CD11b expression was significantly decreased at M0 (p = 0.0220), M3 (p = 0.0504) among PWH/OUD+ as compared to controls (**Fig. 2e**). This association was also observed at periods of high (p=0.0079) and low (p=0.0258) HIV VL as compared to controls (**Fig. 3c**). CD86 expression was increased when comparisons were made between periods of high (p=0.0111) and low (p=0.0005) HIV VL and the control group (**Fig. 3c**). Among PWH/OUD+, nonclassical CD14⁺CD16⁺⁺ monocytes demonstrated reduced CD38 expression at M0 (p = 0.0201), M3 (p = 0.0354), and M6 (p = 0.0162) compared to controls, and reduced CCR2 expression at M0 (adj. p = 0.0005), M3 (adj. p = 0.0002), and M6 (adj. p = 0.0002) **(Fig. 2f**). These associations were not observed when comparisons were made between periods of high and low HIV VL and the control group (**Sup table 5**). CD11b expression was reduced when stratified comparisons were made between periods of high (p=0.0266) and low (p=0.0152) HIV VL and the control group **(Fig 3c**). CD86 expression was elevated when stratified comparisons were made between periods of high (p=0.0111) and low (p=0.0005) HIV VL and the control group (**Fig. 3c**).

Longitudinal analysis of CD14+ monocytes was performed among PWH/OUD+ over a 6-month period (**Fig. 2a**). Expression of CD86, CD16, and HLA-DR decreased over 6-months, while CD11b, CCR7, CD38, and CCR2 expression were lower at M0 but increased by M6 (**Sup Table 4**). Classical monocytes CD14⁺⁺CD16⁻ were lower at M0 but increased by M6 (**Sup Table 6**), whereas nonclassical monocytes CD14⁺CD16⁺⁺ were elevated at M0 and decreased at M6 (**Sup Table 7**). Myeloid-derived suppressor cell (MDSC) populations (CD14⁺/CD15⁻/HLA-DR⁻) were significantly increased at M0 in PWH/OUD+ and decreased by M6 (**Sup Table 4**). When examining classical CD14⁺⁺CD16⁻ monocytes in PWH/OUD+ over time, CCR7, CD86, and HLA-DR expression decreased over 6 months (**Sup Table 6**). Among CD14⁺⁺CD16⁺ monocytes, CD86 expression also decreased over time, and CD36, CD27, CCR5, and CCR2 expression increased (**Sup Table 7**). Longitudinal analysis of CD14⁺CD16⁺⁺ revealed decreased expression of CD86 and CXCR4 over time, although CCR7 expression increased during the same time period (**Sup Table 8**).

### OUD is associated with limited cytokine production from CD14+ monocytes

Prior work has shown that monocyte-derived macrophages exposed to opioids exhibit altered cytokine responses to innate stimuli (Limiroli et al., 2002; Martucci et al., 2007; Odunayo et al., 2010), and we previously reported limited cytokine responses to LPS from PBMC isolated from PWH/OUD+ with viral suppression (Underwood et al., 2020). To assess the impact of COE on monocyte cytokine directly, CD14+ monocytes were isolated from PBMC of PWH/OUD+ and control participants, and cytokine responses to LPS stimulation quantified (**Fig 3d; Sup Table 9**). Overall, CD14+ monocytes from PWH/OUD+ demonstrated reduced cytokine responses compared with monocytes from controls, regardless of HIV VL. Stratified analysis by HIV VL demonstrated significant reductions in production of Interleukin-10 (IL-10) at periods of high (p=0.0125) and low (p=0.0549) HIV VL, when compared to the control group (**Sup Table 10**). Similarly, tumor necrosis factor-α (TNF-α) production was significantly lower at periods of high (p=0.0016) and low (p=0.0115) HIV VL when compared to the control group. Finally, Macrophage inflammatory protein-3 alpha (MIP-3α) production was significantly reduced at periods of high (p=0.0159) and low (p=0.0021) HIV VL when compared to the control group. However, Interleukin-1beta (IL-1β) was significantly decreased only when comparisons were made between the period of low HIV VL (p=0.0418) among PWH/OUD+ and the control group (**Fig. 3d**). These cytokine profiles suggest that, regardless of HIV viremia, OUD is associated with dampened cytokine response to LPS, a potent activator of innate immunity.

### OUD is associated with altered T, NK, and B cells phenotypes among PWH

Detailed phenotyping of T, NK, and B cells was also performed using spectral flow cytometry and data analyzed using both unsupervised (**Figure 4a**) and supervised (**Figure 4b-e; Sup Table 11-14**) approaches. Among CD3⁺CD4⁺ T cells, CCR5 expression was reduced at M0 (p=0.0290), M3 (p=0.0240), and M6 (p=0.0384) when compared to the control group (**Fig. 4a,b).** However, this association was not observed when stratified comparisons were made between periods of high and low HIV VL, and the control group (**Sup table 5**). CD57 showed decreased expression at M0 (p=0.0086), M3 (p=0.0136), and M6 (p=0.0215) when compared to controls (**Fig. 4a,b)**, and when comparisons were made during high HIV VL timepoints among PWH/OUD+, and the control group (p=0.0009) (**Fig. 5a**). Expression of CD27 was elevated at M3 (p=0.0458) and M6 (p=0.0498) when compared to the control group (**Fig. 3b**); this association was also observed when comparisons were made between periods of high (p=0.0206) and low (p=0.0283) HIV VL and the control group (**Fig. 5a**). CD3⁺CD8⁺ T cells from PWH/OUD+ participants exhibited increased GLUT1 expression at M0 (p = 0.0188), M3 (p = 0.0492), and M6 (p = 0.0130) (**Fig. 3c**) when compared to controls. This association was also observed when comparisons were made between periods of high (p=0.0161) and low (p=0.0343) HIV VL and the control group (**Fig. 5a**). When examining CD56⁺ NK cells, HLA-DR expression was significantly elevated at M0 (p = 0.0161), M3 (p =0.0260), M6 (p = 0.0201) when compared to controls (**Fig. 3d**). This association was also observed when comparisons were made between periods of high (p=0.0152) and low (p=0.0062) HIV VL and the control group (**Fig. 5b)**. Conversely, NK cell expression of CD57 was reduced at M0 (p = 0.0138), M3 (p = 0.0213), M6 (p = 0.0118) when compared to controls (**Fig. 3d**). This association was also observed when comparisons were made between periods of high (p=0.0274) and low (p=0.0031) HIV VL, and the control group (**Fig. 5b)**. CD86 was elevated at M0 (p = 0.0109) and M6 (p = 0.0473) among PWH/OUD+, when compared to the control group (**Fig. 3d);** however, this association was not maintained when comparisons were made between periods of high and low HIV VL and the control group (**Sup table 5**). Finally, CD19⁺ B cells exhibited reduced CCR7 expression at M0 (adj. p = 0.0011), M3 (adj. p = 0.0004), and M6 (adj. p = 0.0008) among PWH/OUD+, when compared to the control group (**Fig. 3e**). This association was also observed when comparisons were made between periods of high (p=0.0001) and low (p=0.0008) HIV VL, and the control group (**Fig. 5c)**.

**Figure 4.**
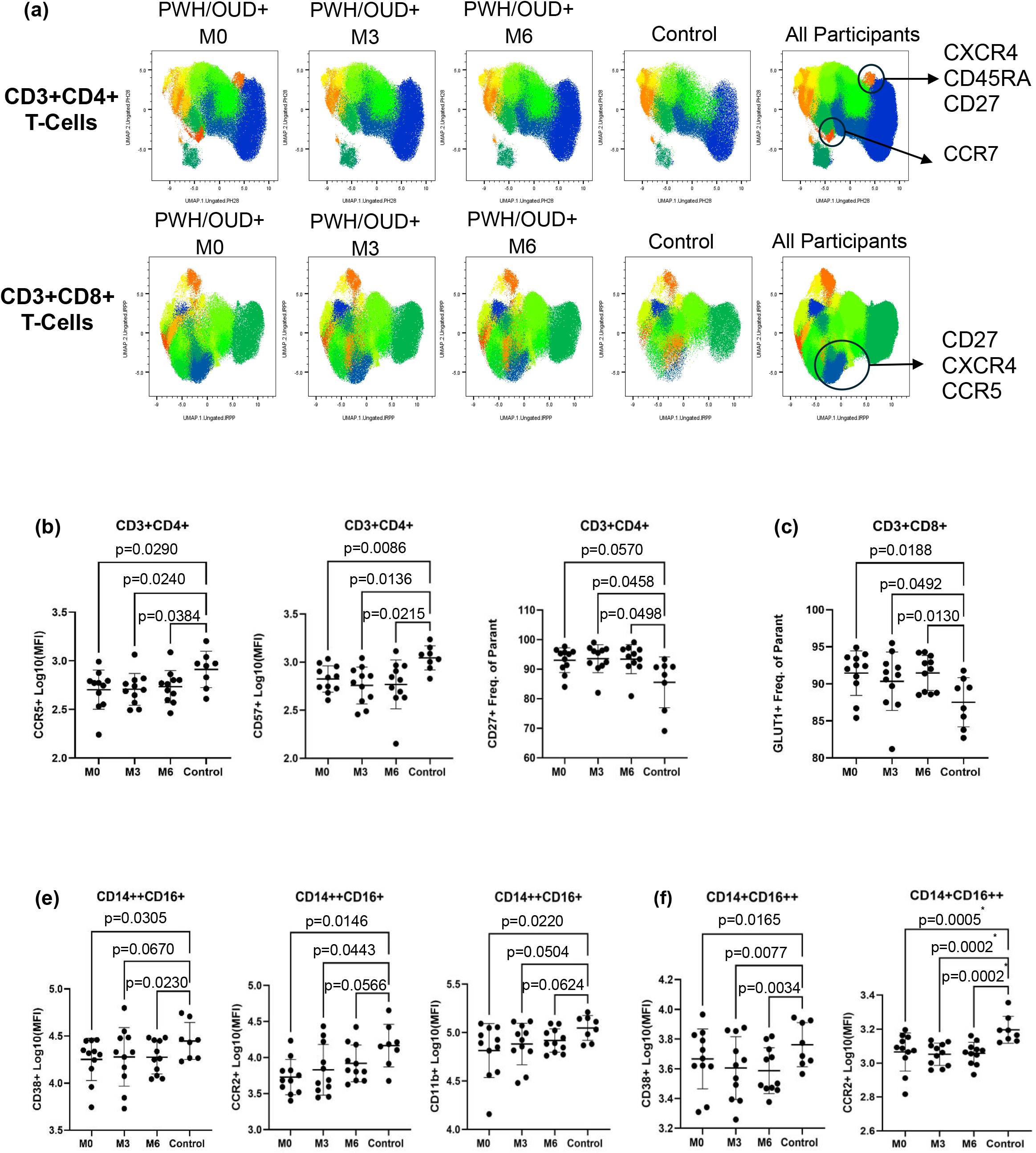
Altered lymphocyte phenotypes among PWH/OUD+. **(a)** UMAP visualization of CD3⁺CD4⁺ and CD3⁺CD8⁺ T cells over time among PWH/OUD+ (n=11) and control participants (n=8). Annotation on CD3⁺CD4⁺ T cells for markers CXCR4, CD45RA, CD27 and CCR7. Annotation of CD3⁺CD8⁺ T cells for markers CD27, CXCR4 and CCR5. **(b)** MFI of CCR5 and CD27 expression, and frequency of CD3⁺CD4⁺CD27⁺ T cells, from PWH/OUD+ across timepoints M0, M3, and M6 compared to controls. Frequencies and MFI were calculated from total live CD14⁻CD19⁻CD56⁻CD3⁺CD4⁺ cells (Sup Fig 2). **(c)** Frequency of GLUT1 expression on CD3⁺CD8⁺ T cells from PWH/OUD+ across M0, M3, and M6 timepoints compared to controls. Frequencies were calculated from total live CD14⁻CD19⁻CD56⁻CD3⁺CD8⁺ cells (Sup Fig 2). **(d)** MFI of HLA-DR and CD86 on CD56⁺ NK cells from PWH/OUD+ across all three timepoints M0, M3, and M6 compared to controls. MFI was calculated from total live CD3⁻CD14⁻CD19⁻CD56⁺ cells (Sup Fig 3). **(e)** MFI of CCR7 on CD19⁺ B cells across all three timepoints M0, M3, and M6 in PWH/OUD+ participants compared to controls. MFI was calculated from total live CD3⁻CD14⁻CD56⁻CD19⁺ cells (Sup Fig 4). Cross-sectional analyses were performed using mixed-effects repeated-measures ANOVA with compound symmetry to account for within-subject correlation. HIV viral load (log₁₀[viral load + 1]) was included as a covariate; Tukey correction was applied for multiple comparisons. Shown are log10 transformed mean fluorescence intensity (MFI) with SD, and mean frequencies with SD. P values adjusted for multiple comparisons are denoted with *.

**Figure 5.**
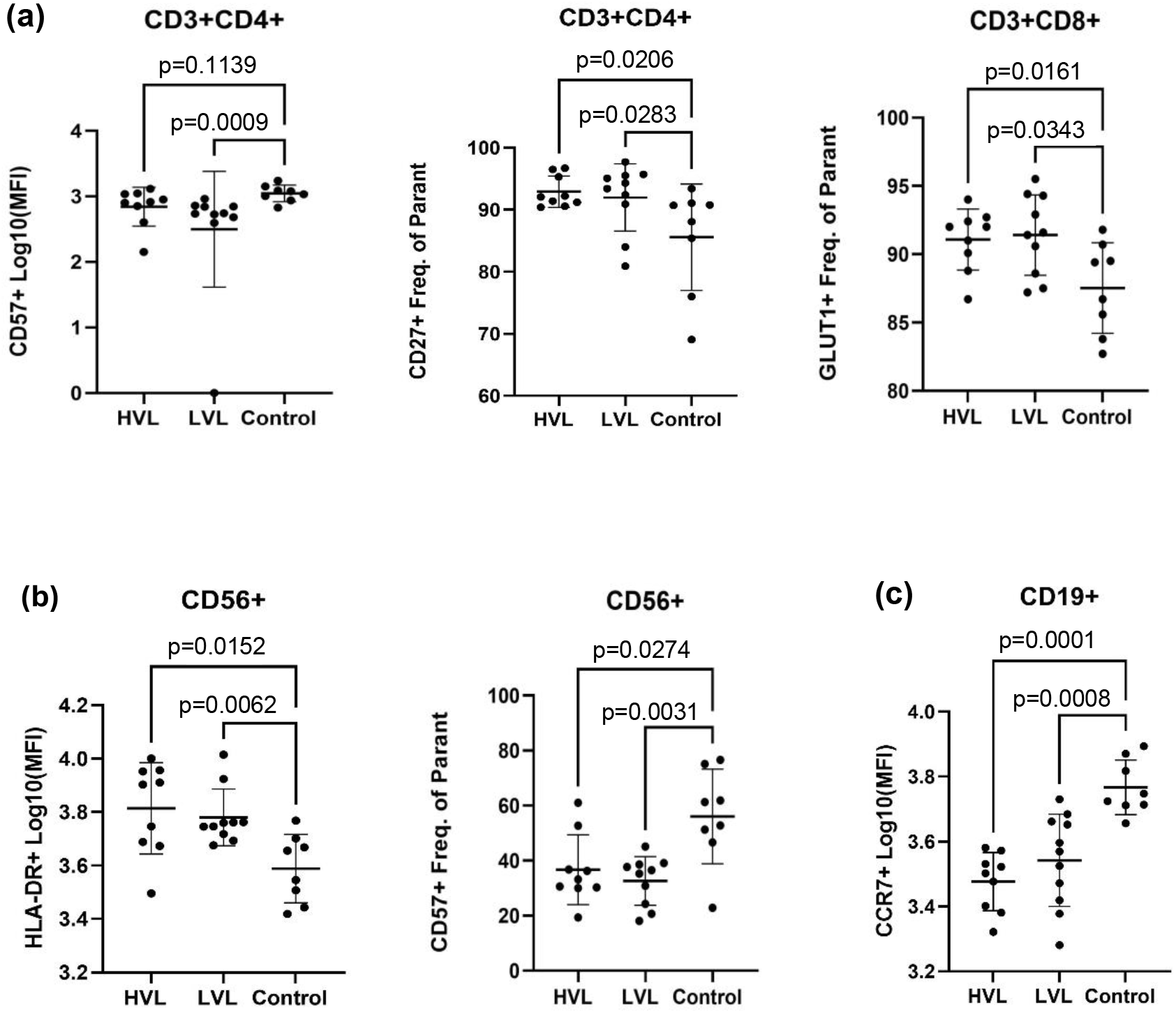
Stratification by HIV Viral load supports association between OUD and altered immune cellular phenotypes and functional responses. **(a)** The frequency of CD27 and MFI of CD57 and on CD3⁺CD4⁺ T cells, and the frequency of GLUT1 on CD3+CD8+ T cells of both HVL and LVL groups compared to the control. **(b)** Shown are MFI of HLA-DR and frequencies of CD57 in CD56⁺ NK cells among HVL and LVL groups compared to control. **(c)** Shown is the MFI of CCR7 on CD19⁺ B cells in PWH/OUD+ in both HVL and LVL groups compared to control. P values adjusted for multiple comparisons are denoted with *.

In a longitudinal analysis over 6 months in PWH/OUD+, CD3⁺CD4⁺ T cells showed increased expression of CD36, and co-expression of CD28+CD57+, while there was decreased expression of CD28, CCR2, HLA-DR, CXCR4, and co-expression of CD38+HLADR+, over time (**Sup table 11**). CD3⁺CD8⁺ T cells showed increased expression of CCR7 over 6 months, while expression of CD57, CXCR4, and PD-1 significantly decreased (**Sup table 12**). CD56⁺ NK cells showed elevated levels of CCR7 and PD-1 over 6 months, and decreased expression of CD16 and CXCR4 over time **(Sup table 13**). In CD19+ B Cells, there was increased expression of PD-1 and decreased expression of CXCR4 over 6 months (**Sup table 14**).

## Discussion

PWH are at increased risk for OUD, a comorbidity associated with worse health outcomes. However, the consequences of long-term opioid use on immunity and inflammation among PWH remain poorly understood. Although ART is highly effective at controlling HIV viremia, many PWH still show signs of ongoing immune imbalance and chronic inflammation that have been linked to increased risk of chronic co-comorbidities such as cardiovascular disease, HAND, and some malignancies. It remains unclear, however, if long-term opioid exposure is an independent driver of immune imbalance among PWH and could contribute to risk of these chronic diseases. Here, we identified numerous alterations in systemic and innate and adaptive cellular immune responses associated with COE. First, we demonstrated that PWH/OUD+ have sustained alterations in systemic biomarkers of innate immune activation and altered cytokine profiles, even with successful viral suppression, when compared to PWH/OUD-. PWH/OUD+ also exhibited expansion of CD16+ intermediate and non-classical monocyte populations, a finding we had previously reported among a cohort of PWH/OUD+ individuals with well-controlled HIV VL. Building on these findings, we have shown that monocytes from PWH/OUD+ have dysregulated functional responses to innate immune stimuli. We also identified opioid-associated alterations in expression of proteins supporting adhesion and migration in a variety of cell types among PWH/OUD+. Collectively, our results suggest that COE is associated with broad changes in cellular immunity that are not restricted to the innate immune system. Notably, the immunologic features distinguishing PWH with COE were observed among individuals with both poorly and well controlled HIV VL. Overall, our findings suggest that COE is associated with broad and persistent changes to human peripheral immunity among PWH, and that OUD should be considered a significant risk factor for immune dysregulation regardless of control of HIV viremia.

Sustained elevation of biomarkers reflecting innate immune activation have been strongly associated with HIV disease progression, increased mortality, as well as increased risk of chronic diseases driven by immune dysregulation (Dubrow et al., 2012; He et al., 2025; McGuire et al., 2015; Wallis & Williams, 2022; Zicari et al., 2019). Here we quantified systemic biomarkers reflective of innate immune activation associated with poor outcomes among PWH, sCD163 and sCD14. We found that both sCD163 and sCD14 were persistently elevated among PWH/OUD+ compared with PWH/OUD-. The association between COE and elevation of sCD163 and sCD14 has been previously reported by both our and other groups among PWH with well-controlled HIV VL (Ahmed et al., 2025; Azzoni et al., 2022; Dang et al., 2023; Hileman et al., 2022; Kholodnaia et al., 2022; Monnig et al., 2019; Underwood et al., 2020). Here, we are able to follow individuals who remained exposed to opioids as ART was optimized, to demonstrate that clinically relevant biomarkers of innate immune activation persist among individuals with OUD despite successful reductions in HIV VL.

Several mechanisms may drive systemic innate immune activation among PWH, including the negative impact of HIV infection on intestinal epithelial barrier function, leading to ongoing microbial translocation (Balena et al., 2020; Ellis et al., 2021; Monnig et al., 2019). However, emerging research also suggests that opioids themselves may lead to disruption of the intestinal epithelium (Hileman et al., 2022; Wang & Roy, 2017). Here, we aimed to determine if there was a relationship between COE and changes in systemic biomarkers of gut epithelial damage and microbial translocation, specifically I-FABP, IL-37, and LBP. In our cohort, no significant differences in these biomarkers were identified between PWH with and without OUD, suggesting that intestinal barrier dysfunction is unlikely to be the primary driver of the innate immune activation observed among individuals with OUD.

Peripheral blood cytokines and chemokines play an important role in orchestrating proper immune responses, and disruptions in their production or regulation can be a key factor in driving dysfunctional responses to acute infection and chronic disease. Research into plasma cytokine signatures among PWH/OUD+ is limited. Here, we broadly examined systemic cytokine profiles among PWH with and without COE and identified elevated levels of two cytokines key to governing chemotaxis, fractalkine and ITAC, as well as reduced levels of the key regulatory cytokine, TGF-β1, among PWH/OUD+. Fractalkine facilitates leukocyte adhesion and chemotaxis in monocytes and T cells (Rodriguez et al., 2024) and has a well-documented association with comorbidities in PWH, specifically HIV-associated neurocognitive disorders (HAND) (Chamera et al., 2019; Mountford et al., 2018; Panigrahi et al., 2020). ITAC (CXCL11) is a chemokine induced by IFN-γ, and acts as a ligand for CXCR3 (Groom & Luster, 2011; Tokunaga et al., 2018). CXCR3 has been well documented as an important ligand in neuroinflammation and recruitment of immune cells to the CNS during HIV infection (Hauser et al., 2007; White et al., 2005). TGF-β1 is essential for immune regulation and maintenance of homeostasis (Wang et al., 2023), but contributes to tissue remodeling and fibrosis that can worsen underlying organ disease. In HIV, chronic immune activation has been shown to sustain TGF-β1 production, thereby contributing to both overall immunosuppression and fibrotic remodeling in several organs (Laurence et al., 2017; Theron et al., 2017). Taken together, this cytokine pattern suggests that the peripheral plasma environment in PWH/OUD+ is enriched for signals promoting immune cell trafficking and adhesion, while simultaneously exhibiting reduced regulatory balance. Our results are the first to demonstrate such complex alterations in systemic plasma cytokine levels among PWH/OUD+ that persist despite successful reductions in HIV VL.

Alterations in monocyte subtypes and inflammatory responses are associated with numerous chronic diseases, such as rheumatoid arthritis, cardiovascular disease, atherosclerosis, and dementia, among individuals living with and without HIV (Chistiakov et al., 2018; Cooper et al., 2012; Czepluch et al., 2014; Genkel et al., 2023; Iwahashi et al., 2004). In the context of HIV infection, CD16+ monocytes (nonclassical and intermediate) are more vulnerable to cellular HIV infection, permissive to viral replication, and exhibit distinctive inflammatory and migratory responses to innate stimuli (Campbell et al., 2014; Cattin et al., 2021; Nabatanzi et al., 2019; Nowlin et al., 2018). We previously reported that PWH/OUD+ with well-controlled HIV VL demonstrate expansion of CD16+ nonclassical and intermediate monocyte subtypes (Underwood et al., 2020). We confirm this finding here, where expansions of these populations were found to persist as individuals with ongoing COE underwent optimization of ART and reductions in HIV VL. We further identified a marked shift in the expression of additional surface proteins on monocytes that support migratory, inflammatory, and antigen-presenting capacities in individuals with COE. For example, CD86 expression was persistently elevated on classical monocytes among PWH/OUD+. CD86 is a co-stimulatory molecule that plays an important role in T-cell activation and immune signaling, and its upregulation is commonly associated with monocyte activation, suggesting enhanced antigen-presenting capacity and a potential shift toward increased inflammatory responsiveness (Funderburg et al., 2011; Guest et al., 2008; Kapellos et al., 2019; Souza et al., 2007). HIV viremia has been shown to drive CD86 expression (Beuria et al., 2005; Papasavvas et al., 2008; Zhao et al., 2002), but the role of opioid exposure has not been previously reported. Intermediate and nonclassical monocytes exhibited reduced expression of CCR2 and CD38 across all time points. Expression of CCR2 on monocytes facilitates transmigration across the blood–brain barrier and contributes to neuroinflammatory processes in HIV-associated (White et al., 2005; Williams et al., 2014) but little is known about CCR2 expression in the context of opioid exposure. Previous studies on CCR2 expression on monocytes indicate increased expression in the context of HIV infection and an association with HAND (Veenstra et al., 2019; Williams et al., 2013, 2014). Here, CCR2 expression was significantly reduced on monocytes in PWH/OUD+. This finding was unexpected due to the association between OUD and dementia, and calls for further study of how opioids alter the propensity for monocytes to drive neuroinflammation and cognitive disorders. CD38 is associated with monocyte differentiation and activation (García-Rodríguez et al., 2018; Schneider et al., 2015) and is associated with heightened immune responses during chronic inflammatory states such as HIV (Kelesidis et al., 2016). OUD has been associated with the upregulation of CD38 on CD28-CD57+ senescent CD8+ T cells (Dang et al., 2023), but no association between opioid exposure and monocyte CD38 expression has been previously reported. CD11b was also significantly decreased in intermediate and nonclassical monocytes. CD11b is an integrin involved in cell adhesion and transmigration that facilitates interactions between monocytes and endothelial cells during migration into inflamed tissues (Thaler et al., 2016), and in the context of HIV, CD11b+ monocytes contribute to heightened inflammation. Although the mechanism of opioid-induced changes in monocyte CD11b expression is unknown, prior *in vitro* studies have shown that morphine can induce alterations in expression of cellular adhesion molecules (CAMs) (Machelska et al., 2004; Machelska & Celik, 2020; Stein & Machelska, 2011), such as vascular cell adhesion molecule 1 (VCAM-1) in cancer (Ghasemi et al., 2023) and endothelial cells (Strazza et al., 2016). Taken together, our data indicates that PWH/OUD+ exhibit a redistribution of monocyte subsets toward populations commonly associated with inflammatory responses, while simultaneously displaying reduced expression of molecules involved in adhesion, migration, and activation. These findings broadly support the conclusion that COE is associated with persistent alterations in the functional capacities of monocytes, key effector cells of the innate immune system.

Beyond alterations in monocyte subsets, PWH/OUD+ also exhibited phenotypic changes across multiple immune cell populations, further supporting disruption of immune trafficking and intercellular communication. Although the overall frequencies of CD4⁺ T cells, CD8⁺ T cells, CD19⁺ B cells, and CD56⁺ NK cells were comparable between individuals with and without OUD, differences in cellular expression of key surface markers involved in migration, activation, and tissue homing were observed. These findings suggest that opioid exposure alters immune cell phenotypes without substantially affecting overall population abundance.

Within the CD3⁺CD4⁺ T cell compartment, reduced CD57 expression and increased CD27 expression were observed. CD57 is commonly associated with terminal differentiation, replicative senescence, and cytotoxic potential in T cells (Chen et al., 2020; Hassouneh et al., 2017), and alterations in CD57⁺CD28⁻ T-cell populations have been reported in PWH/OUD+ (Dang et al., 2023). The reduced CD57 expression observed here suggests a shift away from terminally differentiated phenotypes. In contrast, CD27 is a costimulatory molecule whose loss is typically associated with progression toward effector differentiation (Schuetz et al., 2011). In HIV infection, CD27 expression is often reduced and linked to impaired proliferative capacity. The increased CD27 expression observed in PWH/OUD+, therefore, contrasts with canonical HIV-associated patterns and suggests that opioid exposure may alter CD4⁺ T-cell differentiation states. Additional alterations in CD3⁺CD4⁺ T cells were observed in CCR5 expression, which was reduced. CCR5 is a chemokine receptor that mediates trafficking of Th1-type cells to sites of inflammation and also functions as a key co-receptor for HIV entry into CD4⁺ T cells (Björndal et al., 1997; Connor et al., 1997; Wu et al., 1997). While ex vivo studies have shown that opioid exposure can increase CCR5 expression in activated CD4⁺ T cells (Banerjee et al., 2011; Madhuravasal Krishnan et al., 2023; Trunfio et al., 2023) the reduced CCR5 expression observed in PWH/OUD+ highlights the complexity of opioid immune interactions *in vivo* and suggests that chronic exposure may differentially regulate trafficking associated receptors as compared to short term *in vitro* exposures.

In CD3⁺CD8⁺ T cells, increased expression of the glucose transporter GLUT1 was observed. GLUT1 is a key regulator of cellular metabolism and is upregulated in activated T cells to support glycolytic demand (Cammann et al., 2016; Palmer et al., 2015). Increased GLUT1 expression has been reported in CD8⁺ T cells during HIV infection and is associated with altered metabolic programming (Kang & Tang, 2020; Masson et al., 2017; Palmer et al., 2016). While the effects of OUD on CD8⁺ T cell metabolism have not been well characterized, opioid mediated alterations in GLUT1 expression have been described in other cell types, including microglia (Olianas et al., 2011; Shivling Mali et al., 2023). These findings suggest that opioid exposure may contribute to metabolic reprogramming of CD8⁺ T cells in PWH/OUD+.

Analysis of CD56⁺ NK cells revealed increased HLA-DR and CD86 expression alongside reduced CD57 expression in PWH/OUD+. Elevated HLA-DR expression on NK cells has been reported in PWH despite ART (Hong et al., 2010; Naidoo & Altfeld, 2024), and our findings suggest that OUD may further enhance this activated phenotype. CD86, a co-stimulatory molecule typically associated with antigen-presenting cells, is minimally expressed on resting NK cells but increases upon activation (Cruz-González et al., 2018; Senju et al., 2018; Skak et al., 2008). The increased expression of CD86 observed here supports an activated NK cell phenotype in PWH/OUD+. In contrast, CD57, a marker of terminal maturation and cytotoxic potential in NK cells (Cao et al., 2015; Morice, 2007), was reduced; this contrasts with the increased CD57 expression typically observed during well treated HIV-infection (Abou Hassan et al., 2019; Affandi et al., 2015; Meier et al., 2005). Together, these findings suggest that NK cells in PWH/OUD+ exhibit features of activation while displaying altered maturation states.

In CD19⁺ B cells, CCR7 expression was persistently increased among PWH/OUD+. CCR7 is a chemokine receptor that directs B-cell homing to secondary lymphoid tissues and is critical for migration to lymph nodes, where germinal centers support antigen presentation (Coraglia et al., 2016; Kealy et al., 2025; Payne et al., 2009). In the context of HIV infection, CCR7 expression is often reduced and associated with disrupted B cell trafficking and the expansion of atypical or exhausted memory B cell populations, with partial restoration following ART (Boswell et al., 2014; Liechti et al., 2019; Rakhmanov et al., 2009). Elevated CCR7 expression may indicate enhanced homing to lymphoid tissues or retention within lymphoid compartments, potentially influencing antigen presentation and B-cell activation states. These findings suggest that OUD may modify B cell migration and differentiation pathways in PWH, contributing to altered immune coordination.

Our earlier work demonstrated reduced cytokine production from LPS-stimulated PBMCs isolated from individuals with PWH/OUD+ and well-controlled HIV VL (Underwood et al., 2020). Here, using isolated CD14+ monocytes, we demonstrated that COE is associated with a reduced capacity of monocytes to generate both pro- and anti-inflammatory cytokines in response to innate stimuli, regardless of HIV VL, including IL-1β, TNF-α, IL-10, and MIP-3α. IL-1β production in CD14⁺ monocytes is tightly linked to activation of the NLRP3 inflammasome, and prior studies have demonstrated that NLRP3 signaling is dysregulated in HIV-infected monocytes (Guerville et al., 2023; Rolfes et al., 2020; Stunnenberg et al., 2021). The reduced IL-1β production observed in our study is consistent with impaired inflammasome activation, suggesting that opioid exposure further interferes with NLRP3 signaling among PWH. Although production of TNF-α and IL-10 are mediated through distinct cellular processes, altered production of these cytokines has also been reported among PWH on ART (Ramendra et al., 2019; Thurman et al., 2020). Although the specific effects of COE on monocyte cytokine production in PWH remain incompletely defined, opioid exposure has been broadly shown to modulate cytokine signaling pathways, including TNF-α and IL-10 regulation in microglial cells (Azzoni et al., 2022; Bekhbat et al., 2018; Trunfio et al., 2023; Xiao et al., 2022). In addition to inflammatory cytokines, the reduced MIP-3α (CCL20) production by isolated monocytes suggests altered chemokine signaling, as MIP-3α binds CCR6 and plays an important role in directing immune cell trafficking to mucosal and central nervous system compartments during inflammation (Alessia Verani et al., 1997; Geonnotti et al., 2010; Rock et al., 2004). In HIV infection, dysregulated chemokine signaling contributes to altered immune cell recruitment to the CNS and has been implicated in the development of HAND (Jaureguiberry-Bravo et al., 2018; Lu & Wang, 2020; McGuire et al., 2015; Trunfio et al., 2023). The observed reduction in MIP-3α production suggests that opioid exposure may impair chemokine-mediated immune trafficking in addition to altering inflammatory signaling and supports our observation of altered systemic levels of several chemokines in the context of COE. Taken together, these findings demonstrate that monocytes from PWH/OUD+ exhibit impaired functional responses to pathogen-associated stimulation despite evidence of systemic immune activation. More specifically, we found that opioid exposure induced a state of innate immune miscoordination, where monocytes are chronically activated yet have reduced capacity to mount appropriate migratory, inflammatory, and regulatory responses.

Several limitations should be considered when interpreting these findings. The reference population (PWH/OUD−) was predominantly male-sex and exclusively HCV-negative, whereas the PWH/OUD+ cohort included individuals of both sexes and with varying HCV status, potentially influencing some of the observed phenotypes and inflammatory biomarkers. CTN-067 participants were followed for only six months after starting and/or optimizing ART, and may not yet have resolved the inflammatory changes induced by HIV viremia. The relatively small sample size for flow cytometric and functional studies may have limited the ability to detect more subtle immune alterations and limited our statistical power. Finally, as an observational study, our findings demonstrate associations between chronic opioid exposure and immune dysregulation but cannot establish causality or define underlying mechanisms. Despite these limitations, this study leverages a unique longitudinal cohort of PWH/OUD+ undergoing ART treatment and integrates plasma biomarkers and cytokines, immune phenotyping, and functional monocyte assays. The consistency of the observed findings across multiple time points, together with multiple analytic approaches to tease apart the impact of HIV viremia from opioid exposure, strengthens the conclusion that chronic opioid exposure is associated with persistent immune dysregulation in treated HIV infection.

Overall, our findings demonstrate that OUD is associated with numerous changes in mediators of systemic inflammation, as well as alterations across multiple immune cell types. Across monocytes, T cells, NK cells, and B cells, a consistent pattern emerges in which immune cells exhibit features of activation or redistribution while simultaneously displaying altered expression of key molecules required for proper migration and functional coordination. This supports a broader model in which chronic opioid exposure disrupts immune communication networks, contributing to dysregulated immune responses in PWH. Importantly, these phenotypic and functional changes persisted among individuals with OUD, regardless of control of HIV VL, suggesting that OUD serves as an independent factor contributing to immune dysregulation among PWH.

## Declaration of generative AI and AI-assisted technologies in the manuscript preparation process

Grammarly (Superhuman Platform Inc., San Francisco, CA) was used for grammatical correction.

Scite.ai (Research Solutions, Henderson, NV) was used for literature review.

## CRediT authorship contribution statement

Brent Bever: Writing original draft, Data curation, visualization, investigation, methodology, funding acquisition, Review and editing

Michelle Underwood-Ditto: Methodology, investigation, review, and editing

Byung Park: Formal analysis, review, and editing

Susan Pereira Ribeiro: Data curation, review, and editing

Ryan Cook: Data curation, review, and editing

Lynn Kunkel: Project administration, review, and editing

Todd Korthuis: Conceptualization, funding acquisition, review, and editing

Christina Lancioni: Conceptualization, methodology, supervision, funding acquisition, resources, Review and Editing.

## Funding

National Institutes of Health: NIDA: UG1DA015815, UG1DA013732, R03DA039731, R01DA046229 and R36DA061652, the Oregon Clinical & Translational Research Institute, and the Collins Medical Trust. The research reported in this publication used computational infrastructure supported by the Office of Research Infrastructure Programs, Office of the Director, of the National Institutes of Health under Award Number S10OD034224. The content is solely the responsibility of the authors and does not necessarily represent the official views of the National Institutes of Health. This project used the OHSU Flow Cytometry and Monoclonal Antibody Shared Resource Core Facility (RRID:SCR_009974).

## Declaration of competing interests

Authors have no competing interests to declare.

## Acknowledgments

We would like to thank all study participants for their valuable contribution to this work. We are grateful to the Oregon Clinical and Translational Research Institute and the Collins Medical trust. We are grateful to the CTN network for their technical assistance and EMMES for CTN data management. We would also like to thank the CTN network and all participating CTN-067 study sites for their support. We acknowledge and thank the staff of the OHSU HIV-comprehensive care clinic.

## Appendix A: Supplementary Material

## Data availability

The de-identified data supporting the findings of this study are available from the corresponding author upon reasonable request.

## Notes

### Competing Interest Statement

The authors have declared no competing interest.

