## Supplemental Figure 1 for "Chronic opioid-associated immune dysregulation among people living with HIV"

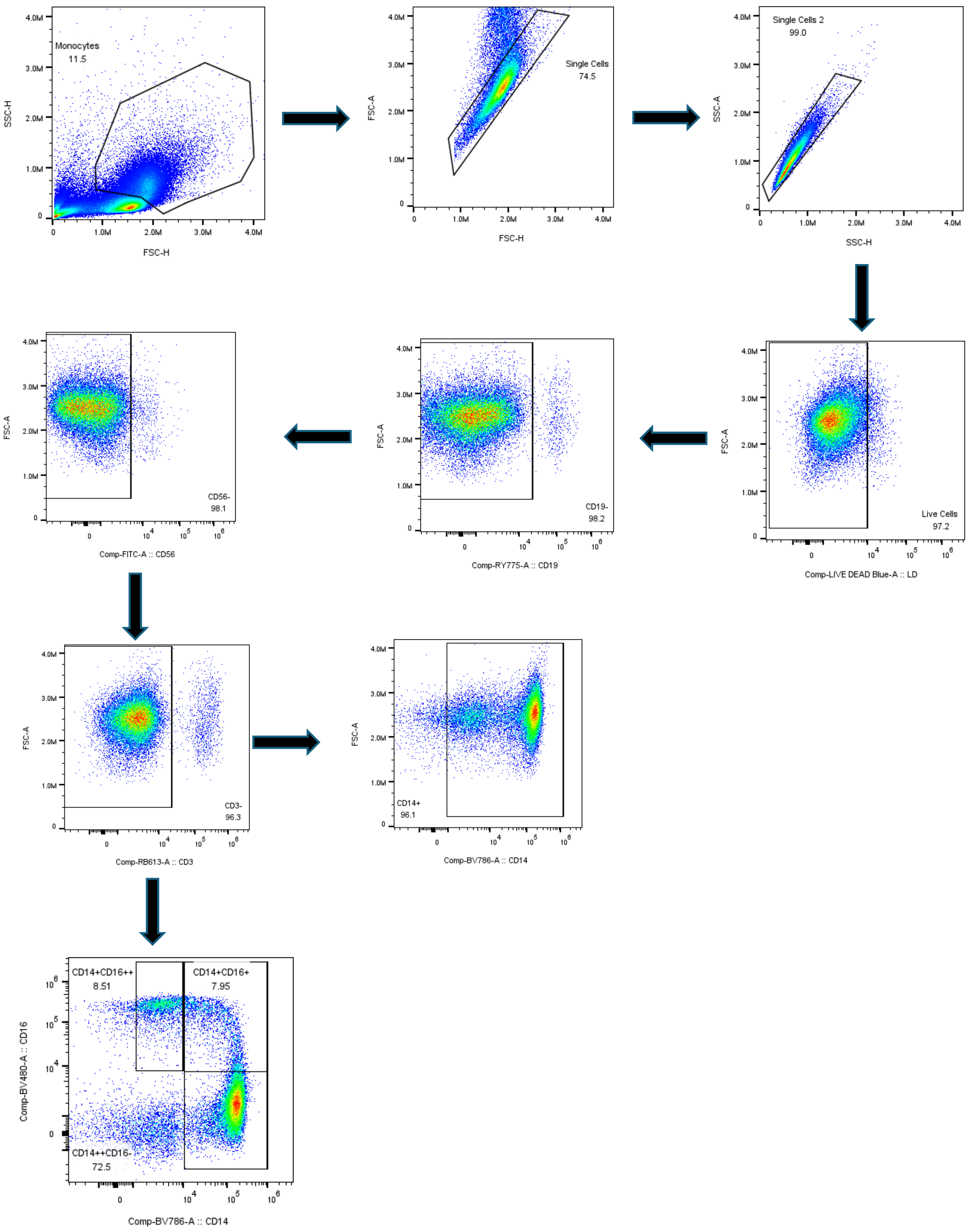


**Supplementary Figure 1.** Gating Strategy for CD14+ Cells, Classical (CD14++CD16-), Intermediate (Cd14++CD16+), and non-classical (CD14+CD16++) Monocytes.
