## Supplementary figures and images for "Chronic opioid-associated immune dysregulation among people living with HIV"

### Supplemental Figure 2

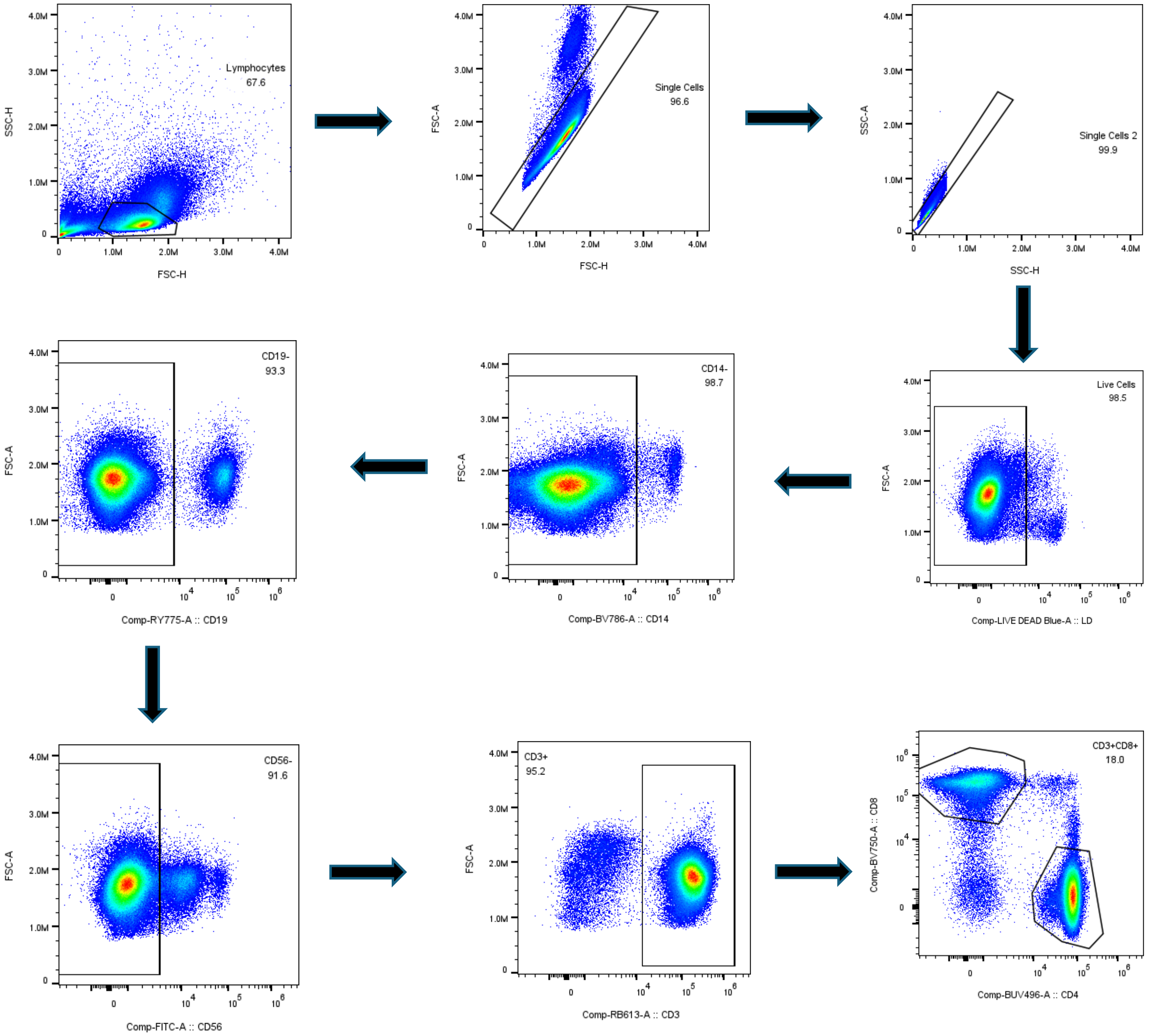


**Supplementary Figure 2.** Gating Strategy for CD3+CD4+ and CD3+CD8+ T Cells

### Supplemental Figure 3

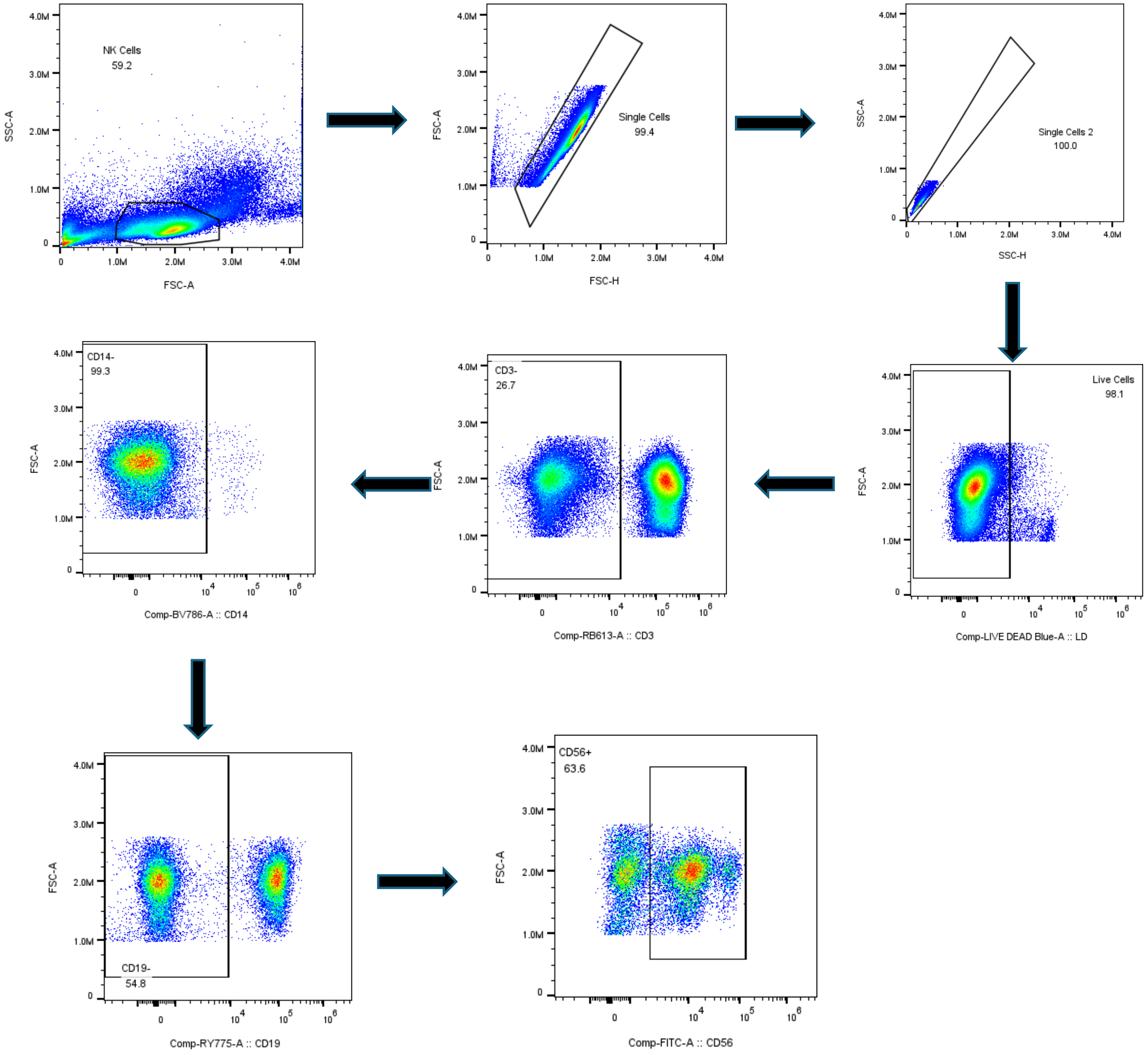


**Supplementary Figure 3.** Gating Strategy for CD56+ NK Cells

### Supplemental Figure 4

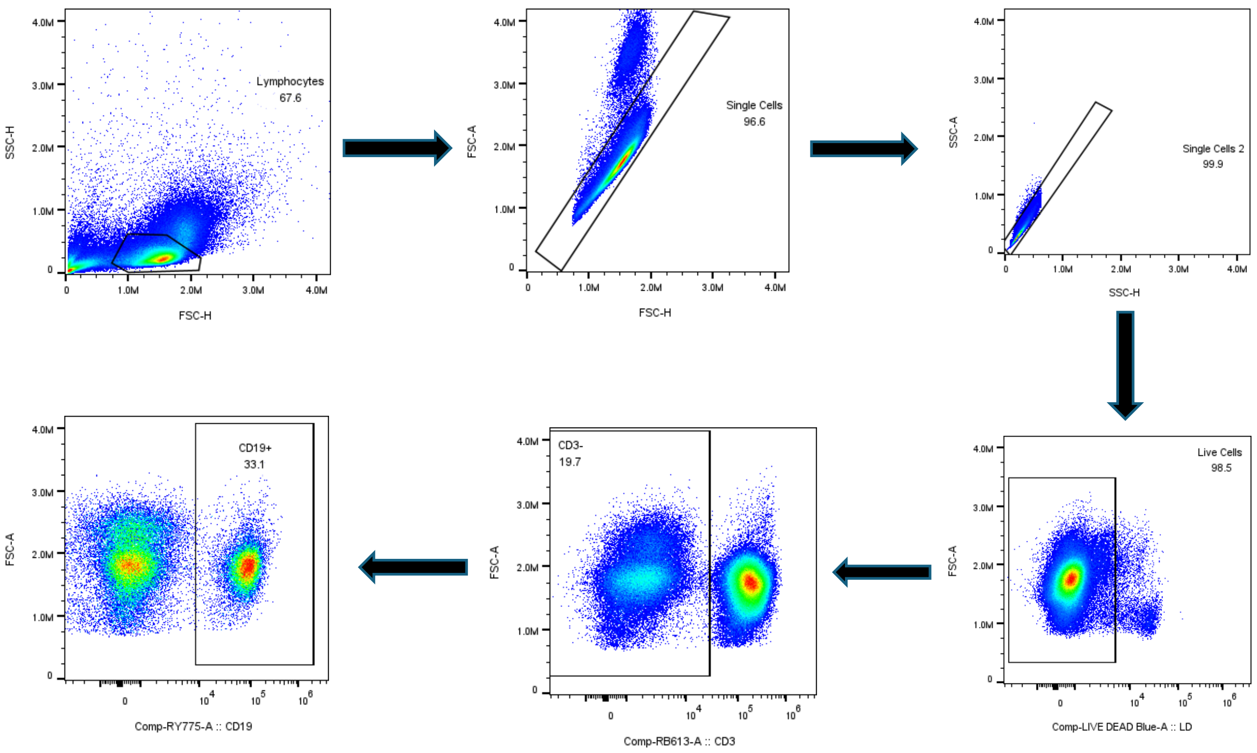


**Supplementary Figure 4.** Gating Strategy for CD19+ B Cells
