## Supplemental Table 1 for "Chronic opioid-associated immune dysregulation among people living with HIV"

**Supplementary Table 1.** Flow cytometry antibodies and reagents

| Antibody | Clone | Conjugate | Catalog # | Lot # | Manufacture |
|---|---|---|---|---|---|
| CD14 | M5E2 | BV786 | 563698 | 4053877 | BD Biosciences |
| CD16 | 3G8 | BV480 | 566106 | 4136809 | |
| CD15 | HI98 | BV650 | 564232 | 4113069 | |
| CD11b | ICRF44 | BUV615 | 752298 | 5083153 | |
| CCR2 (CD192) | LS132.1D9 | RB705 | 757633 | 5083150 | |
| CCR5 (CD195) | 2D7/CCR5 | R718 | 752091 | 5035058 | |
| HLA-DR | G46-6 | RB780 | 568767 | 4200389 | |
| CD86 | 2331 | BV711 | 563158 | 4323864 | |
| CXCR4 (CD184) | 12G5 | RY586 | 569646 | 4221075 | |
| CCR7 (CD197) | 2-L1-A | BV510 | 566760 | 4134264 | |
| GLUT1 | 202915 | Alexa Flour-647 | 566580 | 3240481 | |
| CD8 | RPA-T8 | BV750 | 747385 | 5035055 | |
| CD3 | UCHT1 | RB613 | 571085 | 4002470 | |
| CD4 | RPA-T4 | BUV496 | 569179 | 5063531 | |
| CD38 | HIT2 | PE | 560981 | 1153290 | |
| CD28 | CD28.2 | BUV395 | 569160 | 5010440 | |
| CD19 | SJ25C1 | RY775 | 571374 | 4100316 | |
| CD57 | HNK-1 | RB545 | 569740 | 4179206 | |
| PD1 | EH12.1 | RY703 | 571439 | 4106698 | |
| CD27 | M-T271 | BUV661 | 741609 | 5083926 | |
| CD45RA | HI100 | BV605 | 562886 | 4051131 | |
| CD69 | FN50 | RY610 | 571167 | 4267078 | |
| TLR4 (CD284) | HTA125 | BV421 | 743392 | 5083928 | |
| FC Block | NA | NA | 564220 | 5080041 | |
| Brilliant Stain Buffer Plus | NA | NA | 566385 | 4304047 | |
| Cell Viability | NA | Live Dead Fixable Blue | 2729825 | L34962 | ThermoFisher |
| CD56 | HCD56 | FITC | 318304 | B441180 | BioLegend |
| CD36 | 5-271 | APC-Fire750 | 336220 | B423582 | |
