## Supplemental Table 2 for "Chronic opioid-associated immune dysregulation among people living with HIV"

**Supplementary Table 2**. Peripheral blood ELISA Biomarkers

| Biomarker | Comparison (A vs. B) | Group A | | | Group B | | | Viral load adjusted<br>Group A - Group B | | | |
|---|---|---|---|---|---|---|---|---|---|---|---|
| | | N | Mean | SD | N | Mean | SD | Estimate | SE | p | adj p |
| Log10 (I-FABP) | Control vs. CTN-067 M0 | 44 | 2.874 | 0.239 | 59 | 2.954 | 0.274 | -0.077 | 0.056 | 0.1673 | 0.5076 |
| | Control vs. CTN-067 M3 | 44 | 2.874 | 0.239 | 50 | 2.990 | 0.317 | -0.112 | 0.056 | 0.0496 | 0.1993 |
| | Control vs. CTN-067 M6 | 44 | 2.874 | 0.239 | 53 | 2.945 | 0.271 | -0.068 | 0.056 | 0.2270 | 0.6180 |
| | CTN-067 M0 vs. CTN-067 M3 | 59 | 2.954 | 0.274 | 50 | 2.990 | 0.317 | -0.035 | 0.045 | 0.4360 | 0.8622 |
| | CTN-067 M0 vs. CTN-067 M6 | 59 | 2.954 | 0.274 | 53 | 2.945 | 0.271 | 0.009 | 0.044 | 0.8352 | 0.9968 |
| | CTN-067 M3 vs. CTN-067 M6 | 50 | 2.990 | 0.317 | 53 | 2.945 | 0.271 | 0.044 | 0.045 | 0.3337 | 0.7657 |
| Log10 (IL37) | Control vs. CTN-067 M0 | 36 | 1.683 | 1.261 | 57 | 1.100 | 1.249 | 0.513 | 0.265 | 0.0567 | 0.2223 |
| | Control vs. CTN-067 M3 | 36 | 1.683 | 1.261 | 47 | 1.205 | 1.236 | 0.420 | 0.267 | 0.1202 | 0.4011 |
| | Control vs. CTN-067 M6 | 36 | 1.683 | 1.261 | 48 | 1.033 | 1.256 | 0.559 | 0.267 | 0.0393 | 0.1636 |
| | CTN-067 M0 vs. CTN-067 M3 | 57 | 1.100 | 1.249 | 47 | 1.205 | 1.236 | -0.093 | 0.114 | 0.4156 | 0.8457 |
| | CTN-067 M0 vs. CTN-067 M6 | 57 | 1.100 | 1.249 | 48 | 1.033 | 1.256 | 0.047 | 0.114 | 0.6840 | 0.9768 |

| | | | | | | | | | | | |
|---|---|---|---|---|---|---|---|---|---|---|---|
| | CTN-067 M3 vs. CTN-067 M6 | 47 | 1.205 | 1.236 | 48 | 1.033 | 1.256 | 0.140 | 0.118 | 0.2374 | 0.6347 |
| Log10 (LBP) | Control vs. CTN-067 M0 | 44 | 3.645 | 0.100 | 55 | 3.628 | 0.725 | 0.003 | 0.097 | 0.9719 | 1.0000 |
| | Control vs. CTN-067 M3 | 44 | 3.645 | 0.100 | 47 | 3.651 | 0.571 | -0.042 | 0.098 | 0.6673 | 0.9729 |
| | Control vs. CTN-067 M6 | 44 | 3.645 | 0.100 | 53 | 3.659 | 0.547 | -0.029 | 0.097 | 0.7656 | 0.9906 |
| | CTN-067 M0 vs. CTN-067 M3 | 55 | 3.628 | 0.725 | 47 | 3.651 | 0.571 | -0.046 | 0.056 | 0.4206 | 0.8499 |
| | CTN-067 M0 vs. CTN-067 M6 | 55 | 3.628 | 0.725 | 53 | 3.659 | 0.547 | -0.032 | 0.055 | 0.5554 | 0.9342 |
| | CTN-067 M3 vs. CTN-067 M6 | 47 | 3.651 | 0.571 | 53 | 3.659 | 0.547 | 0.013 | 0.057 | 0.8149 | 0.9954 |
| Log10 (sCD14) | Control vs. CTN-067 M0 | 44 | 5.949 | 0.091 | 59 | 6.128 | 0.109 | -0.177 | 0.022 | <.0001 | <.0001 |
| | Control vs. CTN-067 M3 | 44 | 5.949 | 0.091 | 50 | 6.121 | 0.117 | -0.178 | 0.022 | <.0001 | <.0001 |
| | Control vs. CTN-067 M6 | 44 | 5.949 | 0.091 | 54 | 6.092 | 0.119 | -0.144 | 0.022 | <.0001 | <.0001 |
| | CTN-067 M0 vs. CTN-067 M3 | 59 | 6.128 | 0.109 | 50 | 6.121 | 0.117 | 0.000 | 0.013 | 0.9882 | 1.0000 |
| | CTN-067 M0 vs. CTN-067 M6 | 59 | 6.128 | 0.109 | 54 | 6.092 | 0.119 | 0.033 | 0.013 | 0.0113 | 0.0542 |
| | CTN-067 M3 vs. CTN-067 M6 | 50 | 6.121 | 0.117 | 54 | 6.092 | 0.119 | 0.033 | 0.013 | 0.0130 | 0.0615 |
| | Control vs. CTN-067 M0 | 44 | 4.552 | 0.190 | 59 | 4.908 | 0.286 | -0.345 | 0.052 | <.0001 | <.0001 |

| | | | | | | | | | | | |
|---|---|---|---|---|---|---|---|---|---|---|---|
| Log10 (aCD163) | Control vs. CTN-067 M3 | 44 | 4.552 | 0.190 | 49 | 4.866 | 0.298 | -0.306 | 0.052 | <.0001 | <.0001 |
| | Control vs. CTN-067 M6 | 44 | 4.552 | 0.190 | 54 | 4.835 | 0.299 | -0.275 | 0.052 | <.0001 | <.0001 |
| | CTN-067 M0 vs. CTN-067 M3 | 59 | 4.908 | 0.286 | 49 | 4.866 | 0.298 | 0.039 | 0.026 | 0.1365 | 0.4404 |
| | CTN-067 M0 vs. CTN-067 M6 | 59 | 4.908 | 0.286 | 54 | 4.835 | 0.299 | 0.069 | 0.025 | 0.0070 | 0.0345 |
| | CTN-067 M3 vs. CTN-067 M6 | 49 | 4.866 | 0.298 | 54 | 4.835 | 0.299 | 0.031 | 0.026 | 0.2430 | 0.6439 |
