## Supplemental Table 3 for "Chronic opioid-associated immune dysregulation among people living with HIV"

**Supplementary Table 3.** Peripheral Blood Cytokine Results

| Cytokine | Comparision | Exp | | Ref | | Difference | | p-value | Adj. p-value |
|---|---|---|---|---|---|---|---|---|---|
| | | mean | SE | Mean | SE | Mean | SE | | |
| Fraktalkine | Control vs. CTN-067 M0 | 4891.550 | 411.090 | 3349.030 | 526.030 | 1542.520 | 746.140 | 0.0410 | 0.1703 |
| | Control vs. CTN-067 M3 | 5221.280 | 383.490 | 3349.030 | 411.090 | 1872.250 | 657.340 | 0.0052 | 0.0265 |
| | Control vs. CTN-067 M6 | 4940.900 | 378.160 | 3349.030 | 383.490 | 1591.860 | 626.770 | 0.0125 | 0.0593 |
| | CTN-067 M0 vs. CTN-067 M3 | 4891.550 | 411.090 | 5221.280 | 383.490 | -329.730 | 508.630 | 0.5181 | 0.9160 |
| | CTN-067 M0 vs. CTN-067 M6 | 4891.550 | 411.090 | 4940.900 | 378.160 | -49.346 | 560.890 | 0.9301 | 0.9998 |
| | CTN-067 M3 vs. CTN-067 M6 | 5221.280 | 383.490 | 4940.900 | 378.160 | 280.380 | 488.970 | 0.5675 | 0.9398 |
| GMCSF | Control vs. CTN-067 M0 | 0.248 | 0.139 | 0.061 | 0.195 | 0.187 | 0.264 | 0.4838 | 0.8941 |
| | Control vs. CTN-067 M3 | 0.384 | 0.128 | 0.061 | 0.139 | 0.323 | 0.235 | 0.1769 | 0.5230 |
| | Control vs. CTN-067 M6 | 0.233 | 0.148 | 0.061 | 0.128 | 0.171 | 0.233 | 0.4672 | 0.8831 |
| | CTN-067 M0 vs. CTN-067 M3 | 0.248 | 0.139 | 0.384 | 0.128 | -0.136 | 0.150 | 0.3685 | 0.8003 |
| | CTN-067 M0 vs. CTN-067 M6 | 0.248 | 0.139 | 0.233 | 0.148 | 0.016 | 0.195 | 0.9371 | 0.9998 |
| | CTN-067 M3 vs. CTN-067 M6 | 0.384 | 0.128 | 0.233 | 0.148 | 0.152 | 0.165 | 0.3631 | 0.7948 |
| ITAC | Control vs. CTN-067 M0 | 141.900 | 14.275 | 81.701 | 18.334 | 60.202 | 25.660 | 0.0208 | 0.0940 |
| | Control vs. CTN-067 M3 | 145.490 | 13.272 | 81.701 | 14.275 | 63.785 | 22.872 | 0.0062 | 0.0312 |
| | Control vs. CTN-067 M6 | 124.370 | 13.167 | 81.701 | 13.272 | 42.673 | 21.999 | 0.0550 | 0.2176 |
| | CTN-067 M0 vs. CTN-067 M3 | 141.900 | 14.275 | 145.490 | 13.272 | -3.582 | 14.985 | 0.8115 | 0.9952 |
| | CTN-067 M0 vs. CTN-067 M6 | 141.900 | 14.275 | 124.370 | 13.167 | 17.530 | 17.749 | 0.3255 | 0.7568 |
| | CTN-067 M3 vs. CTN-067 M6 | 145.490 | 13.272 | 124.370 | 13.167 | 21.112 | 14.219 | 0.1405 | 0.4501 |

| | | | | | | | | | |
|---|---|---|---|---|---|---|---|---|---|
| IFN-β | Control vs. CTN-067 M0 | 83.601 | 57.040 | 142.610 | 76.079 | -59.005 | 106.000 | 0.5816 | 0.9440 |
| | Control vs. CTN-067 M3 | 56.112 | 48.704 | 142.610 | 57.040 | -86.493 | 91.242 | 0.3503 | 0.7794 |
| | Control vs. CTN-067 M6 | 172.470 | 53.024 | 142.610 | 48.704 | 29.861 | 88.860 | 0.7390 | 0.9867 |
| | CTN-067 M0 vs. CTN-067 M3 | 83.601 | 57.040 | 56.112 | 48.704 | 27.488 | 69.067 | 0.6933 | 0.9783 |
| | CTN-067 M0 vs. CTN-067 M6 | 83.601 | 57.040 | 172.470 | 53.024 | -88.866 | 79.366 | 0.2712 | 0.6803 |
| | CTN-067 M3 vs. CTN-067 M6 | 56.112 | 48.704 | 172.470 | 53.024 | -116.350 | 66.010 | 0.0875 | 0.3000 |
| IFN-α | Control vs. CTN-067 M0 | 7.353 | 2.177 | 6.126 | 2.640 | 1.226 | 3.788 | 0.7470 | 0.9882 |
| | Control vs. CTN-067 M3 | 8.391 | 1.910 | 6.126 | 2.177 | 2.265 | 3.270 | 0.4904 | 0.8996 |
| | Control vs. CTN-067 M6 | 5.422 | 1.998 | 6.126 | 1.910 | -0.705 | 3.213 | 0.8269 | 0.9962 |
| | CTN-067 M0 vs. CTN-067 M3 | 7.353 | 2.177 | 8.391 | 1.910 | -1.039 | 2.948 | 0.7254 | 0.9849 |
| | CTN-067 M0 vs. CTN-067 M6 | 7.353 | 2.177 | 5.422 | 1.998 | 1.931 | 3.051 | 0.5285 | 0.9212 |
| | CTN-067 M3 vs. CTN-067 M6 | 8.391 | 1.910 | 5.422 | 1.998 | 2.970 | 2.836 | 0.2980 | 0.7223 |
| IFN-γ | Control vs. CTN-067 M0 | 106.160 | 36.837 | -2.676 | 53.796 | 108.830 | 71.850 | 0.1333 | 0.4330 |
| | Control vs. CTN-067 M3 | 115.650 | 37.207 | -2.676 | 36.837 | 118.320 | 65.082 | 0.0723 | 0.2716 |
| | Control vs. CTN-067 M6 | 76.455 | 35.288 | -2.676 | 37.207 | 79.132 | 62.275 | 0.2071 | 0.5838 |
| | CTN-067 M0 vs. CTN-067 M3 | 106.160 | 36.837 | 115.650 | 37.207 | -9.489 | 47.022 | 0.8405 | 0.9971 |
| | CTN-067 M0 vs. CTN-067 M6 | 106.160 | 36.837 | 76.455 | 35.288 | 29.703 | 50.469 | 0.5576 | 0.9353 |
| | CTN-067 M3 vs. CTN-067 M6 | 115.650 | 37.207 | 76.455 | 35.288 | 39.192 | 45.185 | 0.3880 | 0.8216 |
| IL-10 | Control vs. CTN-067 M0 | 3.491 | 1.834 | -0.254 | 2.670 | 3.745 | 3.610 | 0.3027 | 0.7281 |
| | Control vs. CTN-067 M3 | 5.710 | 1.866 | -0.254 | 1.834 | 5.964 | 3.250 | 0.0701 | 0.2645 |

| | | | | | | | | | |
|---|---|---|---|---|---|---|---|---|---|
| | Control vs. CTN-067 M6 | 1.554 | 1.773 | -0.254 | 1.866 | 1.808 | 3.094 | 0.5606 | 0.9365 |
| | CTN-067 M0 vs. CTN-067 M3 | 3.491 | 1.834 | 5.710 | 1.866 | -2.219 | 2.609 | 0.3975 | 0.8300 |
| | CTN-067 M0 vs. CTN-067 M6 | 3.491 | 1.834 | 1.554 | 1.773 | 1.937 | 2.636 | 0.4646 | 0.8828 |
| | CTN-067 M3 vs. CTN-067 M6 | 5.710 | 1.866 | 1.554 | 1.773 | 4.156 | 2.557 | 0.1079 | 0.3703 |
| IL-12p70 | Control vs. CTN-067 M0 | 3.749 | 1.340 | 0.949 | 1.732 | 2.800 | 2.445 | 0.2567 | 0.6634 |
| | Control vs. CTN-067 M3 | 1.639 | 1.209 | 0.949 | 1.340 | 0.690 | 2.158 | 0.7503 | 0.9886 |
| | Control vs. CTN-067 M6 | 1.433 | 1.284 | 0.949 | 1.209 | 0.484 | 2.076 | 0.8164 | 0.9955 |
| | CTN-067 M0 vs. CTN-067 M3 | 3.749 | 1.340 | 1.639 | 1.209 | 2.110 | 1.450 | 0.1508 | 0.4707 |
| | CTN-067 M0 vs. CTN-067 M6 | 3.749 | 1.340 | 1.433 | 1.284 | 2.316 | 1.781 | 0.1984 | 0.5664 |
| | CTN-067 M3 vs. CTN-067 M6 | 1.639 | 1.209 | 1.433 | 1.284 | 0.206 | 1.516 | 0.8926 | 0.9991 |
| IL-15 | Control vs. CTN-067 M0 | 7.653 | 1.949 | 5.030 | 2.476 | 2.624 | 3.518 | 0.4574 | 0.8783 |
| | Control vs. CTN-067 M3 | 8.219 | 1.824 | 5.030 | 1.949 | 3.190 | 3.081 | 0.3028 | 0.7291 |
| | Control vs. CTN-067 M6 | 7.611 | 1.810 | 5.030 | 1.824 | 2.582 | 2.968 | 0.3863 | 0.8204 |
| | CTN-067 M0 vs. CTN-067 M3 | 7.653 | 1.949 | 8.219 | 1.824 | -0.566 | 2.695 | 0.8341 | 0.9967 |
| | CTN-067 M0 vs. CTN-067 M6 | 7.653 | 1.949 | 7.611 | 1.810 | 0.042 | 2.751 | 0.9878 | 1.0000 |
| | CTN-067 M3 vs. CTN-067 M6 | 8.219 | 1.824 | 7.611 | 1.810 | 0.608 | 2.603 | 0.8158 | 0.9955 |
| IL-17A | Control vs. CTN-067 M0 | 66.125 | 25.556 | 89.227 | 39.134 | -23.102 | 51.756 | 0.6568 | 0.9701 |
| | Control vs. CTN-067 M3 | 44.181 | 24.737 | 89.227 | 25.556 | -45.046 | 45.938 | 0.3303 | 0.7609 |
| | Control vs. CTN-067 M6 | 97.974 | 24.848 | 89.227 | 24.737 | 8.747 | 44.516 | 0.8448 | 0.9973 |
| | CTN-067 M0 vs. CTN-067 M3 | 66.125 | 25.556 | 44.181 | 24.737 | 21.944 | 34.921 | 0.5319 | 0.9226 |

| | | | | | | | | | |
|---|---|---|---|---|---|---|---|---|---|
| | CTN-067 M0 vs. CTN-067 M6 | 66.125 | 25.556 | 97.974 | 24.848 | -31.849 | 36.906 | 0.3912 | 0.8238 |
| | CTN-067 M3 vs. CTN-067 M6 | 44.181 | 24.737 | 97.974 | 24.848 | -53.793 | 33.956 | 0.1178 | 0.3944 |
| IL-18 | Control vs. CTN-067 M0 | 680.390 | 51.707 | 633.300 | 67.218 | 47.088 | 92.033 | 0.6099 | 0.9562 |
| | Control vs. CTN-067 M3 | 699.910 | 48.870 | 633.300 | 51.707 | 66.608 | 83.718 | 0.4279 | 0.8563 |
| | Control vs. CTN-067 M6 | 668.920 | 48.712 | 633.300 | 48.870 | 35.612 | 81.542 | 0.6632 | 0.9720 |
| | CTN-067 M0 vs. CTN-067 M3 | 680.390 | 51.707 | 699.910 | 48.870 | -19.521 | 47.051 | 0.6790 | 0.9758 |
| | CTN-067 M0 vs. CTN-067 M6 | 680.390 | 51.707 | 668.920 | 48.712 | 11.476 | 57.295 | 0.8416 | 0.9971 |
| | CTN-067 M3 vs. CTN-067 M6 | 699.910 | 48.870 | 668.920 | 48.712 | 30.996 | 43.900 | 0.4816 | 0.8945 |
| IL-1β | Control vs. CTN-067 M0 | 17.764 | 32.117 | -9.375 | 48.966 | 27.138 | 64.693 | 0.6759 | 0.9750 |
| | Control vs. CTN-067 M3 | 66.667 | 31.743 | -9.375 | 32.117 | 76.041 | 58.387 | 0.1962 | 0.5639 |
| | Control vs. CTN-067 M6 | 26.542 | 31.184 | -9.375 | 31.743 | 35.916 | 56.057 | 0.5234 | 0.9185 |
| | CTN-067 M0 vs. CTN-067 M3 | 17.764 | 32.117 | 66.667 | 31.743 | -48.903 | 44.879 | 0.2788 | 0.6967 |
| | CTN-067 M0 vs. CTN-067 M6 | 17.764 | 32.117 | 26.542 | 31.184 | -8.778 | 46.230 | 0.8498 | 0.9976 |
| | CTN-067 M3 vs. CTN-067 M6 | 66.667 | 31.743 | 26.542 | 31.184 | 40.125 | 44.266 | 0.3672 | 0.8014 |
| IL-2 | Control vs. CTN-067 M0 | 5.110 | 1.507 | 0.819 | 7.168 | 4.290 | 7.374 | 0.5683 | 0.9362 |
| | Control vs. CTN-067 M3 | 2.488 | 1.574 | 0.819 | 1.507 | 1.668 | 7.320 | 0.8224 | 0.9957 |
| | Control vs. CTN-067 M6 | 3.064 | 1.709 | 0.819 | 1.574 | 2.245 | 7.327 | 0.7630 | 0.9897 |
| | CTN-067 M0 vs. CTN-067 M3 | 5.110 | 1.507 | 2.488 | 1.574 | 2.622 | 2.241 | 0.2581 | 0.6529 |
| | CTN-067 M0 vs. CTN-067 M6 | 5.110 | 1.507 | 3.064 | 1.709 | 2.045 | 2.396 | 0.4051 | 0.8281 |
| | CTN-067 M3 vs. CTN-067 M6 | 2.488 | 1.574 | 3.064 | 1.709 | -0.577 | 2.282 | 0.8035 | 0.9941 |

| | | | | | | | | | |
|---|---|---|---|---|---|---|---|---|---|
| IL-21 | Control vs. CTN-067 M0 | 215.950 | 87.512 | 146.230 | 106.860 | 69.722 | 152.800 | 0.6497 | 0.9682 |
| | Control vs. CTN-067 M3 | 227.320 | 75.025 | 146.230 | 87.512 | 81.085 | 130.720 | 0.5372 | 0.9252 |
| | Control vs. CTN-067 M6 | 160.830 | 86.591 | 146.230 | 75.025 | 14.595 | 132.470 | 0.9126 | 0.9995 |
| | CTN-067 M0 vs. CTN-067 M3 | 215.950 | 87.512 | 227.320 | 75.025 | -11.362 | 117.760 | 0.9234 | 0.9997 |
| | CTN-067 M0 vs. CTN-067 M6 | 215.950 | 87.512 | 160.830 | 86.591 | 55.128 | 128.140 | 0.6684 | 0.9731 |
| | CTN-067 M3 vs. CTN-067 M6 | 227.320 | 75.025 | 160.830 | 86.591 | 66.490 | 117.620 | 0.5738 | 0.9420 |
| IL-22 | Control vs. CTN-067 M0 | 8.138 | 4.223 | 1.249 | 5.453 | 6.889 | 7.674 | 0.3714 | 0.8060 |
| | Control vs. CTN-067 M3 | 13.567 | 3.943 | 1.249 | 4.223 | 12.318 | 6.735 | 0.0703 | 0.2658 |
| | Control vs. CTN-067 M6 | 2.705 | 4.014 | 1.249 | 3.943 | 1.456 | 6.549 | 0.8245 | 0.9961 |
| | CTN-067 M0 vs. CTN-067 M3 | 8.138 | 4.223 | 13.567 | 3.943 | -5.429 | 5.804 | 0.3518 | 0.7859 |
| | CTN-067 M0 vs. CTN-067 M6 | 8.138 | 4.223 | 2.705 | 4.014 | 5.433 | 6.023 | 0.3692 | 0.8038 |
| | CTN-067 M3 vs. CTN-067 M6 | 13.567 | 3.943 | 2.705 | 4.014 | 10.861 | 5.661 | 0.0578 | 0.2267 |
| IL-23 | Control vs. CTN-067 M0 | 7.716 | 3.464 | 1.737 | 6.043 | 5.979 | 7.377 | 0.4199 | 0.8493 |
| | Control vs. CTN-067 M3 | 6.325 | 3.272 | 1.737 | 3.464 | 4.589 | 6.845 | 0.5044 | 0.9080 |
| | Control vs. CTN-067 M6 | 5.505 | 3.402 | 1.737 | 3.272 | 3.769 | 6.724 | 0.5766 | 0.9434 |
| | CTN-067 M0 vs. CTN-067 M3 | 7.716 | 3.464 | 6.325 | 3.272 | 1.391 | 3.449 | 0.6878 | 0.9777 |
| | CTN-067 M0 vs. CTN-067 M6 | 7.716 | 3.464 | 5.505 | 3.402 | 2.210 | 4.310 | 0.6094 | 0.9558 |
| | CTN-067 M3 vs. CTN-067 M6 | 6.325 | 3.272 | 5.505 | 3.402 | 0.820 | 3.281 | 0.8033 | 0.9945 |
| IL-27 | Control vs. CTN-067 M0 | 267.970 | 60.898 | 259.290 | 76.926 | 8.685 | 109.750 | 0.9371 | 0.9998 |
| | Control vs. CTN-067 M3 | 325.110 | 56.383 | 259.290 | 60.898 | 65.823 | 95.704 | 0.4931 | 0.9016 |

| | | | | | | | | | |
|---|---|---|---|---|---|---|---|---|---|
| | Control vs. CTN-067 M6 | 244.630 | 55.908 | 259.290 | 56.383 | -14.660 | 92.149 | 0.8739 | 0.9986 |
| | CTN-067 M0 vs. CTN-067 M3 | 267.970 | 60.898 | 325.110 | 56.383 | -57.138 | 80.223 | 0.4779 | 0.8920 |
| | CTN-067 M0 vs. CTN-067 M6 | 267.970 | 60.898 | 244.630 | 55.908 | 23.345 | 84.639 | 0.7832 | 0.9926 |
| | CTN-067 M3 vs. CTN-067 M6 | 325.110 | 56.383 | 244.630 | 55.908 | 80.483 | 76.768 | 0.2968 | 0.7214 |
| IL-29 | Control vs. CTN-067 M0 | 17.820 | 5.382 | 7.756 | 7.157 | 10.064 | 9.931 | 0.3134 | 0.7420 |
| | Control vs. CTN-067 M3 | 21.689 | 5.013 | 7.756 | 5.382 | 13.933 | 8.785 | 0.1160 | 0.3914 |
| | Control vs. CTN-067 M6 | 13.761 | 5.202 | 7.756 | 5.013 | 6.004 | 8.593 | 0.4864 | 0.8973 |
| | CTN-067 M0 vs. CTN-067 M3 | 17.820 | 5.382 | 21.689 | 5.013 | -3.869 | 6.211 | 0.5348 | 0.9245 |
| | CTN-067 M0 vs. CTN-067 M6 | 17.820 | 5.382 | 13.761 | 5.202 | 4.060 | 7.192 | 0.5738 | 0.9423 |
| | CTN-067 M3 vs. CTN-067 M6 | 21.689 | 5.013 | 13.761 | 5.202 | 7.928 | 6.019 | 0.1909 | 0.5543 |
| IL-4 | Control vs. CTN-067 M0 | 3.355 | 1.152 | 4.495 | 2.152 | -1.140 | 2.666 | 0.6703 | 0.9736 |
| | Control vs. CTN-067 M3 | 2.675 | 1.152 | 4.495 | 1.152 | -1.820 | 2.393 | 0.4494 | 0.8717 |
| | Control vs. CTN-067 M6 | 5.572 | 1.148 | 4.495 | 1.152 | 1.077 | 2.343 | 0.6475 | 0.9676 |
| | CTN-067 M0 vs. CTN-067 M3 | 3.355 | 1.152 | 2.675 | 1.152 | 0.681 | 1.575 | 0.6672 | 0.9728 |
| | CTN-067 M0 vs. CTN-067 M6 | 3.355 | 1.152 | 5.572 | 1.148 | -2.216 | 1.676 | 0.1906 | 0.5523 |
| | CTN-067 M3 vs. CTN-067 M6 | 2.675 | 1.152 | 5.572 | 1.148 | -2.897 | 1.496 | 0.0571 | 0.2230 |
| IL-6 | Control vs. CTN-067 M0 | 23.695 | 32.235 | -5.382 | 41.912 | 29.077 | 59.056 | 0.6235 | 0.9606 |
| | Control vs. CTN-067 M3 | 66.814 | 30.289 | -5.382 | 32.235 | 72.196 | 52.056 | 0.1684 | 0.5103 |
| | Control vs. CTN-067 M6 | 35.031 | 30.554 | -5.382 | 30.289 | 40.413 | 50.128 | 0.4219 | 0.8514 |
| | CTN-067 M0 vs. CTN-067 M3 | 23.695 | 32.235 | 66.814 | 30.289 | -43.120 | 43.883 | 0.3281 | 0.7596 |

| | | | | | | | | | |
|---|---|---|---|---|---|---|---|---|---|
| | CTN-067 M0 vs. CTN-067 M6 | 23.695 | 32.235 | 35.031 | 30.554 | -11.337 | 45.856 | 0.8052 | 0.9947 |
| | CTN-067 M3 vs. CTN-067 M6 | 66.814 | 30.289 | 35.031 | 30.554 | 31.783 | 43.050 | 0.4620 | 0.8814 |
| IL-7 | Control vs. CTN-067 M0 | 18.917 | 4.671 | 15.897 | 5.917 | 3.020 | 8.423 | 0.7206 | 0.9841 |
| | Control vs. CTN-067 M3 | 21.178 | 4.416 | 15.897 | 4.671 | 5.281 | 7.432 | 0.4789 | 0.8927 |
| | Control vs. CTN-067 M6 | 18.644 | 4.312 | 15.897 | 4.416 | 2.747 | 7.065 | 0.6982 | 0.9799 |
| | CTN-067 M0 vs. CTN-067 M3 | 18.917 | 4.671 | 21.178 | 4.416 | -2.261 | 6.544 | 0.7305 | 0.9858 |
| | CTN-067 M0 vs. CTN-067 M6 | 18.917 | 4.671 | 18.644 | 4.312 | 0.273 | 6.599 | 0.9671 | 1.0000 |
| | CTN-067 M3 vs. CTN-067 M6 | 21.178 | 4.416 | 18.644 | 4.312 | 2.534 | 6.357 | 0.6910 | 0.9784 |
| IL-8 | Control vs. CTN-067 M0 | 1363.380 | 248.400 | 92.246 | 315.420 | 1271.130 | 448.910 | 0.0055 | 0.0278 |
| | Control vs. CTN-067 M3 | 621.730 | 229.830 | 92.246 | 248.400 | 529.480 | 394.210 | 0.1820 | 0.5376 |
| | Control vs. CTN-067 M6 | 978.570 | 226.700 | 92.246 | 229.830 | 886.330 | 375.740 | 0.0201 | 0.0913 |
| | CTN-067 M0 vs. CTN-067 M3 | 1363.380 | 248.400 | 621.730 | 229.830 | 741.650 | 304.550 | 0.0165 | 0.0764 |
| | CTN-067 M0 vs. CTN-067 M6 | 1363.380 | 248.400 | 978.570 | 226.700 | 384.800 | 337.540 | 0.2567 | 0.6656 |
| | CTN-067 M3 vs. CTN-067 M6 | 621.730 | 229.830 | 978.570 | 226.700 | -356.840 | 291.690 | 0.2238 | 0.6134 |
| IL-9 | Control vs. CTN-067 M0 | 4.528 | 1.369 | 2.413 | 1.971 | 2.115 | 2.622 | 0.4223 | 0.8513 |
| | Control vs. CTN-067 M3 | 3.320 | 1.321 | 2.413 | 1.369 | 0.907 | 2.370 | 0.7030 | 0.9808 |
| | Control vs. CTN-067 M6 | 5.358 | 1.312 | 2.413 | 1.321 | 2.945 | 2.291 | 0.2021 | 0.5747 |
| | CTN-067 M0 vs. CTN-067 M3 | 4.528 | 1.369 | 3.320 | 1.321 | 1.208 | 1.958 | 0.5390 | 0.9265 |
| | CTN-067 M0 vs. CTN-067 M6 | 4.528 | 1.369 | 5.358 | 1.312 | -0.830 | 1.967 | 0.6739 | 0.9745 |
| | CTN-067 M3 vs. CTN-067 M6 | 3.320 | 1.321 | 5.358 | 1.312 | -2.038 | 1.918 | 0.2909 | 0.7130 |

| | | | | | | | | | |
|---|---|---|---|---|---|---|---|---|---|
| IP-10 | Control vs. CTN-067 M0 | 2090.740 | 267.370 | 1717.540 | 339.220 | 373.200 | 483.130 | 0.4415 | 0.8667 |
| | Control vs. CTN-067 M3 | 2732.240 | 247.390 | 1717.540 | 267.370 | 1014.700 | 424.010 | 0.0184 | 0.0844 |
| | Control vs. CTN-067 M6 | 2031.570 | 243.950 | 1717.540 | 247.390 | 314.040 | 404.020 | 0.4387 | 0.8646 |
| | CTN-067 M0 vs. CTN-067 M3 | 2090.740 | 267.370 | 2732.240 | 247.390 | -641.500 | 331.250 | 0.0554 | 0.2189 |
| | CTN-067 M0 vs. CTN-067 M6 | 2090.740 | 267.370 | 2031.570 | 243.950 | 59.165 | 364.920 | 0.8715 | 0.9985 |
| | CTN-067 M3 vs. CTN-067 M6 | 2732.240 | 247.390 | 2031.570 | 243.950 | 700.670 | 317.490 | 0.0294 | 0.1278 |
| MCP-3 | Control vs. CTN-067 M0 | 133.090 | 49.394 | 67.010 | 63.118 | 66.081 | 89.666 | 0.4628 | 0.8820 |
| | Control vs. CTN-067 M3 | 48.424 | 46.067 | 67.010 | 49.394 | -18.586 | 78.464 | 0.8132 | 0.9953 |
| | Control vs. CTN-067 M6 | 135.180 | 45.755 | 67.010 | 46.067 | 68.174 | 75.285 | 0.3672 | 0.8019 |
| | CTN-067 M0 vs. CTN-067 M3 | 133.090 | 49.394 | 48.424 | 46.067 | 84.667 | 64.733 | 0.1937 | 0.5600 |
| | CTN-067 M0 vs. CTN-067 M6 | 133.090 | 49.394 | 135.180 | 45.755 | -2.094 | 69.042 | 0.9759 | 1.0000 |
| | CTN-067 M3 vs. CTN-067 M6 | 48.424 | 46.067 | 135.180 | 45.755 | -86.760 | 62.278 | 0.1665 | 0.5065 |
| MIP-1α | Control vs. CTN-067 M0 | 104.880 | 44.601 | 12.197 | 59.137 | 92.679 | 82.270 | 0.2625 | 0.6740 |
| | Control vs. CTN-067 M3 | 98.904 | 41.312 | 12.197 | 44.601 | 86.706 | 72.441 | 0.2340 | 0.6301 |
| | Control vs. CTN-067 M6 | 149.190 | 40.514 | 12.197 | 41.312 | 136.990 | 68.975 | 0.0496 | 0.1997 |
| | CTN-067 M0 vs. CTN-067 M3 | 104.880 | 44.601 | 98.904 | 41.312 | 5.973 | 59.339 | 0.9200 | 0.9996 |
| | CTN-067 M0 vs. CTN-067 M6 | 104.880 | 44.601 | 149.190 | 40.514 | -44.314 | 62.394 | 0.4791 | 0.8928 |
| | CTN-067 M3 vs. CTN-067 M6 | 98.904 | 41.312 | 149.190 | 40.514 | -50.287 | 56.710 | 0.3772 | 0.8117 |
| MIP-3α | Control vs. CTN-067 M0 | 60.598 | 16.742 | 16.011 | 21.566 | 44.586 | 30.425 | 0.1458 | 0.4619 |
| | Control vs. CTN-067 M3 | 88.699 | 15.571 | 16.011 | 16.742 | 72.688 | 26.581 | 0.0073 | 0.0363 |

| | | | | | | | | | |
|---|---|---|---|---|---|---|---|---|---|
| | Control vs. CTN-067 M6 | 36.359 | 15.448 | 16.011 | 15.571 | 20.348 | 25.689 | 0.4301 | 0.8579 |
| | CTN-067 M0 vs. CTN-067 M3 | 60.598 | 16.742 | 88.699 | 15.571 | -28.102 | 22.765 | 0.2198 | 0.6065 |
| | CTN-067 M0 vs. CTN-067 M6 | 60.598 | 16.742 | 36.359 | 15.448 | 24.238 | 23.495 | 0.3046 | 0.7313 |
| | CTN-067 M3 vs. CTN-067 M6 | 88.699 | 15.571 | 36.359 | 15.448 | 52.340 | 21.810 | 0.0181 | 0.0833 |
| SDF-1α | Control vs. CTN-067 M0 | 503.950 | 51.919 | 371.460 | 68.944 | 132.500 | 94.246 | 0.1628 | 0.4986 |
| | Control vs. CTN-067 M3 | 550.510 | 48.794 | 371.460 | 51.919 | 179.060 | 84.822 | 0.0372 | 0.1565 |
| | Control vs. CTN-067 M6 | 433.300 | 48.080 | 371.460 | 48.794 | 61.839 | 82.073 | 0.4529 | 0.8750 |
| | CTN-067 M0 vs. CTN-067 M3 | 503.950 | 51.919 | 550.510 | 48.794 | -46.560 | 51.417 | 0.3673 | 0.8019 |
| | CTN-067 M0 vs. CTN-067 M6 | 503.950 | 51.919 | 433.300 | 48.080 | 70.658 | 60.939 | 0.2489 | 0.6536 |
| | CTN-067 M3 vs. CTN-067 M6 | 550.510 | 48.794 | 433.300 | 48.080 | 117.220 | 47.546 | 0.0153 | 0.0715 |
| TGF-β1 | Control vs. CTN-067 M0 | 16372.00 | 934.25 | 20644.00 | 1191.85 | -4271.63 | 1686.36 | 0.0127 | 0.0604 |
| | Control vs. CTN-067 M3 | 16648.00 | 866.73 | 20644.00 | 934.25 | -3995.53 | 1489.86 | 0.0085 | 0.0415 |
| | Control vs. CTN-067 M6 | 16889.00 | 862.42 | 20644.00 | 866.73 | -3754.72 | 1427.46 | 0.0098 | 0.0473 |
| | CTN-067 M0 vs. CTN-067 M3 | 16372.00 | 934.25 | 16648.00 | 866.73 | -276.10 | 1083.32 | 0.7993 | 0.9942 |
| | CTN-067 M0 vs. CTN-067 M6 | 16372.00 | 934.25 | 16889.00 | 862.42 | -516.91 | 1235.11 | 0.6764 | 0.9752 |
| | CTN-067 M3 vs. CTN-067 M6 | 16648.00 | 866.73 | 16889.00 | 862.42 | -240.81 | 1034.96 | 0.8164 | 0.9955 |
| TGF-β2 | Control vs. CTN-067 M0 | 58.062 | 4.742 | 64.311 | 6.169 | -6.250 | 8.283 | 0.4522 | 0.8746 |
| | Control vs. CTN-067 M3 | 60.888 | 4.527 | 64.311 | 4.742 | -3.424 | 7.711 | 0.6579 | 0.9706 |
| | Control vs. CTN-067 M6 | 64.339 | 4.555 | 64.311 | 4.527 | 0.028 | 7.564 | 0.9971 | 1.0000 |
| | CTN-067 M0 vs. CTN-067 M3 | 58.062 | 4.742 | 60.888 | 4.527 | -2.826 | 3.573 | 0.4307 | 0.8584 |

| | | | | | | | | | |
|---|---|---|---|---|---|---|---|---|---|
| | CTN-067 M0 vs. CTN-067 M6 | 58.062 | 4.742 | 64.339 | 4.555 | -6.278 | 4.512 | 0.1670 | 0.5075 |
| | CTN-067 M3 vs. CTN-067 M6 | 60.888 | 4.527 | 64.339 | 4.555 | -3.452 | 3.363 | 0.3070 | 0.7345 |
| TGF-β3 | Control vs. CTN-067 M0 | 0.509 | 0.054 | 0.455 | 0.075 | 0.054 | 0.103 | 0.5983 | 0.9518 |
| | Control vs. CTN-067 M3 | 0.509 | 0.048 | 0.455 | 0.054 | 0.054 | 0.090 | 0.5486 | 0.9311 |
| | Control vs. CTN-067 M6 | 0.502 | 0.049 | 0.455 | 0.048 | 0.047 | 0.086 | 0.5829 | 0.9460 |
| | CTN-067 M0 vs. CTN-067 M3 | 0.509 | 0.054 | 0.509 | 0.048 | 0.000 | 0.066 | 0.9954 | 1.0000 |
| | CTN-067 M0 vs. CTN-067 M6 | 0.509 | 0.054 | 0.502 | 0.049 | 0.007 | 0.075 | 0.9217 | 0.9997 |
| | CTN-067 M3 vs. CTN-067 M6 | 0.509 | 0.048 | 0.502 | 0.049 | 0.007 | 0.064 | 0.9132 | 0.9995 |
| TNF-α | Control vs. CTN-067 M0 | 25.047 | 5.680 | 11.434 | 7.657 | 13.613 | 10.616 | 0.2028 | 0.5763 |
| | Control vs. CTN-067 M3 | 18.217 | 5.570 | 11.434 | 5.680 | 6.783 | 9.539 | 0.4787 | 0.8925 |
| | Control vs. CTN-067 M6 | 23.263 | 5.337 | 11.434 | 5.570 | 11.829 | 8.995 | 0.1916 | 0.5557 |
| | CTN-067 M0 vs. CTN-067 M3 | 25.047 | 5.680 | 18.217 | 5.570 | 6.830 | 7.788 | 0.3826 | 0.8168 |
| | CTN-067 M0 vs. CTN-067 M6 | 25.047 | 5.680 | 23.263 | 5.337 | 1.784 | 8.051 | 0.8251 | 0.9961 |
| | CTN-067 M3 vs. CTN-067 M6 | 18.217 | 5.570 | 23.263 | 5.337 | -5.046 | 7.617 | 0.5093 | 0.9109 |
| IL-33 | Control vs. CTN-067 M0 | 1.617 | 0.453 | 1.582 | 0.539 | 0.035 | 0.784 | 0.9646 | 1.0000 |
| | Control vs. CTN-067 M3 | 0.988 | 0.396 | 1.582 | 0.453 | -0.593 | 0.675 | 0.3819 | 0.8154 |
| | Control vs. CTN-067 M6 | 1.483 | 0.427 | 1.582 | 0.396 | -0.098 | 0.671 | 0.8838 | 0.9989 |
| | CTN-067 M0 vs. CTN-067 M3 | 1.617 | 0.453 | 0.988 | 0.396 | 0.628 | 0.574 | 0.2774 | 0.6942 |
| | CTN-067 M0 vs. CTN-067 M6 | 1.617 | 0.453 | 1.483 | 0.427 | 0.133 | 0.632 | 0.8337 | 0.9967 |
| | CTN-067 M3 vs. CTN-067 M6 | 0.988 | 0.396 | 1.483 | 0.427 | -0.495 | 0.558 | 0.3775 | 0.8112 |
