## Supplemental Table 4 for "Chronic opioid-associated immune dysregulation among people living with HIV"

**Supplementary Table 4.** CD14+ flow cytometry results

| Marker | Comparison (A vs B) | Group A | | | Group B | | | Viral Load adjusted Group A – Group B | | | |
|---|---|---|---|---|---|---|---|---|---|---|---|
| | | n | Mean | SD | N | Mean | SD | Estimate | SE | p | adj p |
| CD14+ \| Geometric Mean (APC-Fire 750-A :: CD36) | Control vs. CTN-067 M0 | 8 | 138733.88 | 53563.65 | 11 | 116541.64 | 42389.29 | 37864.00 | 25882.00 | 0.1598 | 0.4780 |
| | Control vs. CTN-067 M3 | 8 | 138733.88 | 53563.65 | 11 | 135080.36 | 43300.14 | 19582.00 | 23222.00 | 0.4096 | 0.8332 |
| | Control vs. CTN-067 M6 | 8 | 138733.88 | 53563.65 | 11 | 138790.73 | 40138.75 | 15884.00 | 23122.00 | 0.5004 | 0.9008 |
| | CTN-067 M0 vs. CTN-067 M3 | 11 | 116541.64 | 42389.29 | 11 | 135080.36 | 43300.14 | -18282.00 | 12421.00 | 0.1574 | 0.4728 |
| | CTN-067 M0 vs. CTN-067 M6 | 11 | 116541.64 | 42389.29 | 11 | 138790.73 | 40138.75 | -21980.00 | 12567.00 | 0.0964 | 0.3273 |
| | CTN-067 M3 vs. CTN-067 M6 | 11 | 135080.36 | 43300.14 | 11 | 138790.73 | 40138.75 | -3697.62 | 10891.00 | 0.7379 | 0.9861 |
| CD14+/CD36+ \| Freq. of Parent | Control vs. CTN-067 M0 | 8 | 97.21 | 1.96 | 11 | 94.01 | 3.88 | 37864.00 | 25882.00 | 0.1598 | 0.4780 |
| | Control vs. CTN-067 M3 | 8 | 97.21 | 1.96 | 11 | 94.53 | 2.66 | 19582.00 | 23222.00 | 0.4096 | 0.8332 |
| | Control vs. CTN-067 M6 | 8 | 97.21 | 1.96 | 11 | 95.58 | 1.82 | 15884.00 | 23122.00 | 0.5004 | 0.9008 |
| | CTN-067 M0 vs. CTN-067 M3 | 11 | 94.01 | 3.88 | 11 | 94.53 | 2.66 | -18282.00 | 12421.00 | 0.1574 | 0.4728 |
| | CTN-067 M0 vs. CTN-067 M6 | 11 | 94.01 | 3.88 | 11 | 95.58 | 1.82 | -21980.00 | 12567.00 | 0.0964 | 0.3273 |
| | CTN-067 M3 vs. CTN-067 M6 | 11 | 94.53 | 2.66 | 11 | 95.58 | 1.82 | -3697.62 | 10891.00 | 0.7379 | 0.9861 |
| CD14+ \| Geometric Mean (Alexa Fluor 647-A :: GLUT1) | Control vs. CTN-067 M0 | 8 | 395.10 | 378.46 | 11 | 396.89 | 460.73 | 202.89 | 233.68 | 0.3961 | 0.8210 |
| | Control vs. CTN-067 M3 | 8 | 395.10 | 378.46 | 11 | 268.77 | 399.98 | 270.01 | 218.24 | 0.2311 | 0.6118 |
| | Control vs. CTN-067 M6 | 8 | 395.10 | 378.46 | 11 | 286.05 | 401.19 | 249.70 | 217.67 | 0.2656 | 0.6659 |
| | CTN-067 M0 vs. CTN-067 M3 | 11 | 396.89 | 460.73 | 11 | 268.77 | 399.98 | 67.12 | 88.93 | 0.4596 | 0.8735 |

| | | | | | | | | | | |
|---|---|---|---|---|---|---|---|---|---|---|
| | CTN-067 M0 vs. CTN-067 M6 | 11 | 396.89 | 460.73 | 11 | 286.05 | 401.19 | 46.82 | 90.01 | 0.6090 | 0.9532 |
| | CTN-067 M3 vs. CTN-067 M6 | 11 | 268.77 | 399.98 | 11 | 286.05 | 401.19 | -20.31 | 77.47 | 0.7961 | 0.9935 |
| CD14+/GLUT1+ \| Freq. of Parent | Control vs. CTN-067 M0 | 8 | 2.05 | 1.54 | 11 | 3.94 | 4.97 | -4.29 | 2.02 | 0.0467 | 0.1802 |
| | Control vs. CTN-067 M3 | 8 | 2.05 | 1.54 | 11 | 2.26 | 2.18 | -1.41 | 1.69 | 0.4143 | 0.8374 |
| | Control vs. CTN-067 M6 | 8 | 2.05 | 1.54 | 11 | 2.43 | 2.72 | -1.52 | 1.68 | 0.3760 | 0.8016 |
| | CTN-067 M0 vs. CTN-067 M3 | 11 | 3.94 | 4.97 | 11 | 2.26 | 2.18 | 2.89 | 1.37 | 0.0485 | 0.1861 |
| | CTN-067 M0 vs. CTN-067 M6 | 11 | 3.94 | 4.97 | 11 | 2.43 | 2.72 | 2.78 | 1.38 | 0.0588 | 0.2191 |
| | CTN-067 M3 vs. CTN-067 M6 | 11 | 2.26 | 2.18 | 11 | 2.43 | 2.72 | -0.11 | 1.24 | 0.9304 | 0.9997 |
| CD14+ \| Geometric Mean (BUV615-A :: CD11b) | Control vs. CTN-067 M0 | 8 | 102672.38 | 14562.71 | 11 | 73878.82 | 28529.75 | 25244.00 | 15208.00 | 0.1133 | 0.3710 |
| | Control vs. CTN-067 M3 | 8 | 102672.38 | 14562.71 | 11 | 82174.73 | 34147.32 | 22591.00 | 13258.00 | 0.1047 | 0.3490 |
| | Control vs. CTN-067 M6 | 8 | 102672.38 | 14562.71 | 11 | 88322.64 | 26871.63 | 16724.00 | 13183.00 | 0.2199 | 0.5930 |
| | CTN-067 M0 vs. CTN-067 M3 | 11 | 73878.82 | 28529.75 | 11 | 82174.73 | 34147.32 | -2653.45 | 8294.53 | 0.7525 | 0.9883 |
| | CTN-067 M0 vs. CTN-067 M6 | 11 | 73878.82 | 28529.75 | 11 | 88322.64 | 26871.63 | -8520.83 | 8387.35 | 0.3224 | 0.7423 |
| | CTN-067 M3 vs. CTN-067 M6 | 11 | 82174.73 | 34147.32 | 11 | 88322.64 | 26871.63 | -5867.38 | 7324.06 | 0.4330 | 0.8530 |
| CD14+/CD11b+ \| Freq. of Parent | Control vs. CTN-067 M0 | 8 | 94.93 | 2.34 | 11 | 88.97 | 6.74 | 7.80 | 2.65 | 0.0084 | 0.0383 |
| | Control vs. CTN-067 M3 | 8 | 94.93 | 2.34 | 11 | 90.44 | 4.12 | 5.99 | 2.31 | 0.0179 | 0.0772 |
| | Control vs. CTN-067 M6 | 8 | 94.93 | 2.34 | 11 | 92.78 | 2.79 | 3.62 | 2.30 | 0.1311 | 0.4139 |
| | CTN-067 M0 vs. | 11 | 88.97 | 6.74 | 11 | 90.44 | 4.12 | -1.81 | 1.44 | 0.2258 | 0.6029 |

| | | | | | | | | | | | |
|---|---|---|---|---|---|---|---|---|---|---|---|
| | CTN-067 M3 | | | | | | | | | | |
| | CTN-067 M0 vs. CTN-067 M6 | 1 1 | 88.97 | 6.74 | 1 1 | 92.78 | 2.79 | -4.17 | 1.46 | 0.010 1 | 0.045 8 |
| | CTN-067 M3 vs. CTN-067 M6 | 1 1 | 90.44 | 4.12 | 1 1 | 92.78 | 2.79 | -2.36 | 1.28 | 0.079 6 | 0.281 1 |
| CD14+ \| Geometric Mean (BUV661-A :: CD27) | Control vs. CTN-067 M0 | 8 | 1397.25 | 489.89 | 1 1 | 1302.09 | 495.94 | -93.37 | 282.03 | 0.744 2 | 0.987 1 |
| | Control vs. CTN-067 M3 | 8 | 1397.25 | 489.89 | 1 1 | 1456.00 | 525.94 | -165.53 | 258.97 | 0.530 3 | 0.918 0 |
| | Control vs. CTN-067 M6 | 8 | 1397.25 | 489.89 | 1 1 | 1530.73 | 374.52 | -236.19 | 258.11 | 0.371 6 | 0.797 1 |
| | CTN-067 M0 vs. CTN-067 M3 | 1 1 | 1302.09 | 495.94 | 1 1 | 1456.00 | 525.94 | -72.16 | 119.80 | 0.554 1 | 0.930 1 |
| | CTN-067 M0 vs. CTN-067 M6 | 1 1 | 1302.09 | 495.94 | 1 1 | 1530.73 | 374.52 | -142.82 | 121.24 | 0.253 3 | 0.647 5 |
| | CTN-067 M3 vs. CTN-067 M6 | 1 1 | 1456.00 | 525.94 | 1 1 | 1530.73 | 374.52 | -70.66 | 104.62 | 0.507 5 | 0.905 1 |
| CD14+ \| Geometric Mean (BV510-A :: CCR7) | Control vs. CTN-067 M0 | 8 | 140.63 | 783.12 | 1 1 | -409.07 | 567.16 | 808.21 | 410.19 | 0.063 6 | 0.233 7 |
| | Control vs. CTN-067 M3 | 8 | 140.63 | 783.12 | 1 1 | -157.00 | 695.74 | 433.23 | 368.60 | 0.254 4 | 0.649 1 |
| | Control vs. CTN-067 M6 | 8 | 140.63 | 783.12 | 1 1 | -11.18 | 578.42 | 281.30 | 367.04 | 0.452 9 | 0.868 5 |
| | CTN-067 M0 vs. CTN-067 M3 | 1 1 | -409.07 | 567.16 | 1 1 | -157.00 | 695.74 | -374.98 | 195.40 | 0.070 1 | 0.253 5 |
| | CTN-067 M0 vs. CTN-067 M6 | 1 1 | -409.07 | 567.16 | 1 1 | -11.18 | 578.42 | -526.91 | 197.70 | 0.015 3 | 0.066 9 |
| | CTN-067 M3 vs. CTN-067 M6 | 1 1 | -157.00 | 695.74 | 1 1 | -11.18 | 578.42 | -151.93 | 171.28 | 0.386 2 | 0.811 6 |
| CD14+/CCR7+ \| Freq. of Parent | Control vs. CTN-067 M0 | 8 | 1.23 | 1.44 | 1 1 | 1.90 | 1.41 | -0.24 | 1.21 | 0.842 2 | 0.997 0 |
| | Control vs. CTN-067 M3 | 8 | 1.23 | 1.44 | 1 1 | 3.08 | 2.41 | -1.59 | 1.02 | 0.135 5 | 0.424 1 |
| | Control vs. CTN-067 M6 | 8 | 1.23 | 1.44 | 1 1 | 2.27 | 1.71 | -0.79 | 1.01 | 0.443 2 | 0.861 2 |

| | | | | | | | | | | |
|---|---|---|---|---|---|---|---|---|---|---|
| | CTN-067 M0 vs. CTN-067 M3 | 11 | 1.90 | 1.41 | 11 | 3.08 | 2.41 | -1.35 | 0.78 | 0.1021 | 0.3422 |
| | CTN-067 M0 vs. CTN-067 M6 | 11 | 1.90 | 1.41 | 11 | 2.27 | 1.71 | -0.55 | 0.79 | 0.4967 | 0.8985 |
| | CTN-067 M3 vs. CTN-067 M6 | 11 | 3.08 | 2.41 | 11 | 2.27 | 1.71 | 0.80 | 0.70 | 0.2717 | 0.6749 |
| CD14+ \| Geometric Mean (BV605-A :: CD45RA) | Control vs. CTN-067 M0 | 8 | 7227.13 | 3238.18 | 11 | 15807.09 | 10183.60 | -9782.07 | 4090.10 | 0.0273 | 0.1128 |
| | Control vs. CTN-067 M3 | 8 | 7227.13 | 3238.18 | 11 | 12802.09 | 6670.61 | -6606.86 | 3710.14 | 0.0909 | 0.3126 |
| | Control vs. CTN-067 M6 | 8 | 7227.13 | 3238.18 | 11 | 12379.91 | 6164.03 | -6176.21 | 3695.96 | 0.1111 | 0.3654 |
| | CTN-067 M0 vs. CTN-067 M3 | 11 | 15807.09 | 10183.60 | 11 | 12802.09 | 6670.61 | 3175.22 | 1858.48 | 0.1038 | 0.3468 |
| | CTN-067 M0 vs. CTN-067 M6 | 11 | 15807.09 | 10183.60 | 11 | 12379.91 | 6164.03 | 3605.86 | 1880.59 | 0.0703 | 0.2541 |
| | CTN-067 M3 vs. CTN-067 M6 | 11 | 12802.09 | 6670.61 | 11 | 12379.91 | 6164.03 | 430.64 | 1626.22 | 0.7940 | 0.9933 |
| CD14+/CD45RA+ \| Freq. of Parent | Control vs. CTN-067 M0 | 8 | 56.90 | 21.84 | 11 | 70.51 | 19.50 | -5.66 | 11.72 | 0.6346 | 0.9619 |
| | Control vs. CTN-067 M3 | 8 | 56.90 | 21.84 | 11 | 69.61 | 20.88 | -8.96 | 11.14 | 0.4312 | 0.8516 |
| | Control vs. CTN-067 M6 | 8 | 56.90 | 21.84 | 11 | 69.57 | 19.80 | -9.14 | 11.12 | 0.4217 | 0.8437 |
| | CTN-067 M0 vs. CTN-067 M3 | 11 | 70.51 | 19.50 | 11 | 69.61 | 20.88 | -3.30 | 3.84 | 0.4002 | 0.8248 |
| | CTN-067 M0 vs. CTN-067 M6 | 11 | 70.51 | 19.50 | 11 | 69.57 | 19.80 | -3.47 | 3.88 | 0.3823 | 0.8078 |
| | CTN-067 M3 vs. CTN-067 M6 | 11 | 69.61 | 20.88 | 11 | 69.57 | 19.80 | -0.17 | 3.34 | 0.9593 | 0.9999 |
| CD14+ \| Geometric Mean (BV650-A :: CD15) | Control vs. CTN-067 M0 | 8 | 2136.13 | 778.20 | 11 | 1709.00 | 745.28 | 1093.04 | 814.90 | 0.1956 | 0.5493 |
| | Control vs. CTN-067 M3 | 8 | 2136.13 | 778.20 | 11 | 2664.00 | 2360.66 | 56.86 | 681.58 | 0.9344 | 0.9998 |

| | | | | | | | | | | | |
|---|---|---|---|---|---|---|---|---|---|---|---|
| | Control vs. CTN-067 M6 | 8 | 2136.13 | 778.20 | 11 | 2131.45 | 582.74 | 585.37 | 676.37 | 0.3976 | 0.8224 |
| | CTN-067 M0 vs. CTN-067 M3 | 11 | 1709.00 | 745.28 | 11 | 2664.00 | 2360.66 | -1036.18 | 593.47 | 0.0970 | 0.3288 |
| | CTN-067 M0 vs. CTN-067 M6 | 11 | 1709.00 | 745.28 | 11 | 2131.45 | 582.74 | -507.67 | 598.14 | 0.4066 | 0.8306 |
| | CTN-067 M3 vs. CTN-067 M6 | 11 | 2664.00 | 2360.66 | 11 | 2131.45 | 582.74 | 528.51 | 545.64 | 0.3449 | 0.7685 |
| CD14+/CD15+ \| Freq. of Parent | Control vs. CTN-067 M0 | 8 | 37.03 | 11.95 | 11 | 30.88 | 10.48 | 10.04 | 7.06 | 0.1714 | 0.5019 |
| | Control vs. CTN-067 M3 | 8 | 37.03 | 11.95 | 11 | 38.14 | 17.29 | 3.25 | 6.16 | 0.6036 | 0.9512 |
| | Control vs. CTN-067 M6 | 8 | 37.03 | 11.95 | 11 | 37.40 | 7.96 | 4.02 | 6.13 | 0.5200 | 0.9123 |
| | CTN-067 M0 vs. CTN-067 M3 | 11 | 30.88 | 10.48 | 11 | 38.14 | 17.29 | -6.78 | 3.84 | 0.0930 | 0.3183 |
| | CTN-067 M0 vs. CTN-067 M6 | 11 | 30.88 | 10.48 | 11 | 37.40 | 7.96 | -6.02 | 3.88 | 0.1371 | 0.4279 |
| | CTN-067 M3 vs. CTN-067 M6 | 11 | 38.14 | 17.29 | 11 | 37.40 | 7.96 | 0.76 | 3.39 | 0.8242 | 0.9958 |
| CD14+ \| Geometric Mean (BV711-A :: CD86) | Control vs. CTN-067 M0 | 8 | 10108.00 | 1181.97 | 11 | 18757.55 | 4155.46 | -9658.42 | 2293.27 | 0.0005 | 0.0024 |
| | Control vs. CTN-067 M3 | 8 | 10108.00 | 1181.97 | 11 | 13544.00 | 4971.63 | -4402.46 | 1922.64 | 0.0336 | 0.1357 |
| | Control vs. CTN-067 M6 | 8 | 10108.00 | 1181.97 | 11 | 14469.73 | 2617.87 | -5326.08 | 1908.19 | 0.0116 | 0.0521 |
| | CTN-067 M0 vs. CTN-067 M3 | 11 | 18757.55 | 4155.46 | 11 | 13544.00 | 4971.63 | 5255.96 | 1712.94 | 0.0063 | 0.0295 |
| | CTN-067 M0 vs. CTN-067 M6 | 11 | 18757.55 | 4155.46 | 11 | 14469.73 | 2617.87 | 4332.34 | 1725.62 | 0.0213 | 0.0903 |
| | CTN-067 M3 vs. CTN-067 M6 | 11 | 13544.00 | 4971.63 | 11 | 14469.73 | 2617.87 | -923.62 | 1583.50 | 0.5666 | 0.9359 |
| CD14+/CD86+ \| Freq. of Parent | Control vs. CTN-067 M0 | 8 | 94.41 | 2.45 | 11 | 94.75 | 3.61 | -1.06 | 4.01 | 0.7946 | 0.9933 |

| | | | | | | | | | | | |
|---|---|---|---|---|---|---|---|---|---|---|---|
| | Control vs. CTN-067 M3 | 8 | 94.41 | 2.45 | 11 | 91.46 | 11.19 | 1.78 | 3.36 | 0.6021 | 0.9506 |
| | Control vs. CTN-067 M6 | 8 | 94.41 | 2.45 | 11 | 96.07 | 2.74 | -2.85 | 3.33 | 0.4030 | 0.8274 |
| | CTN-067 M0 vs. CTN-067 M3 | 11 | 94.75 | 3.61 | 11 | 91.46 | 11.19 | 2.84 | 2.73 | 0.3109 | 0.7279 |
| | CTN-067 M0 vs. CTN-067 M6 | 11 | 94.75 | 3.61 | 11 | 96.07 | 2.74 | -1.79 | 2.75 | 0.5232 | 0.9141 |
| | CTN-067 M3 vs. CTN-067 M6 | 11 | 91.46 | 11.19 | 11 | 96.07 | 2.74 | -4.63 | 2.47 | 0.0767 | 0.2728 |
| CD14+ \| Geometric Mean (PE-A :: CD38) | Control vs. CTN-067 M0 | 8 | 44862.25 | 6977.41 | 11 | 33436.64 | 12269.49 | 15830.00 | 6356.17 | 0.0222 | 0.0938 |
| | Control vs. CTN-067 M3 | 8 | 44862.25 | 6977.41 | 11 | 39621.27 | 20534.87 | 11247.00 | 5331.38 | 0.0484 | 0.1858 |
| | Control vs. CTN-067 M6 | 8 | 44862.25 | 6977.41 | 11 | 35845.27 | 10489.06 | 15103.00 | 5291.44 | 0.0101 | 0.0459 |
| | CTN-067 M0 vs. CTN-067 M3 | 11 | 33436.64 | 12269.49 | 11 | 39621.27 | 20534.87 | -4582.90 | 4198.78 | 0.2887 | 0.6988 |
| | CTN-067 M0 vs. CTN-067 M6 | 11 | 33436.64 | 12269.49 | 11 | 35845.27 | 10489.06 | -727.26 | 4238.39 | 0.8656 | 0.9981 |
| | CTN-067 M3 vs. CTN-067 M6 | 11 | 39621.27 | 20534.87 | 11 | 35845.27 | 10489.06 | 3855.63 | 3789.33 | 0.3217 | 0.7414 |
| CD14+/CD38+ \| Freq. of Parent | Control vs. CTN-067 M0 | 8 | 95.53 | 1.81 | 11 | 87.73 | 6.07 | 11.26 | 2.56 | 0.0003 | 0.0016 |
| | Control vs. CTN-067 M3 | 8 | 95.53 | 1.81 | 11 | 88.08 | 5.99 | 10.35 | 2.16 | 0.0001 | 0.0007 |
| | Control vs. CTN-067 M6 | 8 | 95.53 | 1.81 | 11 | 90.99 | 2.96 | 7.41 | 2.14 | 0.0027 | 0.0130 |
| | CTN-067 M0 vs. CTN-067 M3 | 11 | 87.73 | 6.07 | 11 | 88.08 | 5.99 | -0.92 | 1.63 | 0.5814 | 0.9424 |
| | CTN-067 M0 vs. CTN-067 M6 | 11 | 87.73 | 6.07 | 11 | 90.99 | 2.96 | -3.85 | 1.65 | 0.0306 | 0.1248 |
| | CTN-067 M3 vs. CTN-067 M6 | 11 | 88.08 | 5.99 | 11 | 90.99 | 2.96 | -2.94 | 1.47 | 0.0595 | 0.2214 |

| | | | | | | | | | | | |
|---|---|---|---|---|---|---|---|---|---|---|---|
| CD14+ \| Geometric Mean (R718-A :: CCR5) | Control vs. CTN-067 M0 | 8 | 1532.63 | 620.81 | 11 | 1881.73 | 519.48 | 440.07 | 355.73 | 0.2311 | 0.6119 |
| | Control vs. CTN-067 M3 | 8 | 1532.63 | 620.81 | 11 | 1699.55 | 572.66 | 318.83 | 315.31 | 0.3246 | 0.7450 |
| | Control vs. CTN-067 M6 | 8 | 1532.63 | 620.81 | 11 | 1899.91 | 829.82 | 103.39 | 313.79 | 0.7454 | 0.9873 |
| | CTN-067 M0 vs. CTN-067 M3 | 11 | 1881.73 | 519.48 | 11 | 1699.55 | 572.66 | -121.23 | 180.57 | 0.5100 | 0.9066 |
| | CTN-067 M0 vs. CTN-067 M6 | 11 | 1881.73 | 519.48 | 11 | 1899.91 | 829.82 | -336.68 | 182.65 | 0.0809 | 0.2848 |
| | CTN-067 M3 vs. CTN-067 M6 | 11 | 1699.55 | 572.66 | 11 | 1899.91 | 829.82 | -215.45 | 158.74 | 0.1906 | 0.5398 |
| CD14+/CCR5+ \| Freq. of Parent | Control vs. CTN-067 M0 | 8 | 12.77 | 8.27 | 11 | 20.08 | 9.83 | -1.19 | 5.25 | 0.8232 | 0.9958 |
| | Control vs. CTN-067 M3 | 8 | 12.77 | 8.27 | 11 | 17.45 | 7.51 | -0.89 | 4.59 | 0.8484 | 0.9973 |
| | Control vs. CTN-067 M6 | 8 | 12.77 | 8.27 | 11 | 18.64 | 8.02 | -2.20 | 4.57 | 0.6363 | 0.9625 |
| | CTN-067 M0 vs. CTN-067 M3 | 11 | 20.08 | 9.83 | 11 | 17.45 | 7.51 | 0.30 | 2.82 | 0.9165 | 0.9996 |
| | CTN-067 M0 vs. CTN-067 M6 | 11 | 20.08 | 9.83 | 11 | 18.64 | 8.02 | -1.01 | 2.85 | 0.7284 | 0.9845 |
| | CTN-067 M3 vs. CTN-067 M6 | 11 | 17.45 | 7.51 | 11 | 18.64 | 8.02 | -1.31 | 2.49 | 0.6060 | 0.9521 |
| CD14+ \| Geometric Mean (RB545-A :: CD57) | Control vs. CTN-067 M0 | 8 | 1017.25 | 682.69 | 11 | 730.91 | 551.04 | 199.89 | 371.73 | 0.5970 | 0.9487 |
| | Control vs. CTN-067 M3 | 8 | 1017.25 | 682.69 | 11 | 722.32 | 768.25 | 214.34 | 358.27 | 0.5567 | 0.9313 |
| | Control vs. CTN-067 M6 | 8 | 1017.25 | 682.69 | 11 | 801.56 | 633.46 | 135.39 | 357.78 | 0.7093 | 0.9810 |
| | CTN-067 M0 vs. CTN-067 M3 | 11 | 730.91 | 551.04 | 11 | 722.32 | 768.25 | 14.46 | 104.38 | 0.8913 | 0.9990 |
| | CTN-067 M0 vs. CTN-067 M6 | 11 | 730.91 | 551.04 | 11 | 801.56 | 633.46 | -64.50 | 105.69 | 0.5489 | 0.9276 |
| | CTN-067 M3 vs. | 11 | 722.32 | 768.25 | 11 | 801.56 | 633.46 | -78.95 | 90.62 | 0.3945 | 0.8195 |

| | | | | | | | | | | | |
|---|---|---|---|---|---|---|---|---|---|---|---|
| | CTN-067 M6 | | | | | | | | | | |
| CD14+ \| Geometric Mean (RB705-A :: CCR2) | Control vs. CTN-067 M0 | 8 | 72874.25 | 26794.83 | 11 | 30784.18 | 18972.18 | 52820.00 | 12767.00 | 0.0006 | 0.0029 |
| | Control vs. CTN-067 M3 | 8 | 72874.25 | 26794.83 | 11 | 38231.18 | 17759.71 | 43433.00 | 11540.00 | 0.0013 | 0.0066 |
| | Control vs. CTN-067 M6 | 8 | 72874.25 | 26794.83 | 11 | 44130.27 | 21909.16 | 37438.00 | 11495.00 | 0.0041 | 0.0198 |
| | CTN-067 M0 vs. CTN-067 M3 | 11 | 30784.18 | 18972.18 | 11 | 38231.18 | 17759.71 | -9387.04 | 5905.23 | 0.1284 | 0.4077 |
| | CTN-067 M0 vs. CTN-067 M6 | 11 | 30784.18 | 18972.18 | 11 | 44130.27 | 21909.16 | -15383.00 | 5975.21 | 0.0186 | 0.0799 |
| | CTN-067 M3 vs. CTN-067 M6 | 11 | 38231.18 | 17759.71 | 11 | 44130.27 | 21909.16 | -5995.54 | 5170.41 | 0.2606 | 0.6585 |
| CD14+/CCR2+ \| Freq. of Parent | Control vs. CTN-067 M0 | 8 | 92.53 | 3.03 | 11 | 72.30 | 14.35 | 23.58 | 5.56 | 0.0004 | 0.0023 |
| | Control vs. CTN-067 M3 | 8 | 92.53 | 3.03 | 11 | 80.85 | 8.06 | 14.75 | 4.81 | 0.0064 | 0.0297 |
| | Control vs. CTN-067 M6 | 8 | 92.53 | 3.03 | 11 | 83.40 | 7.45 | 12.18 | 4.78 | 0.0197 | 0.0842 |
| | CTN-067 M0 vs. CTN-067 M3 | 11 | 72.30 | 14.35 | 11 | 80.85 | 8.06 | -8.82 | 3.13 | 0.0109 | 0.0489 |
| | CTN-067 M0 vs. CTN-067 M6 | 11 | 72.30 | 14.35 | 11 | 83.40 | 7.45 | -11.39 | 3.16 | 0.0019 | 0.0093 |
| | CTN-067 M3 vs. CTN-067 M6 | 11 | 80.85 | 8.06 | 11 | 83.40 | 7.45 | -2.57 | 2.77 | 0.3648 | 0.7901 |
| CD14+ \| Geometric Mean (RB780-A :: HLA-DR) | Control vs. CTN-067 M0 | 8 | 114434.88 | 26444.80 | 11 | 234291.18 | 73187.11 | -88746.00 | 34787.00 | 0.0195 | 0.0835 |
| | Control vs. CTN-067 M3 | 8 | 114434.88 | 26444.80 | 11 | 167965.64 | 64858.02 | -33029.00 | 31275.00 | 0.3042 | 0.7194 |
| | Control vs. CTN-067 M6 | 8 | 114434.88 | 26444.80 | 11 | 161418.27 | 62464.73 | -27009.00 | 31143.00 | 0.3966 | 0.8215 |
| | CTN-067 M0 vs. CTN-067 M3 | 11 | 234291.18 | 73187.11 | 11 | 167965.64 | 64858.02 | 55717.00 | 16533.00 | 0.0032 | 0.0156 |
| | CTN-067 M0 vs. CTN-067 M6 | 11 | 234291.18 | 73187.11 | 11 | 161418.27 | 62464.73 | 61737.00 | 16727.00 | 0.0016 | 0.0077 |

| | | | | | | | | | | | |
|---|---|---|---|---|---|---|---|---|---|---|---|
| | CTN-067 M3 vs. CTN-067 M6 | 11 | 167965.64 | 64858.02 | 11 | 161418.27 | 62464.73 | 6019.93 | 14490.00 | 0.6825 | 0.9752 |
| CD14+/HLA-DR+ \| Freq. of Parent | Control vs. CTN-067 M0 | 8 | 94.36 | 1.06 | 11 | 94.60 | 1.81 | -0.54 | 1.41 | 0.7064 | 0.9804 |
| | Control vs. CTN-067 M3 | 8 | 94.36 | 1.06 | 11 | 92.63 | 3.63 | 1.25 | 1.23 | 0.3200 | 0.7394 |
| | Control vs. CTN-067 M6 | 8 | 94.36 | 1.06 | 11 | 94.70 | 2.15 | -0.83 | 1.22 | 0.5045 | 0.9033 |
| | CTN-067 M0 vs. CTN-067 M3 | 11 | 94.60 | 1.81 | 11 | 92.63 | 3.63 | 1.79 | 0.77 | 0.0318 | 0.1290 |
| | CTN-067 M0 vs. CTN-067 M6 | 11 | 94.60 | 1.81 | 11 | 94.70 | 2.15 | -0.29 | 0.78 | 0.7137 | 0.9818 |
| | CTN-067 M3 vs. CTN-067 M6 | 11 | 92.63 | 3.63 | 11 | 94.70 | 2.15 | -2.08 | 0.68 | 0.0066 | 0.0307 |
| CD14+ \| Geometric Mean (RY586-A :: CXCR4) | Control vs. CTN-067 M0 | 8 | 12470.50 | 4536.97 | 11 | 39278.55 | 17444.27 | -2730.67 | 13517.00 | 0.8420 | 0.9970 |
| | Control vs. CTN-067 M3 | 8 | 12470.50 | 4536.97 | 11 | 25920.36 | 27284.27 | 3861.57 | 11318.00 | 0.7367 | 0.9859 |
| | Control vs. CTN-067 M6 | 8 | 12470.50 | 4536.97 | 11 | 27246.27 | 31982.00 | 2199.27 | 11232.00 | 0.8468 | 0.9972 |
| | CTN-067 M0 vs. CTN-067 M3 | 11 | 39278.55 | 17444.27 | 11 | 25920.36 | 27284.27 | 6592.24 | 9982.25 | 0.5169 | 0.9106 |
| | CTN-067 M0 vs. CTN-067 M6 | 11 | 39278.55 | 17444.27 | 11 | 27246.27 | 31982.00 | 4929.95 | 10058.00 | 0.6296 | 0.9603 |
| | CTN-067 M3 vs. CTN-067 M6 | 11 | 25920.36 | 27284.27 | 11 | 27246.27 | 31982.00 | -1662.29 | 9205.17 | 0.8586 | 0.9978 |
| CD14+/CXCR4+ \| Freq. of Parent | Control vs. CTN-067 M0 | 8 | 83.68 | 5.20 | 11 | 90.90 | 8.41 | -6.79 | 4.44 | 0.1426 | 0.4403 |
| | Control vs. CTN-067 M3 | 8 | 83.68 | 5.20 | 11 | 84.81 | 9.58 | 0.36 | 3.81 | 0.9259 | 0.9997 |
| | Control vs. CTN-067 M6 | 8 | 83.68 | 5.20 | 11 | 87.88 | 6.97 | -2.66 | 3.79 | 0.4906 | 0.8946 |
| | CTN-067 M0 vs. CTN-067 M3 | 11 | 90.90 | 8.41 | 11 | 84.81 | 9.58 | 7.15 | 2.59 | 0.0124 | 0.0554 |
| | CTN-067 M0 vs. | 11 | 90.90 | 8.41 | 11 | 87.88 | 6.97 | 4.13 | 2.62 | 0.1312 | 0.4141 |

| | | | | | | | | | | | |
|---|---|---|---|---|---|---|---|---|---|---|---|
| | CTN-067 M6 | | | | | | | | | | |
| | CTN-067 M3 vs. CTN-067 M6 | 11 | 84.81 | 9.58 | 11 | 87.88 | 6.97 | -3.02 | 2.30 | 0.2047 | 0.5660 |
| CD14+/CD15-/HLA-DR_Low \| Freq. of Parent | Control vs. CTN-067 M0 | 8 | 9.19 | 1.99 | 11 | 9.69 | 4.67 | 1.21 | 3.28 | 0.7157 | 0.9822 |
| | Control vs. CTN-067 M3 | 8 | 9.19 | 1.99 | 11 | 13.32 | 8.46 | -2.32 | 2.78 | 0.4145 | 0.8376 |
| | Control vs. CTN-067 M6 | 8 | 9.19 | 1.99 | 11 | 8.48 | 4.21 | 2.52 | 2.77 | 0.3732 | 0.7987 |
| | CTN-067 M0 vs. CTN-067 M3 | 11 | 9.69 | 4.67 | 11 | 13.32 | 8.46 | -3.54 | 2.01 | 0.0953 | 0.3243 |
| | CTN-067 M0 vs. CTN-067 M6 | 11 | 9.69 | 4.67 | 11 | 8.48 | 4.21 | 1.31 | 2.03 | 0.5276 | 0.9165 |
| | CTN-067 M3 vs. CTN-067 M6 | 11 | 13.32 | 8.46 | 11 | 8.48 | 4.21 | 4.85 | 1.80 | 0.0144 | 0.0633 |
| CD14+/CD15-/HLA-DR_Low \| Geometric Mean (RB780-A :: HLA-DR) | Control vs. CTN-067 M0 | 8 | 5427.13 | 1701.80 | 11 | 7508.09 | 2548.64 | -1363.96 | 1205.65 | 0.2720 | 0.6753 |
| | Control vs. CTN-067 M3 | 8 | 5427.13 | 1701.80 | 11 | 5073.18 | 973.09 | 748.17 | 1027.37 | 0.4753 | 0.8846 |
| | Control vs. CTN-067 M6 | 8 | 5427.13 | 1701.80 | 11 | 5419.09 | 1659.40 | 386.21 | 1020.48 | 0.7093 | 0.9810 |
| | CTN-067 M0 vs. CTN-067 M3 | 11 | 7508.09 | 2548.64 | 11 | 5073.18 | 973.09 | 2112.12 | 726.41 | 0.0090 | 0.0412 |
| | CTN-067 M0 vs. CTN-067 M6 | 11 | 7508.09 | 2548.64 | 11 | 5419.09 | 1659.40 | 1750.17 | 734.02 | 0.0277 | 0.1143 |
| | CTN-067 M3 vs. CTN-067 M6 | 11 | 5073.18 | 973.09 | 11 | 5419.09 | 1659.40 | -361.96 | 647.27 | 0.5826 | 0.9429 |
| CD14++CD16- \| Freq. of Parent | Control vs. CTN-067 M0 | 8 | 86.05 | 4.63 | 11 | 64.05 | 14.72 | 22.57 | 6.16 | 0.0017 | 0.0082 |
| | Control vs. CTN-067 M3 | 8 | 86.05 | 4.63 | 11 | 70.91 | 11.86 | 16.40 | 5.54 | 0.0080 | 0.0369 |
| | Control vs. CTN-067 M6 | 8 | 86.05 | 4.63 | 11 | 72.15 | 8.08 | 15.19 | 5.52 | 0.0126 | 0.0562 |
| | CTN-067 M0 vs. CTN-067 M3 | 11 | 64.05 | 14.72 | 11 | 70.91 | 11.86 | -6.17 | 2.93 | 0.0490 | 0.1880 |

| | | | | | | | | | | | |
|---|---|---|---|---|---|---|---|---|---|---|---|
| | CTN-067 M0 vs. CTN-067 M6 | 11 | 64.05 | 14.72 | 11 | 72.15 | 8.08 | -7.38 | 2.97 | 0.0224 | 0.0945 |
| | CTN-067 M3 vs. CTN-067 M6 | 11 | 70.91 | 11.86 | 11 | 72.15 | 8.08 | -1.21 | 2.57 | 0.6432 | 0.9646 |
| CD14++CD16+ \| Freq. of Parent | Control vs. CTN-067 M0 | 8 | 3.24 | 1.49 | 11 | 14.67 | 8.80 | -8.88 | 4.04 | 0.0407 | 0.1601 |
| | Control vs. CTN-067 M3 | 8 | 3.24 | 1.49 | 11 | 9.29 | 4.42 | -4.63 | 3.42 | 0.1911 | 0.5407 |
| | Control vs. CTN-067 M6 | 8 | 3.24 | 1.49 | 11 | 12.62 | 6.24 | -8.01 | 3.39 | 0.0290 | 0.1192 |
| | CTN-067 M0 vs. CTN-067 M3 | 11 | 14.67 | 8.80 | 11 | 9.29 | 4.42 | 4.25 | 2.55 | 0.1115 | 0.3665 |
| | CTN-067 M0 vs. CTN-067 M6 | 11 | 14.67 | 8.80 | 11 | 12.62 | 6.24 | 0.87 | 2.57 | 0.7381 | 0.9861 |
| | CTN-067 M3 vs. CTN-067 M6 | 11 | 9.29 | 4.42 | 11 | 12.62 | 6.24 | -3.38 | 2.28 | 0.1552 | 0.4682 |
| CD14++CD16+ \| Freq. of Parent | Control vs. CTN-067 M0 | 8 | 1.16 | 0.77 | 11 | 7.74 | 7.58 | -10.33 | 2.64 | 0.0009 | 0.0047 |
| | Control vs. CTN-067 M3 | 8 | 1.16 | 0.77 | 11 | 5.18 | 4.23 | -6.94 | 2.27 | 0.0065 | 0.0301 |
| | Control vs. CTN-067 M6 | 8 | 1.16 | 0.77 | 11 | 4.27 | 3.12 | -5.99 | 2.26 | 0.0157 | 0.0684 |
| | CTN-067 M0 vs. CTN-067 M3 | 11 | 7.74 | 7.58 | 11 | 5.18 | 4.23 | 3.39 | 1.52 | 0.0373 | 0.1486 |
| | CTN-067 M0 vs. CTN-067 M6 | 11 | 7.74 | 7.58 | 11 | 4.27 | 3.12 | 4.35 | 1.53 | 0.0105 | 0.0474 |
| | CTN-067 M3 vs. CTN-067 M6 | 11 | 5.18 | 4.23 | 11 | 4.27 | 3.12 | 0.95 | 1.34 | 0.4859 | 0.8916 |
| CD14++CD16+ \| Freq. of Parent | Control vs. CTN-067 M0 | 8 | 1659.75 | 300.71 | 11 | 5429.45 | 2845.28 | -4497.58 | 1100.30 | 0.0006 | 0.0032 |
| | Control vs. CTN-067 M3 | 8 | 1659.75 | 300.71 | 11 | 3420.64 | 1051.66 | -2271.96 | 928.90 | 0.0244 | 0.1020 |
| | Control vs. CTN-067 M6 | 8 | 1659.75 | 300.71 | 11 | 3666.00 | 1236.01 | -2506.54 | 922.24 | 0.0137 | 0.0603 |
| | CTN-067 M0 vs. | 11 | 5429.45 | 2845.28 | 11 | 3420.64 | 1051.66 | 2225.62 | 695.41 | 0.0047 | 0.0223 |

**Commented [CL1]:** the name of the populations in the last 3 sections is the same, so must be a typo or copy error?

| | | | | | | | | | | | |
|---|---|---|---|---|---|---|---|---|---|---|---|
| | CTN-067 M3 | | | | | | | | | | |
| | CTN-067 M0 vs. CTN-067 M6 | 1 1 | 5429.45 | 2845.28 | 1 1 | 3666.00 | 1236.01 | 1991.04 | 702.36 | 0.010 6 | 0.047 7 |
| | CTN-067 M3 vs. CTN-067 M6 | 1 1 | 3420.64 | 1051.66 | 1 1 | 3666.00 | 1236.01 | -234.58 | 623.39 | 0.710 9 | 0.981 3 |
