## Supplemental Table 5 for "Chronic opioid-associated immune dysregulation among people living with HIV"

**Supplementary Table 5.** Flow data stratified by HIV viral load

| Marker | Column B | Vs | Column A | P value | Median of Column A | Median of Column B | Comparison | Minimum | 25% percentile | Median | 75% percentile | Maximum | Mean | Std. Deviation | Std. Error of mean | Lower 95% CL | Upper 95% CL |
|---|---|---|---|---|---|---|---|---|---|---|---|---|---|---|---|---|---|
| CD14+CD11b+ (MFI) | Control | vs. | LVL | 0.122 | 4.857, n=10 | 5.002, n=8 | LVL | 4.769 | 4.804 | 4.857 | 5.076 | 5.105 | 4.907 | 0.1318 | 0.04167 | 4.813 | 5.002 |
| | | | | | | | Control | 4.906 | 4.961 | 5.002 | 5.061 | 5.099 | 5.008 | 0.06213 | 0.02197 | 4.956 | 5.06 |
| | Control | vs. | HVL | 0.0927 | 4.876, n=9 | 5.002, n=8 | HVL | 4.625 | 4.78 | 4.876 | 5.017 | 5.091 | 4.885 | 0.1524 | 0.0508 | 4.767 | 5.002 |
| | | | | | | | Control | 4.906 | 4.961 | 5.002 | 5.061 | 5.099 | 5.008 | 0.06213 | 0.02197 | 4.956 | 5.06 |
| CD14+CD86+ (MFI) | Control | vs. | LVL | 0.0085 | 4.158, n=10 | 4.009, n=8 | LVL | 3.413 | 4.078 | 4.158 | 4.195 | 4.298 | 4.092 | 0.2508 | 0.07929 | 3.913 | 4.271 |
| | | | | | | | Control | 3.933 | 3.948 | 4.009 | 4.052 | 4.066 | 4.002 | 0.05124 | 0.01812 | 3.959 | 4.045 |
| | Control | vs. | HVL | <0.0001 | 4.205, n=9 | 4.009, n=8 | HVL | 4.128 | 4.176 | 4.205 | 4.233 | 4.353 | 4.214 | 0.0625 | 0.02083 | 4.166 | 4.262 |
| | | | | | | | Control | 3.933 | 3.948 | 4.009 | 4.052 | 4.066 | 4.002 | 0.05124 | 0.01812 | 3.959 | 4.045 |
| CD14+CD38+ (MFI) | Control | vs. | LVL | 0.1011 | 4.509, n=10 | 4.655, n=8 | LVL | 4.301 | 4.423 | 4.509 | 4.686 | 4.896 | 4.546 | 0.18 | 0.05692 | 4.418 | 4.675 |
| | | | | | | | Control | 4.535 | 4.584 | 4.655 | 4.709 | 4.73 | 4.647 | 0.06974 | 0.02466 | 4.589 | 4.705 |
| | Control | vs. | HVL | 0.1388 | 4.588, n=9 | 4.655, n=8 | HVL | 4.345 | 4.4 | 4.588 | 4.654 | 4.738 | 4.539 | 0.1388 | 0.04627 | 4.432 | 4.645 |
| | | | | | | | Control | 4.535 | 4.584 | 4.655 | 4.709 | 4.73 | 4.647 | 0.06974 | 0.02466 | 4.589 | 4.705 |
| CD14+CCR2+ (MFI) | Control | vs. | LVL | 0.0155 | 4.570, n=10 | 4.912, n=8 | LVL | 4.179 | 4.371 | 4.57 | 4.687 | 4.795 | 4.534 | 0.1991 | 0.06295 | 4.391 | 4.676 |
| | | | | | | | Control | 4.531 | 4.6 | 4.912 | 4.983 | 4.998 | 4.829 | 0.194 | 0.06859 | 4.667 | 4.991 |
| | Control | vs. | HVL | 0.0152 | 4.546, n=9 | 4.912, n=8 | HVL | 3.848 | 4.255 | 4.546 | 4.641 | 4.838 | 4.452 | 0.3061 | 0.102 | 4.217 | 4.687 |
| | | | | | | | Control | 4.531 | 4.6 | 4.912 | 4.983 | 4.998 | 4.829 | 0.194 | 0.06859 | 4.667 | 4.991 |
| CD14++CD16- (Freq. of Parent) | Control | vs. | LVL | 0.0002 | 72.25, n=10 | 87.40, n=8 | LVL | 53.4 | 64.98 | 72.25 | 77.25 | 80.3 | 70.36 | 8.361 | 2.644 | 64.38 | 76.34 |
| | | | | | | | Control | 77 | 83.15 | 87.4 | 89.88 | 90.9 | 86.05 | 4.626 | 1.636 | 82.18 | 89.92 |
| | Control | vs. | HVL | 0.001 | 69.10, n=9 | 87.40, n=8 | HVL | 46.2 | 61.3 | 69.1 | 77.3 | 87 | 68.31 | 11.86 | 3.955 | 59.19 | 77.43 |
| | | | | | | | Control | 77 | 83.15 | 87.4 | 89.88 | 90.9 | 86.05 | 4.626 | 1.636 | 82.18 | 89.92 |
| CD14++CD16+ (Freq. of Parent) | Control | vs. | LVL | <0.0001 | 11.25, n=10 | 2.875, n=8 | LVL | 5.28 | 7.49 | 11.25 | 14.55 | 28 | 12.31 | 6.802 | 2.151 | 7.439 | 17.17 |
| | | | | | | | Control | 1.36 | 1.89 | 2.875 | 4.603 | 5.55 | 3.239 | 1.494 | 0.5282 | 1.99 | 4.488 |
| | Control | vs. | HVL | 0.0006 | 10.40, n=9 | 2.875, n=8 | HVL | 4.11 | 8.675 | 10.4 | 20.1 | 31.6 | 13.82 | 9.445 | 3.148 | 6.565 | 21.08 |
| | | | | | | | Control | 1.36 | 1.89 | 2.875 | 4.603 | 5.55 | 3.239 | 1.494 | 0.5282 | 1.99 | 4.488 |
| CD14+CD16++ (Freq. of Parent) | Control | vs. | LVL | 0.0003 | 3.915, n=10 | 1.050, n=8 | LVL | 1.55 | 2.68 | 3.915 | 8.52 | 11.9 | 5.136 | 3.36 | 1.062 | 2.733 | 7.539 |
| | | | | | | | Control | 0.39 | 0.4625 | 1.05 | 1.725 | 2.6 | 1.16 | 0.7667 | 0.2711 | 0.519 | 1.801 |
| | Control | vs. | HVL | 0.0016 | 6.350, n=9 | 1.050, n=8 | HVL | 1.45 | 1.855 | 6.35 | 7.985 | 9.93 | 5.346 | 3.137 | 1.046 | 2.934 | 7.757 |
| | | | | | | | Control | 0.39 | 0.4625 | 1.05 | 1.725 | 2.6 | 1.16 | 0.7667 | 0.2711 | 0.519 | 1.801 |
| CD14++CD16-CD86+ (MFI) | Control | vs. | LVL | 0.0005 | 4.174, n=10 | 4.035, n=8 | LVL | 4.065 | 4.132 | 4.174 | 4.331 | 4.434 | 4.215 | 0.1179 | 0.0373 | 4.131 | 4.3 |
| | | | | | | | Control | 3.977 | 4.001 | 4.035 | 4.078 | 4.144 | 4.042 | 0.05455 | 0.01929 | 3.996 | 4.087 |
| | Control | vs. | HVL | 0.0079 | 4.217, n=9 | 4.035, n=8 | HVL | 3.948 | 4.132 | 4.217 | 4.365 | 4.376 | 4.226 | 0.146 | 0.04867 | 4.113 | 4.338 |
| | | | | | | | Control | 3.977 | 4.001 | 4.035 | 4.078 | 4.144 | 4.042 | 0.05455 | 0.01929 | 3.996 | 4.087 |
| CD14++CD16-CD11b+ (MFI) | Control | vs. | LVL | 0.3599 | 5.165, n=10 | 5.090, n=8 | LVL | 5.005 | 5.048 | 5.165 | 5.239 | 5.318 | 5.156 | 0.1065 | 0.03368 | 5.08 | 5.232 |
| | | | | | | | Control | 5.028 | 5.063 | 5.09 | 5.171 | 5.215 | 5.109 | 0.06521 | 0.02306 | 5.055 | 5.164 |
| | Control | vs. | HVL | 0.6058 | 5.094, n=9 | 5.090, n=8 | HVL | 4.975 | 5.084 | 5.094 | 5.189 | 5.314 | 5.125 | 0.09514 | 0.03171 | 5.052 | 5.198 |
| | | | | | | | Control | 5.028 | 5.063 | 5.09 | 5.171 | 5.215 | 5.109 | 0.06521 | 0.02306 | 5.055 | 5.164 |
| CD14++CD16-CD38+ (MFI) | Control | vs. | LVL | 0.3154 | 4.795, n=10 | 4.703, n=8 | LVL | 4.518 | 4.606 | 4.795 | 4.834 | 4.998 | 4.749 | 0.1458 | 0.04611 | 4.645 | 4.854 |
| | | | | | | | Control | 4.597 | 4.661 | 4.703 | 4.744 | 4.773 | 4.698 | 0.05801 | 0.02051 | 4.65 | 4.747 |
| | Control | vs. | HVL | 0.5414 | 4.710, n=9 | 4.703, n=8 | HVL | 4.487 | 4.575 | 4.71 | 4.742 | 4.77 | 4.664 | 0.09917 | 0.03306 | 4.588 | 4.74 |
| | | | | | | | Control | 4.597 | 4.661 | 4.703 | 4.744 | 4.773 | 4.698 | 0.05801 | 0.02051 | 4.65 | 4.747 |
| CD14++CD16-CCR2+ (MFI) | Control | vs. | LVL | >0.9999 | 4.828, n=10 | 4.784, n=8 | LVL | 3.31 | 4.535 | 4.828 | 4.983 | 5.169 | 4.673 | 0.5229 | 0.1653 | 4.299 | 5.047 |
| | | | | | | | Control | 3.193 | 3.658 | 4.784 | 5.046 | 5.143 | 4.485 | 0.7746 | 0.2739 | 3.838 | 5.133 |
| | Control | vs. | HVL | 0.2766 | 4.226, n=9 | 4.784, n=8 | HVL | 3.248 | 3.27 | 4.226 | 4.872 | 5.113 | 4.085 | 0.7937 | 0.2646 | 3.475 | 4.695 |
| | | | | | | | Control | 3.193 | 3.658 | 4.784 | 5.046 | 5.143 | 4.485 | 0.7746 | 0.2739 | 3.838 | 5.133 |
| CD14++CD16+CD11b+ (MFI) | Control | vs. | LVL | 0.0266 | 4.887, n=10 | 5.072, n=8 | LVL | 4.561 | 4.811 | 4.887 | 4.984 | 5.056 | 4.874 | 0.1405 | 0.04442 | 4.774 | 4.975 |
| | | | | | | | Control | 4.868 | 4.907 | 5.072 | 5.148 | 5.211 | 5.047 | 0.1278 | 0.04517 | 4.941 | 5.154 |
| | Control | vs. | HVL | 0.0152 | 4.849, n=9 | 5.072, n=8 | HVL | 4.158 | 4.649 | 4.849 | 5.016 | 5.092 | 4.787 | 0.2888 | 0.09625 | 4.565 | 5.009 |
| | | | | | | | Control | 4.868 | 4.907 | 5.072 | 5.148 | 5.211 | 5.047 | 0.1278 | 0.04517 | 4.941 | 5.154 |
| CD14++CD16+CD86+ (MFI) | Control | vs. | LVL | 0.0005 | 4.472, n=10 | 4.342, n=8 | LVL | 4.378 | 4.409 | 4.472 | 4.562 | 4.632 | 4.488 | 0.08466 | 0.02677 | 4.428 | 4.549 |
| | | | | | | | Control | 4.107 | 4.266 | 4.342 | 4.38 | 4.446 | 4.318 | 0.102 | 0.03605 | 4.233 | 4.403 |

| | | | | | | | | | | | | | | | | | |
|---|---|---|---|---|---|---|---|---|---|---|---|---|---|---|---|---|---|
| | Control | vs. | HVL | 0.0111 | 4.557, n=9 | 4.342, n=8 | HVL | 4.276 | 4.384 | 4.557 | 4.57 | 4.712 | 4.503 | 0.1319 | 0.04397 | 4.402 | 4.605 |
| | | | | | | | Control | 4.107 | 4.266 | 4.342 | 4.38 | 4.446 | 4.318 | 0.102 | 0.03605 | 4.233 | 4.403 |
| CD14++CD16+CD38+ (MFI) | Control | vs. | LVL | 0.5726 | 4.451, n=10 | 4.446, n=8 | LVL | 4.079 | 4.222 | 4.451 | 4.464 | 4.484 | 4.359 | 0.1495 | 0.04726 | 4.252 | 4.466 |
| | | | | | | | Control | 4.23 | 4.266 | 4.446 | 4.648 | 4.745 | 4.447 | 0.1964 | 0.06945 | 4.282 | 4.611 |
| | Control | vs. | HVL | 0.0274 | 4.250, n=9 | 4.446, n=8 | HVL | 3.745 | 3.946 | 4.25 | 4.32 | 4.35 | 4.147 | 0.2216 | 0.07387 | 3.976 | 4.317 |
| | | | | | | | Control | 4.23 | 4.266 | 4.446 | 4.648 | 4.745 | 4.447 | 0.1964 | 0.06945 | 4.282 | 4.611 |
| CD14++CD16+CCR2+ (MFI) | Control | vs. | LVL | 0.0676 | 3.682, n=10 | 4.170, n=8 | LVL | 3.487 | 3.621 | 3.682 | 4.076 | 4.424 | 3.821 | 0.3079 | 0.09737 | 3.601 | 4.041 |
| | | | | | | | Control | 3.675 | 3.961 | 4.17 | 4.37 | 4.654 | 4.165 | 0.2963 | 0.1048 | 3.918 | 4.413 |
| | Control | vs. | HVL | 0.0025 | 3.688, n=9 | 4.170, n=8 | HVL | 3.4 | 3.519 | 3.688 | 3.813 | 3.917 | 3.669 | 0.1746 | 0.05821 | 3.534 | 3.803 |
| | | | | | | | Control | 3.675 | 3.961 | 4.17 | 4.37 | 4.654 | 4.165 | 0.2963 | 0.1048 | 3.918 | 4.413 |
| CD14+CD16++CD11b+ (MFI) | Control | vs. | LVL | 0.0266 | 4.887, n=10 | 5.072, n=8 | LVL | 4.561 | 4.811 | 4.887 | 4.984 | 5.056 | 4.874 | 0.1405 | 0.04442 | 4.774 | 4.975 |
| | | | | | | | Control | 4.868 | 4.907 | 5.072 | 5.148 | 5.211 | 5.047 | 0.1278 | 0.04517 | 4.941 | 5.154 |
| | Control | vs. | HVL | 0.0152 | 4.849, n=9 | 5.072, n=8 | HVL | 4.158 | 4.649 | 4.849 | 5.016 | 5.092 | 4.787 | 0.2888 | 0.09625 | 4.565 | 5.009 |
| | | | | | | | Control | 4.868 | 4.907 | 5.072 | 5.148 | 5.211 | 5.047 | 0.1278 | 0.04517 | 4.941 | 5.154 |
| CD14+CD16++CD86+ (MFI) | Control | vs. | LVL | 0.0005 | 4.472, n=10 | 4.342, n=8 | LVL | 4.378 | 4.409 | 4.472 | 4.562 | 4.632 | 4.488 | 0.08466 | 0.02677 | 4.428 | 4.549 |
| | | | | | | | Control | 4.107 | 4.266 | 4.342 | 4.38 | 4.446 | 4.318 | 0.102 | 0.03605 | 4.233 | 4.403 |
| | Control | vs. | HVL | 0.0111 | 4.557, n=9 | 4.342, n=8 | HVL | 4.276 | 4.384 | 4.557 | 4.57 | 4.712 | 4.503 | 0.1319 | 0.04397 | 4.402 | 4.605 |
| | | | | | | | Control | 4.107 | 4.266 | 4.342 | 4.38 | 4.446 | 4.318 | 0.102 | 0.03605 | 4.233 | 4.403 |
| CD14+CD16++CD38+ (MFI) | Control | vs. | LVL | 0.5726 | 4.451, n=10 | 4.446, n=8 | LVL | 4.079 | 4.222 | 4.451 | 4.464 | 4.484 | 4.359 | 0.1495 | 0.04726 | 4.252 | 4.466 |
| | | | | | | | Control | 4.23 | 4.266 | 4.446 | 4.648 | 4.745 | 4.447 | 0.1964 | 0.06945 | 4.282 | 4.611 |
| | Control | vs. | HVL | 0.0274 | 4.250, n=9 | 4.446, n=8 | HVL | 3.745 | 3.946 | 4.25 | 4.32 | 4.35 | 4.147 | 0.2216 | 0.07387 | 3.976 | 4.317 |
| | | | | | | | Control | 4.23 | 4.266 | 4.446 | 4.648 | 4.745 | 4.447 | 0.1964 | 0.06945 | 4.282 | 4.611 |
| CD14+CD16++CCR2+ (MFI) | Control | vs. | LVL | 0.0676 | 3.682, n=10 | 4.170, n=8 | LVL | 3.487 | 3.621 | 3.682 | 4.076 | 4.424 | 3.821 | 0.3079 | 0.09737 | 3.601 | 4.041 |
| | | | | | | | Control | 3.675 | 3.961 | 4.17 | 4.37 | 4.654 | 4.165 | 0.2963 | 0.1048 | 3.918 | 4.413 |
| | Control | vs. | HVL | 0.0025 | 3.688, n=9 | 4.170, n=8 | HVL | 3.4 | 3.519 | 3.688 | 3.813 | 3.917 | 3.669 | 0.1746 | 0.05821 | 3.534 | 3.803 |
| | | | | | | | Control | 3.675 | 3.961 | 4.17 | 4.37 | 4.654 | 4.165 | 0.2963 | 0.1048 | 3.918 | 4.413 |
| CD3+CD4+CD27+ (Freq. of Parent) | Control | vs. | LVL | 0.0283 | 93.85, n=10 | 89.40, n=8 | LVL | 80.9 | 89.18 | 93.85 | 95.55 | 97.7 | 91.99 | 5.415 | 1.712 | 88.12 | 95.86 |
| | | | | | | | Control | 69.1 | 78.35 | 89.4 | 91.03 | 93.4 | 85.58 | 8.576 | 3.032 | 78.41 | 92.74 |
| | Control | vs. | HVL | 0.0206 | 92.10, n=9 | 89.40, n=8 | HVL | 90.4 | 90.9 | 92.1 | 95.9 | 96.7 | 92.93 | 2.524 | 0.8413 | 90.99 | 94.87 |
| | | | | | | | Control | 69.1 | 78.35 | 89.4 | 91.03 | 93.4 | 85.58 | 8.576 | 3.032 | 78.41 | 92.74 |
| CD3+CD4+CCR5+ (MFI) | Control | vs. | LVL | 0.0676 | 2.762, n=10 | 2.960, n=8 | LVL | 2.241 | 2.636 | 2.762 | 2.797 | 3.088 | 2.713 | 0.2165 | 0.06847 | 2.558 | 2.868 |
| | | | | | | | Control | 2.61 | 2.744 | 2.96 | 3.023 | 3.203 | 2.911 | 0.1877 | 0.06637 | 2.755 | 3.068 |
| | Control | vs. | HVL | 0.0433 | 2.792, n=9 | 2.960, n=8 | HVL | 2.551 | 2.655 | 2.792 | 2.81 | 2.875 | 2.747 | 0.1035 | 0.03448 | 2.667 | 2.826 |
| | | | | | | | Control | 2.61 | 2.744 | 2.96 | 3.023 | 3.203 | 2.911 | 0.1877 | 0.06637 | 2.755 | 3.068 |
| CD3+CD4+CD57+ (MFI) | Control | vs. | LVL | 0.0016 | 2.741, n=9 | 3.055, n=8 | LVL | 2.594 | 2.704 | 2.741 | 2.851 | 2.961 | 2.776 | 0.1101 | 0.03669 | 2.691 | 2.86 |
| | | | | | | | Control | 2.829 | 2.956 | 3.055 | 3.136 | 3.237 | 3.045 | 0.1261 | 0.04457 | 2.94 | 3.15 |
| | Control | vs. | HVL | 0.1139 | 2.914, n=9 | 3.055, n=8 | HVL | 2.152 | 2.729 | 2.914 | 3.039 | 3.115 | 2.841 | 0.2968 | 0.09894 | 2.613 | 3.069 |
| | | | | | | | Control | 2.829 | 2.956 | 3.055 | 3.136 | 3.237 | 3.045 | 0.1261 | 0.04457 | 2.94 | 3.15 |
| CD3+CD8+GLUT1+ (Freq. of Parent) | Control | vs. | LVL | 0.0343 | 91.50, n=10 | 88.05, n=8 | LVL | 87.2 | 88.33 | 91.5 | 94.33 | 95.5 | 91.4 | 2.941 | 0.93 | 89.3 | 93.5 |
| | | | | | | | Control | 82.7 | 84.25 | 88.05 | 90.4 | 91.8 | 87.53 | 3.323 | 1.175 | 84.75 | 90.3 |
| | Control | vs. | HVL | 0.0161 | 92.00, n=9 | 88.05, n=8 | HVL | 86.7 | 89.45 | 92 | 92.55 | 94 | 91.08 | 2.234 | 0.7448 | 89.36 | 92.8 |
| | | | | | | | Control | 82.7 | 84.25 | 88.05 | 90.4 | 91.8 | 87.53 | 3.323 | 1.175 | 84.75 | 90.3 |
| CD56+HLA-DR+ (MFI) | Control | vs. | LVL | 0.0062 | 3.753, n=10 | 3.600, n=8 | LVL | 3.675 | 3.712 | 3.753 | 3.802 | 4.015 | 3.78 | 0.1063 | 0.03361 | 3.704 | 3.856 |
| | | | | | | | Control | 3.419 | 3.459 | 3.6 | 3.693 | 3.768 | 3.588 | 0.1276 | 0.04511 | 3.482 | 3.695 |
| | Control | vs. | HVL | 0.0152 | 3.903, n=9 | 3.600, n=8 | HVL | 3.496 | 3.68 | 3.903 | 3.955 | 4 | 3.814 | 0.1708 | 0.05692 | 3.683 | 3.945 |
| | | | | | | | Control | 3.419 | 3.459 | 3.6 | 3.693 | 3.768 | 3.588 | 0.1276 | 0.04511 | 3.482 | 3.695 |
| CD56+CD57+ (Freq. of Parent) | Control | vs. | LVL | 0.0031 | 35.90, n=10 | 57.05, n=8 | LVL | 18.1 | 23.4 | 35.9 | 38.73 | 45.1 | 32.63 | 8.861 | 2.802 | 26.29 | 38.97 |
| | | | | | | | Control | 22.9 | 47.78 | 57.05 | 71.8 | 76.6 | 56.06 | 17.19 | 6.077 | 41.69 | 70.43 |
| | Control | vs. | HVL | 0.0274 | 33.20, n=9 | 57.05, n=8 | HVL | 19.4 | 30.15 | 33.2 | 44.75 | 61 | 36.7 | 12.66 | 4.218 | 26.97 | 46.43 |
| | | | | | | | Control | 22.9 | 47.78 | 57.05 | 71.8 | 76.6 | 56.06 | 17.19 | 6.077 | 41.69 | 70.43 |
| CD56+CD86+ (MFI) | Control | vs. | LVL | 0.5576 | 2.734, n=10 | 2.683, n=8 | LVL | 2.473 | 2.623 | 2.734 | 2.802 | 2.892 | 2.71 | 0.1319 | 0.04172 | 2.616 | 2.804 |
| | | | | | | | Control | 2.554 | 2.624 | 2.683 | 2.755 | 2.799 | 2.686 | 0.07972 | 0.02819 | 2.619 | 2.752 |
| | Control | vs. | HVL | 0.4807 | 2.678, n=9 | 2.683, n=8 | HVL | 2.58 | 2.646 | 2.678 | 2.885 | 3.025 | 2.754 | 0.1504 | 0.05013 | 2.639 | 2.87 |
| | | | | | | | Control | 2.554 | 2.624 | 2.683 | 2.755 | 2.799 | 2.686 | 0.07972 | 0.02819 | 2.619 | 2.752 |

| | | | | | | | | | | | | | | | | | |
|---|---|---|---|---|---|---|---|---|---|---|---|---|---|---|---|---|---|
| CD19+CCR7+ (MFI) | Contr ol | vs . | LVL | 0.000 8 | 3.569, n= 11 | 3.736, n =8 | LVL | 3.281 | 3.419 | 3.569 | 3.662 | 3.731 | 3.54 3 | 0.1421 | 0.042 83 | 3.44 7 | 3.63 8 |
| | | | | | | | Control | 3.656 | 3.712 | 3.736 | 3.857 | 3.894 | 3.76 7 | 0.0842 6 | 0.029 79 | 3.69 6 | 3.83 7 |
| | Contr ol | vs . | HVL | <0.00 01 | 3.502, n= 9 | 3.736, n =8 | HVL | 3.321 | 3.391 | 3.502 | 3.552 | 3.581 | 3.47 7 | 0.0898 5 | 0.029 95 | 3.40 8 | 3.54 6 |
| | | | | | | | Control | 3.656 | 3.712 | 3.736 | 3.857 | 3.894 | 3.76 7 | 0.0842 6 | 0.029 79 | 3.69 6 | 3.83 7 |
