## Supplemental Table 6 for "Chronic opioid-associated immune dysregulation among people living with HIV"

**Supplementary Table 6.** CD14++CD16- Flow cytometry results

| Marker | Comparison (A vs B) | Group A | | | Group B | | | Viral Load adjusted Group A – Group B | | | |
|---|---|---|---|---|---|---|---|---|---|---|---|
| | | n | Mean | SD | N | Mean | SD | Estimate | SE | p | adj p |
| CD14++CD16- \| Geometric Mean (APC-Fire 750-A :: CD36) | Control vs. CTN-067 M0 | 8 | 164406.38 | 61409.79 | 11 | 180063.91 | 47891.64 | -198.43 | 26948.00 | 0.9942 | 1.0000 |
| | Control vs. CTN-067 M3 | 8 | 164406.38 | 61409.79 | 11 | 197087.00 | 48773.33 | -17222.00 | 26948.00 | 0.5300 | 0.9181 |
| | Control vs. CTN-067 M6 | 8 | 164406.38 | 61409.79 | 11 | 193164.27 | 52305.61 | -13299.00 | 26948.00 | 0.6270 | 0.9596 |
| | CTN-067 M0 vs. CTN-067 M3 | 11 | 180063.91 | 47891.64 | 11 | 197087.00 | 48773.33 | -17023.00 | 9933.16 | 0.1020 | 0.3429 |
| | CTN-067 M0 vs. CTN-067 M6 | 11 | 180063.91 | 47891.64 | 11 | 193164.27 | 52305.61 | -13100.00 | 9933.16 | 0.2021 | 0.5622 |
| | CTN-067 M3 vs. CTN-067 M6 | 11 | 197087.00 | 48773.33 | 11 | 193164.27 | 52305.61 | 3922.73 | 9933.16 | 0.6971 | 0.9785 |
| CD14++CD16- \| Geometric Mean (Alexa Fluor 647-A :: GLUT1) | Control vs. CTN-067 M0 | 8 | 411.68 | 403.01 | 11 | 417.26 | 563.37 | 74.67 | 243.52 | 0.7623 | 0.9897 |
| | Control vs. CTN-067 M3 | 8 | 411.68 | 403.01 | 11 | 295.00 | 480.01 | 196.94 | 243.52 | 0.4282 | 0.8496 |
| | Control vs. CTN-067 M6 | 8 | 411.68 | 403.01 | 11 | 336.24 | 463.80 | 155.70 | 243.52 | 0.5298 | 0.9180 |
| | CTN-067 M0 vs. CTN-067 M3 | 11 | 417.26 | 563.37 | 11 | 295.00 | 480.01 | 122.26 | 86.46 | 0.1727 | 0.5055 |
| | CTN-067 M0 vs. CTN-067 M6 | 11 | 417.26 | 563.37 | 11 | 336.24 | 463.80 | 81.03 | 86.46 | 0.3598 | 0.7855 |
| | CTN-067 M3 vs. CTN-067 M6 | 11 | 295.00 | 480.01 | 11 | 336.24 | 463.80 | -41.24 | 86.46 | 0.6386 | 0.9633 |
| CD14++CD16- \| Geometric Mean (BUV615-A :: CD11b) | Control vs. CTN-067 M0 | 8 | 129931.63 | 20061.34 | 11 | 141049.00 | 32019.84 | -5469.58 | 16537.00 | 0.7443 | 0.9871 |
| | Control vs. CTN-067 M3 | 8 | 129931.63 | 20061.34 | 11 | 142731.55 | 39330.44 | -7152.13 | 16537.00 | 0.6700 | 0.9722 |
| | Control vs. CTN-067 M6 | 8 | 129931.63 | 20061.34 | 11 | 139891.55 | 38102.99 | -4312.13 | 16537.00 | 0.7969 | 0.9936 |
| | CTN-067 M0 vs. CTN-067 M3 | 11 | 141049.00 | 32019.84 | 11 | 142731.55 | 39330.44 | -1682.55 | 7779.03 | 0.8310 | 0.9963 |

| | | | | | | | | | | |
|---|---|---|---|---|---|---|---|---|---|---|
| | CTN-067 M0 vs. CTN-067 M6 | 11 | 141049.00 | 32019.84 | 11 | 139891.55 | 38102.99 | 1157.45 | 7779.03 | 0.8832 | 0.9988 |
| | CTN-067 M3 vs. CTN-067 M6 | 11 | 142731.55 | 39330.44 | 11 | 139891.55 | 38102.99 | 2840.00 | 7779.03 | 0.7189 | 0.9829 |
| CD14++CD16- \| Geometric Mean (BUV661-A :: CD27) | Control vs. CTN-067 M0 | 8 | 1446.13 | 528.48 | 11 | 1566.91 | 786.77 | -162.42 | 321.75 | 0.6192 | 0.9570 |
| | Control vs. CTN-067 M3 | 8 | 1446.13 | 528.48 | 11 | 1622.00 | 686.10 | -217.52 | 321.75 | 0.5068 | 0.9049 |
| | Control vs. CTN-067 M6 | 8 | 1446.13 | 528.48 | 11 | 1691.45 | 447.22 | -286.97 | 321.75 | 0.3830 | 0.8091 |
| | CTN-067 M0 vs. CTN-067 M3 | 11 | 1566.91 | 786.77 | 11 | 1622.00 | 686.10 | -55.09 | 147.31 | 0.7124 | 0.9817 |
| | CTN-067 M0 vs. CTN-067 M6 | 11 | 1566.91 | 786.77 | 11 | 1691.45 | 447.22 | -124.55 | 147.31 | 0.4079 | 0.8322 |
| | CTN-067 M3 vs. CTN-067 M6 | 11 | 1622.00 | 686.10 | 11 | 1691.45 | 447.22 | -69.45 | 147.31 | 0.6424 | 0.9645 |
| CD14++CD16- \| Geometric Mean (BV510-A :: CCR7) | Control vs. CTN-067 M0 | 8 | 60.23 | 890.94 | 11 | -825.29 | 732.04 | 870.17 | 418.32 | 0.0506 | 0.1936 |
| | Control vs. CTN-067 M3 | 8 | 60.23 | 890.94 | 11 | -390.18 | 887.93 | 435.06 | 418.32 | 0.3107 | 0.7285 |
| | Control vs. CTN-067 M6 | 8 | 60.23 | 890.94 | 11 | -228.52 | 621.23 | 273.40 | 418.32 | 0.5208 | 0.9131 |
| | CTN-067 M0 vs. CTN-067 M3 | 11 | -825.29 | 732.04 | 11 | -390.18 | 887.93 | -435.11 | 188.51 | 0.0318 | 0.1297 |
| | CTN-067 M0 vs. CTN-067 M6 | 11 | -825.29 | 732.04 | 11 | -228.52 | 621.23 | -596.77 | 188.51 | 0.0049 | 0.0231 |
| | CTN-067 M3 vs. CTN-067 M6 | 11 | -390.18 | 887.93 | 11 | -228.52 | 621.23 | -161.66 | 188.51 | 0.4013 | 0.8263 |
| CD14++CD16- \| Geometric Mean (BV605-A :: CD45RA) | Control vs. CTN-067 M0 | 8 | 7021.88 | 3019.31 | 11 | 11558.00 | 6706.88 | -4302.05 | 5911911.00 | 0.9994 | 1.0000 |
| | Control vs. CTN-067 M3 | 8 | 7021.88 | 3019.31 | 11 | 11098.82 | 6751.94 | -3842.87 | 5911911.00 | 0.9995 | 1.0000 |
| | Control vs. CTN-067 M6 | 8 | 7021.88 | 3019.31 | 11 | 10345.27 | 6093.99 | -3089.32 | 5911911.00 | 0.9996 | 1.0000 |
| | CTN-067 M0 vs. | 11 | 11558.00 | 6706.88 | 11 | 11098.82 | 6751.94 | 459.18 | 763.24 | 0.5542 | 0.9304 |

| | | | | | | | | | | |
|---|---|---|---|---|---|---|---|---|---|---|
| | CTN-067 M3 | | | | | | | | | |
| | CTN-067 M0 vs. CTN-067 M6 | 11 | 11558.00 | 6706.88 | 11 | 10345.27 | 6093.99 | 1212.73 | 763.24 | 0.1278 | 0.4070 |
| | CTN-067 M3 vs. CTN-067 M6 | 11 | 11098.82 | 6751.94 | 11 | 10345.27 | 6093.99 | 753.55 | 763.24 | 0.3353 | 0.7582 |
| CD14++CD16- \| Geometric Mean (BV650-A :: CD15) | Control vs. CTN-067 M0 | 8 | 2287.13 | 800.65 | 11 | 2229.91 | 865.57 | 623.79 | 829.39 | 0.4607 | 0.8747 |
| | Control vs. CTN-067 M3 | 8 | 2287.13 | 800.65 | 11 | 3399.00 | 3249.50 | -545.30 | 829.39 | 0.5184 | 0.9117 |
| | Control vs. CTN-067 M6 | 8 | 2287.13 | 800.65 | 11 | 2725.09 | 787.20 | 128.61 | 829.39 | 0.8783 | 0.9986 |
| | CTN-067 M0 vs. CTN-067 M3 | 11 | 2229.91 | 865.57 | 11 | 3399.00 | 3249.50 | -1169.09 | 712.19 | 0.1163 | 0.3792 |
| | CTN-067 M0 vs. CTN-067 M6 | 11 | 2229.91 | 865.57 | 11 | 2725.09 | 787.20 | -495.18 | 712.19 | 0.4949 | 0.8977 |
| | CTN-067 M3 vs. CTN-067 M6 | 11 | 3399.00 | 3249.50 | 11 | 2725.09 | 787.20 | 673.91 | 712.19 | 0.3553 | 0.7806 |
| CD14++CD16- \| Geometric Mean (BV711-A :: CD86) | Control vs. CTN-067 M0 | 8 | 11087.00 | 1451.42 | 11 | 20038.00 | 4561.15 | -9498.10 | 1890.76 | <.0001 | 0.0004 |
| | Control vs. CTN-067 M3 | 8 | 11087.00 | 1451.42 | 11 | 14584.45 | 5200.03 | -4044.55 | 1890.76 | 0.0449 | 0.1750 |
| | Control vs. CTN-067 M6 | 8 | 11087.00 | 1451.42 | 11 | 15075.82 | 2626.02 | -4535.92 | 1890.76 | 0.0263 | 0.1096 |
| | CTN-067 M0 vs. CTN-067 M3 | 11 | 20038.00 | 4561.15 | 11 | 14584.45 | 5200.03 | 5453.55 | 1600.87 | 0.0028 | 0.0137 |
| | CTN-067 M0 vs. CTN-067 M6 | 11 | 20038.00 | 4561.15 | 11 | 15075.82 | 2626.02 | 4962.18 | 1600.87 | 0.0056 | 0.0267 |
| | CTN-067 M3 vs. CTN-067 M6 | 11 | 14584.45 | 5200.03 | 11 | 15075.82 | 2626.02 | -491.36 | 1600.87 | 0.7621 | 0.9897 |
| CD14++CD16- \| Geometric Mean (PE-A :: CD38) | Control vs. CTN-067 M0 | 8 | 50283.88 | 6548.84 | 11 | 52414.64 | 13009.09 | 6776.26 | 5939.17 | 0.2674 | 0.6693 |
| | Control vs. CTN-067 M3 | 8 | 50283.88 | 6548.84 | 11 | 57116.45 | 24793.10 | 2074.44 | 5939.17 | 0.7305 | 0.9849 |
| | Control vs. CTN-067 M6 | 8 | 50283.88 | 6548.84 | 11 | 48964.82 | 13234.31 | 10226.00 | 5939.17 | 0.1005 | 0.3390 |

| | | | | | | | | | | | |
|---|---|---|---|---|---|---|---|---|---|---|---|
| | CTN-067 M0 vs. CTN-067 M3 | 11 | 52414.64 | 13009.09 | 11 | 57116.45 | 24793.10 | -4701.82 | 4753.74 | 0.3344 | 0.7572 |
| | CTN-067 M0 vs. CTN-067 M6 | 11 | 52414.64 | 13009.09 | 11 | 48964.82 | 13234.31 | 3449.82 | 4753.74 | 0.4764 | 0.8857 |
| | CTN-067 M3 vs. CTN-067 M6 | 11 | 57116.45 | 24793.10 | 11 | 48964.82 | 13234.31 | 8151.64 | 4753.74 | 0.1018 | 0.3424 |
| CD14++CD16- \| Geometric Mean (R718-A :: CCR5) | Control vs. CTN-067 M0 | 8 | 29735.88 | 52561.05 | 11 | 24566.64 | 39474.95 | 6033.19 | 24288.00 | 0.8064 | 0.9944 |
| | Control vs. CTN-067 M3 | 8 | 29735.88 | 52561.05 | 11 | 24014.09 | 38747.47 | 6585.74 | 24288.00 | 0.7891 | 0.9928 |
| | Control vs. CTN-067 M6 | 8 | 29735.88 | 52561.05 | 11 | 25835.27 | 41665.08 | 4764.55 | 24288.00 | 0.8465 | 0.9972 |
| | CTN-067 M0 vs. CTN-067 M3 | 11 | 24566.64 | 39474.95 | 11 | 24014.09 | 38747.47 | 552.55 | 1498.01 | 0.7161 | 0.9824 |
| | CTN-067 M0 vs. CTN-067 M6 | 11 | 24566.64 | 39474.95 | 11 | 25835.27 | 41665.08 | -1268.64 | 1498.01 | 0.4071 | 0.8315 |
| | CTN-067 M3 vs. CTN-067 M6 | 11 | 24014.09 | 38747.47 | 11 | 25835.27 | 41665.08 | -1821.18 | 1498.01 | 0.2382 | 0.6244 |
| CD14++CD16- \| Geometric Mean (RB705-A :: CCR2) | Control vs. CTN-067 M0 | 8 | 64903.63 | 51094.06 | 11 | 49275.82 | 38676.68 | 20907.00 | 24855.00 | 0.4102 | 0.8343 |
| | Control vs. CTN-067 M3 | 8 | 64903.63 | 51094.06 | 11 | 56438.91 | 42952.89 | 13744.00 | 24855.00 | 0.5864 | 0.9447 |
| | Control vs. CTN-067 M6 | 8 | 64903.63 | 51094.06 | 11 | 59544.82 | 51366.22 | 10638.00 | 24855.00 | 0.6732 | 0.9730 |
| | CTN-067 M0 vs. CTN-067 M3 | 11 | 49275.82 | 38676.68 | 11 | 56438.91 | 42952.89 | -7163.09 | 6064.08 | 0.2514 | 0.6452 |
| | CTN-067 M0 vs. CTN-067 M6 | 11 | 49275.82 | 38676.68 | 11 | 59544.82 | 51366.22 | -10269.00 | 6064.08 | 0.1059 | 0.3530 |
| | CTN-067 M3 vs. CTN-067 M6 | 11 | 56438.91 | 42952.89 | 11 | 59544.82 | 51366.22 | -3105.91 | 6064.08 | 0.6141 | 0.9552 |
| CD14++CD16- \| Geometric Mean (RB780-A :: HLA-DR) | Control vs. CTN-067 M0 | 8 | 121744.13 | 32607.80 | 11 | 254645.64 | 114340.01 | -114937.00 | 37211.00 | 0.0058 | 0.0273 |
| | Control vs. CTN-067 M3 | 8 | 121744.13 | 32607.80 | 11 | 179223.64 | 76939.59 | -39515.00 | 37211.00 | 0.3009 | 0.7158 |

| | | | | | | | | | | | |
|---|---|---|---|---|---|---|---|---|---|---|---|
| | Control vs. CTN-067 M6 | 8 | 121744.13 | 32607.80 | 11 | 151563.09 | 61116.65 | -11854.00 | 37211.00 | 0.7534 | 0.9885 |
| | CTN-067 M0 vs. CTN-067 M3 | 11 | 254645.64 | 114340.01 | 11 | 179223.64 | 76939.59 | 75422.00 | 16889.00 | 0.0002 | 0.0012 |
| | CTN-067 M0 vs. CTN-067 M6 | 11 | 254645.64 | 114340.01 | 11 | 151563.09 | 61116.65 | 103083.00 | 16889.00 | <.0001 | <.0001 |
| | CTN-067 M3 vs. CTN-067 M6 | 11 | 179223.64 | 76939.59 | 11 | 151563.09 | 61116.65 | 27661.00 | 16889.00 | 0.1171 | 0.3812 |
| CD14++CD16- \| Geometric Mean (RY586-A :: CXCR4) | Control vs. CTN-067 M0 | 8 | 13224.75 | 4696.63 | 11 | 51590.18 | 21947.29 | -28024.00 | 14226.00 | 0.0629 | 0.2322 |
| | Control vs. CTN-067 M3 | 8 | 13224.75 | 4696.63 | 11 | 33562.91 | 41301.03 | -9996.45 | 14226.00 | 0.4903 | 0.8948 |
| | Control vs. CTN-067 M6 | 8 | 13224.75 | 4696.63 | 11 | 31731.45 | 39461.39 | -8165.00 | 14226.00 | 0.5724 | 0.9387 |
| | CTN-067 M0 vs. CTN-067 M3 | 11 | 51590.18 | 21947.29 | 11 | 33562.91 | 41301.03 | 18027.00 | 12760.00 | 0.1731 | 0.5063 |
| | CTN-067 M0 vs. CTN-067 M6 | 11 | 51590.18 | 21947.29 | 11 | 31731.45 | 39461.39 | 19859.00 | 12760.00 | 0.1353 | 0.4247 |
| | CTN-067 M3 vs. CTN-067 M6 | 11 | 33562.91 | 41301.03 | 11 | 31731.45 | 39461.39 | 1831.45 | 12760.00 | 0.8873 | 0.9989 |
