## Supplemental Table 7 for "Chronic opioid-associated immune dysregulation among people living with HIV"

**Supplementary Table 7.** CD14++CD16+ Flow cytometry results

<table>
<tr><th rowspan="3">Marker</th><th rowspan="3">Comparison (A vs B)</th><th colspan="3" rowspan="2">Group A</th><th colspan="3" rowspan="2">Group B</th><th colspan="4">Viral Load adjusted</th></tr>
<tr><th colspan="4">Group A – Group B</th></tr>
<tr><th>n</th><th>Mean</th><th>SD</th><th>N</th><th>Mean</th><th>SD</th><th>Estimate</th><th>SE</th><th>p</th><th>adj p</th></tr>
<tr><td rowspan="6">CD14++CD16+ | Geometric Mean (APC-Fire 750-A :: CD36)</td><td>Control vs. CTN-067 M0</td><td>8</td><td>120578.00</td><td>63083.89</td><td>11</td><td>90521.36</td><td>36470.34</td><td>55570.00</td><td>27896.00</td><td>0.0609</td><td>0.2257</td></tr>
<tr><td>Control vs. CTN-067 M3</td><td>8</td><td>120578.00</td><td>63083.89</td><td>11</td><td>103029.73</td><td>50150.06</td><td>34414.00</td><td>25988.00</td><td>0.2011</td><td>0.5595</td></tr>
<tr><td>Control vs. CTN-067 M6</td><td>8</td><td>120578.00</td><td>63083.89</td><td>11</td><td>109491.27</td><td>36083.48</td><td>27522.00</td><td>25918.00</td><td>0.3016</td><td>0.7160</td></tr>
<tr><td>CTN-067 M0 vs. CTN-067 M3</td><td>11</td><td>90521.36</td><td>36470.34</td><td>11</td><td>103029.73</td><td>50150.06</td><td>-21156.00</td><td>10807.00</td><td>0.0651</td><td>0.2385</td></tr>
<tr><td>CTN-067 M0 vs. CTN-067 M6</td><td>11</td><td>90521.36</td><td>36470.34</td><td>11</td><td>109491.27</td><td>36083.48</td><td>-28048.00</td><td>10939.00</td><td>0.0190</td><td>0.0815</td></tr>
<tr><td>CTN-067 M3 vs. CTN-067 M6</td><td>11</td><td>103029.73</td><td>50150.06</td><td>11</td><td>109491.27</td><td>36083.48</td><td>-6891.49</td><td>9418.34</td><td>0.4733</td><td>0.8832</td></tr>
<tr><td rowspan="6">CD14++CD16+ | Geometric Mean (Alexa Fluor 647-A :: GLUT1)</td><td>Control vs. CTN-067 M0</td><td>8</td><td>454.31</td><td>490.81</td><td>11</td><td>381.38</td><td>400.08</td><td>286.15</td><td>247.49</td><td>0.2619</td><td>0.6605</td></tr>
<tr><td>Control vs. CTN-067 M3</td><td>8</td><td>454.31</td><td>490.81</td><td>11</td><td>218.11</td><td>387.86</td><td>376.96</td><td>228.31</td><td>0.1152</td><td>0.3755</td></tr>
<tr><td>Control vs. CTN-067 M6</td><td>8</td><td>454.31</td><td>490.81</td><td>11</td><td>228.59</td><td>342.23</td><td>362.87</td><td>227.60</td><td>0.1274</td><td>0.4052</td></tr>
<tr><td>CTN-067 M0 vs. CTN-067 M3</td><td>11</td><td>381.38</td><td>400.08</td><td>11</td><td>218.11</td><td>387.86</td><td>90.81</td><td>102.22</td><td>0.3855</td><td>0.8109</td></tr>
<tr><td>CTN-067 M0 vs. CTN-067 M6</td><td>11</td><td>381.38</td><td>400.08</td><td>11</td><td>228.59</td><td>342.23</td><td>76.72</td><td>103.46</td><td>0.4674</td><td>0.8791</td></tr>
<tr><td>CTN-067 M3 vs. CTN-067 M6</td><td>11</td><td>218.11</td><td>387.86</td><td>11</td><td>228.59</td><td>342.23</td><td>-14.08</td><td>89.21</td><td>0.8762</td><td>0.9985</td></tr>
<tr><td rowspan="4">CD14++CD16+ | Geometric Mean (BUV615-A :: CD11b)</td><td>Control vs. CTN-067 M0</td><td>8</td><td>115674.13</td><td>32317.08</td><td>11</td><td>75469.91</td><td>34963.80</td><td>47422.00</td><td>19015.00</td><td>0.0220</td><td>0.0932</td></tr>
<tr><td>Control vs. CTN-067 M3</td><td>8</td><td>115674.13</td><td>32317.08</td><td>11</td><td>83875.64</td><td>34658.41</td><td>36132.00</td><td>17298.00</td><td>0.0504</td><td>0.1925</td></tr>
<tr><td>Control vs. CTN-067 M6</td><td>8</td><td>115674.13</td><td>32317.08</td><td>11</td><td>85750.09</td><td>24152.49</td><td>34114.00</td><td>17234.00</td><td>0.0624</td><td>0.2303</td></tr>
<tr><td>CTN-067 M0 vs. CTN-067 M3</td><td>11</td><td>75469.91</td><td>34963.80</td><td>11</td><td>83875.64</td><td>34658.41</td><td>-11290.00</td><td>8508.59</td><td>0.2003</td><td>0.5579</td></tr>
</table>

| | | | | | | | | | | | |
|---|---|---|---|---|---|---|---|---|---|---|---|
| | CTN-067 M0 vs. CTN-067 M6 | 11 | 75469.91 | 34963.80 | 11 | 85750.09 | 24152.49 | -133308.00 | 8610.16 | 0.1387 | 0.4315 |
| | CTN-067 M3 vs. CTN-067 M6 | 11 | 83875.64 | 34658.41 | 11 | 85750.09 | 24152.49 | -2017.84 | 7441.55 | 0.7892 | 0.9928 |
| CD14++CD16+ \| Geometric Mean (BUV661-A :: CD27) | Control vs. CTN-067 M0 | 8 | 1732.38 | 497.94 | 11 | 1235.55 | 476.74 | 624.36 | 283.80 | 0.0404 | 0.1591 |
| | Control vs. CTN-067 M3 | 8 | 1732.38 | 497.94 | 11 | 1441.36 | 574.69 | 375.11 | 266.93 | 0.1761 | 0.5114 |
| | Control vs. CTN-067 M6 | 8 | 1732.38 | 497.94 | 11 | 1475.00 | 389.48 | 339.32 | 266.32 | 0.2180 | 0.5896 |
| | CTN-067 M0 vs. CTN-067 M3 | 11 | 1235.55 | 476.74 | 11 | 1441.36 | 574.69 | -249.25 | 102.32 | 0.0249 | 0.1039 |
| | CTN-067 M0 vs. CTN-067 M6 | 11 | 1235.55 | 476.74 | 11 | 1475.00 | 389.48 | -285.05 | 103.58 | 0.0127 | 0.0564 |
| | CTN-067 M3 vs. CTN-067 M6 | 11 | 1441.36 | 574.69 | 11 | 1475.00 | 389.48 | -35.80 | 89.07 | 0.6922 | 0.9774 |
| CD14++CD16+ \| Geometric Mean (BV510-A :: CCR7) | Control vs. CTN-067 M0 | 8 | 637.18 | 736.97 | 11 | 228.36 | 806.06 | 629.09 | 550.73 | 0.2675 | 0.6688 |
| | Control vs. CTN-067 M3 | 8 | 637.18 | 736.97 | 11 | 381.45 | 1070.64 | 407.32 | 477.40 | 0.4042 | 0.8284 |
| | Control vs. CTN-067 M6 | 8 | 637.18 | 736.97 | 11 | 653.15 | 679.46 | 132.21 | 474.59 | 0.7836 | 0.9922 |
| | CTN-067 M0 vs. CTN-067 M3 | 11 | 228.36 | 806.06 | 11 | 381.45 | 1070.64 | -221.77 | 307.71 | 0.4799 | 0.8876 |
| | CTN-067 M0 vs. CTN-067 M6 | 11 | 228.36 | 806.06 | 11 | 653.15 | 679.46 | -496.88 | 311.11 | 0.1267 | 0.4037 |
| | CTN-067 M3 vs. CTN-067 M6 | 11 | 381.45 | 1070.64 | 11 | 653.15 | 679.46 | -275.11 | 272.21 | 0.3249 | 0.7452 |
| CD14++CD16+ \| Geometric Mean (BV605-A :: CD45RA) | Control vs. CTN-067 M0 | 8 | 25698.38 | 19008.70 | 11 | 38265.82 | 27096.87 | -17882.00 | 12369.00 | 0.1646 | 0.4879 |
| | Control vs. CTN-067 M3 | 8 | 25698.38 | 19008.70 | 11 | 33937.09 | 21058.75 | -11675.00 | 11181.00 | 0.3095 | 0.7262 |
| | Control vs. CTN-067 M6 | 8 | 25698.38 | 19008.70 | 11 | 31414.36 | 12394.86 | -9059.37 | 11137.00 | 0.4260 | 0.8473 |
| | CTN-067 M0 vs. | 11 | 38265.82 | 27096.87 | 11 | 33937.09 | 21058.75 | 6206.73 | 5721.98 | 0.2916 | 0.7028 |

| | | | | | | | | | | | |
|---|---|---|---|---|---|---|---|---|---|---|---|
| | CTN-067 M3 | | | | | | | | | | |
| | CTN-067 M0 vs. CTN-067 M6 | 11 | 38265.82 | 27096.87 | 11 | 31414.36 | 12394.86 | 8822.83 | 5789.79 | 0.1440 | 0.4435 |
| | CTN-067 M3 vs. CTN-067 M6 | 11 | 33937.09 | 21058.75 | 11 | 31414.36 | 12394.86 | 2616.10 | 5009.99 | 0.6076 | 0.9527 |
| CD14++CD16+ \| Geometric Mean (BV650-A :: CD15) | Control vs. CTN-067 M0 | 8 | 2678.25 | 2296.01 | 11 | 903.73 | 433.26 | 2715.76 | 1122.01 | 0.0257 | 0.1069 |
| | Control vs. CTN-067 M3 | 8 | 2678.25 | 2296.01 | 11 | 1830.91 | 2742.69 | 1534.46 | 946.02 | 0.1213 | 0.3905 |
| | Control vs. CTN-067 M6 | 8 | 2678.25 | 2296.01 | 11 | 1129.27 | 612.70 | 2223.47 | 939.18 | 0.0287 | 0.1179 |
| | CTN-067 M0 vs. CTN-067 M3 | 11 | 903.73 | 433.26 | 11 | 1830.91 | 2742.69 | -1181.29 | 868.76 | 0.1898 | 0.5383 |
| | CTN-067 M0 vs. CTN-067 M6 | 11 | 903.73 | 433.26 | 11 | 1129.27 | 612.70 | -492.29 | 874.58 | 0.5801 | 0.9418 |
| | CTN-067 M3 vs. CTN-067 M6 | 11 | 1830.91 | 2742.69 | 11 | 1129.27 | 612.70 | 689.00 | 809.54 | 0.4053 | 0.8294 |
| CD14++CD16+ \| Geometric Mean (BV711-A :: CD86) | Control vs. CTN-067 M0 | 8 | 21262.63 | 4491.24 | 11 | 35629.09 | 7441.43 | -13434.00 | 4196.36 | 0.0047 | 0.0223 |
| | Control vs. CTN-067 M3 | 8 | 21262.63 | 4491.24 | 11 | 27779.27 | 8730.59 | -6413.24 | 3535.43 | 0.0855 | 0.2976 |
| | Control vs. CTN-067 M6 | 8 | 21262.63 | 4491.24 | 11 | 27883.73 | 4537.98 | -6558.93 | 3509.73 | 0.0771 | 0.2740 |
| | CTN-067 M0 vs. CTN-067 M3 | 11 | 35629.09 | 7441.43 | 11 | 27779.27 | 8730.59 | 7020.42 | 3236.08 | 0.0429 | 0.1678 |
| | CTN-067 M0 vs. CTN-067 M6 | 11 | 35629.09 | 7441.43 | 11 | 27883.73 | 4537.98 | 6874.73 | 3258.03 | 0.0483 | 0.1857 |
| | CTN-067 M3 vs. CTN-067 M6 | 11 | 27779.27 | 8730.59 | 11 | 27883.73 | 4537.98 | -145.69 | 3012.75 | 0.9619 | 1.0000 |
| CD14++CD16+ \| Geometric Mean (PE-A :: CD38) | Control vs. CTN-067 M0 | 8 | 30707.13 | 14843.25 | 11 | 19712.91 | 7679.75 | 16410.00 | 7022.26 | 0.0305 | 0.1247 |
| | Control vs. CTN-067 M3 | 8 | 30707.13 | 14843.25 | 11 | 23592.82 | 16190.90 | 12143.00 | 6251.07 | 0.0670 | 0.2443 |
| | Control vs. CTN-067 M6 | 8 | 30707.13 | 14843.25 | 11 | 20327.45 | 7913.07 | 15389.00 | 6221.99 | 0.0230 | 0.0968 |

| | | | | | | | | | | | |
|---|---|---|---|---|---|---|---|---|---|---|---|
| | CTN-067 M0 vs. CTN-067 M3 | 11 | 19712.91 | 7679.75 | 11 | 23592.82 | 16190.90 | -4266.88 | 3496.28 | 0.2372 | 0.6220 |
| | CTN-067 M0 vs. CTN-067 M6 | 11 | 19712.91 | 7679.75 | 11 | 20327.45 | 7913.07 | -1020.75 | 3536.89 | 0.7760 | 0.9913 |
| | CTN-067 M3 vs. CTN-067 M6 | 11 | 23592.82 | 16190.90 | 11 | 20327.45 | 7913.07 | 3246.12 | 3070.61 | 0.3037 | 0.7188 |
| CD14++CD16+ \| Geometric Mean (R718-A :: CCR5) | Control vs. CTN-067 M0 | 8 | 1372.50 | 619.00 | 11 | 1874.64 | 1106.37 | 380.78 | 586.84 | 0.5242 | 0.9146 |
| | Control vs. CTN-067 M3 | 8 | 1372.50 | 619.00 | 11 | 1872.45 | 1235.01 | 53.59 | 545.06 | 0.9227 | 0.9996 |
| | Control vs. CTN-067 M6 | 8 | 1372.50 | 619.00 | 11 | 2128.73 | 1482.08 | -219.06 | 543.52 | 0.6914 | 0.9772 |
| | CTN-067 M0 vs. CTN-067 M3 | 11 | 1874.64 | 1106.37 | 11 | 1872.45 | 1235.01 | -327.19 | 232.04 | 0.1747 | 0.5086 |
| | CTN-067 M0 vs. CTN-067 M6 | 11 | 1874.64 | 1106.37 | 11 | 2128.73 | 1482.08 | -599.84 | 234.87 | 0.0194 | 0.0831 |
| | CTN-067 M3 vs. CTN-067 M6 | 11 | 1872.45 | 1235.01 | 11 | 2128.73 | 1482.08 | -272.65 | 202.31 | 0.1936 | 0.5455 |
| CD14++CD16+ \| Geometric Mean (RB705-A :: CCR2) | Control vs. CTN-067 M0 | 8 | 17885.75 | 12689.68 | 11 | 6273.27 | 4251.56 | 14070.00 | 5237.55 | 0.0146 | 0.0642 |
| | Control vs. CTN-067 M3 | 8 | 17885.75 | 12689.68 | 11 | 9280.09 | 8292.02 | 10654.00 | 4947.09 | 0.0443 | 0.1724 |
| | Control vs. CTN-067 M6 | 8 | 17885.75 | 12689.68 | 11 | 9894.27 | 6880.66 | 10020.00 | 4936.48 | 0.0566 | 0.2123 |
| | CTN-067 M0 vs. CTN-067 M3 | 11 | 6273.27 | 4251.56 | 11 | 9280.09 | 8292.02 | -3415.70 | 1823.03 | 0.0764 | 0.2720 |
| | CTN-067 M0 vs. CTN-067 M6 | 11 | 6273.27 | 4251.56 | 11 | 9894.27 | 6880.66 | -4050.21 | 1845.52 | 0.0408 | 0.1606 |
| | CTN-067 M3 vs. CTN-067 M6 | 11 | 9280.09 | 8292.02 | 11 | 9894.27 | 6880.66 | -634.51 | 1586.06 | 0.6936 | 0.9777 |
| CD14++CD16+ \| Geometric Mean (RB780-A :: HLA-DR) | Control vs. CTN-067 M0 | 8 | 312674.75 | 84547.50 | 11 | 509617.73 | 148884.59 | -28358.00 | 78974.00 | 0.7235 | 0.9837 |
| | Control vs. CTN-067 M3 | 8 | 312674.75 | 84547.50 | 11 | 497422.73 | 186122.60 | -68578.00 | 70762.00 | 0.3447 | 0.7683 |

| | | | | | | | | | | | |
|---|---|---|---|---|---|---|---|---|---|---|---|
| | Control vs. CTN-067 M6 | 8 | 312674.75 | 84547.50 | 11 | 463488.45 | 155182.02 | -37250.00 | 70454.00 | 0.6031 | 0.9510 |
| | CTN-067 M0 vs. CTN-067 M3 | 11 | 509617.73 | 148884.59 | 11 | 497422.73 | 186122.60 | -40221.00 | 38142.00 | 0.3049 | 0.7203 |
| | CTN-067 M0 vs. CTN-067 M6 | 11 | 509617.73 | 148884.59 | 11 | 463488.45 | 155182.02 | -8892.48 | 38589.00 | 0.8202 | 0.9955 |
| | CTN-067 M3 vs. CTN-067 M6 | 11 | 497422.73 | 186122.60 | 11 | 463488.45 | 155182.02 | 31328.00 | 33452.00 | 0.3608 | 0.7858 |
| CD14++CD16+ \| Geometric Mean (RY586-A :: CXCR4) | Control vs. CTN-067 M0 | 8 | 15195.00 | 8281.62 | 11 | 35358.64 | 16565.35 | 7924.45 | 13292.00 | 0.5581 | 0.9320 |
| | Control vs. CTN-067 M3 | 8 | 15195.00 | 8281.62 | 11 | 25627.82 | 27244.27 | 8616.02 | 11166.00 | 0.4498 | 0.8662 |
| | Control vs. CTN-067 M6 | 8 | 15195.00 | 8281.62 | 11 | 29777.00 | 31579.44 | 4017.43 | 11084.00 | 0.7210 | 0.9832 |
| | CTN-067 M0 vs. CTN-067 M3 | 11 | 35358.64 | 16565.35 | 11 | 25627.82 | 27244.27 | 691.57 | 10078.00 | 0.9460 | 0.9999 |
| | CTN-067 M0 vs. CTN-067 M6 | 11 | 35358.64 | 16565.35 | 11 | 29777.00 | 31579.44 | -3907.02 | 10149.00 | 0.7045 | 0.9800 |
| | CTN-067 M3 vs. CTN-067 M6 | 11 | 25627.82 | 27244.27 | 11 | 29777.00 | 31579.44 | -4598.59 | 9346.72 | 0.6284 | 0.9599 |
