## Supplemental Table 8 for "Chronic opioid-associated immune dysregulation among people living with HIV"

**Supplementary Table 8.** CD14+CD16++ Flow Data

| Marker | Comparison (A vs B) | Group A | | | Group B | | | Viral Load adjusted Group A – Group B | | | |
|---|---|---|---|---|---|---|---|---|---|---|---|
| | | n | Mean | SD | N | Mean | SD | Estimate | SE | p | adj p |
| CD14+CD16++ \| Geometric Mean (APC-Fire 750-A :: CD36) | Control vs. CTN-067 M0 | 8 | 12058.75 | 10640.40 | 11 | 12747.73 | 5140.79 | 3157.31 | 4338.98 | 0.4757 | 0.8848 |
| | Control vs. CTN-067 M3 | 8 | 12058.75 | 10640.40 | 11 | 11711.27 | 6310.91 | 2356.95 | 4138.11 | 0.5756 | 0.9399 |
| | Control vs. CTN-067 M6 | 8 | 12058.75 | 10640.40 | 11 | 10227.82 | 4747.90 | 3749.08 | 4130.81 | 0.3755 | 0.8010 |
| | CTN-067 M0 vs. CTN-067 M3 | 11 | 12747.73 | 5140.79 | 11 | 11711.27 | 6310.91 | -800.36 | 1378.48 | 0.5683 | 0.9367 |
| | CTN-067 M0 vs. CTN-067 M6 | 11 | 12747.73 | 5140.79 | 11 | 10227.82 | 4747.90 | 591.78 | 1395.61 | 0.6763 | 0.9737 |
| | CTN-067 M3 vs. CTN-067 M6 | 11 | 11711.27 | 6310.91 | 11 | 10227.82 | 4747.90 | 1392.13 | 1198.00 | 0.2596 | 0.6570 |
| CD14+CD16++ \| Geometric Mean (Alexa Fluor 647-A :: GLUT1) | Control vs. CTN-067 M0 | 8 | -3.48 | 451.77 | 11 | 4.12 | 333.18 | -29.77 | 234.55 | 0.9003 | 0.9992 |
| | Control vs. CTN-067 M3 | 8 | -3.48 | 451.77 | 11 | -96.88 | 356.84 | 37.52 | 213.51 | 0.8624 | 0.9980 |
| | Control vs. CTN-067 M6 | 8 | -3.48 | 451.77 | 11 | -226.57 | 428.95 | 165.54 | 212.73 | 0.4460 | 0.8633 |
| | CTN-067 M0 vs. CTN-067 M3 | 11 | 4.12 | 333.18 | 11 | -96.88 | 356.84 | 67.29 | 104.60 | 0.5277 | 0.9166 |
| | CTN-067 M0 vs. CTN-067 M6 | 11 | 4.12 | 333.18 | 11 | -226.57 | 428.95 | 195.31 | 105.85 | 0.0807 | 0.2840 |
| | CTN-067 M3 vs. CTN-067 M6 | 11 | -96.88 | 356.84 | 11 | -226.57 | 428.95 | 128.02 | 91.47 | 0.1778 | 0.5148 |
| CD14+CD16++ \| Geometric Mean (BUV615-A :: CD11b) | Control vs. CTN-067 M0 | 8 | 14380.50 | 4680.30 | 11 | 10040.09 | 6099.98 | 5203.33 | 3184.74 | 0.1188 | 0.3844 |
| | Control vs. CTN-067 M3 | 8 | 14380.50 | 4680.30 | 11 | 9850.55 | 7106.93 | 5748.03 | 3047.51 | 0.0747 | 0.2668 |
| | Control vs. CTN-067 M6 | 8 | 14380.50 | 4680.30 | 11 | 10257.00 | 6099.91 | 5359.23 | 3042.54 | 0.0942 | 0.3215 |
| | CTN-067 M0 vs. CTN-067 M3 | 11 | 10040.09 | 6099.98 | 11 | 9850.55 | 7106.93 | 544.70 | 975.87 | 0.5832 | 0.9432 |

| | | | | | | | | | | | |
|---|---|---|---|---|---|---|---|---|---|---|---|
| | CTN-067 M0 vs. CTN-067 M6 | 11 | 10040.09 | 6099.98 | 11 | 10257.00 | 6099.91 | 155.90 | 988.02 | 0.8763 | 0.9985 |
| | CTN-067 M3 vs. CTN-067 M6 | 11 | 9850.55 | 7106.93 | 11 | 10257.00 | 6099.91 | -388.80 | 847.80 | 0.6517 | 0.9671 |
| CD14+CD16++ \| Geometric Mean (BUV661-A :: CD27) | Control vs. CTN-067 M0 | 8 | 699.43 | 411.84 | 11 | 597.00 | 269.06 | 154.40 | 191.71 | 0.4306 | 0.8511 |
| | Control vs. CTN-067 M3 | 8 | 699.43 | 411.84 | 11 | 740.36 | 255.77 | -25.46 | 177.79 | 0.8876 | 0.9989 |
| | Control vs. CTN-067 M6 | 8 | 699.43 | 411.84 | 11 | 672.67 | 294.75 | 40.41 | 177.27 | 0.8221 | 0.9957 |
| | CTN-067 M0 vs. CTN-067 M3 | 11 | 597.00 | 269.06 | 11 | 740.36 | 255.77 | -179.87 | 76.59 | 0.0298 | 0.1221 |
| | CTN-067 M0 vs. CTN-067 M6 | 11 | 597.00 | 269.06 | 11 | 672.67 | 294.75 | -113.99 | 77.52 | 0.1578 | 0.4736 |
| | CTN-067 M3 vs. CTN-067 M6 | 11 | 740.36 | 255.77 | 11 | 672.67 | 294.75 | 65.88 | 66.79 | 0.3363 | 0.7588 |
| CD14+CD16++ \| Geometric Mean (BV510-A :: CCR7) | Control vs. CTN-067 M0 | 8 | 1144.63 | 506.39 | 11 | 521.76 | 517.51 | 829.05 | 371.96 | 0.0381 | 0.1513 |
| | Control vs. CTN-067 M3 | 8 | 1144.63 | 506.39 | 11 | 764.96 | 569.44 | 451.85 | 317.72 | 0.1712 | 0.5015 |
| | Control vs. CTN-067 M6 | 8 | 1144.63 | 506.39 | 11 | 1045.98 | 718.61 | 164.17 | 315.63 | 0.6090 | 0.9532 |
| | CTN-067 M0 vs. CTN-067 M3 | 11 | 521.76 | 517.51 | 11 | 764.96 | 569.44 | -377.20 | 221.62 | 0.1051 | 0.3500 |
| | CTN-067 M0 vs. CTN-067 M6 | 11 | 521.76 | 517.51 | 11 | 1045.98 | 718.61 | -664.88 | 223.97 | 0.0079 | 0.0363 |
| | CTN-067 M3 vs. CTN-067 M6 | 11 | 764.96 | 569.44 | 11 | 1045.98 | 718.61 | -287.68 | 197.23 | 0.1610 | 0.4805 |
| CD14+CD16++ \| Geometric Mean (BV605-A :: CD45RA) | Control vs. CTN-067 M0 | 8 | 69496.50 | 57346.71 | 11 | 77865.09 | 52404.36 | -46447.00 | 29830.00 | 0.1360 | 0.4252 |
| | Control vs. CTN-067 M3 | 8 | 69496.50 | 57346.71 | 11 | 77252.82 | 63793.08 | -37884.00 | 26888.00 | 0.1750 | 0.5093 |
| | Control vs. CTN-067 M6 | 8 | 69496.50 | 57346.71 | 11 | 69137.55 | 40966.13 | -29374.00 | 26778.00 | 0.2864 | 0.6956 |
| | CTN-067 M0 vs. | 11 | 77865.09 | 52404.36 | 11 | 77252.82 | 63793.08 | 8562.78 | 13996.00 | 0.5479 | 0.9271 |

| | | | | | | | | | | |
|---|---|---|---|---|---|---|---|---|---|---|
| | CTN-067 M3 | | | | | | | | | |
| | CTN-067 M0 vs. CTN-067 M6 | 11 | 77865.09 | 52404.36 | 11 | 69137.55 | 40966.13 | 17073.00 | 14161.00 | 0.2428 | 0.6309 |
| | CTN-067 M3 vs. CTN-067 M6 | 11 | 77252.82 | 63793.08 | 11 | 69137.55 | 40966.13 | 8510.55 | 12261.00 | 0.4960 | 0.8981 |
| CD14+CD16++ \| Geometric Mean (BV650-A :: CD15) | Control vs. CTN-067 M0 | 8 | 8542.75 | 14460.87 | 11 | 3373.09 | 5586.65 | 6558.37 | 5923.10 | 0.2820 | 0.6895 |
| | Control vs. CTN-067 M3 | 8 | 8542.75 | 14460.87 | 11 | 3684.64 | 5305.72 | 6378.91 | 5398.33 | 0.2519 | 0.6453 |
| | Control vs. CTN-067 M6 | 8 | 8542.75 | 14460.87 | 11 | 6009.82 | 8987.36 | 4060.29 | 5378.80 | 0.4596 | 0.8735 |
| | CTN-067 M0 vs. CTN-067 M3 | 11 | 3373.09 | 5586.65 | 11 | 3684.64 | 5305.72 | -179.46 | 2624.46 | 0.9462 | 0.9999 |
| | CTN-067 M0 vs. CTN-067 M6 | 11 | 3373.09 | 5586.65 | 11 | 6009.82 | 8987.36 | -2498.08 | 2655.84 | 0.3587 | 0.7837 |
| | CTN-067 M3 vs. CTN-067 M6 | 11 | 3684.64 | 5305.72 | 11 | 6009.82 | 8987.36 | -2318.62 | 2294.63 | 0.3250 | 0.7454 |
| CD14+CD16++ \| Geometric Mean (BV711-A :: CD86) | Control vs. CTN-067 M0 | 8 | 11335.75 | 8068.84 | 11 | 15457.18 | 9271.26 | -9877.36 | 4659.12 | 0.0474 | 0.1826 |
| | Control vs. CTN-067 M3 | 8 | 11335.75 | 8068.84 | 11 | 12838.09 | 9542.85 | -6272.36 | 4128.17 | 0.1451 | 0.4460 |
| | Control vs. CTN-067 M6 | 8 | 11335.75 | 8068.84 | 11 | 11143.73 | 7080.21 | -4528.98 | 4108.10 | 0.2840 | 0.6924 |
| | CTN-067 M0 vs. CTN-067 M3 | 11 | 15457.18 | 9271.26 | 11 | 12838.09 | 9542.85 | 3605.00 | 2368.90 | 0.1445 | 0.4447 |
| | CTN-067 M0 vs. CTN-067 M6 | 11 | 15457.18 | 9271.26 | 11 | 11143.73 | 7080.21 | 5348.38 | 2396.22 | 0.0379 | 0.1504 |
| | CTN-067 M3 vs. CTN-067 M6 | 11 | 12838.09 | 9542.85 | 11 | 11143.73 | 7080.21 | 1743.38 | 2082.69 | 0.4130 | 0.8362 |
| CD14+CD16++ \| Geometric Mean (PE-A :: CD38) | Control vs. CTN-067 M0 | 8 | 6085.00 | 2065.12 | 11 | 5075.27 | 2123.05 | 2800.63 | 1065.11 | 0.0165 | 0.0718 |
| | Control vs. CTN-067 M3 | 8 | 6085.00 | 2065.12 | 11 | 4470.27 | 2069.80 | 2895.03 | 971.23 | 0.0077 | 0.0354 |
| | Control vs. CTN-067 M6 | 8 | 6085.00 | 2065.12 | 11 | 4097.27 | 1497.07 | 3242.65 | 967.74 | 0.0034 | 0.0162 |

| | | | | | | | | | | | |
|---|---|---|---|---|---|---|---|---|---|---|---|
| | CTN-067 M0 vs. CTN-067 M3 | 11 | 5075.27 | 2123.05 | 11 | 4470.27 | 2069.80 | 94.40 | 470.65 | 0.8432 | 0.9970 |
| | CTN-067 M0 vs. CTN-067 M6 | 11 | 5075.27 | 2123.05 | 11 | 4097.27 | 1497.07 | 442.01 | 476.28 | 0.3650 | 0.7903 |
| | CTN-067 M3 vs. CTN-067 M6 | 11 | 4470.27 | 2069.80 | 11 | 4097.27 | 1497.07 | 347.61 | 411.47 | 0.4087 | 0.8325 |
| CD14+CD16++ \| Geometric Mean (R718-A :: CCR5) | Control vs. CTN-067 M0 | 8 | 396.63 | 279.73 | 11 | 452.55 | 337.41 | 290.35 | 163.08 | 0.0910 | 0.3127 |
| | Control vs. CTN-067 M3 | 8 | 396.63 | 279.73 | 11 | 459.19 | 261.20 | 157.78 | 147.88 | 0.2994 | 0.7131 |
| | Control vs. CTN-067 M6 | 8 | 396.63 | 279.73 | 11 | 403.33 | 348.37 | 207.39 | 147.32 | 0.1754 | 0.5100 |
| | CTN-067 M0 vs. CTN-067 M3 | 11 | 452.55 | 337.41 | 11 | 459.19 | 261.20 | -132.57 | 74.21 | 0.0900 | 0.3101 |
| | CTN-067 M0 vs. CTN-067 M6 | 11 | 452.55 | 337.41 | 11 | 403.33 | 348.37 | -82.97 | 75.10 | 0.2830 | 0.6910 |
| | CTN-067 M3 vs. CTN-067 M6 | 11 | 459.19 | 261.20 | 11 | 403.33 | 348.37 | 49.60 | 64.94 | 0.4544 | 0.8696 |
| CD14+CD16++ \| Geometric Mean (RB705-A :: CCR2) | Control vs. CTN-067 M0 | 8 | 1593.13 | 318.58 | 11 | 1193.27 | 270.70 | 683.01 | 139.80 | 0.0001 | 0.0005 |
| | Control vs. CTN-067 M3 | 8 | 1593.13 | 318.58 | 11 | 1137.36 | 169.11 | 637.94 | 118.29 | <.0001 | 0.0002 |
| | Control vs. CTN-067 M6 | 8 | 1593.13 | 318.58 | 11 | 1152.18 | 169.72 | 618.10 | 117.45 | <.0001 | 0.0002 |
| | CTN-067 M0 vs. CTN-067 M3 | 11 | 1193.27 | 270.70 | 11 | 1137.36 | 169.11 | -45.08 | 87.26 | 0.6114 | 0.9541 |
| | CTN-067 M0 vs. CTN-067 M6 | 11 | 1193.27 | 270.70 | 11 | 1152.18 | 169.72 | -64.92 | 88.15 | 0.4704 | 0.8812 |
| | CTN-067 M3 vs. CTN-067 M6 | 11 | 1137.36 | 169.11 | 11 | 1152.18 | 169.72 | -19.84 | 78.09 | 0.8022 | 0.9941 |
| CD14+CD16++ \| Geometric Mean (RB780-A :: HLA-DR) | Control vs. CTN-067 M0 | 8 | 87223.13 | 65727.27 | 11 | 123701.82 | 97923.57 | -70245.00 | 47358.00 | 0.1544 | 0.4664 |
| | Control vs. CTN-067 M3 | 8 | 87223.13 | 65727.27 | 11 | 105093.45 | 88746.88 | -47694.00 | 42419.00 | 0.2749 | 0.6795 |

| | | | | | | | | | | | |
|---|---|---|---|---|---|---|---|---|---|---|---|
| | Control vs. CTN-067 M6 | 8 | 87223.13 | 65727.27 | 11 | 91789.55 | 73626.63 | -34194.00 | 42233.00 | 0.4282 | 0.8491 |
| | CTN-067 M0 vs. CTN-067 M3 | 11 | 123701.82 | 97923.57 | 11 | 105093.45 | 88746.88 | 22551.00 | 22911.00 | 0.3373 | 0.7599 |
| | CTN-067 M0 vs. CTN-067 M6 | 11 | 123701.82 | 97923.57 | 11 | 91789.55 | 73626.63 | 36051.00 | 23179.00 | 0.1364 | 0.4262 |
| | CTN-067 M3 vs. CTN-067 M6 | 11 | 105093.45 | 88746.88 | 11 | 91789.55 | 73626.63 | 13500.00 | 20095.00 | 0.5098 | 0.9064 |
| CD14+CD16++ \| Geometric Mean (RY586-A :: CXCR4) | Control vs. CTN-067 M0 | 8 | 4527.75 | 3069.60 | 11 | 13188.27 | 6877.01 | -6374.61 | 3056.51 | 0.0507 | 0.1935 |
| | Control vs. CTN-067 M3 | 8 | 4527.75 | 3069.60 | 11 | 6243.09 | 3190.58 | 201.88 | 2553.70 | 0.9378 | 0.9998 |
| | Control vs. CTN-067 M6 | 8 | 4527.75 | 3069.60 | 11 | 7300.00 | 4977.30 | -873.36 | 2534.06 | 0.7341 | 0.9855 |
| | CTN-067 M0 vs. CTN-067 M3 | 11 | 13188.27 | 6877.01 | 11 | 6243.09 | 3190.58 | 6576.48 | 2143.54 | 0.0063 | 0.0295 |
| | CTN-067 M0 vs. CTN-067 M6 | 11 | 13188.27 | 6877.01 | 11 | 7300.00 | 4977.30 | 5501.24 | 2161.81 | 0.0198 | 0.0846 |
| | CTN-067 M3 vs. CTN-067 M6 | 11 | 6243.09 | 3190.58 | 11 | 7300.00 | 4977.30 | -1075.24 | 1955.61 | 0.5888 | 0.9455 |
