## Supplemental Table 9 for "Chronic opioid-associated immune dysregulation among people living with HIV"

### **Supplementary Table 9.** Monocyte cytokine production

| Marker | Colum n B | Vs | Colum n A | P value | Median of Column A | Median of Column B | Compariso n | Minimu m | 25% percentil e | Median | 75% percentil e | Maximu m | Mean | Std. Deviation | Std. Error of mean | Lower 95% CL | Upper 95% CL |
|---|---|---|---|---|---|---|---|---|---|---|---|---|---|---|---|---|---|
| IL-10 | Control | vs. | LVL | 0.0549 | 95.83, n=9 | 246.0, n=7 | LVL | 0.001 | 27.91 | 95.83 | 159.7 | 441.3 | 120.5 | 132.8 | 44.26 | 18.4 | 222.5 |
| | | | | | | | Control | 85.5 | 149.3 | 246 | 367.1 | 370.9 | 250.2 | 105.9 | 40.02 | 152.3 | 348.2 |
| | Control | vs. | HVL | 0.0125 | 118.6, n=14 | 246.0, n=7 | HVL | 0 | 56.46 | 118.6 | 181.8 | 240.2 | 121.9 | 74.08 | 19.8 | 79.18 | 164.7 |
| | | | | | | | Control | 85.5 | 149.3 | 246 | 367.1 | 370.9 | 250.2 | 105.9 | 40.02 | 152.3 | 348.2 |
| IL-1β | Control | vs. | LVL | 0.0418 | 114.9, n=9 | 194.2, n=7 | LVL | 0.001 | 45.31 | 114.9 | 144.8 | 219.7 | 100 | 68.43 | 22.81 | 47.42 | 152.6 |
| | | | | | | | Control | 81.8 | 147.5 | 194.2 | 201.9 | 343.9 | 189 | 79.95 | 30.22 | 115.1 | 263 |
| | Control | vs. | HVL | 0.1101 | 142.7, n=14 | 194.2, n=7 | HVL | 0 | 104.5 | 142.7 | 167.2 | 254.1 | 136.8 | 67.08 | 17.93 | 98.03 | 175.5 |
| | | | | | | | Control | 81.8 | 147.5 | 194.2 | 201.9 | 343.9 | 189 | 79.95 | 30.22 | 115.1 | 263 |
| IL-6 | Control | vs. | LVL | 0.1416 | 1769, n=9 | 2285, n=7 | LVL | 1102 | 1551 | 1769 | 2861 | 6651 | 2445 | 1705 | 568.4 | 1134 | 3756 |
| | | | | | | | Control | 1914 | 2055 | 2285 | 5043 | 5838 | 3144 | 1603 | 606 | 1662 | 4627 |
| | Control | vs. | HVL | 0.5353 | 3407, n=14 | 2285, n=7 | HVL | 0 | 1804 | 3407 | 6005 | 11396 | 4124 | 3196 | 854 | 2279 | 5969 |
| | | | | | | | Control | 1914 | 2055 | 2285 | 5043 | 5838 | 3144 | 1603 | 606 | 1662 | 4627 |
| MIP-3α | Control | vs. | LVL | 0.0021 | 80.48, n=9 | 583.2, n=7 | LVL | 40.95 | 61.28 | 80.48 | 158.1 | 430.9 | 128.1 | 125.4 | 41.79 | 31.78 | 224.5 |
| | | | | | | | Control | 122.3 | 206.9 | 583.2 | 658 | 792.3 | 508.5 | 247.8 | 93.68 | 279.2 | 737.7 |
| | Control | vs. | HVL | 0.0159 | 126.1, n=14 | 583.2, n=7 | HVL | 6.5 | 74.55 | 126.1 | 220.7 | 648 | 187.4 | 201.9 | 53.96 | 70.79 | 303.9 |
| | | | | | | | Control | 122.3 | 206.9 | 583.2 | 658 | 792.3 | 508.5 | 247.8 | 93.68 | 279.2 | 737.7 |
| TNF-α | Control | vs. | LVL | 0.0115 | 278.3, n=9 | 1302, n=7 | LVL | 0.001 | 103.7 | 278.3 | 828.2 | 927.4 | 436.7 | 361.6 | 120.5 | 158.7 | 714.6 |
| | | | | | | | Control | 545.4 | 608.1 | 1302 | 1392 | 1639 | 1106 | 419.3 | 158.5 | 718.5 | 1494 |
| | Control | vs. | HVL | 0.0016 | 414.7, n=14 | 1302, n=7 | HVL | 0 | 179.1 | 414.7 | 633.3 | 811.8 | 422.7 | 259.6 | 69.38 | 272.8 | 572.6 |
| | | | | | | | Control | 545.4 | 608.1 | 1302 | 1392 | 1639 | 1106 | 419.3 | 158.5 | 718.5 | 1494 |
| Fraktalkin e | Control | vs. | LVL | 0.1142 | 90.82, n=9 | 142.8, n=7 | LVL | 0.001 | 12.61 | 90.82 | 166.4 | 222.5 | 89.46 | 82.65 | 27.55 | 25.93 | 153 |
| | | | | | | | Control | 85.78 | 98.22 | 142.8 | 274 | 274.2 | 168 | 78.11 | 29.52 | 95.72 | 240.2 |
| | Control | vs. | HVL | 0.0756 | 104.0, n=14 | 142.8, n=7 | HVL | 0.001 | 0.001 | 104 | 148.9 | 191.7 | 87.32 | 71.72 | 19.17 | 45.91 | 128.7 |
| | | | | | | | Control | 85.78 | 98.22 | 142.8 | 274 | 274.2 | 168 | 78.11 | 29.52 | 95.72 | 240.2 |
| GMCSF | Control | vs. | LVL | 0.4698 | 13.17, n=9 | 14.65, n=7 | LVL | 0.2 | 5.99 | 13.17 | 34.39 | 50.41 | 20.12 | 17.05 | 5.684 | 7.009 | 33.22 |
| | | | | | | | Control | 0.859 | 1.554 | 14.65 | 23.18 | 26.35 | 13.03 | 10.55 | 3.986 | 3.275 | 22.78 |
| | Control | vs. | HVL | 0.5353 | 11.58, n=14 | 14.65, n=7 | HVL | 1.203 | 3.72 | 11.58 | 30.32 | 50.05 | 17.9 | 16.9 | 4.518 | 8.142 | 27.66 |
| | | | | | | | Control | 0.859 | 1.554 | 14.65 | 23.18 | 26.35 | 13.03 | 10.55 | 3.986 | 3.275 | 22.78 |
| ITAC | Control | vs. | LVL | 0.9182 | 53.32, n=9 | 39.48, n=7 | LVL | 13.79 | 28.59 | 53.32 | 58.17 | 62.77 | 43.87 | 17.34 | 5.781 | 30.54 | 57.2 |
| | | | | | | | Control | 29.63 | 32.99 | 39.48 | 44.56 | 69.01 | 42.41 | 12.82 | 4.846 | 30.55 | 54.26 |
| | Control | vs. | HVL | 0.2545 | 32.03, n=14 | 39.48, n=7 | HVL | 0.001 | 23.65 | 32.03 | 41.85 | 71.68 | 32.3 | 18.08 | 4.831 | 21.86 | 42.73 |
| | | | | | | | Control | 29.63 | 32.99 | 39.48 | 44.56 | 69.01 | 42.41 | 12.82 | 4.846 | 30.55 | 54.26 |
| IFN-β | Control | vs. | LVL | 0.0031 | 13.48, n=9 | 0.001000, n=7 | LVL | 0.001 | 0.2034 | 13.48 | 212.3 | 778.6 | 138.6 | 267.5 | 89.17 | -67 | 344.2 |
| | | | | | | | Control | 0 | 0 | 0.001 | 0.001 | 0.001 | 0.000714 3 | 0.000488 | 0.000184 4 | 0.00026 3 | 0.001166 |
| | Control | vs. | HVL | 0.0066 | 48.31, n=14 | 0.001000, n=7 | HVL | 0.001 | 0.001 | 48.31 | 144.3 | 194.1 | 69.28 | 76.69 | 20.5 | 25 | 113.6 |
| | | | | | | | Control | 0 | 0 | 0.001 | 0.001 | 0.001 | 0.000714 3 | 0.000488 | 0.000184 4 | 0.00026 3 | 0.001166 |
| IFN-α | Control | vs. | LVL | 0.4991 | 0.001000, n=9 | 0.001000, n=7 | LVL | 0.001 | 0.001 | 0.001 | 27.93 | 106.6 | 18.29 | 37.41 | 12.47 | -10.47 | 47.05 |
| | | | | | | | Control | 0 | 0.001 | 0.001 | 5.791 | 82.28 | 12.58 | 30.81 | 11.65 | -15.91 | 41.08 |
| | Control | vs. | HVL | 0.1937 | 4.618, n=14 | 0.001000, n=7 | HVL | 0.001 | 0.001 | 4.618 | 23.95 | 141.2 | 21.29 | 39.93 | 10.67 | -1.76 | 44.35 |
| | | | | | | | Control | 0 | 0.001 | 0.001 | 5.791 | 82.28 | 12.58 | 30.81 | 11.65 | -15.91 | 41.08 |
| IFN-γ | Control | vs. | LVL | 0.0032 | 0.001000, n=9 | 33.51, n=7 | LVL | 0 | 0 | 0.001 | 19.08 | 35.61 | 8.198 | 14.11 | 4.703 | -2.648 | 19.04 |
| | | | | | | | Control | 28.03 | 29.56 | 33.51 | 82.52 | 208.3 | 64.32 | 66.28 | 25.05 | 3.019 | 125.6 |
| | Control | vs. | HVL | 0.277 | 23.75, n=14 | 33.51, n=7 | HVL | 0 | 0.001 | 23.75 | 43.68 | 473.4 | 61.86 | 126.2 | 33.73 | -11.01 | 134.7 |
| | | | | | | | Control | 28.03 | 29.56 | 33.51 | 82.52 | 208.3 | 64.32 | 66.28 | 25.05 | 3.019 | 125.6 |
| IL12p70 | Control | vs. | LVL | 0.8365 | 0.1232, n=9 | 0.1401, n=7 | LVL | 0.001 | 0.03834 | 0.1232 | 0.2215 | 0.2953 | 0.1381 | 0.1009 | 0.03364 | 0.06051 | 0.2156 |
| | | | | | | | Control | 0.001 | 0.001 | 0.1401 | 0.266 | 0.2848 | 0.1282 | 0.1182 | 0.04468 | 0.01885 | 0.2375 |
| | Control | vs. | HVL | 0.8428 | 0.07919, n=14 | 0.1401, n=7 | HVL | 0.001 | 0.001 | 0.0791 9 | 0.2068 | 0.4075 | 0.1196 | 0.1241 | 0.03316 | 0.04798 | 0.1912 |
| | | | | | | | Control | 0.001 | 0.001 | 0.1401 | 0.266 | 0.2848 | 0.1282 | 0.1182 | 0.04468 | 0.01885 | 0.2375 |
| IL-15 | Control | vs. | LVL | 0.1909 | 0.5003, n=9 | 0.001000, n=7 | LVL | 0.001 | 0.001 | 0.5003 | 39.37 | 56.55 | 15.34 | 23.06 | 7.686 | -2.388 | 33.06 |
| | | | | | | | Control | 0 | 0.001 | 0.001 | 8.391 | 33.51 | 5.987 | 12.53 | 4.738 | -5.606 | 17.58 |
| | Control | vs. | HVL | 0.133 | 6.372, n=14 | 0.001000, n=7 | HVL | 0.001 | 0.001 | 6.372 | 24.73 | 70.98 | 14.67 | 20.83 | 5.566 | 2.644 | 26.69 |
| | | | | | | | Control | 0 | 0.001 | 0.001 | 8.391 | 33.51 | 5.987 | 12.53 | 4.738 | -5.606 | 17.58 |
| IL-17a | Control | vs. | LVL | 0.0705 | 12.37, n=9 | 19.94, n=7 | LVL | 0.001 | 0.001 | 12.37 | 20.69 | 30.7 | 12.13 | 11.18 | 3.726 | 3.537 | 20.72 |
| | | | | | | | Control | 11.38 | 18.64 | 19.94 | 28.03 | 40.87 | 22.61 | 9.391 | 3.549 | 13.92 | 31.29 |
| | Control | vs. | HVL | 0.488 | 17.78, n=14 | 19.94, n=7 | HVL | 0 | 15.37 | 17.78 | 23.69 | 36.08 | 19.33 | 8.634 | 2.308 | 14.34 | 24.31 |
| | | | | | | | Control | 11.38 | 18.64 | 19.94 | 28.03 | 40.87 | 22.61 | 9.391 | 3.549 | 13.92 | 31.29 |
| IL-18 | Control | vs. | LVL | 0.0559 | 4.403, n=9 | 0.001000, n=7 | LVL | 0.001 | 0.001 | 4.403 | 17.9 | 84.66 | 15.06 | 27.31 | 9.103 | -5.931 | 36.05 |
| | | | | | | | Control | 0 | 0.001 | 0.001 | 0.001 | 18.02 | 2.575 | 6.81 | 2.574 | -3.723 | 8.873 |
| | Control | vs. | HVL | 0.0394 | 14.37, n=14 | 0.001000, n=7 | HVL | 0.001 | 0.001 | 14.37 | 36.53 | 66.35 | 19.23 | 21.67 | 5.793 | 6.712 | 31.74 |
| | | | | | | | Control | 0 | 0.001 | 0.001 | 0.001 | 18.02 | 2.575 | 6.81 | 2.574 | -3.723 | 8.873 |
| IL-2 | Control | vs. | LVL | 0.022 | 0.8171, n=9 | 4.752, n=7 | LVL | 0 | 0.001 | 0.8171 | 3.147 | 29.39 | 4.257 | 9.539 | 3.18 | -3.076 | 11.59 |
| | | | | | | | Control | 3.134 | 3.487 | 4.752 | 7.88 | 14.17 | 6.418 | 3.86 | 1.459 | 2.849 | 9.988 |
| | Control | vs. | HVL | 0.5846 | 4.750, n=14 | 4.752, n=7 | HVL | 0 | 1.49 | 4.75 | 6.762 | 39.48 | 7.834 | 11.22 | 2.999 | 1.356 | 14.31 |
| | | | | | | | Control | 3.134 | 3.487 | 4.752 | 7.88 | 14.17 | 6.418 | 3.86 | 1.459 | 2.849 | 9.988 |
| IL-21 | Control | vs. | LVL | 0.773 | 0.001000, n=9 | 0.001000, n=7 | LVL | 0 | 0 | 0.001 | 690.6 | 2716 | 455.2 | 955.4 | 318.5 | -279.2 | 1190 |
| | | | | | | | Control | 0 | 0 | 0.001 | 355.7 | 1515 | 267.3 | 566 | 213.9 | -256.2 | 790.8 |
| | Control | vs. | HVL | 0.2225 | 5.760, n=14 | 0.001000, n=7 | HVL | 0 | 0.001 | 5.76 | 522.8 | 2437 | 353.1 | 684.4 | 182.9 | -42.07 | 748.3 |
| | | | | | | | Control | 0 | 0 | 0.001 | 355.7 | 1515 | 267.3 | 566 | 213.9 | -256.2 | 790.8 |
| IL-22 | Control | vs. | LVL | 0.361 | 0.3319, n=9 | 0.001000, n=7 | LVL | 0.001 | 0.001 | 0.3319 | 22.94 | 63.93 | 12.42 | 23.73 | 7.909 | -5.819 | 30.66 |
| | | | | | | | Control | 0 | 0.001 | 0.001 | 3.862 | 51.79 | 7.951 | 19.38 | 7.327 | -9.977 | 25.88 |
| | Control | vs. | HVL | 0.2199 | 1.644, n=14 | 0.001000, n=7 | HVL | 0.001 | 0.001 | 1.644 | 15.73 | 72.3 | 12.12 | 21.24 | 5.677 | -0.1399 | 24.39 |
| | | | | | | | Control | 0 | 0.001 | 0.001 | 3.862 | 51.79 | 7.951 | 19.38 | 7.327 | -9.977 | 25.88 |
| IL-23 | Control | vs. | LVL | 0.2105 | 95.95, n=9 | 110.8, n=7 | LVL | 8.786 | 53.83 | 95.95 | 153.5 | 249.4 | 107.8 | 72.76 | 24.25 | 51.84 | 163.7 |
| | | | | | | | Control | 82.95 | 89.07 | 110.8 | 306.4 | 388.9 | 185.5 | 122.1 | 46.16 | 72.52 | 298.4 |
| | Control | vs. | HVL | 0.0056 | 28.72, n=14 | 110.8, n=7 | HVL | 3.601 | 9.452 | 28.72 | 79.15 | 240.2 | 58.2 | 71.25 | 19.04 | 17.06 | 99.34 |
| | | | | | | | Control | 82.95 | 89.07 | 110.8 | 306.4 | 388.9 | 185.5 | 122.1 | 46.16 | 72.52 | 298.4 |
| IL-27 | Control | vs. | LVL | 0.6754 | 2.468, n=9 | 0.001000, n=7 | LVL | 0 | 0.001 | 2.468 | 1212 | 3939 | 708.5 | 1431 | 477 | -391.5 | 1809 |
| | | | | | | | Control | 0 | 0 | 0.001 | 344.4 | 517.9 | 155 | 211.5 | 79.93 | -40.54 | 350.6 |
| | Control | vs. | HVL | 0.3129 | 70.75, n=14 | 0.001000, n=7 | HVL | 0 | 0.001 | 70.75 | 725.6 | 1472 | 372.1 | 515 | 137.6 | 74.72 | 669.4 |
| | | | | | | | Control | 0 | 0 | 0.001 | 344.4 | 517.9 | 155 | 211.5 | 79.93 | -40.54 | 350.6 |
| IL-29 | Control | vs. | LVL | 0.6108 | 0.000, n=9 | 0.000, n=7 | LVL | 0 | 0 | 0 | 4.003 | 26.06 | 3.785 | 8.626 | 2.875 | -2.846 | 10.42 |
| | | | | | | | Control | 0 | 0 | 0 | 0.001 | 0.001 | 0.000428 6 | 0.000534 5 | 0.000202 | -6.578E-05 | 0.000922 9 |
| | Control | vs. | HVL | 0.085 | 0.001000, n=14 | 0.000, n=7 | HVL | 0 | 0 | 0.001 | 33.22 | 55.13 | 15.04 | 21.43 | 5.729 | 2.66 | 27.41 |
| | | | | | | | Control | 0 | 0 | 0 | 0.001 | 0.001 | 0.000428 6 | 0.000534 5 | 0.000202 | -6.578E-05 | 0.000922 9 |
| IL-33 | Control | vs. | LVL | 0.0031 | 1.222, n=9 | 0.001000, n=7 | LVL | 0.001 | 0.008391 | 1.222 | 10.19 | 33.03 | 6.333 | 11.6 | 3.868 | -2.587 | 15.25 |
| | | | | | | | Control | 0 | 0 | 0.001 | 0.001 | 0.001 | 0.000714 3 | 0.000488 | 0.000184 4 | 0.00026 3 | 0.001166 |
| | Control | vs. | HVL | 0.0066 | 0.7363, n=14 | 0.001000, n=7 | HVL | 0.001 | 0.001 | 0.7363 | 6.133 | 12.44 | 3.324 | 4.397 | 1.175 | 0.7849 | 5.863 |
| | | | | | | | Control | 0 | 0 | 0.001 | 0.001 | 0.001 | 0.000714 3 | 0.000488 | 0.000184 4 | 0.00026 3 | 0.001166 |
| IL-4 | Control | vs. | LVL | 0.1724 | 5.008, n=9 | 10.89, n=7 | LVL | 0.001 | 0.001 | 5.008 | 30.91 | 33.42 | 11.91 | 15.09 | 5.029 | 0.3155 | 23.51 |
| | | | | | | | Control | 7.875 | 7.924 | 10.89 | 19.08 | 33.71 | 14.58 | 9.414 | 3.558 | 5.871 | 23.28 |
| | Control | vs. | HVL | 0.2245 | 23.00, n=14 | 10.89, n=7 | HVL | 0 | 12.61 | 23 | 31.22 | 42.38 | 20.81 | 12.52 | 3.346 | 13.58 | 28.04 |
| | | | | | | | Control | 7.875 | 7.924 | 10.89 | 19.08 | 33.71 | 14.58 | 9.414 | 3.558 | 5.871 | 23.28 |
| IL-7 | Control | vs. | LVL | 0.4391 | 0.07662, n=9 | 0.001000, n=7 | LVL | 0 | 0.0005 | 0.0766 2 | 27.72 | 112 | 18.65 | 39.09 | 13.03 | -11.4 | 48.7 |
| | | | | | | | Control | 0 | 0 | 0.001 | 6.505 | 89.8 | 13.76 | 33.62 | 12.71 | -17.33 | 44.85 |
| | Control | vs. | HVL | 0.2663 | 0.001000, n=14 | 0.001000, n=7 | HVL | 0 | 0.001 | 0.001 | 39.51 | 146 | 26.17 | 45.13 | 12.06 | 0.1075 | 52.23 |
| | | | | | | | Control | 0 | 0 | 0.001 | 6.505 | 89.8 | 13.76 | 33.62 | 12.71 | -17.33 | 44.85 |
| IL-8 | Control | vs. | LVL | 0.16 | 2779, n=9 | 7244, n=7 | LVL | 0.001 | 0.001 | 2779 | 10254 | 14334 | 4910 | 5681 | 1894 | 543.9 | 9277 |
| | | | | | | | Control | 6742 | 6794 | 7244 | 10461 | 13721 | 8456 | 2665 | 1007 | 5992 | 10920 |
| | Control | vs. | HVL | 0.799 | 7691, n=14 | 7244, n=7 | HVL | 0 | 4520 | 7691 | 11800 | 12913 | 7424 | 4617 | 1234 | 4759 | 10090 |
| | | | | | | | Control | 6742 | 6794 | 7244 | 10461 | 13721 | 8456 | 2665 | 1007 | 5992 | 10920 |
| IL-9 | Control | vs. | LVL | 0.3322 | 1.806, n=9 | 2.904, n=7 | LVL | 0.001 | 0.001 | 1.806 | 3.674 | 4.974 | 1.89 | 1.989 | 0.663 | 0.3607 | 3.418 |
| | | | | | | | Control | 2.359 | 2.544 | 2.904 | 4.091 | 4.103 | 3.126 | 0.7016 | 0.2652 | 2.477 | 3.775 |
| | Control | vs. | HVL | 0.6359 | 3.771, n=14 | 2.904, n=7 | HVL | 0 | 1.686 | 3.771 | 5.038 | 24.45 | 4.68 | 6.05 | 1.617 | 1.186 | 8.173 |
| | | | | | | | Control | 2.359 | 2.544 | 2.904 | 4.091 | 4.103 | 3.126 | 0.7016 | 0.2652 | 2.477 | 3.775 |
| IP-10 | Control | vs. | LVL | >0.9999 | 0.001000, n=9 | 0.001000, n=7 | LVL | 0.001 | 0.001 | 0.001 | 0.001 | 0.001 | 0.001 | 0 | 0 | 0.001 | 0.001 |
| | | | | | | | Control | 0.001 | 0.001 | 0.001 | 0.001 | 0.001 | 0.001 | 0 | 0 | 0.001 | 0.001 |
| | Control | vs. | HVL | >0.9999 | 0.001000, n=14 | 0.001000, n=7 | HVL | 0.001 | 0.001 | 0.001 | 0.001 | 365.4 | 26.1 | 97.65 | 26.1 | -30.28 | 82.48 |
| | | | | | | | Control | 0.001 | 0.001 | 0.001 | 0.001 | 0.001 | 0.001 | 0 | 0 | 0.001 | 0.001 |
| MIP-1α | Control | vs. | LVL | 0.9115 | 0.001000, n=9 | 0.001000, n=7 | LVL | 0 | 0.001 | 0.001 | 0.001 | 15225 | 1692 | 5075 | 1692 | -2209 | 5593 |
| | | | | | | | Control | 0 | 0.001 | 0.001 | 0.001 | 47052 | 6722 | 17784 | 6722 | -9726 | 23169 |
| | Control | | HVL | 0.6398 | | | HVL | 0.001 | 0.001 | 0.001 | 0.001 | 25067 | 2233 | 6776 | 1811 | -1680 | 6145 |

| | | | | | | | | | | | | | | | | | |
|---|---|---|---|---|---|---|---|---|---|---|---|---|---|---|---|---|---|
| | | vs. | | | 0.001000, n=14 | 0.001000, n=7 | Control | 0 | 0.001 | 0.001 | 0.001 | 47052 | 6722 | 17784 | 6722 | -9726 | 23169 |
| SDF-1α | Control | vs. | LVL | >0.9999 | 438.9, n=9 | 351.2, n=7 | LVL | 71.89 | 212.7 | 438.9 | 697.1 | 926 | 453.2 | 283.3 | 94.43 | 235.4 | 670.9 |
| | | | | | | | Control | 221.4 | 265.1 | 351.2 | 572.1 | 767.5 | 425.1 | 196 | 74.1 | 243.8 | 606.4 |
| | Control | vs. | HVL | 0.0442 | 207.1, n=14 | 351.2, n=7 | HVL | 0 | 0.001 | 207.1 | 352.5 | 790.1 | 231.2 | 217.7 | 58.17 | 105.5 | 356.8 |
| | | | | | | | Control | 221.4 | 265.1 | 351.2 | 572.1 | 767.5 | 425.1 | 196 | 74.1 | 243.8 | 606.4 |
| TGF-β1 | Control | vs. | LVL | 0.349 | 0.001000, n=9 | 296.2, n=7 | LVL | 0.001 | 0.001 | 0.001 | 352.3 | 749.1 | 166.3 | 299.6 | 99.87 | -64.03 | 396.6 |
| | | | | | | | Control | 0.001 | 33.53 | 296.2 | 603.2 | 606.5 | 293.5 | 273.3 | 103.3 | 40.81 | 546.3 |
| | Control | vs. | HVL | 0.1117 | 0.001000, n=14 | 296.2, n=7 | HVL | 0.001 | 0.001 | 0.001 | 237.1 | 1376 | 189.9 | 393.8 | 105.2 | -37.5 | 417.2 |
| | | | | | | | Control | 0.001 | 33.53 | 296.2 | 603.2 | 606.5 | 293.5 | 273.3 | 103.3 | 40.81 | 546.3 |
| TGF-β2 | Control | vs. | LVL | >0.9999 | 0.001000, n=9 | 6.974, n=7 | LVL | 0.001 | 0.001 | 0.001 | 54.94 | 103.7 | 25.24 | 40.23 | 13.41 | -5.686 | 56.16 |
| | | | | | | | Control | 0.001 | 0.001 | 6.974 | 17.66 | 49.86 | 13.12 | 17.99 | 6.801 | -3.517 | 29.76 |
| | Control | vs. | HVL | 0.7233 | 0.001000, n=14 | 6.974, n=7 | HVL | 0.001 | 0.001 | 0.001 | 28.65 | 137.7 | 19.41 | 39.65 | 10.6 | -3.486 | 42.3 |
| | | | | | | | Control | 0.001 | 0.001 | 6.974 | 17.66 | 49.86 | 13.12 | 17.99 | 6.801 | -3.517 | 29.76 |
| TGF-β3 | Control | vs. | LVL | 0.7247 | 0.06388, n=9 | 0.001000, n=7 | LVL | 0.001 | 0.001 | 0.06388 | 0.4308 | 0.6938 | 0.1934 | 0.2802 | 0.09342 | -0.02207 | 0.4088 |
| | | | | | | | Control | 0.001 | 0.001 | 0.001 | 0.2621 | 0.3051 | 0.1059 | 0.1368 | 0.0517 | -0.02057 | 0.2324 |
| | Control | vs. | HVL | 0.3015 | 0.1440, n=14 | 0.001000, n=7 | HVL | 0.001 | 0.001 | 0.144 | 0.3668 | 0.94 | 0.2232 | 0.2617 | 0.06995 | 0.07206 | 0.3743 |
| | | | | | | | Control | 0.001 | 0.001 | 0.001 | 0.2621 | 0.3051 | 0.1059 | 0.1368 | 0.0517 | -0.02057 | 0.2324 |
