## Supplemental Table 11 for "Chronic opioid-associated immune dysregulation among people living with HIV"

**Supplementary Table 11.** CD3+CD4+ T cell Flow cytometry results

| Marker | Comparison (A vs B) | Group A | | | Group B | | | Viral Load adjusted Group A – Group B | | | |
|---|---|---|---|---|---|---|---|---|---|---|---|
| | | n | Mean | SD | N | Mean | SD | Estimate | SE | p | adj p |
| CD3+ \| Geometric Mean (BUV496-A :: CD4) | Control vs. CTN-067 M0 | 8 | 9324.63 | 5762.58 | 11 | 7192.18 | 5590.65 | 14.53 | 3342.39 | 0.9966 | 1.0000 |
| | Control vs. CTN-067 M3 | 8 | 9324.63 | 5762.58 | 11 | 8589.82 | 6697.21 | -670.98 | 3271.07 | 0.8397 | 0.9968 |
| | Control vs. CTN-067 M6 | 8 | 9324.63 | 5762.58 | 11 | 8688.09 | 7121.74 | -733.85 | 3268.51 | 0.8247 | 0.9959 |
| | CTN-067 M0 vs. CTN-067 M3 | 11 | 7192.18 | 5590.65 | 11 | 8589.82 | 6697.21 | -685.51 | 719.94 | 0.3530 | 0.7775 |
| | CTN-067 M0 vs. CTN-067 M6 | 11 | 7192.18 | 5590.65 | 11 | 8688.09 | 7121.74 | -748.38 | 729.02 | 0.3175 | 0.7363 |
| | CTN-067 M3 vs. CTN-067 M6 | 11 | 8589.82 | 6697.21 | 11 | 8688.09 | 7121.74 | -62.87 | 624.10 | 0.9208 | 0.9996 |
| CD3+CD4+ \| Freq. of Parent | Control vs. CTN-067 M0 | 8 | 56.91 | 12.51 | 11 | 49.52 | 16.15 | -0.11 | 7.62 | 0.9882 | 1.0000 |
| | Control vs. CTN-067 M3 | 8 | 56.91 | 12.51 | 11 | 53.24 | 15.49 | 0.26 | 7.40 | 0.9719 | 1.0000 |
| | Control vs. CTN-067 M6 | 8 | 56.91 | 12.51 | 11 | 53.83 | 15.53 | -0.12 | 7.39 | 0.9869 | 1.0000 |
| | CTN-067 M0 vs. CTN-067 M3 | 11 | 49.52 | 16.15 | 11 | 53.24 | 15.49 | 0.38 | 1.90 | 0.8444 | 0.9971 |
| | CTN-067 M0 vs. CTN-067 M6 | 11 | 49.52 | 16.15 | 11 | 53.83 | 15.53 | -0.01 | 1.93 | 0.9965 | 1.0000 |
| | CTN-067 M3 vs. CTN-067 M6 | 11 | 53.24 | 15.49 | 11 | 53.83 | 15.53 | -0.39 | 1.65 | 0.8171 | 0.9953 |
| CD3+CD4+ \| Geometric Mean (APC-Fire 750-A :: CD36) | Control vs. CTN-067 M0 | 8 | 631.25 | 204.42 | 11 | 475.82 | 195.67 | 259.32 | 129.33 | 0.0594 | 0.2209 |
| | Control vs. CTN-067 M3 | 8 | 631.25 | 204.42 | 11 | 495.82 | 243.13 | 179.63 | 118.02 | 0.1445 | 0.4446 |
| | Control vs. CTN-067 M6 | 8 | 631.25 | 204.42 | 11 | 557.82 | 218.54 | 114.66 | 117.60 | 0.3418 | 0.7651 |
| | CTN-067 M0 vs. CTN-067 M3 | 11 | 475.82 | 195.67 | 11 | 495.82 | 243.13 | -79.70 | 56.91 | 0.1775 | 0.5142 |

| | | | | | | | | | | | |
|---|---|---|---|---|---|---|---|---|---|---|---|
| | CTN-067 M0 vs. CTN-067 M6 | 11 | 475.82 | 195.67 | 11 | 557.82 | 218.54 | -144.67 | 57.59 | 0.0212 | 0.0900 |
| | CTN-067 M3 vs. CTN-067 M6 | 11 | 495.82 | 243.13 | 11 | 557.82 | 218.54 | -64.97 | 49.75 | 0.2071 | 0.5704 |
| CD3+CD4+ \| Geometric Mean (Alexa Fluor 647-A :: GLUT1) | Control vs. CTN-067 M0 | 8 | 334.49 | 348.13 | 11 | 341.40 | 391.41 | 228.47 | 200.15 | 0.2679 | 0.6693 |
| | Control vs. CTN-067 M3 | 8 | 334.49 | 348.13 | 11 | 369.39 | 290.17 | 113.18 | 186.30 | 0.5507 | 0.9284 |
| | Control vs. CTN-067 M6 | 8 | 334.49 | 348.13 | 11 | 375.91 | 323.30 | 102.32 | 185.79 | 0.5882 | 0.9452 |
| | CTN-067 M0 vs. CTN-067 M3 | 11 | 341.40 | 391.41 | 11 | 369.39 | 290.17 | -115.29 | 77.99 | 0.1558 | 0.4693 |
| | CTN-067 M0 vs. CTN-067 M6 | 11 | 341.40 | 391.41 | 11 | 375.91 | 323.30 | -126.14 | 78.95 | 0.1266 | 0.4033 |
| | CTN-067 M3 vs. CTN-067 M6 | 11 | 369.39 | 290.17 | 11 | 375.91 | 323.30 | -10.86 | 67.98 | 0.8748 | 0.9985 |
| CD3+CD4+/GLUT1+ \| Freq. of Parent | Control vs. CTN-067 M0 | 8 | 0.43 | 0.32 | 11 | 0.51 | 0.37 | 0.08 | 0.21 | 0.7048 | 0.9801 |
| | Control vs. CTN-067 M3 | 8 | 0.43 | 0.32 | 11 | 0.52 | 0.31 | -0.01 | 0.19 | 0.9479 | 0.9999 |
| | Control vs. CTN-067 M6 | 8 | 0.43 | 0.32 | 11 | 0.57 | 0.39 | -0.06 | 0.19 | 0.7531 | 0.9884 |
| | CTN-067 M0 vs. CTN-067 M3 | 11 | 0.51 | 0.37 | 11 | 0.52 | 0.31 | -0.09 | 0.09 | 0.3297 | 0.7511 |
| | CTN-067 M0 vs. CTN-067 M6 | 11 | 0.51 | 0.37 | 11 | 0.57 | 0.39 | -0.14 | 0.09 | 0.1506 | 0.4581 |
| | CTN-067 M3 vs. CTN-067 M6 | 11 | 0.52 | 0.31 | 11 | 0.57 | 0.39 | -0.05 | 0.08 | 0.5624 | 0.9340 |
| CD3+CD4+ \| Geometric Mean (BUV395-A :: CD28) | Control vs. CTN-067 M0 | 8 | 4206.38 | 1120.50 | 11 | 4947.36 | 1452.95 | -488.13 | 700.42 | 0.4943 | 0.8970 |
| | Control vs. CTN-067 M3 | 8 | 4206.38 | 1120.50 | 11 | 4216.73 | 1229.40 | 170.32 | 665.66 | 0.8008 | 0.9939 |
| | Control vs. CTN-067 M6 | 8 | 4206.38 | 1120.50 | 11 | 3979.27 | 1093.60 | 404.19 | 664.39 | 0.5502 | 0.9282 |
| | CTN-067 M0 vs. | 11 | 4947.36 | 1452.95 | 11 | 4216.73 | 1229.40 | 658.45 | 230.48 | 0.0101 | 0.0456 |

| | | | | | | | | | | |
|---|---|---|---|---|---|---|---|---|---|---|
| | CTN-067 M3 | | | | | | | | | |
| | CTN-067 M0 vs. CTN-067 M6 | 11 | 4947.36 | 1452.95 | 11 | 3979.27 | 1093.60 | 892.32 | 233.33 | 0.0011 | 0.0058 |
| | CTN-067 M3 vs. CTN-067 M6 | 11 | 4216.73 | 1229.40 | 11 | 3979.27 | 1093.60 | 233.87 | 200.38 | 0.2576 | 0.6540 |
| CD3+CD4+/CD28 \| Freq. of Parent | Control vs. CTN-067 M0 | 8 | 92.75 | 4.64 | 11 | 95.61 | 1.96 | -2.63 | 1.86 | 0.1746 | 0.5084 |
| | Control vs. CTN-067 M3 | 8 | 92.75 | 4.64 | 11 | 94.84 | 2.49 | -1.72 | 1.83 | 0.3602 | 0.7853 |
| | Control vs. CTN-067 M6 | 8 | 92.75 | 4.64 | 11 | 95.61 | 2.67 | -2.48 | 1.83 | 0.1907 | 0.5399 |
| | CTN-067 M0 vs. CTN-067 M3 | 11 | 95.61 | 1.96 | 11 | 94.84 | 2.49 | 0.91 | 0.37 | 0.0231 | 0.0971 |
| | CTN-067 M0 vs. CTN-067 M6 | 11 | 95.61 | 1.96 | 11 | 95.61 | 2.67 | 0.15 | 0.37 | 0.6990 | 0.9789 |
| | CTN-067 M3 vs. CTN-067 M6 | 11 | 94.84 | 2.49 | 11 | 95.61 | 2.67 | -0.77 | 0.32 | 0.0272 | 0.1123 |
| CD3+CD4+ \| Geometric Mean (BUV661-A :: CD27) | Control vs. CTN-067 M0 | 8 | 18053.38 | 6264.31 | 11 | 29820.82 | 9545.62 | -10451.00 | 7746056.00 | 0.9989 | 1.0000 |
| | Control vs. CTN-067 M3 | 8 | 18053.38 | 6264.31 | 11 | 30595.91 | 10542.16 | -11529.00 | 7746056.00 | 0.9988 | 1.0000 |
| | Control vs. CTN-067 M6 | 8 | 18053.38 | 6264.31 | 11 | 29691.18 | 9731.13 | -10639.00 | 7746056.00 | 0.9989 | 1.0000 |
| | CTN-067 M0 vs. CTN-067 M3 | 11 | 29820.82 | 9545.62 | 11 | 30595.91 | 10542.16 | -1077.22 | 688.73 | 0.1343 | 0.4214 |
| | CTN-067 M0 vs. CTN-067 M6 | 11 | 29820.82 | 9545.62 | 11 | 29691.18 | 9731.13 | -187.52 | 697.51 | 0.7910 | 0.9930 |
| | CTN-067 M3 vs. CTN-067 M6 | 11 | 30595.91 | 10542.16 | 11 | 29691.18 | 9731.13 | 889.71 | 596.00 | 0.1519 | 0.4610 |
| CD3+CD4+/CD27+ \| Freq. of Parent | Control vs. CTN-067 M0 | 8 | 85.58 | 8.58 | 11 | 93.04 | 4.20 | -7.09 | 3.50 | 0.0570 | 0.2133 |
| | Control vs. CTN-067 M3 | 8 | 85.58 | 8.58 | 11 | 93.55 | 4.73 | -7.39 | 3.46 | 0.0458 | 0.1775 |
| | Control vs. CTN-067 M6 | 8 | 85.58 | 8.58 | 11 | 93.41 | 4.92 | -7.25 | 3.46 | 0.0498 | 0.1904 |

| | | | | | | | | | | | |
|---|---|---|---|---|---|---|---|---|---|---|---|
| | CTN-067 M0 vs. CTN-067 M3 | 11 | 93.04 | 4.20 | 11 | 93.55 | 4.73 | -0.30 | 0.55 | 0.5909 | 0.9463 |
| | CTN-067 M0 vs. CTN-067 M6 | 11 | 93.04 | 4.20 | 11 | 93.41 | 4.92 | -0.15 | 0.56 | 0.7854 | 0.9924 |
| | CTN-067 M3 vs. CTN-067 M6 | 11 | 93.55 | 4.73 | 11 | 93.41 | 4.92 | 0.15 | 0.48 | 0.7613 | 0.9895 |
| CD3+CD4+ \| Geometric Mean (BV510-A :: CCR7) | Control vs. CTN-067 M0 | 8 | 7213.25 | 2258.53 | 11 | 9110.09 | 3756.22 | -11.34 | 1681.40 | 0.9947 | 1.0000 |
| | Control vs. CTN-067 M3 | 8 | 7213.25 | 2258.53 | 11 | 9755.73 | 3793.48 | -1019.43 | 1566.10 | 0.5229 | 0.9139 |
| | Control vs. CTN-067 M6 | 8 | 7213.25 | 2258.53 | 11 | 9337.00 | 2864.78 | -618.73 | 1561.86 | 0.6964 | 0.9783 |
| | CTN-067 M0 vs. CTN-067 M3 | 11 | 9110.09 | 3756.22 | 11 | 9755.73 | 3793.48 | -1008.09 | 652.12 | 0.1386 | 0.4314 |
| | CTN-067 M0 vs. CTN-067 M6 | 11 | 9110.09 | 3756.22 | 11 | 9337.00 | 2864.78 | -607.39 | 660.08 | 0.3690 | 0.7945 |
| | CTN-067 M3 vs. CTN-067 M6 | 11 | 9755.73 | 3793.48 | 11 | 9337.00 | 2864.78 | 400.71 | 568.34 | 0.4893 | 0.8938 |
| CD3+CD4+/CCR7 \| Freq. of Parent | Control vs. CTN-067 M0 | 8 | 70.58 | 10.79 | 11 | 78.59 | 8.60 | -6.42 | 5.17 | 0.2289 | 0.6082 |
| | Control vs. CTN-067 M3 | 8 | 70.58 | 10.79 | 11 | 78.73 | 9.57 | -6.22 | 5.05 | 0.2333 | 0.6155 |
| | Control vs. CTN-067 M6 | 8 | 70.58 | 10.79 | 11 | 79.13 | 8.67 | -6.60 | 5.05 | 0.2064 | 0.5692 |
| | CTN-067 M0 vs. CTN-067 M3 | 11 | 78.59 | 8.60 | 11 | 78.73 | 9.57 | 0.21 | 1.14 | 0.8591 | 0.9979 |
| | CTN-067 M0 vs. CTN-067 M6 | 11 | 78.59 | 8.60 | 11 | 79.13 | 8.67 | -0.18 | 1.16 | 0.8797 | 0.9987 |
| | CTN-067 M3 vs. CTN-067 M6 | 11 | 78.73 | 9.57 | 11 | 79.13 | 8.67 | -0.38 | 0.99 | 0.7032 | 0.9797 |
| CD3+CD4+ \| Geometric Mean (BV605-A :: CD45RA) | Control vs. CTN-067 M0 | 8 | 6334.13 | 2214.77 | 11 | 8262.45 | 4906.85 | 1545.39 | 3033.45 | 0.6163 | 0.9558 |
| | Control vs. CTN-067 M3 | 8 | 6334.13 | 2214.77 | 11 | 10756.09 | 7865.14 | -1956.87 | 2795.50 | 0.4924 | 0.8958 |

| | | | | | | | | | | | |
|---|---|---|---|---|---|---|---|---|---|---|---|
| | Control vs. CTN-067 M6 | 8 | 6334.13 | 2214.77 | 11 | 9828.00 | 6368.76 | -1078.93 | 2786.70 | 0.7029 | 0.9797 |
| | CTN-067 M0 vs. CTN-067 M3 | 11 | 8262.45 | 4906.85 | 11 | 10756.09 | 7865.14 | -3502.26 | 1261.03 | 0.0120 | 0.0536 |
| | CTN-067 M0 vs. CTN-067 M6 | 11 | 8262.45 | 4906.85 | 11 | 9828.00 | 6368.76 | -2624.31 | 1276.27 | 0.0538 | 0.2032 |
| | CTN-067 M3 vs. CTN-067 M6 | 11 | 10756.09 | 7865.14 | 11 | 9828.00 | 6368.76 | 877.95 | 1100.65 | 0.4349 | 0.8546 |
| CD3+CD4+/CD45RA \| Freq. of Parent | Control vs. CTN-067 M0 | 8 | 31.36 | 8.34 | 11 | 39.30 | 14.46 | -0.19 | 7.07 | 0.9786 | 1.0000 |
| | Control vs. CTN-067 M3 | 8 | 31.36 | 8.34 | 11 | 42.57 | 16.19 | -5.02 | 6.62 | 0.4580 | 0.8723 |
| | Control vs. CTN-067 M6 | 8 | 31.36 | 8.34 | 11 | 40.83 | 14.86 | -3.35 | 6.61 | 0.6181 | 0.9564 |
| | CTN-067 M0 vs. CTN-067 M3 | 11 | 39.30 | 14.46 | 11 | 42.57 | 16.19 | -4.83 | 2.62 | 0.0815 | 0.2865 |
| | CTN-067 M0 vs. CTN-067 M6 | 11 | 39.30 | 14.46 | 11 | 40.83 | 14.86 | -3.16 | 2.66 | 0.2491 | 0.6409 |
| | CTN-067 M3 vs. CTN-067 M6 | 11 | 42.57 | 16.19 | 11 | 40.83 | 14.86 | 1.67 | 2.28 | 0.4742 | 0.8838 |
| CD3+CD4+ \| Geometric Mean (BV711-A :: CD86) | Control vs. CTN-067 M0 | 8 | 120.51 | 59.43 | 11 | -13.24 | 409.41 | 188.05 | 150.16 | 0.2256 | 0.6027 |
| | Control vs. CTN-067 M3 | 8 | 120.51 | 59.43 | 11 | 57.31 | 93.88 | 96.36 | 125.47 | 0.4519 | 0.8678 |
| | Control vs. CTN-067 M6 | 8 | 120.51 | 59.43 | 11 | 76.04 | 94.23 | 76.58 | 124.51 | 0.5458 | 0.9260 |
| | CTN-067 M0 vs. CTN-067 M3 | 11 | -13.24 | 409.41 | 11 | 57.31 | 93.88 | -91.69 | 106.94 | 0.4019 | 0.8263 |
| | CTN-067 M0 vs. CTN-067 M6 | 11 | -13.24 | 409.41 | 11 | 76.04 | 94.23 | -111.47 | 107.82 | 0.3142 | 0.7321 |
| | CTN-067 M3 vs. CTN-067 M6 | 11 | 57.31 | 93.88 | 11 | 76.04 | 94.23 | -19.78 | 97.86 | 0.8420 | 0.9970 |
| CD3+CD4+/CD86+ \| Freq. of Parent | Control vs. CTN-067 M0 | 8 | 0.98 | 0.66 | 11 | 1.29 | 0.74 | -0.25 | 0.38 | 0.5144 | 0.9091 |

| | | | | | | | | | | | |
|---|---|---|---|---|---|---|---|---|---|---|---|
| | Control vs. CTN-067 M3 | 8 | 0.98 | 0.66 | 11 | 1.01 | 0.58 | -0.05 | 0.34 | 0.8931 | 0.9991 |
| | Control vs. CTN-067 M6 | 8 | 0.98 | 0.66 | 11 | 0.90 | 0.49 | 0.06 | 0.33 | 0.8657 | 0.9981 |
| | CTN-067 M0 vs. CTN-067 M3 | 11 | 1.29 | 0.74 | 11 | 1.01 | 0.58 | 0.21 | 0.19 | 0.2960 | 0.7087 |
| | CTN-067 M0 vs. CTN-067 M6 | 11 | 1.29 | 0.74 | 11 | 0.90 | 0.49 | 0.31 | 0.19 | 0.1274 | 0.4053 |
| | CTN-067 M3 vs. CTN-067 M6 | 11 | 1.01 | 0.58 | 11 | 0.90 | 0.49 | 0.10 | 0.17 | 0.5478 | 0.9270 |
| CD3+CD4+ \| Geometric Mean (PE-A :: CD38) | Control vs. CTN-067 M0 | 8 | 3924.63 | 1671.66 | 11 | 6055.18 | 2813.75 | 280.07 | 1512.91 | 0.8551 | 0.9977 |
| | Control vs. CTN-067 M3 | 8 | 3924.63 | 1671.66 | 11 | 6448.64 | 3422.92 | -836.78 | 1345.84 | 0.5415 | 0.9238 |
| | Control vs. CTN-067 M6 | 8 | 3924.63 | 1671.66 | 11 | 5697.73 | 2896.79 | -121.83 | 1339.53 | 0.9285 | 0.9997 |
| | CTN-067 M0 vs. CTN-067 M3 | 11 | 6055.18 | 2813.75 | 11 | 6448.64 | 3422.92 | -1116.85 | 755.62 | 0.1558 | 0.4693 |
| | CTN-067 M0 vs. CTN-067 M6 | 11 | 6055.18 | 2813.75 | 11 | 5697.73 | 2896.79 | -401.91 | 764.39 | 0.6051 | 0.9518 |
| | CTN-067 M3 vs. CTN-067 M6 | 11 | 6448.64 | 3422.92 | 11 | 5697.73 | 2896.79 | 714.94 | 663.73 | 0.2949 | 0.7072 |
| CD3+CD4+/CD38 \| Freq. of Parent | Control vs. CTN-067 M0 | 8 | 42.24 | 7.80 | 11 | 56.95 | 12.91 | -9.20 | 6.36 | 0.1642 | 0.4871 |
| | Control vs. CTN-067 M3 | 8 | 42.24 | 7.80 | 11 | 58.42 | 14.32 | -11.55 | 6.04 | 0.0711 | 0.2564 |
| | Control vs. CTN-067 M6 | 8 | 42.24 | 7.80 | 11 | 55.93 | 13.86 | -9.10 | 6.03 | 0.1477 | 0.4517 |
| | CTN-067 M0 vs. CTN-067 M3 | 11 | 56.95 | 12.91 | 11 | 58.42 | 14.32 | -2.34 | 2.10 | 0.2788 | 0.6850 |
| | CTN-067 M0 vs. CTN-067 M6 | 11 | 56.95 | 12.91 | 11 | 55.93 | 13.86 | 0.10 | 2.13 | 0.9619 | 1.0000 |
| | CTN-067 M3 vs. CTN-067 M6 | 11 | 58.42 | 14.32 | 11 | 55.93 | 13.86 | 2.45 | 1.83 | 0.1965 | 0.5508 |

| | | | | | | | | | | |
|---|---|---|---|---|---|---|---|---|---|---|
| CD3+CD4+ \| Geometric Mean (R718-A :: CCR5) | Control vs. CTN-067 M0 | 8 | 882.63 | 370.43 | 1 1 | 550.36 | 216.84 | 425.93 | 180.36 | 0.02 90 | 0.11 92 |
| | Control vs. CTN-067 M3 | 8 | 882.63 | 370.43 | 1 1 | 546.64 | 235.45 | 383.39 | 156.31 | 0.02 40 | 0.10 07 |
| | Control vs. CTN-067 M6 | 8 | 882.63 | 370.43 | 1 1 | 582.09 | 250.32 | 345.64 | 155.39 | 0.03 84 | 0.15 25 |
| | CTN-067 M0 vs. CTN-067 M3 | 1 1 | 550.36 | 216.84 | 1 1 | 546.64 | 235.45 | -42.54 | 100.89 | 0.67 80 | 0.97 41 |
| | CTN-067 M0 vs. CTN-067 M6 | 1 1 | 550.36 | 216.84 | 1 1 | 582.09 | 250.32 | -80.30 | 102.00 | 0.44 09 | 0.85 93 |
| | CTN-067 M3 vs. CTN-067 M6 | 1 1 | 546.64 | 235.45 | 1 1 | 582.09 | 250.32 | -37.75 | 89.25 | 0.67 70 | 0.97 38 |
| CD3+CD4+/CCR5+ \| Freq. of Parent | Control vs. CTN-067 M0 | 8 | 19.75 | 7.75 | 1 1 | 13.80 | 5.27 | 6.14 | 3.82 | 0.12 40 | 0.39 70 |
| | Control vs. CTN-067 M3 | 8 | 19.75 | 7.75 | 1 1 | 12.31 | 6.23 | 7.27 | 3.61 | 0.05 82 | 0.21 72 |
| | Control vs. CTN-067 M6 | 8 | 19.75 | 7.75 | 1 1 | 12.67 | 5.99 | 6.90 | 3.60 | 0.07 06 | 0.25 50 |
| | CTN-067 M0 vs. CTN-067 M3 | 1 1 | 13.80 | 5.27 | 1 1 | 12.31 | 6.23 | 1.13 | 1.32 | 0.40 11 | 0.82 56 |
| | CTN-067 M0 vs. CTN-067 M6 | 1 1 | 13.80 | 5.27 | 1 1 | 12.67 | 5.99 | 0.75 | 1.33 | 0.57 82 | 0.94 10 |
| | CTN-067 M3 vs. CTN-067 M6 | 1 1 | 12.31 | 6.23 | 1 1 | 12.67 | 5.99 | -0.38 | 1.14 | 0.74 57 | 0.98 73 |
| CD3+CD4+ \| Geometric Mean (RB545-A :: CD57) | Control vs. CTN-067 M0 | 8 | 1149.6 3 | 325.90 | 1 1 | 535.45 | 521.24 | 711.34 | 242.79 | 0.00 86 | 0.03 93 |
| | Control vs. CTN-067 M3 | 8 | 1149.6 3 | 325.90 | 1 1 | 622.64 | 258.07 | 560.19 | 205.98 | 0.01 36 | 0.06 01 |
| | Control vs. CTN-067 M6 | 8 | 1149.6 3 | 325.90 | 1 1 | 667.27 | 314.52 | 512.38 | 204.55 | 0.02 15 | 0.09 13 |
| | CTN-067 M0 vs. CTN-067 M3 | 1 1 | 535.45 | 521.24 | 1 1 | 622.64 | 258.07 | - 151.15 | 149.45 | 0.32 46 | 0.74 49 |
| | CTN-067 M0 vs. CTN-067 M6 | 1 1 | 535.45 | 521.24 | 1 1 | 667.27 | 314.52 | - 198.96 | 150.99 | 0.20 33 | 0.56 34 |
| | CTN-067 M3 vs. | 1 1 | 622.64 | 258.07 | 1 1 | 667.27 | 314.52 | -47.82 | 133.51 | 0.72 42 | 0.98 38 |

| | | | | | | | | | | |
|---|---|---|---|---|---|---|---|---|---|---|
| | CTN-067 M6 | | | | | | | | | |
| CD3+CD4+/CD57 \| Freq. of Parent | Control vs. CTN-067 M0 | 8 | 6.14 | 3.16 | 11 | 4.12 | 1.58 | 2.20 | 1.36 | 0.1229 | 0.3945 |
| | Control vs. CTN-067 M3 | 8 | 6.14 | 3.16 | 11 | 3.91 | 1.65 | 2.25 | 1.32 | 0.1045 | 0.3486 |
| | Control vs. CTN-067 M6 | 8 | 6.14 | 3.16 | 11 | 4.61 | 2.36 | 1.53 | 1.32 | 0.2585 | 0.6553 |
| | CTN-067 M0 vs. CTN-067 M3 | 11 | 4.12 | 1.58 | 11 | 3.91 | 1.65 | 0.04 | 0.37 | 0.9055 | 0.9994 |
| | CTN-067 M0 vs. CTN-067 M6 | 11 | 4.12 | 1.58 | 11 | 4.61 | 2.36 | -0.67 | 0.37 | 0.0903 | 0.3108 |
| | CTN-067 M3 vs. CTN-067 M6 | 11 | 3.91 | 1.65 | 11 | 4.61 | 2.36 | -0.71 | 0.32 | 0.0387 | 0.1535 |
| CD3+CD4+ \| Geometric Mean (RB705-A :: CCR2) | Control vs. CTN-067 M0 | 8 | 1494.75 | 360.84 | 11 | 1282.27 | 1437.94 | -398.92 | 507.26 | 0.4413 | 0.8597 |
| | Control vs. CTN-067 M3 | 8 | 1494.75 | 360.84 | 11 | 833.36 | 332.70 | 266.92 | 428.09 | 0.5404 | 0.9233 |
| | Control vs. CTN-067 M6 | 8 | 1494.75 | 360.84 | 11 | 939.91 | 275.11 | 171.16 | 425.01 | 0.6917 | 0.9773 |
| | CTN-067 M0 vs. CTN-067 M3 | 11 | 1282.27 | 1437.94 | 11 | 833.36 | 332.70 | 665.84 | 321.28 | 0.0521 | 0.1978 |
| | CTN-067 M0 vs. CTN-067 M6 | 11 | 1282.27 | 1437.94 | 11 | 939.91 | 275.11 | 570.08 | 324.48 | 0.0950 | 0.3236 |
| | CTN-067 M3 vs. CTN-067 M6 | 11 | 833.36 | 332.70 | 11 | 939.91 | 275.11 | -95.76 | 288.09 | 0.7432 | 0.9869 |
| CD3+CD4+/CCR2+ \| Freq. of Parent | Control vs. CTN-067 M0 | 8 | 20.65 | 5.86 | 11 | 18.50 | 20.75 | -7.82 | 7.29 | 0.2974 | 0.7105 |
| | Control vs. CTN-067 M3 | 8 | 20.65 | 5.86 | 11 | 11.21 | 5.89 | 2.94 | 6.21 | 0.6412 | 0.9640 |
| | Control vs. CTN-067 M6 | 8 | 20.65 | 5.86 | 11 | 12.65 | 5.75 | 1.67 | 6.16 | 0.7894 | 0.9928 |
| | CTN-067 M0 vs. CTN-067 M3 | 11 | 18.50 | 20.75 | 11 | 11.21 | 5.89 | 10.75 | 4.43 | 0.0252 | 0.1051 |
| | CTN-067 M0 vs. CTN-067 M6 | 11 | 18.50 | 20.75 | 11 | 12.65 | 5.75 | 9.49 | 4.47 | 0.0473 | 0.1822 |

| | | | | | | | | | | |
|---|---|---|---|---|---|---|---|---|---|---|
| | CTN-067 M3 vs. CTN-067 M6 | 11 | 11.21 | 5.89 | 11 | 12.65 | 5.75 | -1.27 | 3.95 | 0.7514 | 0.9881 |
| CD3+CD4+ \| Geometric Mean (RB780-A :: HLA-DR) | Control vs. CTN-067 M0 | 8 | 2262.50 | 570.17 | 11 | 2345.09 | 927.55 | 149.14 | 478.24 | 0.7586 | 0.9891 |
| | Control vs. CTN-067 M3 | 8 | 2262.50 | 570.17 | 11 | 2225.45 | 747.09 | 56.81 | 413.64 | 0.8922 | 0.9990 |
| | Control vs. CTN-067 M6 | 8 | 2262.50 | 570.17 | 11 | 2329.00 | 887.50 | -57.27 | 411.17 | 0.8907 | 0.9990 |
| | CTN-067 M0 vs. CTN-067 M3 | 11 | 2345.09 | 927.55 | 11 | 2225.45 | 747.09 | -92.33 | 269.73 | 0.7359 | 0.9858 |
| | CTN-067 M0 vs. CTN-067 M6 | 11 | 2345.09 | 927.55 | 11 | 2329.00 | 887.50 | -206.41 | 272.70 | 0.4584 | 0.8726 |
| | CTN-067 M3 vs. CTN-067 M6 | 11 | 2225.45 | 747.09 | 11 | 2329.00 | 887.50 | -114.08 | 238.80 | 0.6383 | 0.9631 |
| CD3+CD4+/HLA-DR \| Freq. of Parent | Control vs. CTN-067 M0 | 8 | 11.51 | 5.44 | 11 | 14.07 | 7.27 | -6.28 | 3.40 | 0.0804 | 0.2833 |
| | Control vs. CTN-067 M3 | 8 | 11.51 | 5.44 | 11 | 11.85 | 5.51 | -2.93 | 3.17 | 0.3668 | 0.7922 |
| | Control vs. CTN-067 M6 | 8 | 11.51 | 5.44 | 11 | 11.92 | 4.76 | -2.94 | 3.16 | 0.3633 | 0.7885 |
| | CTN-067 M0 vs. CTN-067 M3 | 11 | 14.07 | 7.27 | 11 | 11.85 | 5.51 | 3.35 | 1.31 | 0.0194 | 0.0830 |
| | CTN-067 M0 vs. CTN-067 M6 | 11 | 14.07 | 7.27 | 11 | 11.92 | 4.76 | 3.33 | 1.33 | 0.0211 | 0.0898 |
| | CTN-067 M3 vs. CTN-067 M6 | 11 | 11.85 | 5.51 | 11 | 11.92 | 4.76 | -0.01 | 1.14 | 0.9904 | 1.0000 |
| CD3+CD4+ \| Geometric Mean (RY586-A :: CXCR4) | Control vs. CTN-067 M0 | 8 | 4483.38 | 853.20 | 11 | 27766.82 | 12507.46 | -20558.00 | 7291.82 | 0.0110 | 0.0493 |
| | Control vs. CTN-067 M3 | 8 | 4483.38 | 853.20 | 11 | 15232.45 | 13454.53 | -8315.72 | 6100.48 | 0.1888 | 0.5363 |
| | Control vs. CTN-067 M6 | 8 | 4483.38 | 853.20 | 11 | 12299.09 | 11017.17 | -5396.90 | 6053.98 | 0.3838 | 0.8093 |
| | CTN-067 M0 vs. CTN-067 M3 | 11 | 27766.82 | 12507.46 | 11 | 15232.45 | 13454.53 | 12242.00 | 4949.89 | 0.0230 | 0.0969 |
| | CTN-067 M0 vs. | 11 | 27766.82 | 12507.46 | 11 | 12299.09 | 11017.17 | 15161.00 | 4994.65 | 0.0068 | 0.0316 |

| | | | | | | | | | | |
|---|---|---|---|---|---|---|---|---|---|---|
| | CTN-067 M6 | | | | | | | | | |
| | CTN-067 M3 vs. CTN-067 M6 | 11 | 15232.45 | 13454.53 | 11 | 12299.09 | 11017.17 | 2918.82 | 4488.25 | 0.5233 | 0.9141 |
| CD3+CD4+/CXCR4+ \| Freq. of Parent | Control vs. CTN-067 M0 | 8 | 58.95 | 4.15 | 11 | 91.17 | 4.91 | -28.58 | 7.93 | 0.0019 | 0.0093 |
| | Control vs. CTN-067 M3 | 8 | 58.95 | 4.15 | 11 | 75.65 | 18.63 | -12.76 | 6.65 | 0.0702 | 0.2538 |
| | Control vs. CTN-067 M6 | 8 | 58.95 | 4.15 | 11 | 75.73 | 15.55 | -12.82 | 6.60 | 0.0672 | 0.2447 |
| | CTN-067 M0 vs. CTN-067 M3 | 11 | 91.17 | 4.91 | 11 | 75.65 | 18.63 | 15.81 | 5.21 | 0.0068 | 0.0317 |
| | CTN-067 M0 vs. CTN-067 M6 | 11 | 91.17 | 4.91 | 11 | 75.73 | 15.55 | 15.75 | 5.26 | 0.0074 | 0.0344 |
| | CTN-067 M3 vs. CTN-067 M6 | 11 | 75.65 | 18.63 | 11 | 75.73 | 15.55 | -0.06 | 4.70 | 0.9903 | 1.0000 |
| CD3+CD4+ \| Geometric Mean (RY610-A :: CD69) | Control vs. CTN-067 M0 | 8 | 181.27 | 439.51 | 11 | 621.56 | 527.63 | 23.06 | 344.78 | 0.9474 | 0.9999 |
| | Control vs. CTN-067 M3 | 8 | 181.27 | 439.51 | 11 | 726.14 | 639.96 | -254.19 | 292.26 | 0.3953 | 0.8203 |
| | Control vs. CTN-067 M6 | 8 | 181.27 | 439.51 | 11 | 765.64 | 582.70 | -302.27 | 290.22 | 0.3107 | 0.7278 |
| | CTN-067 M0 vs. CTN-067 M3 | 11 | 621.56 | 527.63 | 11 | 726.14 | 639.96 | -277.24 | 213.13 | 0.2089 | 0.5735 |
| | CTN-067 M0 vs. CTN-067 M6 | 11 | 621.56 | 527.63 | 11 | 765.64 | 582.70 | -325.32 | 215.30 | 0.1472 | 0.4507 |
| | CTN-067 M3 vs. CTN-067 M6 | 11 | 726.14 | 639.96 | 11 | 765.64 | 582.70 | -48.08 | 190.49 | 0.8035 | 0.9942 |
| CD3+CD4+ \| Geometric Mean (RY703-A :: PD-1) | Control vs. CTN-067 M0 | 8 | 2037.75 | 517.85 | 11 | 2101.73 | 470.24 | -20.36 | 277.73 | 0.9423 | 0.9999 |
| | Control vs. CTN-067 M3 | 8 | 2037.75 | 517.85 | 11 | 1836.55 | 402.23 | 249.61 | 255.49 | 0.3408 | 0.7640 |
| | Control vs. CTN-067 M6 | 8 | 2037.75 | 517.85 | 11 | 1867.45 | 448.52 | 218.94 | 254.67 | 0.4007 | 0.8252 |
| | CTN-067 M0 vs. CTN-067 M3 | 11 | 2101.73 | 470.24 | 11 | 1836.55 | 402.23 | 269.97 | 116.69 | 0.0320 | 0.1301 |

| | | | | | | | | | | | |
|---|---|---|---|---|---|---|---|---|---|---|---|
| | CTN-067 M0 vs. CTN-067 M6 | 11 | 2101.73 | 470.24 | 11 | 1867.45 | 448.52 | 239.30 | 118.10 | 0.0570 | 0.2135 |
| | CTN-067 M3 vs. CTN-067 M6 | 11 | 1836.55 | 402.23 | 11 | 1867.45 | 448.52 | -30.67 | 101.88 | 0.7666 | 0.9902 |
| CD3+CD4+/PD-1+ \| Freq. of Parent | Control vs. CTN-067 M0 | 8 | 36.81 | 9.53 | 11 | 30.80 | 10.13 | 7.39 | 5.45 | 0.1906 | 0.5398 |
| | Control vs. CTN-067 M3 | 8 | 36.81 | 9.53 | 11 | 27.98 | 9.25 | 9.77 | 4.98 | 0.0644 | 0.2363 |
| | Control vs. CTN-067 M6 | 8 | 36.81 | 9.53 | 11 | 27.58 | 6.28 | 10.15 | 4.96 | 0.0548 | 0.2064 |
| | CTN-067 M0 vs. CTN-067 M3 | 11 | 30.80 | 10.13 | 11 | 27.98 | 9.25 | 2.38 | 2.37 | 0.3274 | 0.7483 |
| | CTN-067 M0 vs. CTN-067 M6 | 11 | 30.80 | 10.13 | 11 | 27.58 | 6.28 | 2.76 | 2.40 | 0.2640 | 0.6636 |
| | CTN-067 M3 vs. CTN-067 M6 | 11 | 27.98 | 9.25 | 11 | 27.58 | 6.28 | 0.38 | 2.07 | 0.8571 | 0.9978 |
| CD3+CD4+/CCR7- \| Freq. of Parent | Control vs. CTN-067 M0 | 8 | 29.43 | 10.79 | 11 | 21.41 | 8.60 | 6.42 | 5.17 | 0.2289 | 0.6082 |
| | Control vs. CTN-067 M3 | 8 | 29.43 | 10.79 | 11 | 21.27 | 9.57 | 6.22 | 5.05 | 0.2333 | 0.6155 |
| | Control vs. CTN-067 M6 | 8 | 29.43 | 10.79 | 11 | 20.87 | 8.67 | 6.60 | 5.05 | 0.2064 | 0.5692 |
| | CTN-067 M0 vs. CTN-067 M3 | 11 | 21.41 | 8.60 | 11 | 21.27 | 9.57 | -0.21 | 1.14 | 0.8591 | 0.9979 |
| | CTN-067 M0 vs. CTN-067 M6 | 11 | 21.41 | 8.60 | 11 | 20.87 | 8.67 | 0.18 | 1.16 | 0.8797 | 0.9987 |
| | CTN-067 M3 vs. CTN-067 M6 | 11 | 21.27 | 9.57 | 11 | 20.87 | 8.67 | 0.38 | 0.99 | 0.7032 | 0.9797 |
| CD3+CD4+/CD28- \| Freq. of Parent | Control vs. CTN-067 M0 | 8 | 7.25 | 4.64 | 11 | 4.39 | 1.96 | 2.62 | 1.86 | 0.1760 | 0.5112 |
| | Control vs. CTN-067 M3 | 8 | 7.25 | 4.64 | 11 | 5.17 | 2.49 | 1.70 | 1.83 | 0.3630 | 0.7882 |
| | Control vs. CTN-067 M6 | 8 | 7.25 | 4.64 | 11 | 4.39 | 2.67 | 2.48 | 1.83 | 0.1916 | 0.5417 |
| | CTN-067 M0 vs. | 11 | 4.39 | 1.96 | 11 | 5.17 | 2.49 | -0.91 | 0.37 | 0.0243 | 0.1017 |

| | | | | | | | | | | | |
|---|---|---|---|---|---|---|---|---|---|---|---|
| | CTN-067 M3 | | | | | | | | | | |
| | CTN-067 M0 vs. CTN-067 M6 | 1 1 | 4.39 | 1.96 | 1 1 | 4.39 | 2.67 | -0.14 | 0.38 | 0.70 73 | 0.98 06 |
| | CTN-067 M3 vs. CTN-067 M6 | 1 1 | 5.17 | 2.49 | 1 1 | 4.39 | 2.67 | 0.77 | 0.32 | 0.02 80 | 0.11 56 |
| CD3+CD4+/CD38- \| Freq. of Parent | Control vs. CTN-067 M0 | 8 | 57.76 | 7.80 | 1 1 | 43.05 | 12.91 | 9.20 | 6.36 | 0.16 42 | 0.48 71 |
| | Control vs. CTN-067 M3 | 8 | 57.76 | 7.80 | 1 1 | 41.58 | 14.32 | 11.55 | 6.04 | 0.07 11 | 0.25 64 |
| | Control vs. CTN-067 M6 | 8 | 57.76 | 7.80 | 1 1 | 44.07 | 13.86 | 9.10 | 6.03 | 0.14 77 | 0.45 17 |
| | CTN-067 M0 vs. CTN-067 M3 | 1 1 | 43.05 | 12.91 | 1 1 | 41.58 | 14.32 | 2.34 | 2.10 | 0.27 88 | 0.68 50 |
| | CTN-067 M0 vs. CTN-067 M6 | 1 1 | 43.05 | 12.91 | 1 1 | 44.07 | 13.86 | -0.10 | 2.13 | 0.96 19 | 1.00 00 |
| | CTN-067 M3 vs. CTN-067 M6 | 1 1 | 41.58 | 14.32 | 1 1 | 44.07 | 13.86 | -2.45 | 1.83 | 0.19 65 | 0.55 08 |
| CD3+CD4+/CD45RA- \| Freq. of Parent | Control vs. CTN-067 M0 | 8 | 68.64 | 8.34 | 1 1 | 60.70 | 14.46 | 0.19 | 7.07 | 0.97 86 | 1.00 00 |
| | Control vs. CTN-067 M3 | 8 | 68.64 | 8.34 | 1 1 | 57.43 | 16.19 | 5.02 | 6.62 | 0.45 80 | 0.87 23 |
| | Control vs. CTN-067 M6 | 8 | 68.64 | 8.34 | 1 1 | 59.17 | 14.86 | 3.35 | 6.61 | 0.61 81 | 0.95 64 |
| | CTN-067 M0 vs. CTN-067 M3 | 1 1 | 60.70 | 14.46 | 1 1 | 57.43 | 16.19 | 4.83 | 2.62 | 0.08 15 | 0.28 65 |
| | CTN-067 M0 vs. CTN-067 M6 | 1 1 | 60.70 | 14.46 | 1 1 | 59.17 | 14.86 | 3.16 | 2.66 | 0.24 91 | 0.64 09 |
| | CTN-067 M3 vs. CTN-067 M6 | 1 1 | 57.43 | 16.19 | 1 1 | 59.17 | 14.86 | -1.67 | 2.28 | 0.47 42 | 0.88 38 |
| CD3+CD4+/CD57- \| Freq. of Parent | Control vs. CTN-067 M0 | 8 | 93.86 | 3.15 | 1 1 | 95.88 | 1.58 | -2.21 | 1.36 | 0.11 98 | 0.38 70 |
| | Control vs. CTN-067 M3 | 8 | 93.86 | 3.15 | 1 1 | 96.07 | 1.64 | -2.23 | 1.31 | 0.10 54 | 0.35 08 |
| | Control vs. CTN-067 M6 | 8 | 93.86 | 3.15 | 1 1 | 95.38 | 2.37 | -1.53 | 1.31 | 0.25 68 | 0.65 28 |

| | | | | | | | | | | |
|---|---|---|---|---|---|---|---|---|---|---|
| | CTN-067 M0 vs. CTN-067 M3 | 11 | 95.88 | 1.58 | 11 | 96.07 | 1.64 | -0.02 | 0.37 | 0.9618 | 1.0000 |
| | CTN-067 M0 vs. CTN-067 M6 | 11 | 95.88 | 1.58 | 11 | 95.38 | 2.37 | 0.68 | 0.38 | 0.0857 | 0.2981 |
| | CTN-067 M3 vs. CTN-067 M6 | 11 | 96.07 | 1.64 | 11 | 95.38 | 2.37 | 0.70 | 0.32 | 0.0428 | 0.1674 |
| CD3+CD4+/HLA-DR- \| Freq. of Parent | Control vs. CTN-067 M0 | 8 | 88.49 | 5.44 | 11 | 85.93 | 7.26 | 6.28 | 3.40 | 0.0801 | 0.2824 |
| | Control vs. CTN-067 M3 | 8 | 88.49 | 5.44 | 11 | 88.15 | 5.50 | 2.93 | 3.17 | 0.3659 | 0.7912 |
| | Control vs. CTN-067 M6 | 8 | 88.49 | 5.44 | 11 | 88.08 | 4.76 | 2.94 | 3.16 | 0.3634 | 0.7886 |
| | CTN-067 M0 vs. CTN-067 M3 | 11 | 85.93 | 7.26 | 11 | 88.15 | 5.50 | -3.35 | 1.31 | 0.0192 | 0.0825 |
| | CTN-067 M0 vs. CTN-067 M6 | 11 | 85.93 | 7.26 | 11 | 88.08 | 4.76 | -3.34 | 1.32 | 0.0208 | 0.0885 |
| | CTN-067 M3 vs. CTN-067 M6 | 11 | 88.15 | 5.50 | 11 | 88.08 | 4.76 | 0.01 | 1.14 | 0.9948 | 1.0000 |
| CD3+CD4+/CCR7+CD45RA+ \| Freq. of Parent | Control vs. CTN-067 M0 | 8 | 29.46 | 7.68 | 11 | 37.65 | 14.26 | -0.51 | 6.92 | 0.9422 | 0.9999 |
| | Control vs. CTN-067 M3 | 8 | 29.46 | 7.68 | 11 | 40.99 | 16.37 | -5.31 | 6.50 | 0.4243 | 0.8459 |
| | Control vs. CTN-067 M6 | 8 | 29.46 | 7.68 | 11 | 39.31 | 14.88 | -3.70 | 6.48 | 0.5752 | 0.9397 |
| | CTN-067 M0 vs. CTN-067 M3 | 11 | 37.65 | 14.26 | 11 | 40.99 | 16.37 | -4.80 | 2.52 | 0.0718 | 0.2584 |
| | CTN-067 M0 vs. CTN-067 M6 | 11 | 37.65 | 14.26 | 11 | 39.31 | 14.88 | -3.19 | 2.55 | 0.2259 | 0.6032 |
| | CTN-067 M3 vs. CTN-067 M6 | 11 | 40.99 | 16.37 | 11 | 39.31 | 14.88 | 1.61 | 2.19 | 0.4715 | 0.8819 |
| CD3+CD4+/CCR7+CD45RA- \| Freq. of Parent | Control vs. CTN-067 M0 | 8 | 41.14 | 14.55 | 11 | 40.97 | 9.20 | -5.66 | 6.33 | 0.3822 | 0.8077 |
| | Control vs. CTN-067 M3 | 8 | 41.14 | 14.55 | 11 | 37.74 | 9.47 | -0.74 | 6.02 | 0.9029 | 0.9993 |

| | | | | | | | | | | |
|---|---|---|---|---|---|---|---|---|---|---|
| | Control vs. CTN-067 M6 | 8 | 41.14 | 14.55 | 11 | 39.83 | 8.97 | -2.75 | 6.01 | 0.6521 | 0.9672 |
| | CTN-067 M0 vs. CTN-067 M3 | 11 | 40.97 | 9.20 | 11 | 37.74 | 9.47 | 4.92 | 2.08 | 0.0289 | 0.1186 |
| | CTN-067 M0 vs. CTN-067 M6 | 11 | 40.97 | 9.20 | 11 | 39.83 | 8.97 | 2.91 | 2.11 | 0.1828 | 0.5248 |
| | CTN-067 M3 vs. CTN-067 M6 | 11 | 37.74 | 9.47 | 11 | 39.83 | 8.97 | -2.01 | 1.81 | 0.2811 | 0.6883 |
| CD3+CD4+/CCR7-CD45RA+ \| Freq. of Parent | Control vs. CTN-067 M0 | 8 | 1.92 | 1.36 | 11 | 1.67 | 1.33 | 0.27 | 0.67 | 0.6910 | 0.9771 |
| | Control vs. CTN-067 M3 | 8 | 1.92 | 1.36 | 11 | 1.58 | 0.95 | 0.28 | 0.65 | 0.6696 | 0.9720 |
| | Control vs. CTN-067 M6 | 8 | 1.92 | 1.36 | 11 | 1.50 | 1.06 | 0.36 | 0.65 | 0.5835 | 0.9433 |
| | CTN-067 M0 vs. CTN-067 M3 | 11 | 1.67 | 1.33 | 11 | 1.58 | 0.95 | 0.01 | 0.18 | 0.9521 | 0.9999 |
| | CTN-067 M0 vs. CTN-067 M6 | 11 | 1.67 | 1.33 | 11 | 1.50 | 1.06 | 0.09 | 0.18 | 0.6149 | 0.9553 |
| | CTN-067 M3 vs. CTN-067 M6 | 11 | 1.58 | 0.95 | 11 | 1.50 | 1.06 | 0.08 | 0.15 | 0.6046 | 0.9516 |
| CD3+CD4+/CCR7-CD45RA- \| Freq. of Parent | Control vs. CTN-067 M0 | 8 | 27.48 | 9.90 | 11 | 19.73 | 8.43 | 6.13 | 4.91 | 0.2264 | 0.6041 |
| | Control vs. CTN-067 M3 | 8 | 27.48 | 9.90 | 11 | 19.71 | 9.26 | 5.88 | 4.78 | 0.2334 | 0.6157 |
| | Control vs. CTN-067 M6 | 8 | 27.48 | 9.90 | 11 | 19.36 | 8.35 | 6.23 | 4.78 | 0.2079 | 0.5717 |
| | CTN-067 M0 vs. CTN-067 M3 | 11 | 19.73 | 8.43 | 11 | 19.71 | 9.26 | -0.25 | 1.16 | 0.8316 | 0.9963 |
| | CTN-067 M0 vs. CTN-067 M6 | 11 | 19.73 | 8.43 | 11 | 19.36 | 8.35 | 0.09 | 1.17 | 0.9367 | 0.9998 |
| | CTN-067 M3 vs. CTN-067 M6 | 11 | 19.71 | 9.26 | 11 | 19.36 | 8.35 | 0.34 | 1.00 | 0.7358 | 0.9858 |
| CD3+CD4+/CD28+CD57+ \| Freq. of Parent | Control vs. CTN-067 M0 | 8 | 5.18 | 2.52 | 11 | 3.73 | 1.54 | 1.77 | 1.18 | 0.1515 | 0.4602 |

| | | | | | | | | | | |
|---|---|---|---|---|---|---|---|---|---|---|
| | Control vs. CTN-067 M3 | 8 | 5.18 | 2.52 | 1 1 | 3.55 | 1.57 | 1.78 | 1.14 | 0.13 45 | 0.42 20 |
| | Control vs. CTN-067 M6 | 8 | 5.18 | 2.52 | 1 1 | 4.28 | 2.28 | 1.04 | 1.14 | 0.37 23 | 0.79 79 |
| | CTN-067 M0 vs. CTN-067 M3 | 1 1 | 3.73 | 1.54 | 1 1 | 3.55 | 1.57 | 0.01 | 0.34 | 0.97 80 | 1.00 00 |
| | CTN-067 M0 vs. CTN-067 M6 | 1 1 | 3.73 | 1.54 | 1 1 | 4.28 | 2.28 | -0.73 | 0.35 | 0.04 98 | 0.19 04 |
| | CTN-067 M3 vs. CTN-067 M6 | 1 1 | 3.55 | 1.57 | 1 1 | 4.28 | 2.28 | -0.74 | 0.30 | 0.02 29 | 0.09 66 |
| CD3+CD4+/CD28+CD 57- \| Freq. of Parent | Control vs. CTN-067 M0 | 8 | 87.58 | 5.36 | 1 1 | 91.88 | 2.21 | -4.39 | 2.18 | 0.05 80 | 0.21 65 |
| | Control vs. CTN-067 M3 | 8 | 87.58 | 5.36 | 1 1 | 91.30 | 2.83 | -3.50 | 2.11 | 0.11 41 | 0.37 28 |
| | Control vs. CTN-067 M6 | 8 | 87.58 | 5.36 | 1 1 | 91.32 | 3.32 | -3.50 | 2.11 | 0.11 34 | 0.37 12 |
| | CTN-067 M0 vs. CTN-067 M3 | 1 1 | 91.88 | 2.21 | 1 1 | 91.30 | 2.83 | 0.89 | 0.55 | 0.12 28 | 0.39 43 |
| | CTN-067 M0 vs. CTN-067 M6 | 1 1 | 91.88 | 2.21 | 1 1 | 91.32 | 3.32 | 0.89 | 0.56 | 0.12 83 | 0.40 74 |
| | CTN-067 M3 vs. CTN-067 M6 | 1 1 | 91.30 | 2.83 | 1 1 | 91.32 | 3.32 | 0.00 | 0.48 | 0.99 56 | 1.00 00 |
| CD3+CD4+/CD28-CD57+ \| Freq. of Parent | Control vs. CTN-067 M0 | 8 | 0.96 | 1.09 | 1 1 | 0.40 | 0.42 | 0.44 | 0.40 | 0.28 53 | 0.69 41 |
| | Control vs. CTN-067 M3 | 8 | 0.96 | 1.09 | 1 1 | 0.36 | 0.38 | 0.47 | 0.40 | 0.24 82 | 0.63 95 |
| | Control vs. CTN-067 M6 | 8 | 0.96 | 1.09 | 1 1 | 0.34 | 0.38 | 0.50 | 0.40 | 0.22 49 | 0.60 14 |
| | CTN-067 M0 vs. CTN-067 M3 | 1 1 | 0.40 | 0.42 | 1 1 | 0.36 | 0.38 | 0.03 | 0.06 | 0.61 67 | 0.95 60 |
| | CTN-067 M0 vs. CTN-067 M6 | 1 1 | 0.40 | 0.42 | 1 1 | 0.34 | 0.38 | 0.06 | 0.06 | 0.38 00 | 0.80 55 |
| | CTN-067 M3 vs. CTN-067 M6 | 1 1 | 0.36 | 0.38 | 1 1 | 0.34 | 0.38 | 0.02 | 0.05 | 0.64 83 | 0.96 61 |

| | | | | | | | | | | |
|---|---|---|---|---|---|---|---|---|---|---|
| CD3+CD4+/CD28-CD57- \| Freq. of Parent | Control vs. CTN-067 M0 | 8 | 6.30 | 4.19 | 11 | 4.00 | 1.66 | 2.18 | 1.67 | 0.2076 | 0.5712 |
| | Control vs. CTN-067 M3 | 8 | 6.30 | 4.19 | 11 | 4.81 | 2.14 | 1.23 | 1.64 | 0.4600 | 0.8738 |
| | Control vs. CTN-067 M6 | 8 | 6.30 | 4.19 | 11 | 4.06 | 2.34 | 1.98 | 1.64 | 0.2419 | 0.6295 |
| | CTN-067 M0 vs. CTN-067 M3 | 11 | 4.00 | 1.66 | 11 | 4.81 | 2.14 | -0.94 | 0.34 | 0.0130 | 0.0578 |
| | CTN-067 M0 vs. CTN-067 M6 | 11 | 4.00 | 1.66 | 11 | 4.06 | 2.34 | -0.20 | 0.35 | 0.5683 | 0.9367 |
| | CTN-067 M3 vs. CTN-067 M6 | 11 | 4.81 | 2.14 | 11 | 4.06 | 2.34 | 0.74 | 0.30 | 0.0226 | 0.0952 |
| CD3+CD4+/CD38+HLA-DR+ \| Freq. of Parent | Control vs. CTN-067 M0 | 8 | 4.31 | 2.42 | 11 | 6.80 | 2.28 | -2.93 | 1.55 | 0.0739 | 0.2647 |
| | Control vs. CTN-067 M3 | 8 | 4.31 | 2.42 | 11 | 5.56 | 2.24 | -1.62 | 1.38 | 0.2554 | 0.6506 |
| | Control vs. CTN-067 M6 | 8 | 4.31 | 2.42 | 11 | 5.70 | 2.92 | -1.76 | 1.37 | 0.2162 | 0.5865 |
| | CTN-067 M0 vs. CTN-067 M3 | 11 | 6.80 | 2.28 | 11 | 5.56 | 2.24 | 1.31 | 0.76 | 0.1030 | 0.3446 |
| | CTN-067 M0 vs. CTN-067 M6 | 11 | 6.80 | 2.28 | 11 | 5.70 | 2.92 | 1.17 | 0.77 | 0.1466 | 0.4494 |
| | CTN-067 M3 vs. CTN-067 M6 | 11 | 5.56 | 2.24 | 11 | 5.70 | 2.92 | -0.14 | 0.67 | 0.8379 | 0.9967 |
| CD3+CD4+/CD38+HLA-DR- \| Freq. of Parent | Control vs. CTN-067 M0 | 8 | 37.91 | 6.23 | 11 | 50.15 | 13.68 | -5.75 | 6.14 | 0.3609 | 0.7860 |
| | Control vs. CTN-067 M3 | 8 | 37.91 | 6.23 | 11 | 52.85 | 14.77 | -9.63 | 5.84 | 0.1157 | 0.3769 |
| | Control vs. CTN-067 M6 | 8 | 37.91 | 6.23 | 11 | 50.23 | 13.05 | -7.06 | 5.83 | 0.2408 | 0.6277 |
| | CTN-067 M0 vs. CTN-067 M3 | 11 | 50.15 | 13.68 | 11 | 52.85 | 14.77 | -3.88 | 2.00 | 0.0676 | 0.2461 |
| | CTN-067 M0 vs. CTN-067 M6 | 11 | 50.15 | 13.68 | 11 | 50.23 | 13.05 | -1.31 | 2.03 | 0.5256 | 0.9154 |
| | CTN-067 M3 vs. | 11 | 52.85 | 14.77 | 11 | 50.23 | 13.05 | 2.57 | 1.74 | 0.1563 | 0.4704 |

| | | | | | | | | | | | |
|---|---|---|---|---|---|---|---|---|---|---|---|
| | CTN-067 M6 | | | | | | | | | | |
| CD3+CD4+/CD38-HLA-DR+ \| Freq. of Parent | Control vs. CTN-067 M0 | 8 | 7.22 | 3.49 | 11 | 7.28 | 5.68 | -2.99 | 2.38 | 0.2251 | 0.6017 |
| | Control vs. CTN-067 M3 | 8 | 7.22 | 3.49 | 11 | 6.31 | 4.23 | -1.11 | 2.26 | 0.6291 | 0.9602 |
| | Control vs. CTN-067 M6 | 8 | 7.22 | 3.49 | 11 | 6.22 | 3.57 | -0.98 | 2.25 | 0.6698 | 0.9720 |
| | CTN-067 M0 vs. CTN-067 M3 | 11 | 7.28 | 5.68 | 11 | 6.31 | 4.23 | 1.88 | 0.81 | 0.0316 | 0.1285 |
| | CTN-067 M0 vs. CTN-067 M6 | 11 | 7.28 | 5.68 | 11 | 6.22 | 3.57 | 2.01 | 0.82 | 0.0240 | 0.1006 |
| | CTN-067 M3 vs. CTN-067 M6 | 11 | 6.31 | 4.23 | 11 | 6.22 | 3.57 | 0.13 | 0.70 | 0.8530 | 0.9976 |
| CD3+CD4+/CD38-HLA-DR- \| Freq. of Parent | Control vs. CTN-067 M0 | 8 | 50.54 | 9.80 | 11 | 35.78 | 9.45 | 12.09 | 5.79 | 0.0502 | 0.1918 |
| | Control vs. CTN-067 M3 | 8 | 50.54 | 9.80 | 11 | 35.29 | 11.39 | 12.58 | 5.48 | 0.0332 | 0.1342 |
| | Control vs. CTN-067 M6 | 8 | 50.54 | 9.80 | 11 | 37.85 | 11.60 | 10.02 | 5.47 | 0.0827 | 0.2897 |
| | CTN-067 M0 vs. CTN-067 M3 | 11 | 35.78 | 9.45 | 11 | 35.29 | 11.39 | 0.49 | 1.96 | 0.8063 | 0.9944 |
| | CTN-067 M0 vs. CTN-067 M6 | 11 | 35.78 | 9.45 | 11 | 37.85 | 11.60 | -2.08 | 1.99 | 0.3096 | 0.7263 |
| | CTN-067 M3 vs. CTN-067 M6 | 11 | 35.29 | 11.39 | 11 | 37.85 | 11.60 | -2.56 | 1.71 | 0.1497 | 0.4562 |
