## Supplemental Table 12 for "Chronic opioid-associated immune dysregulation among people living with HIV"

**Supplementary Table 12.** CD3+CD8+ T cell Flow Data

| Marker | Comparison (A vs B) | Group A | | | Group B | | | Viral Load adjusted<br>Group A – Group B | | | |
|---|---|---|---|---|---|---|---|---|---|---|---|
| | | n | Mean | SD | N | Mean | SD | Estimate | SE | p | adj p |
| CD3+ \| Geometric Mean (BV750-A :: CD8) | Control vs. CTN-067 M0 | 8 | 6193.250 | 3098.890 | 11 | 11418.090 | 10174.410 | 81.695 | 3673.700 | 0.9825 | 1.0000 |
| | Control vs. CTN-067 M3 | 8 | 6193.250 | 3098.890 | 11 | 7639.360 | 5522.160 | 1207.750 | 3275.820 | 0.7164 | 0.9824 |
| | Control vs. CTN-067 M6 | 8 | 6193.250 | 3098.890 | 11 | 7176.550 | 4543.340 | 1538.680 | 3260.830 | 0.6424 | 0.9644 |
| | CTN-067 M0 vs. CTN-067 M3 | 11 | 11418.090 | 10174.410 | 11 | 7639.360 | 5522.160 | 1126.050 | 1814.900 | 0.5423 | 0.9243 |
| | CTN-067 M0 vs. CTN-067 M6 | 11 | 11418.090 | 10174.410 | 11 | 7176.550 | 4543.340 | 1456.990 | 1836.020 | 0.4373 | 0.8565 |
| | CTN-067 M3 vs. CTN-067 M6 | 11 | 7639.360 | 5522.160 | 11 | 7176.550 | 4543.340 | 330.930 | 1593.340 | 0.8377 | 0.9967 |
| CD3+CD8+ \| Freq. of Parent | Control vs. CTN-067 M0 | 8 | 38.300 | 12.134 | 11 | 45.164 | 15.950 | -0.246 | 7.488 | 0.9742 | 1.0000 |
| | Control vs. CTN-067 M3 | 8 | 38.300 | 12.134 | 11 | 41.764 | 14.738 | -0.589 | 7.247 | 0.9361 | 0.9998 |
| | Control vs. CTN-067 M6 | 8 | 38.300 | 12.134 | 11 | 41.073 | 14.987 | -0.084 | 7.238 | 0.9909 | 1.0000 |
| | CTN-067 M0 vs. CTN-067 M3 | 11 | 45.164 | 15.950 | 11 | 41.764 | 14.738 | -0.343 | 1.984 | 0.8645 | 0.9981 |
| | CTN-067 M0 vs. CTN-067 M6 | 11 | 45.164 | 15.950 | 11 | 41.073 | 14.987 | 0.161 | 2.009 | 0.9368 | 0.9998 |
| | CTN-067 M3 vs. CTN-067 M6 | 11 | 41.764 | 14.738 | 11 | 41.073 | 14.987 | 0.505 | 1.722 | 0.7726 | 0.9909 |
| CD3+CD8+ \| Geometric Mean (APC-Fire 750-A :: CD36) | Control vs. CTN-067 M0 | 8 | 1202.000 | 308.503 | 11 | 1169.270 | 374.877 | 364.890 | 203.770 | 0.0893 | 0.3081 |
| | Control vs. CTN-067 M3 | 8 | 1202.000 | 308.503 | 11 | 1239.730 | 280.110 | 171.060 | 173.680 | 0.3370 | 0.7596 |
| | Control vs. CTN-067 M6 | 8 | 1202.000 | 308.503 | 11 | 1260.000 | 275.760 | 144.660 | 172.510 | 0.4122 | 0.8355 |
| | CTN-067 M0 vs. CTN-067 M3 | 11 | 1169.270 | 374.877 | 11 | 1239.730 | 280.110 | -193.830 | 122.640 | 0.1305 | 0.4126 |

| | | | | | | | | | | | |
|---|---|---|---|---|---|---|---|---|---|---|---|
| | CTN-067 M0 vs. CTN-067 M6 | 11 | 1169.270 | 374.877 | 11 | 1260.000 | 275.760 | -220.240 | 123.920 | 0.0915 | 0.3142 |
| | CTN-067 M3 vs. CTN-067 M6 | 11 | 1239.730 | 280.110 | 11 | 1260.000 | 275.760 | -26.407 | 109.260 | 0.8116 | 0.9949 |
| CD3+CD8+ \| Geometric Mean (Alexa Fluor 647-A :: GLUT1) | Control vs. CTN-067 M0 | 8 | 6443.630 | 1263.680 | 11 | 6548.640 | 2143.280 | -27.116 | 1584358.000 | 1.0000 | 1.0000 |
| | Control vs. CTN-067 M3 | 8 | 6443.630 | 1263.680 | 11 | 7700.360 | 3346.120 | -1154.000 | 1584358.000 | 0.9994 | 1.0000 |
| | Control vs. CTN-067 M6 | 8 | 6443.630 | 1263.680 | 11 | 7353.640 | 3791.730 | -806.040 | 1584358.000 | 0.9996 | 1.0000 |
| | CTN-067 M0 vs. CTN-067 M3 | 11 | 6548.640 | 2143.280 | 11 | 7700.360 | 3346.120 | -1126.890 | 478.270 | 0.0294 | 0.1204 |
| | CTN-067 M0 vs. CTN-067 M6 | 11 | 6548.640 | 2143.280 | 11 | 7353.640 | 3791.730 | -778.920 | 484.370 | 0.1243 | 0.3978 |
| | CTN-067 M3 vs. CTN-067 M6 | 11 | 7700.360 | 3346.120 | 11 | 7353.640 | 3791.730 | 347.960 | 413.880 | 0.4110 | 0.8345 |
| CD3+CD8+/GLUT1+ \| Freq. of Parent | Control vs. CTN-067 M0 | 8 | 87.525 | 3.323 | 11 | 91.455 | 3.005 | -5.315 | 2.069 | 0.0188 | 0.0808 |
| | Control vs. CTN-067 M3 | 8 | 87.525 | 3.323 | 11 | 90.345 | 3.955 | -3.723 | 1.772 | 0.0492 | 0.1885 |
| | Control vs. CTN-067 M6 | 8 | 87.525 | 3.323 | 11 | 91.473 | 2.362 | -4.826 | 1.760 | 0.0130 | 0.0576 |
| | CTN-067 M0 vs. CTN-067 M3 | 11 | 91.455 | 3.005 | 11 | 90.345 | 3.955 | 1.592 | 1.219 | 0.2072 | 0.5704 |
| | CTN-067 M0 vs. CTN-067 M6 | 11 | 91.455 | 3.005 | 11 | 91.473 | 2.362 | 0.489 | 1.232 | 0.6959 | 0.9782 |
| | CTN-067 M3 vs. CTN-067 M6 | 11 | 90.345 | 3.955 | 11 | 91.473 | 2.362 | -1.103 | 1.084 | 0.3215 | 0.7412 |
| CD3+CD8+ \| Geometric Mean (BUV395-A :: CD28) | Control vs. CTN-067 M0 | 8 | 1839.880 | 741.048 | 11 | 1576.730 | 663.249 | 341.780 | 371.470 | 0.3691 | 0.7945 |
| | Control vs. CTN-067 M3 | 8 | 1839.880 | 741.048 | 11 | 1537.640 | 593.545 | 402.630 | 356.930 | 0.2734 | 0.6773 |
| | Control vs. CTN-067 M6 | 8 | 1839.880 | 741.048 | 11 | 1611.820 | 636.587 | 329.530 | 356.410 | 0.3668 | 0.7922 |
| | CTN-067 M0 | 11 | 1576.730 | 663.249 | 11 | 1537.640 | 593.545 | 60.849 | 108.430 | 0.5812 | 0.9423 |

| | | | | | | | | | | | |
|---|---|---|---|---|---|---|---|---|---|---|---|
| | vs. CTN-067 M3 | | | | | | | | | | |
| | CTN-067 M0 vs. CTN-067 M6 | 11 | 1576.730 | 663.249 | 11 | 1611.820 | 636.587 | -12.251 | 109.790 | 0.9123 | 0.9995 |
| | CTN-067 M3 vs. CTN-067 M6 | 11 | 1537.640 | 593.545 | 11 | 1611.820 | 636.587 | -73.100 | 94.164 | 0.4471 | 0.8642 |
| CD3+CD8+/CD28 \| Freq. of Parent | Control vs. CTN-067 M0 | 8 | 58.763 | 21.363 | 11 | 53.082 | 19.146 | 1.745 | 11.509 | 0.8811 | 0.9987 |
| | Control vs. CTN-067 M3 | 8 | 58.763 | 21.363 | 11 | 60.082 | 16.727 | -1.807 | 10.378 | 0.8636 | 0.9981 |
| | Control vs. CTN-067 M6 | 8 | 58.763 | 21.363 | 11 | 63.136 | 18.985 | -4.690 | 10.336 | 0.6551 | 0.9681 |
| | CTN-067 M0 vs. CTN-067 M3 | 11 | 53.082 | 19.146 | 11 | 60.082 | 16.727 | -3.552 | 5.388 | 0.5177 | 0.9110 |
| | CTN-067 M0 vs. CTN-067 M6 | 11 | 53.082 | 19.146 | 11 | 63.136 | 18.985 | -6.435 | 5.452 | 0.2524 | 0.6461 |
| | CTN-067 M3 vs. CTN-067 M6 | 11 | 60.082 | 16.727 | 11 | 63.136 | 18.985 | -2.883 | 4.720 | 0.5485 | 0.9274 |
| CD3+CD8+ \| Geometric Mean (BUV661-A :: CD27) | Control vs. CTN-067 M0 | 8 | 11121.500 | 4691.740 | 11 | 16519.450 | 6207.430 | -1235.570 | 3556.520 | 0.7321 | 0.9851 |
| | Control vs. CTN-067 M3 | 8 | 11121.500 | 4691.740 | 11 | 17426.730 | 8115.750 | -2993.220 | 3298.680 | 0.3756 | 0.8011 |
| | Control vs. CTN-067 M6 | 8 | 11121.500 | 4691.740 | 11 | 17463.820 | 7307.570 | -3072.590 | 3289.180 | 0.3619 | 0.7871 |
| | CTN-067 M0 vs. CTN-067 M3 | 11 | 16519.450 | 6207.430 | 11 | 17426.730 | 8115.750 | -1757.650 | 1419.420 | 0.2307 | 0.6112 |
| | CTN-067 M0 vs. CTN-067 M6 | 11 | 16519.450 | 6207.430 | 11 | 17463.820 | 7307.570 | -1837.020 | 1436.690 | 0.2164 | 0.5869 |
| | CTN-067 M3 vs. CTN-067 M6 | 11 | 17426.730 | 8115.750 | 11 | 17463.820 | 7307.570 | -79.369 | 1237.770 | 0.9495 | 0.9999 |
| CD3+CD8+/CD27+ \| Freq. of Parent | Control vs. CTN-067 M0 | 8 | 80.775 | 8.117 | 11 | 86.555 | 7.665 | -0.743 | 4.274 | 0.8639 | 0.9981 |
| | Control vs. CTN-067 M3 | 8 | 80.775 | 8.117 | 11 | 86.336 | 8.842 | -1.530 | 4.039 | 0.7090 | 0.9809 |
| | Control vs. CTN-067 M6 | 8 | 80.775 | 8.117 | 11 | 87.145 | 7.440 | -2.389 | 4.030 | 0.5603 | 0.9330 |

| | | | | | | | | | | | |
|---|---|---|---|---|---|---|---|---|---|---|---|
| | CTN-067 M0 vs. CTN-067 M3 | 11 | 86.555 | 7.665 | 11 | 86.336 | 8.842 | -0.787 | 1.482 | 0.6015 | 0.9504 |
| | CTN-067 M0 vs. CTN-067 M6 | 11 | 86.555 | 7.665 | 11 | 87.145 | 7.440 | -1.646 | 1.500 | 0.2862 | 0.6954 |
| | CTN-067 M3 vs. CTN-067 M6 | 11 | 86.336 | 8.842 | 11 | 87.145 | 7.440 | -0.859 | 1.289 | 0.5133 | 0.9085 |
| CD3+CD8+ \| Geometric Mean (BV510-A :: CCR7) | Control vs. CTN-067 M0 | 8 | 2897.250 | 922.702 | 11 | 2381.180 | 1807.620 | 1229.770 | 876.610 | 0.1768 | 0.5128 |
| | Control vs. CTN-067 M3 | 8 | 2897.250 | 922.702 | 11 | 3236.270 | 1948.050 | 292.580 | 808.150 | 0.7213 | 0.9833 |
| | Control vs. CTN-067 M6 | 8 | 2897.250 | 922.702 | 11 | 3444.090 | 1765.220 | 80.683 | 805.610 | 0.9213 | 0.9996 |
| | CTN-067 M0 vs. CTN-067 M3 | 11 | 2381.180 | 1807.620 | 11 | 3236.270 | 1948.050 | -937.190 | 363.600 | 0.0184 | 0.0794 |
| | CTN-067 M0 vs. CTN-067 M6 | 11 | 2381.180 | 1807.620 | 11 | 3444.090 | 1765.220 | -1149.090 | 368.000 | 0.0056 | 0.0263 |
| | CTN-067 M3 vs. CTN-067 M6 | 11 | 3236.270 | 1948.050 | 11 | 3444.090 | 1765.220 | -211.900 | 317.340 | 0.5123 | 0.9079 |
| CD3+CD8+/CCR7 \| Freq. of Parent | Control vs. CTN-067 M0 | 8 | 41.650 | 16.704 | 11 | 39.400 | 14.726 | 0.198 | 8.697 | 0.9820 | 1.0000 |
| | Control vs. CTN-067 M3 | 8 | 41.650 | 16.704 | 11 | 42.964 | 16.575 | -0.375 | 8.415 | 0.9649 | 1.0000 |
| | Control vs. CTN-067 M6 | 8 | 41.650 | 16.704 | 11 | 46.691 | 16.489 | -3.954 | 8.405 | 0.6434 | 0.9647 |
| | CTN-067 M0 vs. CTN-067 M3 | 11 | 39.400 | 14.726 | 11 | 42.964 | 16.575 | -0.574 | 2.312 | 0.8066 | 0.9944 |
| | CTN-067 M0 vs. CTN-067 M6 | 11 | 39.400 | 14.726 | 11 | 46.691 | 16.489 | -4.153 | 2.341 | 0.0921 | 0.3157 |
| | CTN-067 M3 vs. CTN-067 M6 | 11 | 42.964 | 16.575 | 11 | 46.691 | 16.489 | -3.579 | 2.006 | 0.0904 | 0.3112 |
| CD3+CD8+ \| Geometric Mean (BV605-A :: CD45RA) | Control vs. CTN-067 M0 | 8 | 28694.750 | 13682.710 | 11 | 23020.000 | 13473.940 | 8330.880 | 8350.490 | 0.3310 | 0.7525 |
| | Control vs. CTN-067 M3 | 8 | 28694.750 | 13682.710 | 11 | 29609.550 | 19550.480 | 2737.450 | 7912.560 | 0.7332 | 0.9853 |

| | | | | | | | | | | |
|---|---|---|---|---|---|---|---|---|---|---|
| | Control vs. CTN-067 M6 | 8 | 28694.750 | 13682.710 | 11 | 28891.180 | 15358.130 | 3505.340 | 7896.600 | 0.6621 | 0.9700 |
| | CTN-067 M0 vs. CTN-067 M3 | 11 | 23020.000 | 13473.940 | 11 | 29609.550 | 19550.480 | -5593.420 | 2825.420 | 0.0624 | 0.2302 |
| | CTN-067 M0 vs. CTN-067 M6 | 11 | 23020.000 | 13473.940 | 11 | 28891.180 | 15358.130 | -4825.540 | 2860.360 | 0.1079 | 0.3574 |
| | CTN-067 M3 vs. CTN-067 M6 | 11 | 29609.550 | 19550.480 | 11 | 28891.180 | 15358.130 | 767.890 | 2457.250 | 0.7581 | 0.9891 |
| CD3+CD8+/CD45RA \| Freq. of Parent | Control vs. CTN-067 M0 | 8 | 85.488 | 2.932 | 11 | 84.336 | 5.053 | 4.855 | 2.623 | 0.0798 | 0.2817 |
| | Control vs. CTN-067 M3 | 8 | 85.488 | 2.932 | 11 | 84.273 | 5.165 | 3.467 | 2.463 | 0.1754 | 0.5101 |
| | Control vs. CTN-067 M6 | 8 | 85.488 | 2.932 | 11 | 84.327 | 6.187 | 3.340 | 2.457 | 0.1899 | 0.5385 |
| | CTN-067 M0 vs. CTN-067 M3 | 11 | 84.336 | 5.053 | 11 | 84.273 | 5.165 | -1.388 | 0.960 | 0.1644 | 0.4875 |
| | CTN-067 M0 vs. CTN-067 M6 | 11 | 84.336 | 5.053 | 11 | 84.327 | 6.187 | -1.515 | 0.971 | 0.1354 | 0.4240 |
| | CTN-067 M3 vs. CTN-067 M6 | 11 | 84.273 | 5.165 | 11 | 84.327 | 6.187 | -0.127 | 0.836 | 0.8810 | 0.9987 |
| CD3+CD8+ \| Geometric Mean (BV711-A :: CD86) | Control vs. CTN-067 M0 | 8 | 407.625 | 123.235 | 11 | 506.091 | 168.537 | 9.548 | 82.733 | 0.9093 | 0.9994 |
| | Control vs. CTN-067 M3 | 8 | 407.625 | 123.235 | 11 | 440.000 | 103.331 | 30.527 | 75.727 | 0.6914 | 0.9772 |
| | Control vs. CTN-067 M6 | 8 | 407.625 | 123.235 | 11 | 429.818 | 134.860 | 38.466 | 75.467 | 0.6161 | 0.9558 |
| | CTN-067 M0 vs. CTN-067 M3 | 11 | 506.091 | 168.537 | 11 | 440.000 | 103.331 | 20.979 | 35.795 | 0.5647 | 0.9351 |
| | CTN-067 M0 vs. CTN-067 M6 | 11 | 506.091 | 168.537 | 11 | 429.818 | 134.860 | 28.918 | 36.225 | 0.4346 | 0.8543 |
| | CTN-067 M3 vs. CTN-067 M6 | 11 | 440.000 | 103.331 | 11 | 429.818 | 134.860 | 7.939 | 31.275 | 0.8023 | 0.9941 |
| CD3+CD8+/CD86+ \| Freq. of Parent | Control vs. CTN-067 M0 | 8 | 2.118 | 2.093 | 11 | 3.932 | 3.105 | -1.788 | 1.839 | 0.3432 | 0.7666 |

| | | | | | | | | | | |
|---|---|---|---|---|---|---|---|---|---|---|
| | Control vs. CTN-067 M3 | 8 | 2.118 | 2.093 | 11 | 3.696 | 3.273 | -1.380 | 1.715 | 0.4310 | 0.8514 |
| | Control vs. CTN-067 M6 | 8 | 2.118 | 2.093 | 11 | 3.811 | 4.450 | -1.486 | 1.710 | 0.3958 | 0.8207 |
| | CTN-067 M0 vs. CTN-067 M3 | 11 | 3.932 | 3.105 | 11 | 3.696 | 3.273 | 0.408 | 0.707 | 0.5709 | 0.9378 |
| | CTN-067 M0 vs. CTN-067 M6 | 11 | 3.932 | 3.105 | 11 | 3.811 | 4.450 | 0.302 | 0.715 | 0.6780 | 0.9741 |
| | CTN-067 M3 vs. CTN-067 M6 | 11 | 3.696 | 3.273 | 11 | 3.811 | 4.450 | -0.106 | 0.616 | 0.8652 | 0.9981 |
| CD3+CD8+ \| Geometric Mean (PE-A :: CD38) | Control vs. CTN-067 M0 | 8 | 2810.130 | 1372.280 | 11 | 6897.450 | 4457.580 | 218.760 | 1445.090 | 0.8813 | 0.9987 |
| | Control vs. CTN-067 M3 | 8 | 2810.130 | 1372.280 | 11 | 5725.090 | 3155.700 | 82.039 | 1293.070 | 0.9501 | 0.9999 |
| | Control vs. CTN-067 M6 | 8 | 2810.130 | 1372.280 | 11 | 4679.270 | 2318.470 | 1062.770 | 1287.360 | 0.4193 | 0.8417 |
| | CTN-067 M0 vs. CTN-067 M3 | 11 | 6897.450 | 4457.580 | 11 | 5725.090 | 3155.700 | -136.720 | 702.420 | 0.8477 | 0.9973 |
| | CTN-067 M0 vs. CTN-067 M6 | 11 | 6897.450 | 4457.580 | 11 | 4679.270 | 2318.470 | 844.020 | 710.640 | 0.2496 | 0.6417 |
| | CTN-067 M3 vs. CTN-067 M6 | 11 | 5725.090 | 3155.700 | 11 | 4679.270 | 2318.470 | 980.730 | 616.220 | 0.1280 | 0.4067 |
| CD3+CD8+/CD38 \| Freq. of Parent | Control vs. CTN-067 M0 | 8 | 42.813 | 23.965 | 11 | 67.555 | 13.170 | -10.405 | 9.408 | 0.2825 | 0.6903 |
| | Control vs. CTN-067 M3 | 8 | 42.813 | 23.965 | 11 | 65.855 | 14.913 | -12.324 | 9.082 | 0.1907 | 0.5399 |
| | Control vs. CTN-067 M6 | 8 | 42.813 | 23.965 | 11 | 63.300 | 13.427 | -9.949 | 9.070 | 0.2864 | 0.6956 |
| | CTN-067 M0 vs. CTN-067 M3 | 11 | 67.555 | 13.170 | 11 | 65.855 | 14.913 | -1.919 | 2.584 | 0.4666 | 0.8785 |
| | CTN-067 M0 vs. CTN-067 M6 | 11 | 67.555 | 13.170 | 11 | 63.300 | 13.427 | 0.455 | 2.616 | 0.8637 | 0.9981 |
| | CTN-067 M3 vs. CTN-067 M6 | 11 | 65.855 | 14.913 | 11 | 63.300 | 13.427 | 2.375 | 2.243 | 0.3029 | 0.7178 |

| | | | | | | | | | | | |
|---|---|---|---|---|---|---|---|---|---|---|---|
| CD3+CD8+ \| Geometric Mean (R718-A :: CCR5) | Control vs. CTN-067 M0 | 8 | 2389.500 | 954.784 | 11 | 2299.000 | 1016.320 | 884.280 | 550.710 | 0.1248 | 0.3991 |
| | Control vs. CTN-067 M3 | 8 | 2389.500 | 954.784 | 11 | 2025.270 | 772.371 | 850.700 | 493.940 | 0.1013 | 0.3401 |
| | Control vs. CTN-067 M6 | 8 | 2389.500 | 954.784 | 11 | 1922.000 | 731.088 | 938.690 | 491.810 | 0.0715 | 0.2576 |
| | CTN-067 M0 vs. CTN-067 M3 | 11 | 2299.000 | 1016.320 | 11 | 2025.270 | 772.371 | -33.580 | 264.710 | 0.9004 | 0.9992 |
| | CTN-067 M0 vs. CTN-067 M6 | 11 | 2299.000 | 1016.320 | 11 | 1922.000 | 731.088 | 54.414 | 267.810 | 0.8412 | 0.9969 |
| | CTN-067 M3 vs. CTN-067 M6 | 11 | 2025.270 | 772.371 | 11 | 1922.000 | 731.088 | 87.994 | 232.110 | 0.7088 | 0.9809 |
| CD3+CD8+/CCR5+ \| Freq. of Parent | Control vs. CTN-067 M0 | 8 | 44.600 | 12.570 | 11 | 44.700 | 14.272 | 9.081 | 8.458 | 0.2964 | 0.7092 |
| | Control vs. CTN-067 M3 | 8 | 44.600 | 12.570 | 11 | 39.836 | 14.892 | 9.781 | 7.959 | 0.2341 | 0.6168 |
| | Control vs. CTN-067 M6 | 8 | 44.600 | 12.570 | 11 | 38.236 | 16.044 | 11.174 | 7.941 | 0.1755 | 0.5103 |
| | CTN-067 M0 vs. CTN-067 M3 | 11 | 44.700 | 14.272 | 11 | 39.836 | 14.892 | 0.701 | 3.038 | 0.8201 | 0.9955 |
| | CTN-067 M0 vs. CTN-067 M6 | 11 | 44.700 | 14.272 | 11 | 38.236 | 16.044 | 2.094 | 3.076 | 0.5043 | 0.9031 |
| | CTN-067 M3 vs. CTN-067 M6 | 11 | 39.836 | 14.892 | 11 | 38.236 | 16.044 | 1.393 | 2.644 | 0.6044 | 0.9515 |
| CD3+CD8+ \| Geometric Mean (RB545-A :: CD57) | Control vs. CTN-067 M0 | 8 | 2890.750 | 1127.200 | 11 | 2280.450 | 929.055 | 566.240 | 611.670 | 0.3662 | 0.7916 |
| | Control vs. CTN-067 M3 | 8 | 2890.750 | 1127.200 | 11 | 2348.090 | 1093.870 | 383.630 | 562.300 | 0.5033 | 0.9026 |
| | Control vs. CTN-067 M6 | 8 | 2890.750 | 1127.200 | 11 | 2281.730 | 1123.390 | 444.280 | 560.470 | 0.4377 | 0.8569 |
| | CTN-067 M0 vs. CTN-067 M3 | 11 | 2280.450 | 929.055 | 11 | 2348.090 | 1093.870 | -182.610 | 258.090 | 0.4878 | 0.8929 |
| | CTN-067 M0 vs. CTN-067 M6 | 11 | 2280.450 | 929.055 | 11 | 2281.730 | 1123.390 | -121.960 | 261.200 | 0.6459 | 0.9654 |
| | CTN-067 M3 | 11 | 2348.090 | 1093.870 | 11 | 2281.730 | 1123.390 | 60.648 | 225.350 | 0.7907 | 0.9930 |

| | | | | | | | | | | |
|---|---|---|---|---|---|---|---|---|---|---|
| | vs. CTN-067 M6 | | | | | | | | | |
| CD3+CD8+/CD57 \| Freq. of Parent | Control vs. CTN-067 M0 | 8 | 16.429 | 7.155 | 11 | 19.211 | 8.817 | -6.009 | 4.458 | 0.1936 | 0.5454 |
| | Control vs. CTN-067 M3 | 8 | 16.429 | 7.155 | 11 | 17.981 | 9.117 | -4.218 | 4.304 | 0.3393 | 0.7622 |
| | Control vs. CTN-067 M6 | 8 | 16.429 | 7.155 | 11 | 16.812 | 8.821 | -3.021 | 4.298 | 0.4906 | 0.8947 |
| | CTN-067 M0 vs. CTN-067 M3 | 11 | 19.211 | 8.817 | 11 | 17.981 | 9.117 | 1.791 | 1.224 | 0.1599 | 0.4780 |
| | CTN-067 M0 vs. CTN-067 M6 | 11 | 19.211 | 8.817 | 11 | 16.812 | 8.821 | 2.987 | 1.239 | 0.0262 | 0.1089 |
| | CTN-067 M3 vs. CTN-067 M6 | 11 | 17.981 | 9.117 | 11 | 16.812 | 8.821 | 1.197 | 1.062 | 0.2739 | 0.6781 |
| CD3+CD8+ \| Geometric Mean (RB705-A :: CCR2) | Control vs. CTN-067 M0 | 8 | 1507.000 | 595.769 | 11 | 1293.730 | 866.835 | -120.140 | 375.270 | 0.7524 | 0.9883 |
| | Control vs. CTN-067 M3 | 8 | 1507.000 | 595.769 | 11 | 930.909 | 349.182 | 378.090 | 315.740 | 0.2459 | 0.6358 |
| | Control vs. CTN-067 M6 | 8 | 1507.000 | 595.769 | 11 | 1015.180 | 269.030 | 300.550 | 313.430 | 0.3496 | 0.7738 |
| | CTN-067 M0 vs. CTN-067 M3 | 11 | 1293.730 | 866.835 | 11 | 930.909 | 349.182 | 498.230 | 242.190 | 0.0537 | 0.2029 |
| | CTN-067 M0 vs. CTN-067 M6 | 11 | 1293.730 | 866.835 | 11 | 1015.180 | 269.030 | 420.690 | 244.540 | 0.1016 | 0.3411 |
| | CTN-067 M3 vs. CTN-067 M6 | 11 | 930.909 | 349.182 | 11 | 1015.180 | 269.030 | -77.541 | 217.770 | 0.7257 | 0.9840 |
| CD3+CD8+/CCR2+ \| Freq. of Parent | Control vs. CTN-067 M0 | 8 | 18.713 | 11.464 | 11 | 18.118 | 17.506 | -8.085 | 7.186 | 0.2746 | 0.6790 |
| | Control vs. CTN-067 M3 | 8 | 18.713 | 11.464 | 11 | 10.566 | 5.448 | 2.916 | 6.004 | 0.6328 | 0.9614 |
| | Control vs. CTN-067 M6 | 8 | 18.713 | 11.464 | 11 | 12.311 | 5.600 | 1.343 | 5.958 | 0.8241 | 0.9958 |
| | CTN-067 M0 vs. CTN-067 M3 | 11 | 18.118 | 17.506 | 11 | 10.566 | 5.448 | 11.001 | 5.018 | 0.0410 | 0.1612 |
| | CTN-067 M0 vs. CTN-067 M6 | 11 | 18.118 | 17.506 | 11 | 12.311 | 5.600 | 9.428 | 5.061 | 0.0780 | 0.2765 |

| | | | | | | | | | | |
|---|---|---|---|---|---|---|---|---|---|---|
| | CTN-067 M3 vs. CTN-067 M6 | 11 | 10.566 | 5.448 | 11 | 12.311 | 5.600 | -1.573 | 4.574 | 0.7347 | 0.9856 |
| CD3+CD8+ \| Geometric Mean (RB780-A :: HLA-DR) | Control vs. CTN-067 M0 | 8 | 3896.630 | 1623.320 | 11 | 8193.910 | 4063.210 | -1222.990 | 2138.180 | 0.5740 | 0.9392 |
| | Control vs. CTN-067 M3 | 8 | 3896.630 | 1623.320 | 11 | 6649.180 | 3470.760 | -1222.400 | 1895.440 | 0.5267 | 0.9160 |
| | Control vs. CTN-067 M6 | 8 | 3896.630 | 1623.320 | 11 | 6084.550 | 4188.460 | -734.540 | 1886.270 | 0.7013 | 0.9794 |
| | CTN-067 M0 vs. CTN-067 M3 | 11 | 8193.910 | 4063.210 | 11 | 6649.180 | 3470.760 | 0.590 | 1084.760 | 0.9996 | 1.0000 |
| | CTN-067 M0 vs. CTN-067 M6 | 11 | 8193.910 | 4063.210 | 11 | 6084.550 | 4188.460 | 488.460 | 1097.280 | 0.6612 | 0.9698 |
| | CTN-067 M3 vs. CTN-067 M6 | 11 | 6649.180 | 3470.760 | 11 | 6084.550 | 4188.460 | 487.870 | 953.590 | 0.6148 | 0.9553 |
| CD3+CD8+/HLA-DR \| Freq. of Parent | Control vs. CTN-067 M0 | 8 | 46.188 | 22.533 | 11 | 56.045 | 18.152 | -12.156 | 11.643 | 0.3096 | 0.7263 |
| | Control vs. CTN-067 M3 | 8 | 46.188 | 22.533 | 11 | 52.736 | 19.052 | -9.728 | 11.235 | 0.3974 | 0.8222 |
| | Control vs. CTN-067 M6 | 8 | 46.188 | 22.533 | 11 | 51.318 | 23.516 | -8.353 | 11.220 | 0.4657 | 0.8779 |
| | CTN-067 M0 vs. CTN-067 M3 | 11 | 56.045 | 18.152 | 11 | 52.736 | 19.052 | 2.428 | 3.214 | 0.4593 | 0.8732 |
| | CTN-067 M0 vs. CTN-067 M6 | 11 | 56.045 | 18.152 | 11 | 51.318 | 23.516 | 3.802 | 3.254 | 0.2571 | 0.6532 |
| | CTN-067 M3 vs. CTN-067 M6 | 11 | 52.736 | 19.052 | 11 | 51.318 | 23.516 | 1.374 | 2.790 | 0.6279 | 0.9598 |
| CD3+CD8+ \| Geometric Mean (RY586-A :: CXCR4) | Control vs. CTN-067 M0 | 8 | 4625.750 | 1222.490 | 11 | 38111.910 | 19005.350 | -30801.000 | 11617.000 | 0.0158 | 0.0688 |
| | Control vs. CTN-067 M3 | 8 | 4625.750 | 1222.490 | 11 | 20885.550 | 24613.780 | -12624.000 | 9726.100 | 0.2098 | 0.5752 |
| | Control vs. CTN-067 M6 | 8 | 4625.750 | 1222.490 | 11 | 14776.270 | 16753.630 | -6467.890 | 9652.320 | 0.5109 | 0.9071 |
| | CTN-067 M0 vs. CTN-067 M3 | 11 | 38111.910 | 19005.350 | 11 | 20885.550 | 24613.780 | 18176.000 | 7812.300 | 0.0312 | 0.1270 |
| | CTN-067 M0 | 11 | 38111.910 | 19005.350 | 11 | 14776.270 | 16753.630 | 24333.000 | 7884.020 | 0.0061 | 0.0284 |

| | | | | | | | | | | | |
|---|---|---|---|---|---|---|---|---|---|---|---|
| | vs. CTN-067 M6 | | | | | | | | | | |
| | CTN-067 M3 vs. CTN-067 M6 | 11 | 20885.550 | 24613.780 | 11 | 14776.270 | 16753.630 | 6156.490 | 7071.930 | 0.3949 | 0.8198 |
| CD3+CD8+/CXCR4+ \| Freq. of Parent | Control vs. CTN-067 M0 | 8 | 53.988 | 19.498 | 11 | 87.855 | 17.991 | -30.480 | 14.937 | 0.0554 | 0.2085 |
| | Control vs. CTN-067 M3 | 8 | 53.988 | 19.498 | 11 | 57.215 | 34.418 | 4.395 | 12.480 | 0.7286 | 0.9845 |
| | Control vs. CTN-067 M6 | 8 | 53.988 | 19.498 | 11 | 56.245 | 33.662 | 5.575 | 12.385 | 0.6577 | 0.9688 |
| | CTN-067 M0 vs. CTN-067 M3 | 11 | 87.855 | 17.991 | 11 | 57.215 | 34.418 | 34.875 | 10.600 | 0.0039 | 0.0185 |
| | CTN-067 M0 vs. CTN-067 M6 | 11 | 87.855 | 17.991 | 11 | 56.245 | 33.662 | 36.055 | 10.688 | 0.0032 | 0.0154 |
| | CTN-067 M3 vs. CTN-067 M6 | 11 | 57.215 | 34.418 | 11 | 56.245 | 33.662 | 1.180 | 9.694 | 0.9044 | 0.9993 |
| CD3+CD8+ \| Geometric Mean (RY610-A :: CD69) | Control vs. CTN-067 M0 | 8 | 659.000 | 337.221 | 11 | 1133.090 | 415.586 | -19.515 | 327.380 | 0.9531 | 0.9999 |
| | Control vs. CTN-067 M3 | 8 | 659.000 | 337.221 | 11 | 1216.000 | 611.467 | -249.310 | 276.140 | 0.3779 | 0.8035 |
| | Control vs. CTN-067 M6 | 8 | 659.000 | 337.221 | 11 | 1273.910 | 647.588 | -314.520 | 274.150 | 0.2655 | 0.6658 |
| | CTN-067 M0 vs. CTN-067 M3 | 11 | 1133.090 | 415.586 | 11 | 1216.000 | 611.467 | -229.800 | 207.970 | 0.2830 | 0.6909 |
| | CTN-067 M0 vs. CTN-067 M6 | 11 | 1133.090 | 415.586 | 11 | 1273.910 | 647.588 | -295.010 | 210.030 | 0.1763 | 0.5118 |
| | CTN-067 M3 vs. CTN-067 M6 | 11 | 1216.000 | 611.467 | 11 | 1273.910 | 647.588 | -65.212 | 186.560 | 0.7305 | 0.9849 |
| CD3+CD8+/CD69+ \| Freq. of Parent | Control vs. CTN-067 M0 | 8 | 6.074 | 6.321 | 11 | 8.395 | 4.863 | 2.139 | 3.395 | 0.5362 | 0.9211 |
| | Control vs. CTN-067 M3 | 8 | 6.074 | 6.321 | 11 | 8.691 | 6.947 | 0.589 | 3.032 | 0.8482 | 0.9973 |
| | Control vs. CTN-067 M6 | 8 | 6.074 | 6.321 | 11 | 7.385 | 5.680 | 1.832 | 3.019 | 0.5512 | 0.9287 |
| | CTN-067 M0 vs. CTN-067 M3 | 11 | 8.395 | 4.863 | 11 | 8.691 | 6.947 | -1.550 | 1.664 | 0.3631 | 0.7883 |

| | | | | | | | | | | |
|---|---|---|---|---|---|---|---|---|---|---|
| | CTN-067 M0 vs. CTN-067 M6 | 11 | 8.395 | 4.863 | 11 | 7.385 | 5.680 | -0.307 | 1.683 | 0.8571 | 0.9978 |
| | CTN-067 M3 vs. CTN-067 M6 | 11 | 8.691 | 6.947 | 11 | 7.385 | 5.680 | 1.243 | 1.460 | 0.4052 | 0.8293 |
| CD3+CD8+ \| Geometric Mean (RY703-A :: PD-1) | Control vs. CTN-067 M0 | 8 | 2097.750 | 578.789 | 11 | 3244.270 | 2129.490 | 221.200 | 725.810 | 0.7639 | 0.9899 |
| | Control vs. CTN-067 M3 | 8 | 2097.750 | 578.789 | 11 | 2629.360 | 939.905 | 449.590 | 662.260 | 0.5054 | 0.9038 |
| | Control vs. CTN-067 M6 | 8 | 2097.750 | 578.789 | 11 | 2582.640 | 1212.120 | 477.100 | 659.900 | 0.4785 | 0.8867 |
| | CTN-067 M0 vs. CTN-067 M3 | 11 | 3244.270 | 2129.490 | 11 | 2629.360 | 939.905 | 228.390 | 319.600 | 0.4835 | 0.8901 |
| | CTN-067 M0 vs. CTN-067 M6 | 11 | 3244.270 | 2129.490 | 11 | 2582.640 | 1212.120 | 255.900 | 323.420 | 0.4386 | 0.8575 |
| | CTN-067 M3 vs. CTN-067 M6 | 11 | 2629.360 | 939.905 | 11 | 2582.640 | 1212.120 | 27.511 | 279.380 | 0.9226 | 0.9996 |
| CD3+CD8+/PD-1+ \| Freq. of Parent | Control vs. CTN-067 M0 | 8 | 33.313 | 12.467 | 11 | 42.045 | 16.034 | 1.840 | 7.706 | 0.8139 | 0.9950 |
| | Control vs. CTN-067 M3 | 8 | 33.313 | 12.467 | 11 | 35.327 | 13.107 | 5.243 | 7.565 | 0.4967 | 0.8985 |
| | Control vs. CTN-067 M6 | 8 | 33.313 | 12.467 | 11 | 34.682 | 15.520 | 5.724 | 7.560 | 0.4583 | 0.8725 |
| | CTN-067 M0 vs. CTN-067 M3 | 11 | 42.045 | 16.034 | 11 | 35.327 | 13.107 | 3.404 | 1.536 | 0.0391 | 0.1547 |
| | CTN-067 M0 vs. CTN-067 M6 | 11 | 42.045 | 16.034 | 11 | 34.682 | 15.520 | 3.884 | 1.555 | 0.0219 | 0.0925 |
| | CTN-067 M3 vs. CTN-067 M6 | 11 | 35.327 | 13.107 | 11 | 34.682 | 15.520 | 0.481 | 1.331 | 0.7220 | 0.9834 |
| CD3+CD8+/CCR7- \| Freq. of Parent | Control vs. CTN-067 M0 | 8 | 58.350 | 16.704 | 11 | 60.600 | 14.726 | -0.198 | 8.697 | 0.9820 | 1.0000 |
| | Control vs. CTN-067 M3 | 8 | 58.350 | 16.704 | 11 | 57.036 | 16.575 | 0.375 | 8.415 | 0.9649 | 1.0000 |
| | Control vs. CTN-067 M6 | 8 | 58.350 | 16.704 | 11 | 53.309 | 16.489 | 3.954 | 8.405 | 0.6434 | 0.9647 |
| | CTN-067 M0 | 11 | 60.600 | 14.726 | 11 | 57.036 | 16.575 | 0.574 | 2.312 | 0.8066 | 0.9944 |

| | | | | | | | | | | | |
|---|---|---|---|---|---|---|---|---|---|---|---|
| | vs. CTN-067 M3 | | | | | | | | | | |
| | CTN-067 M0 vs. CTN-067 M6 | 11 | 60.600 | 14.726 | 11 | 53.309 | 16.489 | 4.153 | 2.341 | 0.0921 | 0.3157 |
| | CTN-067 M3 vs. CTN-067 M6 | 11 | 57.036 | 16.575 | 11 | 53.309 | 16.489 | 3.579 | 2.006 | 0.0904 | 0.3112 |
| CD3+CD8+/CD28- \| Freq. of Parent | Control vs. CTN-067 M0 | 8 | 41.238 | 21.363 | 11 | 46.918 | 19.146 | -1.745 | 11.509 | 0.8811 | 0.9987 |
| | Control vs. CTN-067 M3 | 8 | 41.238 | 21.363 | 11 | 39.918 | 16.727 | 1.807 | 10.378 | 0.8636 | 0.9981 |
| | Control vs. CTN-067 M6 | 8 | 41.238 | 21.363 | 11 | 36.864 | 18.985 | 4.690 | 10.336 | 0.6551 | 0.9681 |
| | CTN-067 M0 vs. CTN-067 M3 | 11 | 46.918 | 19.146 | 11 | 39.918 | 16.727 | 3.552 | 5.388 | 0.5177 | 0.9110 |
| | CTN-067 M0 vs. CTN-067 M6 | 11 | 46.918 | 19.146 | 11 | 36.864 | 18.985 | 6.435 | 5.452 | 0.2524 | 0.6461 |
| | CTN-067 M3 vs. CTN-067 M6 | 11 | 39.918 | 16.727 | 11 | 36.864 | 18.985 | 2.883 | 4.720 | 0.5485 | 0.9274 |
| CD3+CD8+/CD38- \| Freq. of Parent | Control vs. CTN-067 M0 | 8 | 57.188 | 23.965 | 11 | 32.445 | 13.170 | 10.405 | 9.408 | 0.2825 | 0.6903 |
| | Control vs. CTN-067 M3 | 8 | 57.188 | 23.965 | 11 | 34.145 | 14.913 | 12.324 | 9.082 | 0.1907 | 0.5399 |
| | Control vs. CTN-067 M6 | 8 | 57.188 | 23.965 | 11 | 36.700 | 13.427 | 9.949 | 9.070 | 0.2864 | 0.6956 |
| | CTN-067 M0 vs. CTN-067 M3 | 11 | 32.445 | 13.170 | 11 | 34.145 | 14.913 | 1.919 | 2.584 | 0.4666 | 0.8785 |
| | CTN-067 M0 vs. CTN-067 M6 | 11 | 32.445 | 13.170 | 11 | 36.700 | 13.427 | -0.455 | 2.616 | 0.8637 | 0.9981 |
| | CTN-067 M3 vs. CTN-067 M6 | 11 | 34.145 | 14.913 | 11 | 36.700 | 13.427 | -2.375 | 2.243 | 0.3029 | 0.7178 |
| CD3+CD8+/CD45RA- \| Freq. of Parent | Control vs. CTN-067 M0 | 8 | 14.511 | 2.935 | 11 | 15.662 | 5.055 | -4.856 | 2.624 | 0.0799 | 0.2817 |
| | Control vs. CTN-067 M3 | 8 | 14.511 | 2.935 | 11 | 15.727 | 5.165 | -3.469 | 2.464 | 0.1752 | 0.5097 |
| | Control vs. CTN-067 M6 | 8 | 14.511 | 2.935 | 11 | 15.673 | 6.187 | -3.342 | 2.458 | 0.1898 | 0.5382 |

| | | | | | | | | | | |
|---|---|---|---|---|---|---|---|---|---|---|
| | CTN-067 M0 vs. CTN-067 M3 | 11 | 15.662 | 5.055 | 11 | 15.727 | 5.165 | 1.387 | 0.960 | 0.1649 | 0.4886 |
| | CTN-067 M0 vs. CTN-067 M6 | 11 | 15.662 | 5.055 | 11 | 15.673 | 6.187 | 1.513 | 0.972 | 0.1359 | 0.4251 |
| | CTN-067 M3 vs. CTN-067 M6 | 11 | 15.727 | 5.165 | 11 | 15.673 | 6.187 | 0.127 | 0.836 | 0.8811 | 0.9987 |
| CD3+CD8+/CD57- \| Freq. of Parent | Control vs. CTN-067 M0 | 8 | 83.575 | 7.159 | 11 | 80.791 | 8.819 | 6.008 | 4.460 | 0.1939 | 0.5460 |
| | Control vs. CTN-067 M3 | 8 | 83.575 | 7.159 | 11 | 82.018 | 9.117 | 4.220 | 4.306 | 0.3394 | 0.7623 |
| | Control vs. CTN-067 M6 | 8 | 83.575 | 7.159 | 11 | 83.191 | 8.825 | 3.019 | 4.300 | 0.4912 | 0.8950 |
| | CTN-067 M0 vs. CTN-067 M3 | 11 | 80.791 | 8.819 | 11 | 82.018 | 9.117 | -1.788 | 1.225 | 0.1606 | 0.4797 |
| | CTN-067 M0 vs. CTN-067 M6 | 11 | 80.791 | 8.819 | 11 | 83.191 | 8.825 | -2.989 | 1.240 | 0.0262 | 0.1090 |
| | CTN-067 M3 vs. CTN-067 M6 | 11 | 82.018 | 9.117 | 11 | 83.191 | 8.825 | -1.201 | 1.063 | 0.2728 | 0.6764 |
| CD3+CD8+/HLA-DR- \| Freq. of Parent | Control vs. CTN-067 M0 | 8 | 53.813 | 22.533 | 11 | 43.955 | 18.152 | 12.156 | 11.643 | 0.3096 | 0.7263 |
| | Control vs. CTN-067 M3 | 8 | 53.813 | 22.533 | 11 | 47.264 | 19.052 | 9.728 | 11.235 | 0.3974 | 0.8222 |
| | Control vs. CTN-067 M6 | 8 | 53.813 | 22.533 | 11 | 48.682 | 23.516 | 8.353 | 11.220 | 0.4657 | 0.8779 |
| | CTN-067 M0 vs. CTN-067 M3 | 11 | 43.955 | 18.152 | 11 | 47.264 | 19.052 | -2.428 | 3.214 | 0.4593 | 0.8732 |
| | CTN-067 M0 vs. CTN-067 M6 | 11 | 43.955 | 18.152 | 11 | 48.682 | 23.516 | -3.802 | 3.254 | 0.2571 | 0.6532 |
| | CTN-067 M3 vs. CTN-067 M6 | 11 | 47.264 | 19.052 | 11 | 48.682 | 23.516 | -1.374 | 2.790 | 0.6279 | 0.9598 |
| CD3+CD8+/CCR7+CD45RA+ \| Freq. of Parent | Control vs. CTN-067 M0 | 8 | 37.075 | 14.939 | 11 | 36.173 | 14.138 | 0.822 | 8.213 | 0.9213 | 0.9996 |
| | Control vs. CTN-067 M3 | 8 | 37.075 | 14.939 | 11 | 39.682 | 16.186 | -0.677 | 7.935 | 0.9329 | 0.9998 |

| | | | | | | | | | | | |
|---|---|---|---|---|---|---|---|---|---|---|---|
| | Control vs. CTN-067 M6 | 8 | 37.075 | 14.939 | 11 | 42.464 | 16.163 | -3.358 | 7.925 | 0.6765 | 0.9737 |
| | CTN-067 M0 vs. CTN-067 M3 | 11 | 36.173 | 14.138 | 11 | 39.682 | 16.186 | -1.499 | 2.228 | 0.5091 | 0.9060 |
| | CTN-067 M0 vs. CTN-067 M6 | 11 | 36.173 | 14.138 | 11 | 42.464 | 16.163 | -4.181 | 2.256 | 0.0794 | 0.2804 |
| | CTN-067 M3 vs. CTN-067 M6 | 11 | 39.682 | 16.186 | 11 | 42.464 | 16.163 | -2.682 | 1.933 | 0.1814 | 0.5221 |
| CD3+CD8+/CCR7+CD45RA- \| Freq. of Parent | Control vs. CTN-067 M0 | 8 | 4.566 | 2.583 | 11 | 3.241 | 1.435 | -0.612 | 1.252 | 0.6308 | 0.9607 |
| | Control vs. CTN-067 M3 | 8 | 4.566 | 2.583 | 11 | 3.285 | 2.015 | 0.313 | 1.173 | 0.7922 | 0.9931 |
| | Control vs. CTN-067 M6 | 8 | 4.566 | 2.583 | 11 | 4.215 | 2.913 | -0.570 | 1.170 | 0.6319 | 0.9611 |
| | CTN-067 M0 vs. CTN-067 M3 | 11 | 3.241 | 1.435 | 11 | 3.285 | 2.015 | 0.925 | 0.467 | 0.0621 | 0.2294 |
| | CTN-067 M0 vs. CTN-067 M6 | 11 | 3.241 | 1.435 | 11 | 4.215 | 2.913 | 0.042 | 0.472 | 0.9301 | 0.9997 |
| | CTN-067 M3 vs. CTN-067 M6 | 11 | 3.285 | 2.015 | 11 | 4.215 | 2.913 | -0.883 | 0.406 | 0.0427 | 0.1669 |
| CD3+CD8+/CCR7-CD45RA+ \| Freq. of Parent | Control vs. CTN-067 M0 | 8 | 48.400 | 16.385 | 11 | 48.173 | 10.879 | 3.851 | 7.903 | 0.6317 | 0.9610 |
| | Control vs. CTN-067 M3 | 8 | 48.400 | 16.385 | 11 | 44.609 | 14.365 | 4.025 | 7.555 | 0.6004 | 0.9500 |
| | Control vs. CTN-067 M6 | 8 | 48.400 | 16.385 | 11 | 41.845 | 14.110 | 6.620 | 7.543 | 0.3911 | 0.8163 |
| | CTN-067 M0 vs. CTN-067 M3 | 11 | 48.173 | 10.879 | 11 | 44.609 | 14.365 | 0.174 | 2.448 | 0.9441 | 0.9999 |
| | CTN-067 M0 vs. CTN-067 M6 | 11 | 48.173 | 10.879 | 11 | 41.845 | 14.110 | 2.769 | 2.478 | 0.2777 | 0.6835 |
| | CTN-067 M3 vs. CTN-067 M6 | 11 | 44.609 | 14.365 | 11 | 41.845 | 14.110 | 2.595 | 2.127 | 0.2373 | 0.6220 |
| CD3+CD8+/CCR7-CD45RA- \| Freq. of Parent | Control vs. CTN-067 M0 | 8 | 9.936 | 3.263 | 11 | 12.418 | 5.304 | -4.267 | 2.384 | 0.0894 | 0.3083 |

| | | | | | | | | | | | |
|---|---|---|---|---|---|---|---|---|---|---|---|
| | Control vs. CTN-067 M3 | 8 | 9.936 | 3.263 | 11 | 12.449 | 4.445 | -3.810 | 2.261 | 0.1083 | 0.3582 |
| | Control vs. CTN-067 M6 | 8 | 9.936 | 3.263 | 11 | 11.454 | 4.368 | -2.790 | 2.256 | 0.2313 | 0.6122 |
| | CTN-067 M0 vs. CTN-067 M3 | 11 | 12.418 | 5.304 | 11 | 12.449 | 4.445 | 0.458 | 0.800 | 0.5741 | 0.9392 |
| | CTN-067 M0 vs. CTN-067 M6 | 11 | 12.418 | 5.304 | 11 | 11.454 | 4.368 | 1.477 | 0.810 | 0.0839 | 0.2932 |
| | CTN-067 M3 vs. CTN-067 M6 | 11 | 12.449 | 4.445 | 11 | 11.454 | 4.368 | 1.020 | 0.696 | 0.1591 | 0.4764 |
| CD3+CD8+/CD28+CD57+ \| Freq. of Parent | Control vs. CTN-067 M0 | 8 | 5.021 | 1.710 | 11 | 5.280 | 2.673 | -0.998 | 1.229 | 0.4266 | 0.8479 |
| | Control vs. CTN-067 M3 | 8 | 5.021 | 1.710 | 11 | 5.000 | 2.399 | -0.254 | 1.154 | 0.8285 | 0.9961 |
| | Control vs. CTN-067 M6 | 8 | 5.021 | 1.710 | 11 | 5.161 | 2.167 | -0.391 | 1.151 | 0.7377 | 0.9861 |
| | CTN-067 M0 vs. CTN-067 M3 | 11 | 5.280 | 2.673 | 11 | 5.000 | 2.399 | 0.745 | 0.449 | 0.1133 | 0.3709 |
| | CTN-067 M0 vs. CTN-067 M6 | 11 | 5.280 | 2.673 | 11 | 5.161 | 2.167 | 0.607 | 0.454 | 0.1972 | 0.5522 |
| | CTN-067 M3 vs. CTN-067 M6 | 11 | 5.000 | 2.399 | 11 | 5.161 | 2.167 | -0.138 | 0.391 | 0.7281 | 0.9845 |
| CD3+CD8+/CD28+CD57- \| Freq. of Parent | Control vs. CTN-067 M0 | 8 | 53.725 | 20.671 | 11 | 47.818 | 18.512 | 2.608 | 11.320 | 0.8203 | 0.9955 |
| | Control vs. CTN-067 M3 | 8 | 53.725 | 20.671 | 11 | 55.100 | 17.130 | -1.646 | 10.295 | 0.8746 | 0.9985 |
| | Control vs. CTN-067 M6 | 8 | 53.725 | 20.671 | 11 | 57.973 | 19.153 | -4.369 | 10.257 | 0.6749 | 0.9733 |
| | CTN-067 M0 vs. CTN-067 M3 | 11 | 47.818 | 18.512 | 11 | 55.100 | 17.130 | -4.254 | 5.073 | 0.4122 | 0.8355 |
| | CTN-067 M0 vs. CTN-067 M6 | 11 | 47.818 | 18.512 | 11 | 57.973 | 19.153 | -6.976 | 5.134 | 0.1901 | 0.5388 |
| | CTN-067 M3 vs. CTN-067 M6 | 11 | 55.100 | 17.130 | 11 | 57.973 | 19.153 | -2.722 | 4.437 | 0.5468 | 0.9265 |

| | | | | | | | | | | | |
|---|---|---|---|---|---|---|---|---|---|---|---|
| CD3+CD8+/CD28-CD57+ \| Freq. of Parent | Control vs. CTN-067 M0 | 8 | 11.409 | 7.440 | 11 | 13.929 | 7.139 | -5.066 | 4.059 | 0.2271 | 0.6053 |
| | Control vs. CTN-067 M3 | 8 | 11.409 | 7.440 | 11 | 12.979 | 7.866 | -3.994 | 3.863 | 0.3142 | 0.7321 |
| | Control vs. CTN-067 M6 | 8 | 11.409 | 7.440 | 11 | 11.645 | 7.777 | -2.654 | 3.856 | 0.4997 | 0.9004 |
| | CTN-067 M0 vs. CTN-067 M3 | 11 | 13.929 | 7.139 | 11 | 12.979 | 7.866 | 1.072 | 1.315 | 0.4253 | 0.8467 |
| | CTN-067 M0 vs. CTN-067 M6 | 11 | 13.929 | 7.139 | 11 | 11.645 | 7.777 | 2.412 | 1.332 | 0.0859 | 0.2987 |
| | CTN-067 M3 vs. CTN-067 M6 | 11 | 12.979 | 7.866 | 11 | 11.645 | 7.777 | 1.341 | 1.143 | 0.2555 | 0.6507 |
| CD3+CD8+/CD28-CD57- \| Freq. of Parent | Control vs. CTN-067 M0 | 8 | 29.838 | 16.235 | 11 | 32.964 | 16.536 | 3.588 | 8.751 | 0.6864 | 0.9761 |
| | Control vs. CTN-067 M3 | 8 | 29.838 | 16.235 | 11 | 26.936 | 9.899 | 5.946 | 7.704 | 0.4497 | 0.8661 |
| | Control vs. CTN-067 M6 | 8 | 29.838 | 16.235 | 11 | 25.207 | 12.510 | 7.493 | 7.664 | 0.3405 | 0.7636 |
| | CTN-067 M0 vs. CTN-067 M3 | 11 | 32.964 | 16.536 | 11 | 26.936 | 9.899 | 2.358 | 4.577 | 0.6124 | 0.9544 |
| | CTN-067 M0 vs. CTN-067 M6 | 11 | 32.964 | 16.536 | 11 | 25.207 | 12.510 | 3.905 | 4.630 | 0.4095 | 0.8331 |
| | CTN-067 M3 vs. CTN-067 M6 | 11 | 26.936 | 9.899 | 11 | 25.207 | 12.510 | 1.547 | 4.031 | 0.7054 | 0.9802 |
| CD3+CD8+/CD38+HLA-DR+ \| Freq. of Parent | Control vs. CTN-067 M0 | 8 | 26.736 | 21.119 | 11 | 43.709 | 14.651 | -9.617 | 10.118 | 0.3538 | 0.7784 |
| | Control vs. CTN-067 M3 | 8 | 26.736 | 21.119 | 11 | 37.809 | 13.181 | -6.698 | 9.642 | 0.4957 | 0.8979 |
| | Control vs. CTN-067 M6 | 8 | 26.736 | 21.119 | 11 | 35.673 | 17.479 | -4.710 | 9.625 | 0.6302 | 0.9605 |
| | CTN-067 M0 vs. CTN-067 M3 | 11 | 43.709 | 14.651 | 11 | 37.809 | 13.181 | 2.919 | 3.241 | 0.3790 | 0.8045 |
| | CTN-067 M0 vs. CTN-067 M6 | 11 | 43.709 | 14.651 | 11 | 35.673 | 17.479 | 4.907 | 3.281 | 0.1511 | 0.4593 |
| | CTN-067 M3 | 11 | 37.809 | 13.181 | 11 | 35.673 | 17.479 | 1.988 | 2.817 | 0.4888 | 0.8935 |

| | | | | | | | | | | | |
|---|---|---|---|---|---|---|---|---|---|---|---|
| | vs. CTN-067 M6 | | | | | | | | | | |
| CD3+CD8+/CD38+HLA-DR- \| Freq. of Parent | Control vs. CTN-067 M0 | 8 | 16.075 | 6.972 | 11 | 23.853 | 14.692 | -0.772 | 7.194 | 0.9157 | 0.9995 |
| | Control vs. CTN-067 M3 | 8 | 16.075 | 6.972 | 11 | 28.073 | 17.994 | -5.645 | 6.867 | 0.4212 | 0.8433 |
| | Control vs. CTN-067 M6 | 8 | 16.075 | 6.972 | 11 | 27.591 | 18.124 | -5.196 | 6.855 | 0.4578 | 0.8722 |
| | CTN-067 M0 vs. CTN-067 M3 | 11 | 23.853 | 14.692 | 11 | 28.073 | 17.994 | -4.873 | 2.267 | 0.0447 | 0.1736 |
| | CTN-067 M0 vs. CTN-067 M6 | 11 | 23.853 | 14.692 | 11 | 27.591 | 18.124 | -4.424 | 2.295 | 0.0690 | 0.2502 |
| | CTN-067 M3 vs. CTN-067 M6 | 11 | 28.073 | 17.994 | 11 | 27.591 | 18.124 | 0.449 | 1.970 | 0.8220 | 0.9957 |
| CD3+CD8+/CD38-HLA-DR+ \| Freq. of Parent | Control vs. CTN-067 M0 | 8 | 19.473 | 5.687 | 11 | 12.343 | 9.688 | -2.825 | 3.924 | 0.4804 | 0.8880 |
| | Control vs. CTN-067 M3 | 8 | 19.473 | 5.687 | 11 | 14.955 | 11.247 | -3.214 | 3.720 | 0.3984 | 0.8231 |
| | Control vs. CTN-067 M6 | 8 | 19.473 | 5.687 | 11 | 15.615 | 10.296 | -3.763 | 3.713 | 0.3236 | 0.7437 |
| | CTN-067 M0 vs. CTN-067 M3 | 11 | 12.343 | 9.688 | 11 | 14.955 | 11.247 | -0.389 | 1.323 | 0.7719 | 0.9909 |
| | CTN-067 M0 vs. CTN-067 M6 | 11 | 12.343 | 9.688 | 11 | 15.615 | 10.296 | -0.938 | 1.339 | 0.4923 | 0.8958 |
| | CTN-067 M3 vs. CTN-067 M6 | 11 | 14.955 | 11.247 | 11 | 15.615 | 10.296 | -0.549 | 1.150 | 0.6389 | 0.9633 |
| CD3+CD8+/CD38-HLA-DR- \| Freq. of Parent | Control vs. CTN-067 M0 | 8 | 37.741 | 26.084 | 11 | 20.113 | 10.975 | 13.148 | 10.036 | 0.2058 | 0.5679 |
| | Control vs. CTN-067 M3 | 8 | 37.741 | 26.084 | 11 | 19.207 | 9.881 | 15.491 | 9.832 | 0.1316 | 0.4152 |
| | Control vs. CTN-067 M6 | 8 | 37.741 | 26.084 | 11 | 21.100 | 11.117 | 13.670 | 9.825 | 0.1802 | 0.5196 |
| | CTN-067 M0 vs. CTN-067 M3 | 11 | 20.113 | 10.975 | 11 | 19.207 | 9.881 | 2.343 | 2.109 | 0.2806 | 0.6875 |
| | CTN-067 M0 vs. CTN-067 M6 | 11 | 20.113 | 10.975 | 11 | 21.100 | 11.117 | 0.522 | 2.136 | 0.8097 | 0.9947 |

| | | | | | | | | | | | |
|---|---|---|---|---|---|---|---|---|---|---|---|
| | CTN-067 M3 vs. CTN-067 M6 | 1 1 | 19.207 | 9.881 | 1 1 | 21.100 | 11.117 | -1.821 | 1.829 | 0.33 17 | 0.75 34 |
