## Supplemental Table 13 for "Chronic opioid-associated immune dysregulation among people living with HIV"

**Supplementary Table 13.** CD56+ NK cell Flow Data

| Marker | Comparison (A vs B) | Group A | | | Group B | | | Viral Load adjusted Group A – Group B | | | |
|---|---|---|---|---|---|---|---|---|---|---|---|
| | | n | Mean | SD | N | Mean | SD | Estimate | SE | p | adj p |
| CD19- \| Geometric Mean (FITC-A :: CD56) | Control vs. CTN-067 M0 | 8 | 1328.500 | 1350.860 | 11 | 1275.180 | 564.668 | 131.920 | 526.880 | 0.8050 | 0.9943 |
| | Control vs. CTN-067 M3 | 8 | 1328.500 | 1350.860 | 11 | 1183.550 | 574.771 | 207.900 | 510.610 | 0.6884 | 0.9765 |
| | Control vs. CTN-067 M6 | 8 | 1328.500 | 1350.860 | 11 | 1249.820 | 532.600 | 140.850 | 510.020 | 0.7854 | 0.9924 |
| | CTN-067 M0 vs. CTN-067 M3 | 11 | 1275.180 | 564.668 | 11 | 1183.550 | 574.771 | 75.980 | 136.590 | 0.5845 | 0.9437 |
| | CTN-067 M0 vs. CTN-067 M6 | 11 | 1275.180 | 564.668 | 11 | 1249.820 | 532.600 | 8.929 | 138.300 | 0.9492 | 0.9999 |
| | CTN-067 M3 vs. CTN-067 M6 | 11 | 1183.550 | 574.771 | 11 | 1249.820 | 532.600 | -67.051 | 118.510 | 0.5782 | 0.9410 |
| CD56+ \| Freq. of Parent | Control vs. CTN-067 M0 | 8 | 26.310 | 22.461 | 11 | 27.427 | 11.837 | -1.143 | 9.224 | 0.9027 | 0.9993 |
| | Control vs. CTN-067 M3 | 8 | 26.310 | 22.461 | 11 | 24.027 | 12.230 | 2.725 | 8.939 | 0.7638 | 0.9898 |
| | Control vs. CTN-067 M6 | 8 | 26.310 | 22.461 | 11 | 25.743 | 10.614 | 1.033 | 8.929 | 0.9091 | 0.9994 |
| | CTN-067 M0 vs. CTN-067 M3 | 11 | 27.427 | 11.837 | 11 | 24.027 | 12.230 | 3.868 | 2.391 | 0.1222 | 0.3928 |
| | CTN-067 M0 vs. CTN-067 M6 | 11 | 27.427 | 11.837 | 11 | 25.743 | 10.614 | 2.176 | 2.421 | 0.3801 | 0.8056 |
| | CTN-067 M3 vs. CTN-067 M6 | 11 | 24.027 | 12.230 | 11 | 25.743 | 10.614 | -1.692 | 2.075 | 0.4248 | 0.8463 |
| CD56+ \| Geometric Mean (FITC-A :: CD56) | Control vs. CTN-067 M0 | 8 | 8186.750 | 543.368 | 11 | 9130.180 | 3257.680 | -354.110 | 1168586.000 | 0.9998 | 1.0000 |
| | Control vs. CTN-067 M3 | 8 | 8186.750 | 543.368 | 11 | 10029.550 | 2715.130 | -1118.110 | 1168586.000 | 0.9992 | 1.0000 |
| | Control vs. CTN-067 M6 | 8 | 8186.750 | 543.368 | 11 | 10111.820 | 3561.750 | -1193.650 | 1168586.000 | 0.9992 | 1.0000 |
| | CTN-067 M0 vs. CTN-067 M3 | 11 | 9130.180 | 3257.680 | 11 | 10029.550 | 2715.130 | -764.000 | 411.060 | 0.0786 | 0.2783 |
| | CTN-067 M0 vs. | 11 | 9130.180 | 3257.680 | 11 | 10111.820 | 3561.750 | -839.550 | 416.300 | 0.0581 | 0.2168 |

| | | | | | | | | | | |
|---|---|---|---|---|---|---|---|---|---|---|
| | CTN-067 M6 | | | | | | | | | |
| | CTN-067 M3 vs. CTN-067 M6 | 11 | 10029.550 | 2715.130 | 11 | 10111.820 | 3561.750 | -75.543 | 355.720 | 0.8341 | 0.9965 |
| CD56+ \| Geometric Mean (APC-Fire 750-A :: CD36) | Control vs. CTN-067 M0 | 8 | 1019.630 | 290.243 | 11 | 1083.820 | 502.395 | 95.610 | 243.770 | 0.6993 | 0.9789 |
| | Control vs. CTN-067 M3 | 8 | 1019.630 | 290.243 | 11 | 1093.180 | 409.766 | 13.016 | 216.670 | 0.9527 | 0.9999 |
| | Control vs. CTN-067 M6 | 8 | 1019.630 | 290.243 | 11 | 1200.450 | 377.722 | -97.898 | 215.650 | 0.6550 | 0.9680 |
| | CTN-067 M0 vs. CTN-067 M3 | 11 | 1083.820 | 502.395 | 11 | 1093.180 | 409.766 | -82.595 | 122.190 | 0.5072 | 0.9049 |
| | CTN-067 M0 vs. CTN-067 M6 | 11 | 1083.820 | 502.395 | 11 | 1200.450 | 377.722 | -193.510 | 123.600 | 0.1340 | 0.4206 |
| | CTN-067 M3 vs. CTN-067 M6 | 11 | 1093.180 | 409.766 | 11 | 1200.450 | 377.722 | -110.910 | 107.350 | 0.3145 | 0.7325 |
| CD56+/CD36+ \| Freq. of Parent | Control vs. CTN-067 M0 | 8 | 13.616 | 6.094 | 11 | 15.404 | 7.345 | 0.062 | 4.372 | 0.9888 | 1.0000 |
| | Control vs. CTN-067 M3 | 8 | 13.616 | 6.094 | 11 | 17.256 | 8.333 | -2.794 | 3.986 | 0.4918 | 0.8954 |
| | Control vs. CTN-067 M6 | 8 | 13.616 | 6.094 | 11 | 18.145 | 7.801 | -3.733 | 3.972 | 0.3590 | 0.7840 |
| | CTN-067 M0 vs. CTN-067 M3 | 11 | 15.404 | 7.345 | 11 | 17.256 | 8.333 | -2.856 | 1.935 | 0.1562 | 0.4703 |
| | CTN-067 M0 vs. CTN-067 M6 | 11 | 15.404 | 7.345 | 11 | 18.145 | 7.801 | -3.795 | 1.958 | 0.0676 | 0.2459 |
| | CTN-067 M3 vs. CTN-067 M6 | 11 | 17.256 | 8.333 | 11 | 18.145 | 7.801 | -0.939 | 1.692 | 0.5853 | 0.9440 |
| CD56+ \| Geometric Mean (Alexa Fluor 647-A :: GLUT1) | Control vs. CTN-067 M0 | 8 | 767.500 | 153.082 | 11 | 665.727 | 174.534 | 138.450 | 119.580 | 0.2613 | 0.6595 |
| | Control vs. CTN-067 M3 | 8 | 767.500 | 153.082 | 11 | 714.182 | 236.439 | 52.607 | 110.650 | 0.6399 | 0.9636 |
| | Control vs. CTN-067 M6 | 8 | 767.500 | 153.082 | 11 | 673.182 | 274.032 | 91.748 | 110.320 | 0.4159 | 0.8388 |
| | CTN-067 M0 vs. CTN-067 M3 | 11 | 665.727 | 174.534 | 11 | 714.182 | 236.439 | -85.842 | 48.457 | 0.0925 | 0.3169 |
| | CTN-067 M0 vs. CTN-067 M6 | 11 | 665.727 | 174.534 | 11 | 673.182 | 274.032 | -46.701 | 49.046 | 0.3530 | 0.7775 |
| | CTN-067 M3 vs. CTN-067 M6 | 11 | 714.182 | 236.439 | 11 | 673.182 | 274.032 | 39.141 | 42.270 | 0.3661 | 0.7914 |

| | | | | | | | | | | | |
|---|---|---|---|---|---|---|---|---|---|---|---|
| CD56+/GLUT1+ \| Freq. of Parent | Control vs. CTN-067 M0 | 8 | 18.816 | 6.595 | 11 | 18.313 | 7.032 | -3.702 | 4.729 | 0.4434 | 0.8613 |
| | Control vs. CTN-067 M3 | 8 | 18.816 | 6.595 | 11 | 23.891 | 8.499 | -7.960 | 4.155 | 0.0705 | 0.2547 |
| | Control vs. CTN-067 M6 | 8 | 18.816 | 6.595 | 11 | 22.100 | 8.829 | -6.103 | 4.133 | 0.1561 | 0.4701 |
| | CTN-067 M0 vs. CTN-067 M3 | 11 | 18.313 | 7.032 | 11 | 23.891 | 8.499 | -4.258 | 2.495 | 0.1041 | 0.3476 |
| | CTN-067 M0 vs. CTN-067 M6 | 11 | 18.313 | 7.032 | 11 | 22.100 | 8.829 | -2.402 | 2.523 | 0.3531 | 0.7776 |
| | CTN-067 M3 vs. CTN-067 M6 | 11 | 23.891 | 8.499 | 11 | 22.100 | 8.829 | 1.857 | 2.198 | 0.4088 | 0.8325 |
| CD56+ \| Geometric Mean (BUV615-A :: CD11b) | Control vs. CTN-067 M0 | 8 | 7289.000 | 2165.250 | 11 | 8249.550 | 2402.300 | -909.940 | 1551.970 | 0.5646 | 0.9350 |
| | Control vs. CTN-067 M3 | 8 | 7289.000 | 2165.250 | 11 | 9378.000 | 2679.190 | -2117.810 | 1406.110 | 0.1485 | 0.4534 |
| | Control vs. CTN-067 M6 | 8 | 7289.000 | 2165.250 | 11 | 9256.820 | 2926.270 | -2000.580 | 1400.670 | 0.1694 | 0.4980 |
| | CTN-067 M0 vs. CTN-067 M3 | 11 | 8249.550 | 2402.300 | 11 | 9378.000 | 2679.190 | -1207.880 | 709.580 | 0.1050 | 0.3499 |
| | CTN-067 M0 vs. CTN-067 M6 | 11 | 8249.550 | 2402.300 | 11 | 9256.820 | 2926.270 | -1090.650 | 718.010 | 0.1452 | 0.4463 |
| | CTN-067 M3 vs. CTN-067 M6 | 11 | 9378.000 | 2679.190 | 11 | 9256.820 | 2926.270 | 117.230 | 621.030 | 0.8523 | 0.9975 |
| CD56+/CD11b+ \| Freq. of Parent | Control vs. CTN-067 M0 | 8 | 84.450 | 6.564 | 11 | 86.955 | 5.114 | -1.500 | 3.042 | 0.6276 | 0.9597 |
| | Control vs. CTN-067 M3 | 8 | 84.450 | 6.564 | 11 | 89.009 | 3.582 | -3.925 | 2.847 | 0.1840 | 0.5271 |
| | Control vs. CTN-067 M6 | 8 | 84.450 | 6.564 | 11 | 88.855 | 3.848 | -3.788 | 2.840 | 0.1979 | 0.5536 |
| | CTN-067 M0 vs. CTN-067 M3 | 11 | 86.955 | 5.114 | 11 | 89.009 | 3.582 | -2.424 | 1.142 | 0.0471 | 0.1816 |
| | CTN-067 M0 vs. CTN-067 M6 | 11 | 86.955 | 5.114 | 11 | 88.855 | 3.848 | -2.288 | 1.156 | 0.0624 | 0.2303 |
| | CTN-067 M3 vs. CTN-067 M6 | 11 | 89.009 | 3.582 | 11 | 88.855 | 3.848 | 0.136 | 0.995 | 0.8926 | 0.9991 |
| CD56+ \| Geometric Mean (BUV661-A :: CD27) | Control vs. CTN-067 M0 | 8 | 1011.250 | 222.764 | 11 | 1022.360 | 364.422 | -79.814 | 183.140 | 0.6679 | 0.9715 |
| | Control vs. CTN-067 M3 | 8 | 1011.250 | 222.764 | 11 | 1232.000 | 343.940 | -223.610 | 165.070 | 0.1914 | 0.5413 |

| | | | | | | | | | | | |
|---|---|---|---|---|---|---|---|---|---|---|---|
| | Control vs. CTN-067 M6 | 8 | 1011.250 | 222.764 | 11 | 1254.640 | 373.851 | -242.970 | 164.390 | 0.1558 | 0.4694 |
| | CTN-067 M0 vs. CTN-067 M3 | 11 | 1022.360 | 364.422 | 11 | 1232.000 | 343.940 | -143.790 | 85.952 | 0.1107 | 0.3644 |
| | CTN-067 M0 vs. CTN-067 M6 | 11 | 1022.360 | 364.422 | 11 | 1254.640 | 373.851 | -163.160 | 86.967 | 0.0761 | 0.2709 |
| | CTN-067 M3 vs. CTN-067 M6 | 11 | 1232.000 | 343.940 | 11 | 1254.640 | 373.851 | -19.363 | 75.297 | 0.7998 | 0.9938 |
| CD56+/CD27+ \| Freq. of Parent | Control vs. CTN-067 M0 | 8 | 12.620 | 6.134 | 11 | 14.382 | 7.439 | -3.212 | 4.111 | 0.4443 | 0.8620 |
| | Control vs. CTN-067 M3 | 8 | 12.620 | 6.134 | 11 | 18.253 | 7.628 | -5.464 | 3.747 | 0.1611 | 0.4807 |
| | Control vs. CTN-067 M6 | 8 | 12.620 | 6.134 | 11 | 18.911 | 9.610 | -6.041 | 3.734 | 0.1221 | 0.3926 |
| | CTN-067 M0 vs. CTN-067 M3 | 11 | 14.382 | 7.439 | 11 | 18.253 | 7.628 | -2.252 | 1.820 | 0.2311 | 0.6118 |
| | CTN-067 M0 vs. CTN-067 M6 | 11 | 14.382 | 7.439 | 11 | 18.911 | 9.610 | -2.830 | 1.842 | 0.1410 | 0.4367 |
| | CTN-067 M3 vs. CTN-067 M6 | 11 | 18.253 | 7.628 | 11 | 18.911 | 9.610 | -0.578 | 1.591 | 0.7206 | 0.9831 |
| CD56+ \| Geometric Mean (BV480-A :: CD16) | Control vs. CTN-067 M0 | 8 | 11318.880 | 8331.750 | 11 | 24392.640 | 15393.900 | -20996.000 | 6366.100 | 0.0038 | 0.0182 |
| | Control vs. CTN-067 M3 | 8 | 11318.880 | 8331.750 | 11 | 15329.450 | 11152.850 | -9923.490 | 5522.370 | 0.0883 | 0.3052 |
| | Control vs. CTN-067 M6 | 8 | 11318.880 | 8331.750 | 11 | 12916.180 | 6827.740 | -7410.340 | 5490.090 | 0.1930 | 0.5442 |
| | CTN-067 M0 vs. CTN-067 M3 | 11 | 24392.640 | 15393.900 | 11 | 15329.450 | 11152.850 | 11072.000 | 3546.160 | 0.0056 | 0.0264 |
| | CTN-067 M0 vs. CTN-067 M6 | 11 | 24392.640 | 15393.900 | 11 | 12916.180 | 6827.740 | 13585.000 | 3585.400 | 0.0012 | 0.0062 |
| | CTN-067 M3 vs. CTN-067 M6 | 11 | 15329.450 | 11152.850 | 11 | 12916.180 | 6827.740 | 2513.150 | 3136.250 | 0.4329 | 0.8529 |
| CD56+ \| Geometric Mean (BV510-A :: CCR7) | Control vs. CTN-067 M0 | 8 | 1402.750 | 550.714 | 11 | 1075.820 | 431.654 | 607.430 | 254.360 | 0.0275 | 0.1135 |
| | Control vs. CTN-067 M3 | 8 | 1402.750 | 550.714 | 11 | 1191.910 | 280.784 | 387.060 | 235.950 | 0.1174 | 0.3810 |
| | Control vs. CTN-067 M6 | 8 | 1402.750 | 550.714 | 11 | 1216.360 | 342.806 | 357.420 | 235.270 | 0.1452 | 0.4462 |
| | CTN-067 M0 vs. | 11 | 1075.820 | 431.654 | 11 | 1191.910 | 280.784 | -220.380 | 101.430 | 0.0427 | 0.1668 |

| | | | | | | | | | | | |
|---|---|---|---|---|---|---|---|---|---|---|---|
| | CTN-067 M3 | | | | | | | | | | |
| | CTN-067 M0 vs. CTN-067 M6 | 11 | 1075.820 | 431.654 | 11 | 1216.360 | 342.806 | -250.020 | 102.660 | 0.0249 | 0.1040 |
| | CTN-067 M3 vs. CTN-067 M6 | 11 | 1191.910 | 280.784 | 11 | 1216.360 | 342.806 | -29.639 | 88.447 | 0.7412 | 0.9866 |
| CD56+/CCR7+ \| Freq. of Parent | Control vs. CTN-067 M0 | 8 | 14.840 | 5.216 | 11 | 17.557 | 6.074 | -1.351 | 3.800 | 0.7261 | 0.9841 |
| | Control vs. CTN-067 M3 | 8 | 14.840 | 5.216 | 11 | 15.157 | 6.819 | 0.172 | 3.535 | 0.9618 | 1.0000 |
| | Control vs. CTN-067 M6 | 8 | 14.840 | 5.216 | 11 | 15.874 | 8.203 | -0.589 | 3.525 | 0.8692 | 0.9983 |
| | CTN-067 M0 vs. CTN-067 M3 | 11 | 17.557 | 6.074 | 11 | 15.157 | 6.819 | 1.523 | 1.488 | 0.3190 | 0.7382 |
| | CTN-067 M0 vs. CTN-067 M6 | 11 | 17.557 | 6.074 | 11 | 15.874 | 8.203 | 0.763 | 1.506 | 0.6184 | 0.9566 |
| | CTN-067 M3 vs. CTN-067 M6 | 11 | 15.157 | 6.819 | 11 | 15.874 | 8.203 | -0.760 | 1.297 | 0.5649 | 0.9351 |
| CD56+ \| Geometric Mean (BV605-A :: CD45RA) | Control vs. CTN-067 M0 | 8 | 120482.250 | 28429.300 | 11 | 112094.270 | 34721.400 | 7814.440 | 22633.000 | 0.7337 | 0.9854 |
| | Control vs. CTN-067 M3 | 8 | 120482.250 | 28429.300 | 11 | 116501.000 | 38048.810 | -79.741 | 19580.000 | 0.9968 | 1.0000 |
| | Control vs. CTN-067 M6 | 8 | 120482.250 | 28429.300 | 11 | 111493.090 | 41791.880 | 4754.780 | 19463.000 | 0.8096 | 0.9947 |
| | CTN-067 M0 vs. CTN-067 M3 | 11 | 112094.270 | 34721.400 | 11 | 116501.000 | 38048.810 | -7894.180 | 12752.000 | 0.5432 | 0.9247 |
| | CTN-067 M0 vs. CTN-067 M6 | 11 | 112094.270 | 34721.400 | 11 | 111493.090 | 41791.880 | -3059.660 | 12892.000 | 0.8149 | 0.9951 |
| | CTN-067 M3 vs. CTN-067 M6 | 11 | 116501.000 | 38048.810 | 11 | 111493.090 | 41791.880 | 4834.520 | 11288.000 | 0.6733 | 0.9729 |
| CD56+/CD45RA+ \| Freq. of Parent | Control vs. CTN-067 M0 | 8 | 97.938 | 1.064 | 11 | 95.527 | 3.899 | 2.169 | 2.440 | 0.3852 | 0.8106 |
| | Control vs. CTN-067 M3 | 8 | 97.938 | 1.064 | 11 | 96.245 | 2.745 | 1.478 | 2.141 | 0.4985 | 0.8996 |
| | Control vs. CTN-067 M6 | 8 | 97.938 | 1.064 | 11 | 94.618 | 5.986 | 3.106 | 2.130 | 0.1610 | 0.4805 |
| | CTN-067 M0 vs. CTN-067 M3 | 11 | 95.527 | 3.899 | 11 | 96.245 | 2.745 | -0.692 | 1.295 | 0.5996 | 0.9497 |
| | CTN-067 M0 vs. CTN-067 M6 | 11 | 95.527 | 3.899 | 11 | 94.618 | 5.986 | 0.937 | 1.310 | 0.4830 | 0.8897 |

| | | | | | | | | | | | |
|---|---|---|---|---|---|---|---|---|---|---|---|
| | CTN-067 M3 vs. CTN-067 M6 | 11 | 96.245 | 2.745 | 11 | 94.618 | 5.986 | 1.629 | 1.142 | 0.1699 | 0.4989 |
| CD56+ \| Geometric Mean (BV650-A :: CD15) | Control vs. CTN-067 M0 | 8 | 215.438 | 475.185 | 11 | -75.902 | 454.213 | 108.750 | 286.910 | 0.7089 | 0.9809 |
| | Control vs. CTN-067 M3 | 8 | 215.438 | 475.185 | 11 | 141.773 | 402.748 | -36.484 | 254.940 | 0.8877 | 0.9989 |
| | Control vs. CTN-067 M6 | 8 | 215.438 | 475.185 | 11 | 150.825 | 468.798 | -41.936 | 253.740 | 0.8705 | 0.9983 |
| | CTN-067 M0 vs. CTN-067 M3 | 11 | -75.902 | 454.213 | 11 | 141.773 | 402.748 | -145.240 | 144.020 | 0.3259 | 0.7465 |
| | CTN-067 M0 vs. CTN-067 M6 | 11 | -75.902 | 454.213 | 11 | 150.825 | 468.798 | -150.690 | 145.690 | 0.3140 | 0.7318 |
| | CTN-067 M3 vs. CTN-067 M6 | 11 | 141.773 | 402.748 | 11 | 150.825 | 468.798 | -5.451 | 126.530 | 0.9661 | 1.0000 |
| CD56+/CD15+ \| Freq. of Parent | Control vs. CTN-067 M0 | 8 | 19.163 | 9.207 | 11 | 14.120 | 7.140 | 2.261 | 5.293 | 0.6741 | 0.9731 |
| | Control vs. CTN-067 M3 | 8 | 19.163 | 9.207 | 11 | 17.966 | 8.160 | -0.490 | 4.737 | 0.9187 | 0.9996 |
| | Control vs. CTN-067 M6 | 8 | 19.163 | 9.207 | 11 | 16.705 | 8.790 | 0.827 | 4.716 | 0.8627 | 0.9980 |
| | CTN-067 M0 vs. CTN-067 M3 | 11 | 14.120 | 7.140 | 11 | 17.966 | 8.160 | -2.751 | 2.571 | 0.2981 | 0.7115 |
| | CTN-067 M0 vs. CTN-067 M6 | 11 | 14.120 | 7.140 | 11 | 16.705 | 8.790 | -1.434 | 2.601 | 0.5878 | 0.9450 |
| | CTN-067 M3 vs. CTN-067 M6 | 11 | 17.966 | 8.160 | 11 | 16.705 | 8.790 | 1.316 | 2.256 | 0.5664 | 0.9358 |
| CD56+ \| Geometric Mean (BV711-A :: CD86) | Control vs. CTN-067 M0 | 8 | 492.000 | 88.737 | 11 | 645.364 | 261.627 | -285.060 | 100.970 | 0.0109 | 0.0489 |
| | Control vs. CTN-067 M3 | 8 | 492.000 | 88.737 | 11 | 514.364 | 175.451 | -125.410 | 86.329 | 0.1626 | 0.4838 |
| | Control vs. CTN-067 M6 | 8 | 492.000 | 88.737 | 11 | 572.273 | 118.067 | -181.900 | 85.764 | 0.0473 | 0.1823 |
| | CTN-067 M0 vs. CTN-067 M3 | 11 | 645.364 | 261.627 | 11 | 514.364 | 175.451 | 159.650 | 59.899 | 0.0153 | 0.0669 |
| | CTN-067 M0 vs. CTN-067 M6 | 11 | 645.364 | 261.627 | 11 | 572.273 | 118.067 | 103.170 | 60.534 | 0.1046 | 0.3489 |
| | CTN-067 M3 vs. CTN-067 M6 | 11 | 514.364 | 175.451 | 11 | 572.273 | 118.067 | -56.485 | 53.282 | 0.3024 | 0.7171 |

| | | | | | | | | | | | |
|---|---|---|---|---|---|---|---|---|---|---|---|
| CD56+/CD86+ \| Freq. of Parent | Control vs. CTN-067 M0 | 8 | 5.509 | 2.985 | 11 | 7.885 | 4.986 | -5.067 | 2.078 | 0.0247 | 0.1033 |
| | Control vs. CTN-067 M3 | 8 | 5.509 | 2.985 | 11 | 5.285 | 3.277 | -1.908 | 1.816 | 0.3066 | 0.7225 |
| | Control vs. CTN-067 M6 | 8 | 5.509 | 2.985 | 11 | 6.571 | 3.202 | -3.165 | 1.806 | 0.0957 | 0.3255 |
| | CTN-067 M0 vs. CTN-067 M3 | 11 | 7.885 | 4.986 | 11 | 5.285 | 3.277 | 3.160 | 1.122 | 0.0110 | 0.0495 |
| | CTN-067 M0 vs. CTN-067 M6 | 11 | 7.885 | 4.986 | 11 | 6.571 | 3.202 | 1.902 | 1.134 | 0.1100 | 0.3625 |
| | CTN-067 M3 vs. CTN-067 M6 | 11 | 5.285 | 3.277 | 11 | 6.571 | 3.202 | -1.258 | 0.990 | 0.2192 | 0.5917 |
| CD56+ \| Geometric Mean (BV750-A :: CD8) | Control vs. CTN-067 M0 | 8 | 2904.750 | 545.317 | 11 | 3024.270 | 745.140 | -195.960 | 374.850 | 0.6072 | 0.9526 |
| | Control vs. CTN-067 M3 | 8 | 2904.750 | 545.317 | 11 | 2885.730 | 656.016 | -103.690 | 341.940 | 0.7650 | 0.9900 |
| | Control vs. CTN-067 M6 | 8 | 2904.750 | 545.317 | 11 | 2785.550 | 682.236 | -5.813 | 340.710 | 0.9866 | 1.0000 |
| | CTN-067 M0 vs. CTN-067 M3 | 11 | 3024.270 | 745.140 | 11 | 2885.730 | 656.016 | 92.265 | 165.310 | 0.5833 | 0.9432 |
| | CTN-067 M0 vs. CTN-067 M6 | 11 | 3024.270 | 745.140 | 11 | 2785.550 | 682.236 | 190.150 | 167.290 | 0.2698 | 0.6722 |
| | CTN-067 M3 vs. CTN-067 M6 | 11 | 2885.730 | 656.016 | 11 | 2785.550 | 682.236 | 97.881 | 144.520 | 0.5064 | 0.9044 |
| CD56+CD8+ \| Freq. of Parent | Control vs. CTN-067 M0 | 8 | 11.656 | 8.712 | 11 | 11.309 | 6.223 | -1.721 | 3.809 | 0.6565 | 0.9685 |
| | Control vs. CTN-067 M3 | 8 | 11.656 | 8.712 | 11 | 10.345 | 4.814 | -0.365 | 3.573 | 0.9198 | 0.9996 |
| | Control vs. CTN-067 M6 | 8 | 11.656 | 8.712 | 11 | 10.823 | 4.720 | -0.824 | 3.564 | 0.8197 | 0.9955 |
| | CTN-067 M0 vs. CTN-067 M3 | 11 | 11.309 | 6.223 | 11 | 10.345 | 4.814 | 1.356 | 1.402 | 0.3456 | 0.7694 |
| | CTN-067 M0 vs. CTN-067 M6 | 11 | 11.309 | 6.223 | 11 | 10.823 | 4.720 | 0.898 | 1.420 | 0.5348 | 0.9203 |
| | CTN-067 M3 vs. CTN-067 M6 | 11 | 10.345 | 4.814 | 11 | 10.823 | 4.720 | -0.459 | 1.221 | 0.7113 | 0.9814 |
| CD56+ \| Geometric Mean (PE-A :: CD38) | Control vs. CTN-067 M0 | 8 | 30264.750 | 11640.850 | 11 | 48340.090 | 25094.310 | -4560.580 | 11241.000 | 0.6895 | 0.9768 |
| | Control vs. CTN-067 M3 | 8 | 30264.750 | 11640.850 | 11 | 39374.270 | 19008.410 | -335.030 | 10020.000 | 0.9737 | 1.0000 |

| | | | | | | | | | | | |
|---|---|---|---|---|---|---|---|---|---|---|---|
| | Control vs. CTN-067 M6 | 8 | 30264.750 | 11640.850 | 11 | 35864.730 | 20466.630 | 2938.840 | 9973.700 | 0.7714 | 0.9908 |
| | CTN-067 M0 vs. CTN-067 M3 | 11 | 48340.090 | 25094.310 | 11 | 39374.270 | 19008.410 | 4225.550 | 5563.890 | 0.4569 | 0.8715 |
| | CTN-067 M0 vs. CTN-067 M6 | 11 | 48340.090 | 25094.310 | 11 | 35864.730 | 20466.630 | 7499.420 | 5628.620 | 0.1985 | 0.5546 |
| | CTN-067 M3 vs. CTN-067 M6 | 11 | 39374.270 | 19008.410 | 11 | 35864.730 | 20466.630 | 3273.870 | 4885.100 | 0.5108 | 0.9070 |
| CD56+/CD38+ \| Freq. of Parent | Control vs. CTN-067 M0 | 8 | 92.075 | 3.373 | 11 | 91.855 | 4.149 | 2.180 | 2.123 | 0.3173 | 0.7360 |
| | Control vs. CTN-067 M3 | 8 | 92.075 | 3.373 | 11 | 91.509 | 3.204 | 2.027 | 1.885 | 0.2959 | 0.7085 |
| | Control vs. CTN-067 M6 | 8 | 92.075 | 3.373 | 11 | 90.918 | 3.739 | 2.593 | 1.876 | 0.1831 | 0.5253 |
| | CTN-067 M0 vs. CTN-067 M3 | 11 | 91.855 | 4.149 | 11 | 91.509 | 3.204 | -0.154 | 1.068 | 0.8872 | 0.9989 |
| | CTN-067 M0 vs. CTN-067 M6 | 11 | 91.855 | 4.149 | 11 | 90.918 | 3.739 | 0.413 | 1.081 | 0.7069 | 0.9805 |
| | CTN-067 M3 vs. CTN-067 M6 | 11 | 91.509 | 3.204 | 11 | 90.918 | 3.739 | 0.566 | 0.939 | 0.5536 | 0.9298 |
| CD56+ \| Geometric Mean (R718-A :: CCR5) | Control vs. CTN-067 M0 | 8 | 652.000 | 283.503 | 11 | 1163.730 | 447.012 | -289.340 | 203.800 | 0.1719 | 0.5030 |
| | Control vs. CTN-067 M3 | 8 | 652.000 | 283.503 | 11 | 1097.730 | 272.210 | -265.890 | 182.140 | 0.1607 | 0.4798 |
| | Control vs. CTN-067 M6 | 8 | 652.000 | 283.503 | 11 | 1188.640 | 402.248 | -358.910 | 181.320 | 0.0624 | 0.2303 |
| | CTN-067 M0 vs. CTN-067 M3 | 11 | 1163.730 | 447.012 | 11 | 1097.730 | 272.210 | 23.457 | 99.638 | 0.8164 | 0.9952 |
| | CTN-067 M0 vs. CTN-067 M6 | 11 | 1163.730 | 447.012 | 11 | 1188.640 | 402.248 | -69.567 | 100.800 | 0.4985 | 0.8996 |
| | CTN-067 M3 vs. CTN-067 M6 | 11 | 1097.730 | 272.210 | 11 | 1188.640 | 402.248 | -93.024 | 87.433 | 0.3007 | 0.7148 |
| CD56+/CCR5+ \| Freq. of Parent | Control vs. CTN-067 M0 | 8 | 16.253 | 8.431 | 11 | 29.155 | 12.081 | -7.794 | 5.876 | 0.2004 | 0.5582 |
| | Control vs. CTN-067 M3 | 8 | 16.253 | 8.431 | 11 | 28.682 | 9.605 | -7.824 | 5.425 | 0.1655 | 0.4899 |
| | Control vs. CTN-067 M6 | 8 | 16.253 | 8.431 | 11 | 30.411 | 13.313 | -9.578 | 5.408 | 0.0926 | 0.3171 |
| | CTN-067 M0 vs. | 11 | 29.155 | 12.081 | 11 | 28.682 | 9.605 | -0.030 | 2.416 | 0.9902 | 1.0000 |

| | | | | | | | | | | | |
|---|---|---|---|---|---|---|---|---|---|---|---|
| | CTN-067 M3 | | | | | | | | | | |
| | CTN-067 M0 vs. CTN-067 M6 | 1 1 | 29.155 | 12.081 | 1 1 | 30.411 | 13.313 | -1.784 | 2.44 5 | 0.47 44 | 0.8839 |
| | CTN-067 M3 vs. CTN-067 M6 | 1 1 | 28.682 | 9.605 | 1 1 | 30.411 | 13.313 | -1.754 | 2.10 8 | 0.41 56 | 0.8385 |
| CD56+ \| Geometric Mean (RB545-A :: CD57) | Control vs. CTN-067 M0 | 8 | 19552. 630 | 23939. 200 | 1 1 | 4053.4 50 | 2516.0 90 | 15132.00 0 | 851 9.62 0 | 0.09 17 | 0.3147 |
| | Control vs. CTN-067 M3 | 8 | 19552. 630 | 23939. 200 | 1 1 | 5164.9 10 | 5265.1 40 | 13784.00 0 | 841 9.13 0 | 0.11 80 | 0.3827 |
| | Control vs. CTN-067 M6 | 8 | 19552. 630 | 23939. 200 | 1 1 | 5588.2 70 | 7220.3 60 | 13349.00 0 | 841 5.55 0 | 0.12 92 | 0.4095 |
| | CTN-067 M0 vs. CTN-067 M3 | 1 1 | 4053.4 50 | 2516.0 90 | 1 1 | 5164.9 10 | 5265.1 40 | - 1348.240 | 136 4.25 0 | 0.33 54 | 0.7577 |
| | CTN-067 M0 vs. CTN-067 M6 | 1 1 | 4053.4 50 | 2516.0 90 | 1 1 | 5588.2 70 | 7220.3 60 | - 1783.370 | 138 1.54 0 | 0.21 22 | 0.5795 |
| | CTN-067 M3 vs. CTN-067 M6 | 1 1 | 5164.9 10 | 5265.1 40 | 1 1 | 5588.2 70 | 7220.3 60 | -435.140 | 118 1.67 0 | 0.71 68 | 0.9824 |
| CD56+/CD 57+ \| Freq. of Parent | Control vs. CTN-067 M0 | 8 | 56.063 | 17.189 | 1 1 | 33.200 | 9.913 | 21.431 | 7.89 8 | 0.01 38 | 0.0609 |
| | Control vs. CTN-067 M3 | 8 | 56.063 | 17.189 | 1 1 | 35.655 | 12.281 | 19.025 | 7.58 0 | 0.02 13 | 0.0904 |
| | Control vs. CTN-067 M6 | 8 | 56.063 | 17.189 | 1 1 | 33.609 | 12.924 | 21.073 | 7.56 9 | 0.01 18 | 0.0529 |
| | CTN-067 M0 vs. CTN-067 M3 | 1 1 | 33.200 | 9.913 | 1 1 | 35.655 | 12.281 | -2.405 | 2.33 9 | 0.31 67 | 0.7353 |
| | CTN-067 M0 vs. CTN-067 M6 | 1 1 | 33.200 | 9.913 | 1 1 | 33.609 | 12.924 | -0.357 | 2.36 8 | 0.88 17 | 0.9987 |
| | CTN-067 M3 vs. CTN-067 M6 | 1 1 | 35.655 | 12.281 | 1 1 | 33.609 | 12.924 | 2.048 | 2.03 1 | 0.32 60 | 0.7467 |
| CD56+ \| Geometric Mean (RB705-A :: CCR2) | Control vs. CTN-067 M0 | 8 | 948.50 0 | 187.04 7 | 1 1 | 1139.8 20 | 449.99 0 | -228.720 | 235. 670 | 0.34 40 | 0.7675 |
| | Control vs. CTN-067 M3 | 8 | 948.50 0 | 187.04 7 | 1 1 | 1100.4 50 | 383.30 2 | -160.370 | 201. 320 | 0.43 55 | 0.8551 |
| | Control vs. CTN-067 M6 | 8 | 948.50 0 | 187.04 7 | 1 1 | 1319.2 70 | 352.26 8 | -377.740 | 199. 990 | 0.07 43 | 0.2657 |
| | CTN-067 M0 vs. CTN-067 M3 | 1 1 | 1139.8 20 | 449.99 0 | 1 1 | 1100.4 50 | 383.30 2 | 68.352 | 140. 380 | 0.63 19 | 0.9611 |
| | CTN-067 M0 vs. CTN-067 M6 | 1 1 | 1139.8 20 | 449.99 0 | 1 1 | 1319.2 70 | 352.26 8 | -149.030 | 141. 860 | 0.30 67 | 0.7226 |

| | | | | | | | | | | | |
|---|---|---|---|---|---|---|---|---|---|---|---|
| | CTN-067 M3 vs. CTN-067 M6 | 11 | 1100.450 | 383.302 | 11 | 1319.270 | 352.268 | -217.380 | 124.930 | 0.0980 | 0.3316 |
| CD56+/CCR2+ \| Freq. of Parent | Control vs. CTN-067 M0 | 8 | 11.319 | 4.438 | 11 | 17.390 | 8.312 | -7.051 | 4.632 | 0.1445 | 0.4446 |
| | Control vs. CTN-067 M3 | 8 | 11.319 | 4.438 | 11 | 17.342 | 8.541 | -6.299 | 4.080 | 0.1391 | 0.4324 |
| | Control vs. CTN-067 M6 | 8 | 11.319 | 4.438 | 11 | 20.555 | 8.006 | -9.477 | 4.059 | 0.0307 | 0.1251 |
| | CTN-067 M0 vs. CTN-067 M3 | 11 | 17.390 | 8.312 | 11 | 17.342 | 8.541 | 0.751 | 2.418 | 0.7595 | 0.9893 |
| | CTN-067 M0 vs. CTN-067 M6 | 11 | 17.390 | 8.312 | 11 | 20.555 | 8.006 | -2.427 | 2.446 | 0.3336 | 0.7556 |
| | CTN-067 M3 vs. CTN-067 M6 | 11 | 17.342 | 8.541 | 11 | 20.555 | 8.006 | -3.178 | 2.129 | 0.1520 | 0.4612 |
| CD56+ \| Geometric Mean (RB780-A :: HLA-DR) | Control vs. CTN-067 M0 | 8 | 4022.880 | 1161.950 | 11 | 7015.270 | 1829.710 | -3063.600 | 1159.590 | 0.0161 | 0.0701 |
| | Control vs. CTN-067 M3 | 8 | 4022.880 | 1161.950 | 11 | 6259.180 | 2117.740 | -2471.370 | 1023.520 | 0.0260 | 0.1081 |
| | Control vs. CTN-067 M6 | 8 | 4022.880 | 1161.950 | 11 | 6362.270 | 2430.480 | -2582.610 | 1018.360 | 0.0201 | 0.0860 |
| | CTN-067 M0 vs. CTN-067 M3 | 11 | 7015.270 | 1829.710 | 11 | 6259.180 | 2117.740 | 592.220 | 599.630 | 0.3357 | 0.7581 |
| | CTN-067 M0 vs. CTN-067 M6 | 11 | 7015.270 | 1829.710 | 11 | 6362.270 | 2430.480 | 480.990 | 606.500 | 0.4375 | 0.8567 |
| | CTN-067 M3 vs. CTN-067 M6 | 11 | 6259.180 | 2117.740 | 11 | 6362.270 | 2430.480 | -111.240 | 527.670 | 0.8353 | 0.9966 |
| CD56+/HLA-DR+ \| Freq. of Parent | Control vs. CTN-067 M0 | 8 | 67.150 | 14.281 | 11 | 78.873 | 7.119 | -7.798 | 7.209 | 0.2929 | 0.7045 |
| | Control vs. CTN-067 M3 | 8 | 67.150 | 14.281 | 11 | 77.973 | 12.523 | -8.083 | 6.629 | 0.2376 | 0.6226 |
| | Control vs. CTN-067 M6 | 8 | 67.150 | 14.281 | 11 | 76.773 | 12.903 | -6.942 | 6.607 | 0.3066 | 0.7225 |
| | CTN-067 M0 vs. CTN-067 M3 | 11 | 78.873 | 7.119 | 11 | 77.973 | 12.523 | -0.285 | 3.038 | 0.9263 | 0.9997 |
| | CTN-067 M0 vs. CTN-067 M6 | 11 | 78.873 | 7.119 | 11 | 76.773 | 12.903 | 0.856 | 3.075 | 0.7836 | 0.9922 |
| | CTN-067 M3 vs. CTN-067 M6 | 11 | 77.973 | 12.523 | 11 | 76.773 | 12.903 | 1.141 | 2.653 | 0.6719 | 0.9726 |

| | | | | | | | | | | | |
|---|---|---|---|---|---|---|---|---|---|---|---|
| CD56+ \| Geometric Mean (RY586-A :: CXCR4) | Control vs. CTN-067 M0 | 8 | 2296.130 | 1212.910 | 11 | 10187.090 | 8334.900 | -6211.130 | 3710.400 | 0.1105 | 0.3639 |
| | Control vs. CTN-067 M3 | 8 | 2296.130 | 1212.910 | 11 | 4700.550 | 5764.540 | -1199.580 | 3101.230 | 0.7032 | 0.9797 |
| | Control vs. CTN-067 M6 | 8 | 2296.130 | 1212.910 | 11 | 3686.640 | 3400.190 | -209.280 | 3077.430 | 0.9465 | 0.9999 |
| | CTN-067 M0 vs. CTN-067 M3 | 11 | 10187.090 | 8334.900 | 11 | 4700.550 | 5764.540 | 5011.560 | 2562.260 | 0.0653 | 0.2391 |
| | CTN-067 M0 vs. CTN-067 M6 | 11 | 10187.090 | 8334.900 | 11 | 3686.640 | 3400.190 | 6001.850 | 2584.750 | 0.0315 | 0.1281 |
| | CTN-067 M3 vs. CTN-067 M6 | 11 | 4700.550 | 5764.540 | 11 | 3686.640 | 3400.190 | 990.290 | 2330.680 | 0.6757 | 0.9735 |
| CD56+/CXCR4+ \| Freq. of Parent | Control vs. CTN-067 M0 | 8 | 39.188 | 22.577 | 11 | 69.136 | 19.780 | -15.140 | 14.777 | 0.3184 | 0.7374 |
| | Control vs. CTN-067 M3 | 8 | 39.188 | 22.577 | 11 | 46.118 | 28.953 | 3.927 | 13.009 | 0.7660 | 0.9901 |
| | Control vs. CTN-067 M6 | 8 | 39.188 | 22.577 | 11 | 48.764 | 25.867 | 1.085 | 12.942 | 0.9341 | 0.9998 |
| | CTN-067 M0 vs. CTN-067 M3 | 11 | 69.136 | 19.780 | 11 | 46.118 | 28.953 | 19.066 | 7.729 | 0.0233 | 0.0981 |
| | CTN-067 M0 vs. CTN-067 M6 | 11 | 69.136 | 19.780 | 11 | 48.764 | 25.867 | 16.225 | 7.818 | 0.0518 | 0.1968 |
| | CTN-067 M3 vs. CTN-067 M6 | 11 | 46.118 | 28.953 | 11 | 48.764 | 25.867 | -2.842 | 6.806 | 0.6810 | 0.9748 |
| CD56+ \| Geometric Mean (RY610-A :: CD69) | Control vs. CTN-067 M0 | 8 | 962.500 | 510.005 | 11 | 1377.090 | 898.964 | -32.893 | 485.110 | 0.9466 | 0.9999 |
| | Control vs. CTN-067 M3 | 8 | 962.500 | 510.005 | 11 | 1587.550 | 1081.050 | -322.870 | 419.940 | 0.4514 | 0.8674 |
| | Control vs. CTN-067 M6 | 8 | 962.500 | 510.005 | 11 | 1446.270 | 501.298 | -185.560 | 417.440 | 0.6617 | 0.9699 |
| | CTN-067 M0 vs. CTN-067 M3 | 11 | 1377.090 | 898.964 | 11 | 1587.550 | 1081.050 | -289.980 | 272.620 | 0.3008 | 0.7150 |
| | CTN-067 M0 vs. CTN-067 M6 | 11 | 1377.090 | 898.964 | 11 | 1446.270 | 501.298 | -152.660 | 275.620 | 0.5861 | 0.9443 |
| | CTN-067 M3 vs. CTN-067 M6 | 11 | 1587.550 | 1081.050 | 11 | 1446.270 | 501.298 | 137.320 | 241.280 | 0.5759 | 0.9400 |
| CD56+/CD69+ \| Freq. of Parent | Control vs. CTN-067 M0 | 8 | 22.600 | 7.817 | 11 | 26.085 | 13.819 | 6.220 | 6.976 | 0.3837 | 0.8092 |
| | Control vs. CTN-067 M3 | 8 | 22.600 | 7.817 | 11 | 28.670 | 15.095 | 0.938 | 5.965 | 0.8767 | 0.9986 |

| | | | | | | | | | | | |
|---|---|---|---|---|---|---|---|---|---|---|---|
| | Control vs. CTN-067 M6 | 8 | 22.600 | 7.817 | 11 | 27.791 | 7.638 | 1.683 | 5.926 | 0.7795 | 0.9917 |
| | CTN-067 M0 vs. CTN-067 M3 | 11 | 26.085 | 13.819 | 11 | 28.670 | 15.095 | -5.282 | 4.135 | 0.2168 | 0.5876 |
| | CTN-067 M0 vs. CTN-067 M6 | 11 | 26.085 | 13.819 | 11 | 27.791 | 7.638 | -4.537 | 4.179 | 0.2912 | 0.7022 |
| | CTN-067 M3 vs. CTN-067 M6 | 11 | 28.670 | 15.095 | 11 | 27.791 | 7.638 | 0.745 | 3.678 | 0.8416 | 0.9970 |
| CD56+ \| Geometric Mean (RY703-A :: PD-1) | Control vs. CTN-067 M0 | 8 | 613.000 | 186.496 | 11 | 500.818 | 196.258 | 191.810 | 108.370 | 0.0928 | 0.3175 |
| | Control vs. CTN-067 M3 | 8 | 613.000 | 186.496 | 11 | 607.909 | 193.723 | 66.268 | 100.840 | 0.5190 | 0.9117 |
| | Control vs. CTN-067 M6 | 8 | 613.000 | 186.496 | 11 | 620.364 | 198.345 | 52.896 | 100.570 | 0.6050 | 0.9517 |
| | CTN-067 M0 vs. CTN-067 M3 | 11 | 500.818 | 196.258 | 11 | 607.909 | 193.723 | -125.550 | 42.300 | 0.0079 | 0.0364 |
| | CTN-067 M0 vs. CTN-067 M6 | 11 | 500.818 | 196.258 | 11 | 620.364 | 198.345 | -138.920 | 42.816 | 0.0043 | 0.0203 |
| | CTN-067 M3 vs. CTN-067 M6 | 11 | 607.909 | 193.723 | 11 | 620.364 | 198.345 | -13.372 | 36.870 | 0.7208 | 0.9832 |
| CD56+/PD-1+ \| Freq. of Parent | Control vs. CTN-067 M0 | 8 | 16.201 | 7.885 | 11 | 14.707 | 6.075 | 6.523 | 4.100 | 0.1281 | 0.4069 |
| | Control vs. CTN-067 M3 | 8 | 16.201 | 7.885 | 11 | 17.883 | 7.347 | 1.494 | 3.662 | 0.6879 | 0.9764 |
| | Control vs. CTN-067 M6 | 8 | 16.201 | 7.885 | 11 | 16.099 | 4.875 | 3.185 | 3.645 | 0.3932 | 0.8182 |
| | CTN-067 M0 vs. CTN-067 M3 | 11 | 14.707 | 6.075 | 11 | 17.883 | 7.347 | -5.029 | 2.010 | 0.0216 | 0.0917 |
| | CTN-067 M0 vs. CTN-067 M6 | 11 | 14.707 | 6.075 | 11 | 16.099 | 4.875 | -3.338 | 2.034 | 0.1172 | 0.3805 |
| | CTN-067 M3 vs. CTN-067 M6 | 11 | 17.883 | 7.347 | 11 | 16.099 | 4.875 | 1.692 | 1.764 | 0.3497 | 0.7739 |
| CD56+CD16+ \| Freq. of Parent | Control vs. CTN-067 M0 | 8 | 10.794 | 11.978 | 11 | 15.571 | 8.933 | -6.611 | 5.640 | 0.2556 | 0.6510 |
| | Control vs. CTN-067 M3 | 8 | 10.794 | 11.978 | 11 | 11.825 | 8.727 | -2.215 | 5.201 | 0.6749 | 0.9733 |
| | Control vs. CTN-067 M6 | 8 | 10.794 | 11.978 | 11 | 11.734 | 6.080 | -2.091 | 5.185 | 0.6912 | 0.9772 |
| | CTN-067 M0 vs. | 11 | 15.571 | 8.933 | 11 | 11.825 | 8.727 | 4.396 | 2.334 | 0.0751 | 0.2680 |

| | | | | | | | | | | | |
|---|---|---|---|---|---|---|---|---|---|---|---|
| | CTN-067 M3 | | | | | | | | | | |
| | CTN-067 M0 vs. CTN-067 M6 | 11 | 15.571 | 8.933 | 11 | 11.734 | 6.080 | 4.520 | 2.362 | 0.0709 | 0.2558 |
| | CTN-067 M3 vs. CTN-067 M6 | 11 | 11.825 | 8.727 | 11 | 11.734 | 6.080 | 0.124 | 2.037 | 0.9520 | 0.9999 |
| CD56+CD16- \| Freq. of Parent | Control vs. CTN-067 M0 | 8 | 7.029 | 5.335 | 11 | 6.962 | 6.125 | 1.299 | 2.785 | 0.6462 | 0.9655 |
| | Control vs. CTN-067 M3 | 8 | 7.029 | 5.335 | 11 | 7.053 | 3.692 | 1.316 | 2.600 | 0.6185 | 0.9566 |
| | Control vs. CTN-067 M6 | 8 | 7.029 | 5.335 | 11 | 8.108 | 4.444 | 0.266 | 2.593 | 0.9193 | 0.9996 |
| | CTN-067 M0 vs. CTN-067 M3 | 11 | 6.962 | 6.125 | 11 | 7.053 | 3.692 | 0.018 | 1.061 | 0.9870 | 1.0000 |
| | CTN-067 M0 vs. CTN-067 M6 | 11 | 6.962 | 6.125 | 11 | 8.108 | 4.444 | -1.033 | 1.074 | 0.3486 | 0.7726 |
| | CTN-067 M3 vs. CTN-067 M6 | 11 | 7.053 | 3.692 | 11 | 8.108 | 4.444 | -1.050 | 0.925 | 0.2703 | 0.6728 |
| CD56+CD16dim \| Freq. of Parent | Control vs. CTN-067 M0 | 8 | 9.009 | 7.822 | 11 | 4.375 | 1.646 | 4.989 | 2.798 | 0.0905 | 0.3114 |
| | Control vs. CTN-067 M3 | 8 | 9.009 | 7.822 | 11 | 5.006 | 1.545 | 4.259 | 2.752 | 0.1381 | 0.4302 |
| | Control vs. CTN-067 M6 | 8 | 9.009 | 7.822 | 11 | 5.715 | 2.081 | 3.546 | 2.750 | 0.2127 | 0.5803 |
| | CTN-067 M0 vs. CTN-067 M3 | 11 | 4.375 | 1.646 | 11 | 5.006 | 1.545 | -0.730 | 0.531 | 0.1848 | 0.5286 |
| | CTN-067 M0 vs. CTN-067 M6 | 11 | 4.375 | 1.646 | 11 | 5.715 | 2.081 | -1.443 | 0.537 | 0.0146 | 0.0642 |
| | CTN-067 M3 vs. CTN-067 M6 | 11 | 5.006 | 1.545 | 11 | 5.715 | 2.081 | -0.713 | 0.460 | 0.1373 | 0.4284 |
| CD56+CD57++ \| Freq. of Parent | Control vs. CTN-067 M0 | 8 | 14.201 | 16.094 | 11 | 6.643 | 4.978 | 7.251 | 5.985 | 0.2405 | 0.6273 |
| | Control vs. CTN-067 M3 | 8 | 14.201 | 16.094 | 11 | 6.408 | 6.048 | 7.571 | 5.916 | 0.2161 | 0.5863 |
| | Control vs. CTN-067 M6 | 8 | 14.201 | 16.094 | 11 | 6.425 | 5.262 | 7.558 | 5.914 | 0.2166 | 0.5873 |
| | CTN-067 M0 vs. CTN-067 M3 | 11 | 6.643 | 4.978 | 11 | 6.408 | 6.048 | 0.320 | 0.944 | 0.7384 | 0.9862 |
| | CTN-067 M0 vs. CTN-067 M6 | 11 | 6.643 | 4.978 | 11 | 6.425 | 5.262 | 0.307 | 0.956 | 0.7517 | 0.9882 |

| | | | | | | | | | | |
|---|---|---|---|---|---|---|---|---|---|---|
| | CTN-067 M3 vs. CTN-067 M6 | 11 | 6.408 | 6.048 | 11 | 6.425 | 5.262 | -0.013 | 0.817 | 0.9874 | 1.0000 |
| CD56+CD57dim \| Freq. of Parent | Control vs. CTN-067 M0 | 8 | 5.570 | 3.931 | 11 | 4.109 | 1.712 | 1.308 | 1.575 | 0.4167 | 0.8394 |
| | Control vs. CTN-067 M3 | 8 | 5.570 | 3.931 | 11 | 3.994 | 1.649 | 1.484 | 1.462 | 0.3227 | 0.7426 |
| | Control vs. CTN-067 M6 | 8 | 5.570 | 3.931 | 11 | 3.846 | 1.569 | 1.635 | 1.458 | 0.2761 | 0.6812 |
| | CTN-067 M0 vs. CTN-067 M3 | 11 | 4.109 | 1.712 | 11 | 3.994 | 1.649 | 0.176 | 0.627 | 0.7819 | 0.9920 |
| | CTN-067 M0 vs. CTN-067 M6 | 11 | 4.109 | 1.712 | 11 | 3.846 | 1.569 | 0.326 | 0.635 | 0.6131 | 0.9547 |
| | CTN-067 M3 vs. CTN-067 M6 | 11 | 3.994 | 1.649 | 11 | 3.846 | 1.569 | 0.150 | 0.547 | 0.7865 | 0.9925 |
