## Supplemental Table 14 for "Chronic opioid-associated immune dysregulation among people living with HIV"

**Supplementary Table 14.** CD19+ Flow data

| Marker | Comparison (A vs B) | Group A | | | Group B | | | Viral Load adjusted Group A – Group B | | | |
|---|---|---|---|---|---|---|---|---|---|---|---|
| | | n | Mean | SD | N | Mean | SD | Estimate | SE | p | adj p |
| CD3- \| Geometric Mean (RY775-A :: CD19) | Control vs. CTN-067 M0 | 8 | 4886.500 | 1486.780 | 11 | 5351.000 | 3182.160 | -627.460 | 2149.580 | 0.7735 | 0.9911 |
| | Control vs. CTN-067 M3 | 8 | 4886.500 | 1486.780 | 11 | 6038.000 | 3518.060 | -1471.890 | 1887.590 | 0.4451 | 0.8626 |
| | Control vs. CTN-067 M6 | 8 | 4886.500 | 1486.780 | 11 | 7752.640 | 4838.810 | -3194.360 | 1877.630 | 0.1052 | 0.3503 |
| | CTN-067 M0 vs. CTN-067 M3 | 11 | 5351.000 | 3182.160 | 11 | 6038.000 | 3518.060 | -844.430 | 1136.710 | 0.4666 | 0.8785 |
| | CTN-067 M0 vs. CTN-067 M6 | 11 | 5351.000 | 3182.160 | 11 | 7752.640 | 4838.810 | -2566.900 | 1149.620 | 0.0378 | 0.1502 |
| | CTN-067 M3 vs. CTN-067 M6 | 11 | 6038.000 | 3518.060 | 11 | 7752.640 | 4838.810 | -1722.460 | 1001.630 | 0.1017 | 0.3414 |
| CD19+ \| Freq. of Parent | Control vs. CTN-067 M0 | 8 | 46.363 | 7.301 | 11 | 44.009 | 16.107 | -3.564 | 8.141 | 0.6665 | 0.9712 |
| | Control vs. CTN-067 M3 | 8 | 46.363 | 7.301 | 11 | 46.127 | 15.733 | -4.273 | 7.486 | 0.5749 | 0.9396 |
| | Control vs. CTN-067 M6 | 8 | 46.363 | 7.301 | 11 | 51.964 | 16.488 | -10.039 | 7.462 | 0.1944 | 0.5469 |
| | CTN-067 M0 vs. CTN-067 M3 | 11 | 44.009 | 16.107 | 11 | 46.127 | 15.733 | -0.709 | 3.430 | 0.8385 | 0.9968 |
| | CTN-067 M0 vs. CTN-067 M6 | 11 | 44.009 | 16.107 | 11 | 51.964 | 16.488 | -6.475 | 3.471 | 0.0777 | 0.2755 |
| | CTN-067 M3 vs. CTN-067 M6 | 11 | 46.127 | 15.733 | 11 | 51.964 | 16.488 | -5.766 | 2.995 | 0.0693 | 0.2509 |
| CD19+ \| Geometric Mean (APC-Fire 750-A :: CD36) | Control vs. CTN-067 M0 | 8 | 1101.380 | 282.339 | 11 | 1621.360 | 536.455 | -16.174 | 285.210 | 0.9554 | 0.9999 |
| | Control vs. CTN-067 M3 | 8 | 1101.380 | 282.339 | 11 | 1584.450 | 761.255 | -108.360 | 261.980 | 0.6838 | 0.9755 |
| | Control vs. CTN-067 M6 | 8 | 1101.380 | 282.339 | 11 | 1511.180 | 659.524 | -41.509 | 261.120 | 0.8754 | 0.9985 |

| | | | | | | | | | | |
|---|---|---|---|---|---|---|---|---|---|---|
| | CTN-067 M0 vs. CTN-067 M3 | 11 | 1621.360 | 536.455 | 11 | 1584.450 | 761.255 | -92.190 | 120.900 | 0.4551 | 0.8702 |
| | CTN-067 M0 vs. CTN-067 M6 | 11 | 1621.360 | 536.455 | 11 | 1511.180 | 659.524 | -25.335 | 122.360 | 0.8382 | 0.9967 |
| | CTN-067 M3 vs. CTN-067 M6 | 11 | 1584.450 | 761.255 | 11 | 1511.180 | 659.524 | 66.854 | 105.580 | 0.5341 | 0.9200 |
| CD19+/CD36+ \| Freq. of Parent | Control vs. CTN-067 M0 | 8 | 7.685 | 3.165 | 11 | 15.380 | 8.477 | -2.031 | 3.296 | 0.5451 | 0.9257 |
| | Control vs. CTN-067 M3 | 8 | 7.685 | 3.165 | 11 | 11.418 | 4.025 | 0.153 | 2.903 | 0.9584 | 0.9999 |
| | Control vs. CTN-067 M6 | 8 | 7.685 | 3.165 | 11 | 12.788 | 5.159 | -1.305 | 2.889 | 0.6565 | 0.9685 |
| | CTN-067 M0 vs. CTN-067 M3 | 11 | 15.380 | 8.477 | 11 | 11.418 | 4.025 | 2.184 | 1.720 | 0.2194 | 0.5921 |
| | CTN-067 M0 vs. CTN-067 M6 | 11 | 15.380 | 8.477 | 11 | 12.788 | 5.159 | 0.726 | 1.740 | 0.6811 | 0.9748 |
| | CTN-067 M3 vs. CTN-067 M6 | 11 | 11.418 | 4.025 | 11 | 12.788 | 5.159 | -1.458 | 1.514 | 0.3476 | 0.7715 |
| CD19+ \| Geometric Mean (Alexa Fluor 647-A :: GLUT1) | Control vs. CTN-067 M0 | 8 | 318.963 | 235.399 | 11 | 357.645 | 249.909 | 101.030 | 152.550 | 0.5157 | 0.9099 |
| | Control vs. CTN-067 M3 | 8 | 318.963 | 235.399 | 11 | 384.364 | 290.869 | 2.880 | 140.920 | 0.9839 | 1.0000 |
| | Control vs. CTN-067 M6 | 8 | 318.963 | 235.399 | 11 | 325.427 | 296.471 | 58.266 | 140.490 | 0.6830 | 0.9753 |
| | CTN-067 M0 vs. CTN-067 M3 | 11 | 357.645 | 249.909 | 11 | 384.364 | 290.869 | -98.148 | 62.485 | 0.1327 | 0.4178 |
| | CTN-067 M0 vs. CTN-067 M6 | 11 | 357.645 | 249.909 | 11 | 325.427 | 296.471 | -42.763 | 63.243 | 0.5071 | 0.9048 |
| | CTN-067 M3 vs. CTN-067 M6 | 11 | 384.364 | 290.869 | 11 | 325.427 | 296.471 | 55.385 | 54.520 | 0.3225 | 0.7423 |
| CD19+ \| Geometric Mean (BUV615-A :: CD11b) | Control vs. CTN-067 M0 | 8 | 245.375 | 470.800 | 11 | 844.945 | 581.039 | -310.000 | 279.590 | 0.2814 | 0.6887 |
| | Control vs. CTN-067 M3 | 8 | 245.375 | 470.800 | 11 | 854.327 | 494.717 | -338.100 | 260.000 | 0.2090 | 0.5738 |

| | | | | | | | | | | | |
|---|---|---|---|---|---|---|---|---|---|---|---|
| | Control vs. CTN-067 M6 | 8 | 245.375 | 470.800 | 11 | 790.364 | 662.461 | -275.070 | 259.280 | 0.3020 | 0.7166 |
| | CTN-067 M0 vs. CTN-067 M3 | 11 | 844.945 | 581.039 | 11 | 854.327 | 494.717 | -28.100 | 109.660 | 0.8005 | 0.9939 |
| | CTN-067 M0 vs. CTN-067 M6 | 11 | 844.945 | 581.039 | 11 | 790.364 | 662.461 | 34.934 | 111.000 | 0.7564 | 0.9889 |
| | CTN-067 M3 vs. CTN-067 M6 | 11 | 854.327 | 494.717 | 11 | 790.364 | 662.461 | 63.033 | 95.593 | 0.5176 | 0.9109 |
| CD19+/CD11b+ \| Freq. of Parent | Control vs. CTN-067 M0 | 8 | 24.950 | 8.545 | 11 | 35.773 | 10.234 | -3.832 | 4.625 | 0.4177 | 0.8403 |
| | Control vs. CTN-067 M3 | 8 | 24.950 | 8.545 | 11 | 36.227 | 9.087 | -5.068 | 4.300 | 0.2531 | 0.6471 |
| | Control vs. CTN-067 M6 | 8 | 24.950 | 8.545 | 11 | 34.400 | 11.529 | -3.280 | 4.288 | 0.4537 | 0.8692 |
| | CTN-067 M0 vs. CTN-067 M3 | 11 | 35.773 | 10.234 | 11 | 36.227 | 9.087 | -1.237 | 1.816 | 0.5042 | 0.9031 |
| | CTN-067 M0 vs. CTN-067 M6 | 11 | 35.773 | 10.234 | 11 | 34.400 | 11.529 | 0.552 | 1.839 | 0.7674 | 0.9903 |
| | CTN-067 M3 vs. CTN-067 M6 | 11 | 36.227 | 9.087 | 11 | 34.400 | 11.529 | 1.788 | 1.583 | 0.2728 | 0.6764 |
| CD19+ \| Geometric Mean (BUV661-A :: CD27) | Control vs. CTN-067 M0 | 8 | 1445.130 | 518.986 | 11 | 2069.450 | 1111.780 | -66.757 | 76792.000 | 0.9993 | 1.0000 |
| | Control vs. CTN-067 M3 | 8 | 1445.130 | 518.986 | 11 | 2114.550 | 950.116 | -109.930 | 76792.000 | 0.9989 | 1.0000 |
| | Control vs. CTN-067 M6 | 8 | 1445.130 | 518.986 | 11 | 2201.090 | 1313.920 | -196.380 | 76792.000 | 0.9980 | 1.0000 |
| | CTN-067 M0 vs. CTN-067 M3 | 11 | 2069.450 | 1111.780 | 11 | 2114.550 | 950.116 | -43.175 | 124.230 | 0.7320 | 0.9851 |
| | CTN-067 M0 vs. CTN-067 M6 | 11 | 2069.450 | 1111.780 | 11 | 2201.090 | 1313.920 | -129.630 | 125.810 | 0.3158 | 0.7341 |
| | CTN-067 M3 vs. CTN-067 M6 | 11 | 2114.550 | 950.116 | 11 | 2201.090 | 1313.920 | -86.450 | 107.500 | 0.4312 | 0.8516 |
| CD19+/CD27+ \| Freq. of Parent | Control vs. CTN-067 M0 | 8 | 19.726 | 10.157 | 11 | 30.205 | 16.491 | -1.184 | 5.992 | 0.8454 | 0.9972 |

| | | | | | | | | | | | |
|---|---|---|---|---|---|---|---|---|---|---|---|
| | Control vs. CTN-067 M3 | 8 | 19.726 | 10.157 | 11 | 29.531 | 14.535 | -0.586 | 5.701 | 0.9193 | 0.9996 |
| | Control vs. CTN-067 M6 | 8 | 19.726 | 10.157 | 11 | 30.284 | 18.402 | -1.342 | 5.691 | 0.8161 | 0.9952 |
| | CTN-067 M0 vs. CTN-067 M3 | 11 | 30.205 | 16.491 | 11 | 29.531 | 14.535 | 0.599 | 1.951 | 0.7622 | 0.9896 |
| | CTN-067 M0 vs. CTN-067 M6 | 11 | 30.205 | 16.491 | 11 | 30.284 | 18.402 | -0.158 | 1.975 | 0.9373 | 0.9998 |
| | CTN-067 M3 vs. CTN-067 M6 | 11 | 29.531 | 14.535 | 11 | 30.284 | 18.402 | -0.756 | 1.696 | 0.6606 | 0.9696 |
| CD19+ \| Geometric Mean (BV510-A :: CCR7) | Control vs. CTN-067 M0 | 8 | 5946.000 | 1188.700 | 11 | 3363.820 | 890.522 | 3159.550 | 573.710 | <.0001 | 0.0001 |
| | Control vs. CTN-067 M3 | 8 | 5946.000 | 1188.700 | 11 | 3076.910 | 985.389 | 3356.190 | 530.930 | <.0001 | <.0001 |
| | Control vs. CTN-067 M6 | 8 | 5946.000 | 1188.700 | 11 | 3272.090 | 1021.100 | 3156.520 | 529.350 | <.0001 | <.0001 |
| | CTN-067 M0 vs. CTN-067 M3 | 11 | 3363.820 | 890.522 | 11 | 3076.910 | 985.389 | 196.640 | 232.310 | 0.4078 | 0.8317 |
| | CTN-067 M0 vs. CTN-067 M6 | 11 | 3363.820 | 890.522 | 11 | 3272.090 | 1021.100 | -3.027 | 235.130 | 0.9899 | 1.0000 |
| | CTN-067 M3 vs. CTN-067 M6 | 11 | 3076.910 | 985.389 | 11 | 3272.090 | 1021.100 | -199.670 | 202.640 | 0.3368 | 0.7594 |
| CD19+/CCR7+ \| Freq. of Parent | Control vs. CTN-067 M0 | 8 | 78.750 | 8.439 | 11 | 60.700 | 11.819 | 18.352 | 6.718 | 0.0133 | 0.0587 |
| | Control vs. CTN-067 M3 | 8 | 78.750 | 8.439 | 11 | 60.191 | 12.985 | 20.239 | 6.042 | 0.0034 | 0.0163 |
| | Control vs. CTN-067 M6 | 8 | 78.750 | 8.439 | 11 | 65.391 | 13.441 | 15.108 | 6.017 | 0.0212 | 0.0902 |
| | CTN-067 M0 vs. CTN-067 M3 | 11 | 60.700 | 11.819 | 11 | 60.191 | 12.985 | 1.888 | 3.187 | 0.5607 | 0.9332 |
| | CTN-067 M0 vs. CTN-067 M6 | 11 | 60.700 | 11.819 | 11 | 65.391 | 13.441 | -3.244 | 3.225 | 0.3271 | 0.7479 |
| | CTN-067 M3 vs. CTN-067 M6 | 11 | 60.191 | 12.985 | 11 | 65.391 | 13.441 | -5.132 | 2.794 | 0.0819 | 0.2876 |

| | | | | | | | | | | | |
|---|---|---|---|---|---|---|---|---|---|---|---|
| CD19+ \| Geometric Mean (BV605-A :: CD45RA) | Control vs. CTN-067 M0 | 8 | 194548.380 | 25568.370 | 11 | 214023.270 | 44072.400 | -677.120 | 21825.000 | 0.9756 | 1.0000 |
| | Control vs. CTN-067 M3 | 8 | 194548.380 | 25568.370 | 11 | 211722.550 | 50486.230 | 1052.460 | 19963.000 | 0.9585 | 0.9999 |
| | Control vs. CTN-067 M6 | 8 | 194548.380 | 25568.370 | 11 | 216581.730 | 46121.290 | -3835.120 | 19894.000 | 0.8492 | 0.9974 |
| | CTN-067 M0 vs. CTN-067 M3 | 11 | 214023.270 | 44072.400 | 11 | 211722.550 | 50486.230 | 1729.580 | 9481.240 | 0.8572 | 0.9978 |
| | CTN-067 M0 vs. CTN-067 M6 | 11 | 214023.270 | 44072.400 | 11 | 216581.730 | 46121.290 | -3158.000 | 9595.060 | 0.7457 | 0.9873 |
| | CTN-067 M3 vs. CTN-067 M6 | 11 | 211722.550 | 50486.230 | 11 | 216581.730 | 46121.290 | -4887.580 | 8284.910 | 0.5622 | 0.9339 |
| CD19+/CD45RA+ \| Freq. of Parent | Control vs. CTN-067 M0 | 8 | 99.563 | 0.302 | 11 | 99.255 | 0.425 | 0.400 | 0.256 | 0.1348 | 0.4227 |
| | Control vs. CTN-067 M3 | 8 | 99.563 | 0.302 | 11 | 99.282 | 0.354 | 0.337 | 0.215 | 0.1339 | 0.4206 |
| | Control vs. CTN-067 M6 | 8 | 99.563 | 0.302 | 11 | 99.336 | 0.420 | 0.281 | 0.214 | 0.2045 | 0.5657 |
| | CTN-067 M0 vs. CTN-067 M3 | 11 | 99.255 | 0.425 | 11 | 99.282 | 0.354 | -0.062 | 0.165 | 0.7088 | 0.9809 |
| | CTN-067 M0 vs. CTN-067 M6 | 11 | 99.255 | 0.425 | 11 | 99.336 | 0.420 | -0.119 | 0.166 | 0.4838 | 0.8903 |
| | CTN-067 M3 vs. CTN-067 M6 | 11 | 99.282 | 0.354 | 11 | 99.336 | 0.420 | -0.056 | 0.148 | 0.7077 | 0.9807 |
| CD19+ \| Geometric Mean (BV650-A :: CD15) | Control vs. CTN-067 M0 | 8 | -47.425 | 371.075 | 11 | 15.073 | 508.643 | -78.548 | 269.690 | 0.7740 | 0.9911 |
| | Control vs. CTN-067 M3 | 8 | -47.425 | 371.075 | 11 | 206.491 | 444.957 | -237.040 | 244.170 | 0.3439 | 0.7674 |
| | Control vs. CTN-067 M6 | 8 | -47.425 | 371.075 | 11 | 178.785 | 462.483 | -207.700 | 243.220 | 0.4038 | 0.8280 |
| | CTN-067 M0 vs. CTN-067 M3 | 11 | 15.073 | 508.643 | 11 | 206.491 | 444.957 | -158.490 | 123.750 | 0.2157 | 0.5856 |
| | CTN-067 M0 vs. CTN-067 M6 | 11 | 15.073 | 508.643 | 11 | 178.785 | 462.483 | -129.150 | 125.220 | 0.3153 | 0.7335 |
| | CTN-067 M3 vs. | 11 | 206.491 | 444.957 | 11 | 178.785 | 462.483 | 29.343 | 108.320 | 0.7894 | 0.9928 |

| | | | | | | | | | | | |
|---|---|---|---|---|---|---|---|---|---|---|---|
| | CTN-067 M6 | | | | | | | | | | |
| CD19+/CD15+ \| Freq. of Parent | Control vs. CTN-067 M0 | 8 | 20.763 | 7.224 | 11 | 23.364 | 9.499 | -0.651 | 5.423 | 0.9057 | 0.9994 |
| | Control vs. CTN-067 M3 | 8 | 20.763 | 7.224 | 11 | 26.682 | 8.074 | -4.100 | 4.708 | 0.3947 | 0.8197 |
| | Control vs. CTN-067 M6 | 8 | 20.763 | 7.224 | 11 | 26.618 | 9.392 | -4.043 | 4.681 | 0.3985 | 0.8232 |
| | CTN-067 M0 vs. CTN-067 M3 | 11 | 23.364 | 9.499 | 11 | 26.682 | 8.074 | -3.449 | 3.010 | 0.2661 | 0.6667 |
| | CTN-067 M0 vs. CTN-067 M6 | 11 | 23.364 | 9.499 | 11 | 26.618 | 9.392 | -3.392 | 3.044 | 0.2789 | 0.6853 |
| | CTN-067 M3 vs. CTN-067 M6 | 11 | 26.682 | 8.074 | 11 | 26.618 | 9.392 | 0.057 | 2.662 | 0.9831 | 1.0000 |
| CD19+ \| Geometric Mean (BV711-A :: CD86) | Control vs. CTN-067 M0 | 8 | 653.875 | 185.189 | 11 | 895.909 | 236.316 | -104.650 | 154.190 | 0.5055 | 0.9039 |
| | Control vs. CTN-067 M3 | 8 | 653.875 | 185.189 | 11 | 951.909 | 302.925 | -246.020 | 138.940 | 0.0926 | 0.3172 |
| | Control vs. CTN-067 M6 | 8 | 653.875 | 185.189 | 11 | 943.818 | 338.557 | -242.170 | 138.360 | 0.0962 | 0.3268 |
| | CTN-067 M0 vs. CTN-067 M3 | 11 | 895.909 | 236.316 | 11 | 951.909 | 302.925 | -141.370 | 72.484 | 0.0660 | 0.2413 |
| | CTN-067 M0 vs. CTN-067 M6 | 11 | 895.909 | 236.316 | 11 | 943.818 | 338.557 | -137.520 | 73.340 | 0.0762 | 0.2713 |
| | CTN-067 M3 vs. CTN-067 M6 | 11 | 951.909 | 302.925 | 11 | 943.818 | 338.557 | 3.847 | 63.502 | 0.9523 | 0.9999 |
| CD19+/CD86+ \| Freq. of Parent | Control vs. CTN-067 M0 | 8 | 3.884 | 1.623 | 11 | 7.580 | 4.283 | -2.715 | 1.896 | 0.1685 | 0.4960 |
| | Control vs. CTN-067 M3 | 8 | 3.884 | 1.623 | 11 | 6.927 | 3.268 | -2.708 | 1.698 | 0.1271 | 0.4046 |
| | Control vs. CTN-067 M6 | 8 | 3.884 | 1.623 | 11 | 5.417 | 2.388 | -1.231 | 1.690 | 0.4755 | 0.8847 |
| | CTN-067 M0 vs. CTN-067 M3 | 11 | 7.580 | 4.283 | 11 | 6.927 | 3.268 | 0.006 | 0.919 | 0.9946 | 1.0000 |
| | CTN-067 M0 vs. CTN-067 M6 | 11 | 7.580 | 4.283 | 11 | 5.417 | 2.388 | 1.484 | 0.930 | 0.1269 | 0.4041 |

| | | | | | | | | | | |
|---|---|---|---|---|---|---|---|---|---|---|
| | CTN-067 M3 vs. CTN-067 M6 | 11 | 6.927 | 3.268 | 11 | 5.417 | 2.388 | 1.478 | 0.806 | 0.0825 | 0.2891 |
| CD19+ \| Geometric Mean (PE-A :: CD38) | Control vs. CTN-067 M0 | 8 | 18559.000 | 5640.560 | 11 | 15505.450 | 12156.490 | 4939.030 | 4858.890 | 0.3222 | 0.7420 |
| | Control vs. CTN-067 M3 | 8 | 18559.000 | 5640.560 | 11 | 11283.910 | 4911.550 | 7588.110 | 4228.940 | 0.0887 | 0.3064 |
| | Control vs. CTN-067 M6 | 8 | 18559.000 | 5640.560 | 11 | 11814.270 | 6377.310 | 6979.570 | 4204.890 | 0.1134 | 0.3710 |
| | CTN-067 M0 vs. CTN-067 M3 | 11 | 15505.450 | 12156.490 | 11 | 11283.910 | 4911.550 | 2649.080 | 2668.680 | 0.3334 | 0.7553 |
| | CTN-067 M0 vs. CTN-067 M6 | 11 | 15505.450 | 12156.490 | 11 | 11814.270 | 6377.310 | 2040.530 | 2698.440 | 0.4588 | 0.8729 |
| | CTN-067 M3 vs. CTN-067 M6 | 11 | 11283.910 | 4911.550 | 11 | 11814.270 | 6377.310 | -608.540 | 2357.650 | 0.7991 | 0.9938 |
| CD19+/CD38+ \| Freq. of Parent | Control vs. CTN-067 M0 | 8 | 88.613 | 3.564 | 11 | 82.273 | 7.726 | 5.971 | 5.044 | 0.2511 | 0.6440 |
| | Control vs. CTN-067 M3 | 8 | 88.613 | 3.564 | 11 | 78.264 | 10.539 | 9.892 | 4.624 | 0.0456 | 0.1767 |
| | Control vs. CTN-067 M6 | 8 | 88.613 | 3.564 | 11 | 78.627 | 11.238 | 9.524 | 4.609 | 0.0527 | 0.1997 |
| | CTN-067 M0 vs. CTN-067 M3 | 11 | 82.273 | 7.726 | 11 | 78.264 | 10.539 | 3.921 | 2.164 | 0.0858 | 0.2986 |
| | CTN-067 M0 vs. CTN-067 M6 | 11 | 82.273 | 7.726 | 11 | 78.627 | 11.238 | 3.553 | 2.190 | 0.1212 | 0.3904 |
| | CTN-067 M3 vs. CTN-067 M6 | 11 | 78.264 | 10.539 | 11 | 78.627 | 11.238 | -0.368 | 1.890 | 0.8477 | 0.9973 |
| CD19+ \| Geometric Mean (R718-A :: CCR5) | Control vs. CTN-067 M0 | 8 | 180.350 | 77.104 | 11 | 258.455 | 148.143 | 81.296 | 79.115 | 0.3171 | 0.7357 |
| | Control vs. CTN-067 M3 | 8 | 180.350 | 77.104 | 11 | 223.964 | 168.795 | 37.938 | 70.210 | 0.5952 | 0.9480 |
| | Control vs. CTN-067 M6 | 8 | 180.350 | 77.104 | 11 | 228.836 | 196.833 | 29.195 | 69.874 | 0.6808 | 0.9747 |
| | CTN-067 M0 vs. CTN-067 M3 | 11 | 258.455 | 148.143 | 11 | 223.964 | 168.795 | -43.359 | 39.941 | 0.2913 | 0.7023 |
| | CTN-067 M0 vs. | 11 | 258.455 | 148.143 | 11 | 228.836 | 196.833 | -52.102 | 40.403 | 0.2127 | 0.5803 |

| | | | | | | | | | | |
|---|---|---|---|---|---|---|---|---|---|---|
| | CTN-067 M6 | | | | | | | | | |
| | CTN-067 M3 vs. CTN-067 M6 | 11 | 223.964 | 168.795 | 11 | 228.836 | 196.833 | -8.743 | 35.103 | 0.8060 | 0.9944 |
| CD19+/CCR5+ \| Freq. of Parent | Control vs. CTN-067 M0 | 8 | 2.095 | 1.275 | 11 | 3.744 | 1.772 | -0.103 | 1.062 | 0.9236 | 0.9997 |
| | Control vs. CTN-067 M3 | 8 | 2.095 | 1.275 | 11 | 3.011 | 2.174 | -0.199 | 0.968 | 0.8394 | 0.9968 |
| | Control vs. CTN-067 M6 | 8 | 2.095 | 1.275 | 11 | 3.046 | 2.659 | -0.276 | 0.965 | 0.7783 | 0.9916 |
| | CTN-067 M0 vs. CTN-067 M3 | 11 | 3.744 | 1.772 | 11 | 3.011 | 2.174 | -0.096 | 0.471 | 0.8413 | 0.9969 |
| | CTN-067 M0 vs. CTN-067 M6 | 11 | 3.744 | 1.772 | 11 | 3.046 | 2.659 | -0.172 | 0.476 | 0.7217 | 0.9833 |
| | CTN-067 M3 vs. CTN-067 M6 | 11 | 3.011 | 2.174 | 11 | 3.046 | 2.659 | -0.077 | 0.412 | 0.8542 | 0.9976 |
| CD19+ \| Geometric Mean (RB545-A :: CD57) | Control vs. CTN-067 M0 | 8 | 1148.380 | 328.966 | 11 | 1061.640 | 242.937 | 545.030 | 243.700 | 0.0375 | 0.1492 |
| | Control vs. CTN-067 M3 | 8 | 1148.380 | 328.966 | 11 | 1111.550 | 454.586 | 291.610 | 203.630 | 0.1684 | 0.4958 |
| | Control vs. CTN-067 M6 | 8 | 1148.380 | 328.966 | 11 | 1140.090 | 528.976 | 252.950 | 202.060 | 0.2258 | 0.6030 |
| | CTN-067 M0 vs. CTN-067 M3 | 11 | 1061.640 | 242.937 | 11 | 1111.550 | 454.586 | -253.410 | 169.740 | 0.1519 | 0.4609 |
| | CTN-067 M0 vs. CTN-067 M6 | 11 | 1061.640 | 242.937 | 11 | 1140.090 | 528.976 | -292.080 | 171.210 | 0.1043 | 0.3480 |
| | CTN-067 M3 vs. CTN-067 M6 | 11 | 1111.550 | 454.586 | 11 | 1140.090 | 528.976 | -38.663 | 154.650 | 0.8053 | 0.9943 |
| CD19+ \| Geometric Mean (RB705-A :: CCR2) | Control vs. CTN-067 M0 | 8 | 18.066 | 6.577 | 11 | 10.779 | 6.889 | 15.883 | 5.738 | 0.0122 | 0.0546 |
| | Control vs. CTN-067 M3 | 8 | 18.066 | 6.577 | 11 | 17.715 | 14.492 | 5.348 | 4.928 | 0.2914 | 0.7025 |
| | Control vs. CTN-067 M6 | 8 | 18.066 | 6.577 | 11 | 13.406 | 6.104 | 9.477 | 4.897 | 0.0680 | 0.2471 |
| | CTN-067 M0 vs. CTN-067 M3 | 11 | 10.779 | 6.889 | 11 | 17.715 | 14.492 | -10.535 | 3.335 | 0.0052 | 0.0244 |

| | | | | | | | | | | | |
|---|---|---|---|---|---|---|---|---|---|---|---|
| | CTN-067 M0 vs. CTN-067 M6 | 1 1 | 10.779 | 6.889 | 1 1 | 13.406 | 6.104 | -6.406 | 3.371 | 0.07 27 | 0.26 11 |
| | CTN-067 M3 vs. CTN-067 M6 | 1 1 | 17.715 | 14.492 | 1 1 | 13.406 | 6.104 | 4.129 | 2.961 | 0.17 92 | 0.51 77 |
| CD19+/CCR2 + \| Freq. of Parent | Control vs. CTN-067 M0 | 8 | 534.000 | 158.139 | 1 1 | 618.827 | 311.012 | 193.630 | 139.240 | 0.18 04 | 0.52 00 |
| | Control vs. CTN-067 M3 | 8 | 534.000 | 158.139 | 1 1 | 590.636 | 269.535 | 148.000 | 122.330 | 0.24 12 | 0.62 83 |
| | Control vs. CTN-067 M6 | 8 | 534.000 | 158.139 | 1 1 | 633.000 | 309.662 | 101.970 | 121.690 | 0.41 25 | 0.83 58 |
| | CTN-067 M0 vs. CTN-067 M3 | 1 1 | 618.827 | 311.012 | 1 1 | 590.636 | 269.535 | -45.631 | 73.463 | 0.54 19 | 0.92 40 |
| | CTN-067 M0 vs. CTN-067 M6 | 1 1 | 618.827 | 311.012 | 1 1 | 633.000 | 309.662 | -91.665 | 74.298 | 0.23 23 | 0.61 39 |
| | CTN-067 M3 vs. CTN-067 M6 | 1 1 | 590.636 | 269.535 | 1 1 | 633.000 | 309.662 | -46.034 | 64.724 | 0.48 56 | 0.89 14 |
| CD19+ \| Geometric Mean (RB780-A :: HLA-DR) | Control vs. CTN-067 M0 | 8 | 6.001 | 1.783 | 1 1 | 7.490 | 5.344 | 2.682 | 1.782 | 0.14 87 | 0.45 40 |
| | Control vs. CTN-067 M3 | 8 | 6.001 | 1.783 | 1 1 | 6.206 | 4.355 | 3.354 | 1.611 | 0.05 11 | 0.19 46 |
| | Control vs. CTN-067 M6 | 8 | 6.001 | 1.783 | 1 1 | 7.451 | 5.647 | 2.079 | 1.605 | 0.21 05 | 0.57 65 |
| | CTN-067 M0 vs. CTN-067 M3 | 1 1 | 7.490 | 5.344 | 1 1 | 6.206 | 4.355 | 0.672 | 0.824 | 0.42 49 | 0.84 64 |
| | CTN-067 M0 vs. CTN-067 M6 | 1 1 | 7.490 | 5.344 | 1 1 | 7.451 | 5.647 | -0.603 | 0.834 | 0.47 85 | 0.88 67 |
| | CTN-067 M3 vs. CTN-067 M6 | 1 1 | 6.206 | 4.355 | 1 1 | 7.451 | 5.647 | -1.275 | 0.722 | 0.09 33 | 0.31 90 |
| CD19+/HLA-DR+ \| Freq. of Parent | Control vs. CTN-067 M0 | 8 | 349387.0 00 | 109833.7 80 | 1 1 | 368127.4 50 | 94712.9 00 | 12586.0 00 | 58004.0 00 | 0.83 05 | 0.99 63 |
| | Control vs. CTN-067 M3 | 8 | 349387.0 00 | 109833.7 80 | 1 1 | 331172.1 80 | 75922.1 50 | 46881.0 00 | 49767.0 00 | 0.35 80 | 0.78 29 |
| | Control vs. CTN-067 M6 | 8 | 349387.0 00 | 109833.7 80 | 1 1 | 323215.1 80 | 89287.3 90 | 54706.0 00 | 49450.0 00 | 0.28 24 | 0.69 01 |
| | CTN-067 M0 vs. | 1 1 | 368127.4 50 | 94712.90 0 | 1 1 | 331172.1 80 | 75922.1 50 | 34295.0 00 | 33877.0 00 | 0.32 41 | 0.74 43 |

| | | | | | | | | | | |
|---|---|---|---|---|---|---|---|---|---|---|
| | CTN-067 M3 | | | | | | | | | |
| | CTN-067 M0 vs. CTN-067 M6 | 11 | 368127.450 | 94712.900 | 11 | 323215.180 | 89287.390 | 42120.000 | 34240.000 | 0.2337 | 0.6161 |
| | CTN-067 M3 vs. CTN-067 M6 | 11 | 331172.180 | 75922.150 | 11 | 323215.180 | 89287.390 | 7824.760 | 30086.000 | 0.7976 | 0.9936 |
| CD19+ \| Geometric Mean (RY586-A :: CXCR4) | Control vs. CTN-067 M0 | 8 | 97.988 | 1.362 | 11 | 97.555 | 1.086 | 0.763 | 0.868 | 0.3902 | 0.8154 |
| | Control vs. CTN-067 M3 | 8 | 97.988 | 1.362 | 11 | 96.673 | 1.823 | 1.505 | 0.793 | 0.0730 | 0.2619 |
| | Control vs. CTN-067 M6 | 8 | 97.988 | 1.362 | 11 | 97.555 | 1.445 | 0.616 | 0.790 | 0.4452 | 0.8627 |
| | CTN-067 M0 vs. CTN-067 M3 | 11 | 97.555 | 1.086 | 11 | 96.673 | 1.823 | 0.741 | 0.380 | 0.0662 | 0.2419 |
| | CTN-067 M0 vs. CTN-067 M6 | 11 | 97.555 | 1.086 | 11 | 97.555 | 1.445 | -0.148 | 0.385 | 0.7058 | 0.9803 |
| | CTN-067 M3 vs. CTN-067 M6 | 11 | 96.673 | 1.823 | 11 | 97.555 | 1.445 | -0.889 | 0.333 | 0.0150 | 0.0659 |
| CD19+/CXCR4+ \| Freq. of Parent | Control vs. CTN-067 M0 | 8 | 22232.250 | 8473.590 | 11 | 55362.090 | 23488.870 | -17163.000 | 20242.000 | 0.4070 | 0.8310 |
| | Control vs. CTN-067 M3 | 8 | 22232.250 | 8473.590 | 11 | 31035.910 | 36792.850 | 3574.090 | 17078.000 | 0.8365 | 0.9966 |
| | Control vs. CTN-067 M6 | 8 | 22232.250 | 8473.590 | 11 | 25859.000 | 42986.100 | 8572.550 | 16955.000 | 0.6189 | 0.9568 |
| | CTN-067 M0 vs. CTN-067 M3 | 11 | 55362.090 | 23488.870 | 11 | 31035.910 | 36792.850 | 20737.000 | 12840.000 | 0.1228 | 0.3942 |
| | CTN-067 M0 vs. CTN-067 M6 | 11 | 55362.090 | 23488.870 | 11 | 25859.000 | 42986.100 | 25735.000 | 12968.000 | 0.0618 | 0.2284 |
| | CTN-067 M3 vs. CTN-067 M6 | 11 | 31035.910 | 36792.850 | 11 | 25859.000 | 42986.100 | 4998.460 | 11516.000 | 0.6692 | 0.9719 |
| CD19+ \| Geometric Mean (RY610-A :: CD69) | Control vs. CTN-067 M0 | 8 | 90.875 | 3.220 | 11 | 92.073 | 8.026 | -1.206 | 6.962 | 0.8643 | 0.9981 |
| | Control vs. CTN-067 M3 | 8 | 90.875 | 3.220 | 11 | 79.727 | 14.922 | 12.122 | 5.931 | 0.0551 | 0.2073 |
| | Control vs. CTN-067 M6 | 8 | 90.875 | 3.220 | 11 | 81.991 | 12.869 | 9.908 | 5.891 | 0.1090 | 0.3599 |

| | | | | | | | | | | |
|---|---|---|---|---|---|---|---|---|---|---|
| | CTN-067 M0 vs. CTN-067 M3 | 11 | 92.073 | 8.026 | 11 | 79.727 | 14.922 | 13.328 | 4.200 | 0.0050 | 0.0237 |
| | CTN-067 M0 vs. CTN-067 M6 | 11 | 92.073 | 8.026 | 11 | 81.991 | 12.869 | 11.113 | 4.244 | 0.0169 | 0.0733 |
| | CTN-067 M3 vs. CTN-067 M6 | 11 | 79.727 | 14.922 | 11 | 81.991 | 12.869 | -2.215 | 3.743 | 0.5610 | 0.9333 |
| CD19+/CD69+ \| Freq. of Parent | Control vs. CTN-067 M0 | 8 | 819.750 | 337.588 | 11 | 1308.730 | 651.825 | -188.130 | 339.020 | 0.5854 | 0.9441 |
| | Control vs. CTN-067 M3 | 8 | 819.750 | 337.588 | 11 | 1078.910 | 571.553 | -64.630 | 291.720 | 0.8270 | 0.9960 |
| | Control vs. CTN-067 M6 | 8 | 819.750 | 337.588 | 11 | 1085.730 | 495.251 | -76.734 | 289.900 | 0.7941 | 0.9933 |
| | CTN-067 M0 vs. CTN-067 M3 | 11 | 1308.730 | 651.825 | 11 | 1078.910 | 571.553 | 123.500 | 195.490 | 0.5351 | 0.9205 |
| | CTN-067 M0 vs. CTN-067 M6 | 11 | 1308.730 | 651.825 | 11 | 1085.730 | 495.251 | 111.390 | 197.610 | 0.5795 | 0.9416 |
| | CTN-067 M3 vs. CTN-067 M6 | 11 | 1078.910 | 571.553 | 11 | 1085.730 | 495.251 | -12.104 | 173.410 | 0.9451 | 0.9999 |
| CD19+ \| Geometric Mean (RY703-A :: PD-1) | Control vs. CTN-067 M0 | 8 | 26.013 | 6.906 | 11 | 35.255 | 9.972 | -5.319 | 5.476 | 0.3437 | 0.7671 |
| | Control vs. CTN-067 M3 | 8 | 26.013 | 6.906 | 11 | 28.745 | 9.648 | -0.099 | 4.587 | 0.9829 | 1.0000 |
| | Control vs. CTN-067 M6 | 8 | 26.013 | 6.906 | 11 | 28.182 | 4.940 | 0.400 | 4.552 | 0.9309 | 0.9997 |
| | CTN-067 M0 vs. CTN-067 M3 | 11 | 35.255 | 9.972 | 11 | 28.745 | 9.648 | 5.219 | 3.668 | 0.1710 | 0.5012 |
| | CTN-067 M0 vs. CTN-067 M6 | 11 | 35.255 | 9.972 | 11 | 28.182 | 4.940 | 5.719 | 3.702 | 0.1389 | 0.4320 |
| | CTN-067 M3 vs. CTN-067 M6 | 11 | 28.745 | 9.648 | 11 | 28.182 | 4.940 | 0.500 | 3.318 | 0.8819 | 0.9987 |
| CD19+/PD-1+ \| Freq. of Parent | Control vs. CTN-067 M0 | 8 | 419.500 | 80.637 | 11 | 528.818 | 199.385 | 22.118 | 3845.200 | 0.9955 | 1.0000 |
| | Control vs. CTN-067 M3 | 8 | 419.500 | 80.637 | 11 | 603.091 | 217.740 | -90.239 | 3845.110 | 0.9815 | 1.0000 |

| | | | | | | | | | | | |
|---|---|---|---|---|---|---|---|---|---|---|---|
| | Control vs. CTN-067 M6 | 8 | 419.500 | 80.637 | 11 | 591.455 | 257.646 | -80.496 | 3845.110 | 0.9835 | 1.0000 |
| | CTN-067 M0 vs. CTN-067 M3 | 11 | 528.818 | 199.385 | 11 | 603.091 | 217.740 | -112.360 | 26.754 | 0.0005 | 0.0025 |
| | CTN-067 M0 vs. CTN-067 M6 | 11 | 528.818 | 199.385 | 11 | 591.455 | 257.646 | -102.610 | 27.095 | 0.0012 | 0.0063 |
| | CTN-067 M3 vs. CTN-067 M6 | 11 | 603.091 | 217.740 | 11 | 591.455 | 257.646 | 9.743 | 23.152 | 0.6786 | 0.9742 |
